# Comparative analysis of clonal evolution among patients with right-sided colon cancer, left-sided colon cancer and rectal cancer

**DOI:** 10.1101/2020.07.01.181586

**Authors:** Santasree Banerjee, Xianxiang Zhang, Shan Kuang, Jigang Wang, Lei Li, Guangyi Fan, Yonglun Luo, Shuai Sun, Peng Han, Qingyao Wu, Shujian Yang, Xiaobin Ji, Yong Li, Li Deng, Xiaofen Tian, Zhiwei Wang, Yue Zhang, Kui Wu, Shida Zhu, Lars Bolund, Huanming Yang, Xun Xu, Junnian Liu, Yun Lu, Xin Liu

**Author notes:** Correspondence Xin Liu, BGI-Qingdao, BGI-Shenzhen, Qingdao, 266555, China., Yun Lu, Department of Gastroenterology, General Surgery Center, The Affiliated Hospital of Qingdao University, Qingdao, 266555, China., Junnian Liu, BGI-Qingdao, BGI-Shenzhen, Qingdao, 266555, China. These authors contributed equally to this study (Santasree Banerjee, Xianxiang Zhang, Shan Kuang, Jigang Wang, Lei Li).

## Abstract

**Background:** Tumor multi-region sequencing reveals intratumor heterogeneity (ITH) and clonal evolution which play a key role in progression and metastases of the tumor. However, large-scale high depths multiregional sequencing of colorectal cancer (CRC) has not been well studied. In addition, the comparative analysis among right-sided colon cancer (RCC), left-sided colon cancer (LCC) and rectal cancer (RC) patients as well as the study of lymph node metastasis (LN) with extranodal tumor deposits (ENTD) from evolutionary perspective remain unknown.

**Results:** In this prospective study, we recruited different stages of 68 CRC patients with RCC (18), LCC (20) and RC (30). We performed high-depth whole exome sequencing (WES) of 206 tumor regions including 176 primary tumors, 19 LN and 11 ENTD samples. Our results showed ITH with a Darwinian pattern of evolution. We identified that the evolution pattern of LCC and RC was more complex and divergent than RCC, suggesting the evolutionary diversity in the initiation and progression of LCC and RC. Genetic and evolutionary evidences found that both LN and ENTD were of polyclonal in origin. Moreover, ENTD was a distinct entity from LN and evolved later.

**Conclusions:** In conclusion, our study showed the Darwinian pattern of evolution with differences in clonal evolution between RCC with LCC and RC.

## Background

CRC is the third most common malignancy and the second leading cause of cancer death worldwide [1]. According to the World Health Organization (WHO) GLOBOCAN database, there were 1,849,518 estimated new CRC cases and 880,792 CRC-related deaths in 2018 [2]. In China, CRC is the second most common neoplasia, occupying the fifth position in mortality, accounting for an incidence of 521,490 new cases and 248,400 deaths in 2018 [2].

Tumor multi-region sequencing reveals ITH and clonal evolution which play a key role in progression and metastases of the tumor [3]. The development of effective target-based precision medicine and personalized cancer therapy is based on ITH and the pattern of clonal evolution in colorectal tumors [4]. Therefore, patients with CRC may respond variably to the same treatment, due to ITH and differences in clonal evolution, despite there being no significant differences identified in the tumor histopathology [5]. Hence, study of ITH and comparative analysis of clonal evolution is highly significant from both clinical and biological perspective, to understand the genomic changes driving the malignant process, which is fundamental to developing an effective personalized cancer therapy.

Recently, tumor multi-region sequencing studies of colorectal cancer have demonstrated ITH [6-13]. This multiregional sequencing approach, sequencing DNA samples from geographically separated regions of a single tumor, explores ITH and cancer evolution. Large-scale multiregional sequencing studies have systematically revealed ITH as well as cancer evolution in patients with non-small-cell lung cancer and renal cancer [14-16]. However, large-scale multiregional sequencing studies of CRC have not been well reported. In addition, multiregional sequencing studies in CRC were performed at relatively shallow sequencing depths [6-10], making it difficult to assess ITH, due to inability to detect somatic mutations with low frequencies.

CRC is no longer regarded as a single disease with increasing knowledge of the molecular mechanisms of carcinogenesis. The location of the primary tumor, with respect to the right side or left side of the splenic flexure and rectum, is an important prognostic factor of CRC [17, 18]. LCC and RC patients (originating from splenic flexure, descending colon, sigmoid colon and rectum) survive longer than RCC patients (originating from hepatic flexure, ascending colon and cecum). Clinical symptoms are also different between patients with RCC and LCC/RC [19, 20]. RCC patients tend to be older, female and have advanced stage of tumors with frequent metastasis to peritoneum compared to metastasis to lung and liver in LCC/RC patients. In addition, RCC and LCC/RC patients exhibits different treatment outcomes towards anti-epidermal growth factor receptor (EGFR) therapy [20]. Many studies have been done to explore the possible reasons for clinical heterogeneity between RCC and LCC/RC and found differences in their embryonic origin, blood supplies, genetic mutations, genomic expression profiles, immunological composition and bacterial population in tumor microenvironment [19-23]. However, the understanding of the ITH and clonal evolution that determine the pathogenesis of RCC and LCC/RC is still unclear.

Amongst CRC patients, the stage of the disease is one of the most important prognostic factors which is correlated with the disease survival rate [24]. Tumor Node Metastasis (TNM)/American Joint Committee on Cancer (AJCC) Cancer Staging system is the gold standard for determining the correct cancer stage, helping to make appropriate treatment plans. Among CRC patients, the presence of cancer cells in lymph nodes is defined as stage III disease [25]. In the 7th and 8th editions of TNM staging system, a separate entity, entitled extranodal tumour deposits (ENTD), was included as ‘N1c’ subcategory [26]. However, inclusion of ENTD within nodal staging has worldwide debates in CRC because lack of significant improvement of prognostic value [27-29]. Although, many ITH and evolution studies of CRC focus on spreading routes of lymphatic metastases by sampling paired primary tumors and LN, none of them included ENTD samples [10-13]. Therefore, the molecular signature and evolutionary relationship between LN and ENTD has not been clear till now. Hence, the characterization of the molecular signature and evolution of the primary tumor, LN and ENTD is very significant for TNM staging and therapeutic interventions for the patients with CRC.

In order to overcome the drawbacks of previous studies, we have comprehensively studied the ITH and clonal evolution of CRC, using high depth (median depth of 395×) WES of 206 multi-region tumor samples and 68 matched germline samples from 68 CRC tumors, determined the differences of ITH, and the clonal evolution of CRC in RCC, LCC and RC patients.

## Results

Comprehensive clinical descriptions of these 68 CRC patients were provided in **Table S1**. Tumor multi-region high depth (median depth of 395×, range 179-596) WES was performed with 206 tumor regions (2-7 regions/tumor) including 176 primary tumor regions, 19 LN regions and 11 ENTD regions, as well as 68 matched germline samples from 68 CRC patients. WES identified 6 hypermutated (mutation rates of each tumor region were >10 mutations/1 Mb bases) CRC patients, of these four patients were identified with microsatellite instability (MSI). The remaining 62 CRC patients were microsatellite stable (MSS) and of these, 12 are RCC patients, 20 are LCC patients and 30 are RC patients. Hypermutated patients were analyzed separately.

### ITH in colorectal tumors

WES of 62 tumors with 188 tumor regions identified 19454 somatic mutations including 17560 SNVs (14361 non-silent SNVs) and 1894 INDELs (**Table S2**). The mutation rate identified by the multi-region WES was significantly higher than single sample sequencing due to detection of subclonal mutations (median number of mutations/1MB bases, 4.61 vs. 3.23; P=8.9×10^−9^) (**Figure S3**). In our study, the mutation rate of single sample sequencing was significantly higher than single CRC sample sequencing data from The Cancer Genome Atlas [30] (TCGA), probably due to the higher sequencing depth in our study (median number of mutations/1 MB bases, 3.23 vs. 2.07; P=1.7×10^−22^) (**Figure S3**).

Then, identified somatic mutations were divided into clonal and subclonal mutations (**Fig. 1A**). It is worth noting that 2 patients (CRC32 and CRC36) with LCC and 6 patients (CRC49, CRC42, CRC51, CRC48, CRC52 and CRC60) with RC had not identified with clonal mutations, suggesting that branched evolution was widespread in patients with LCC and RC. In addition, RCC Patients had significantly more clonal mutations than RC patients (median number, 160 vs 119; P=0.035) (**Figure S4**).

**Fig. 1.**
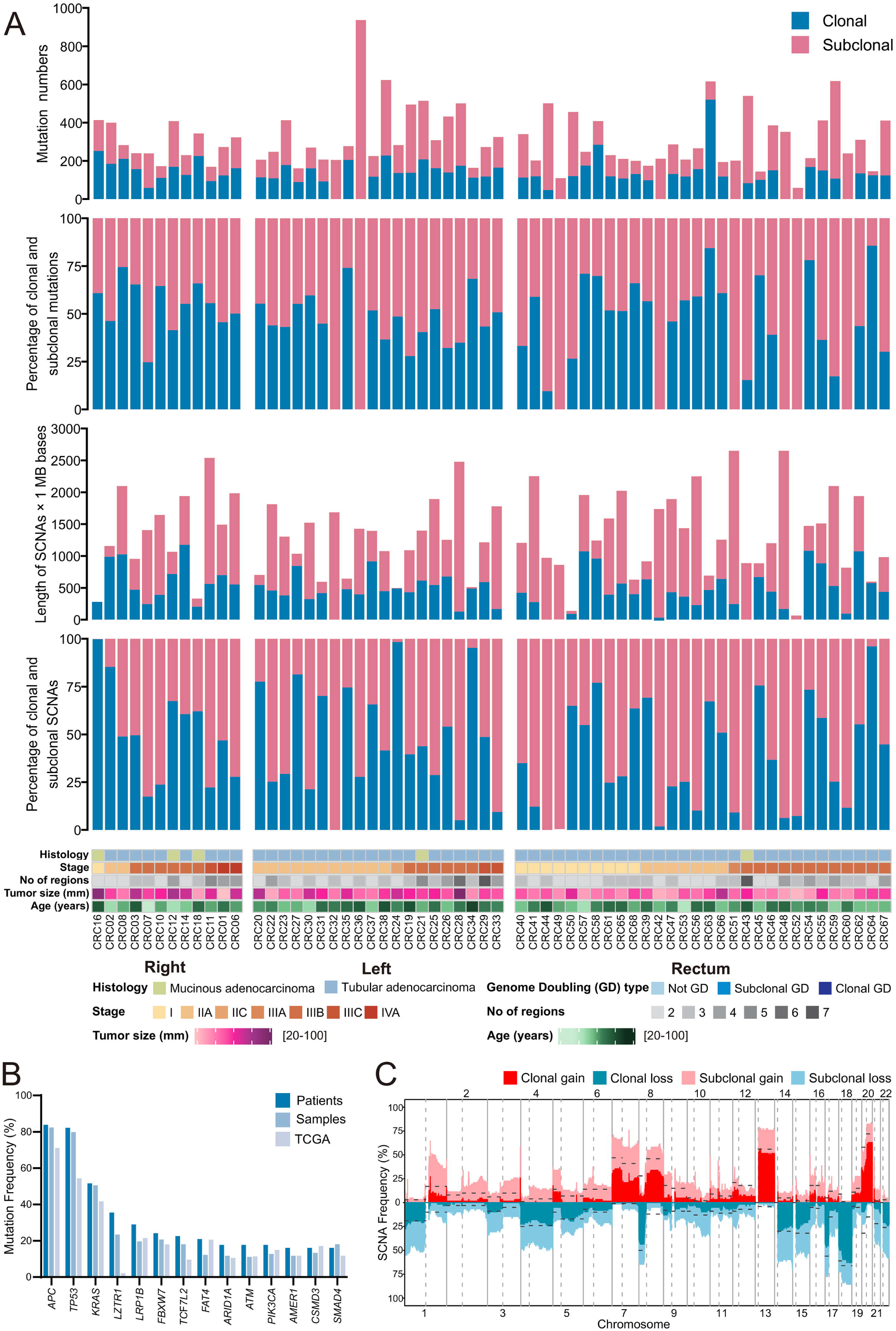
Overview of genomic heterogeneity in CRC tumors. **a** Heterogeneity of mutations and somatic copy-number alterations (SCNAs). Tumors were sorted by location and stage. (1) Number of all SNV and INDEL mutations (including coding and noncoding mutations) in CRC tumors. (2) The percentages of clonal mutations in CRC tumors. (3) Quantification of SCNAs in CRC tumors. (4) The percentages of clonal SCNAs in CRC tumors. (5) Demographic and clinical characteristics of the 62 CRC patients in this study (divided by histology; stage; number of regions; tumor size; age and tumor location). **b** Mutation frequency of driver genes (driver mutations occurred in not less than 10 patients) and comparison with TCGA data. **c** Frequency of SCNAs in CRC tumors. The dotted lines were frequency of SCNAs in TCGA CRC samples.

Somatic copy number alterations (SCNAs) were measured as length of segments affected by either gains or losses (detailed copy number data has been given in **Table S3**). We summarized the total length of the genome that subjected to SCNAs and calculated the percentage of clonal and subclonal SCNAs (**Fig. 1A**). Interestingly, in a RC patient (CRC43), all the identified SCNAs were subclonal. There were no significant differences in the length and percentage of SCNAs among the RCC, LCC and RC patients. (**Figure S5**).

In our study, we identified that the mutation frequency of 14 driver genes (*APC, TP53, KRAS, LZTR1, LRP1B, FBXW7, TCF7L2, FAT4, ARID1A, ATM, PIK3CA, AMER1, CSMD3* and *SMAD4*) were higher at patient-level than at sample-level, except *SMAD4* gene (**Fig. 1B**). In addition, we also found that the mutation frequency was higher at patient-level compared to the TCGA data [30] except *CSMD3* gene (**Fig. 1B**). Notably, the mutation frequency of the *LZTR1* gene was much higher than TCGA data [30] (**Fig. 1B**). Our study also identified higher frequency of SCNAs than TCGA [30] data, probably due to the identification of subclonal SCNAs (**Fig. 1C**).

### Clonal architecture in colorectal tumors

All the mutations (SNVs and INDELs) were clustered according to their CCF values to understand the clonal architecture and evolutionary history of 62 colorectal tumors. Each colored circle in the phylogenetic tree represented one cluster of the tumor (**Fig. 2**). Phylogenetic trees for 62 tumors and 188 regions together with schematic diagram of 100 tumor cells representing distribution of clusters in each tumor region were shown in **Figure S6**. Driver mutations, driver SCNAs and their clusters were annotated beside the phylogenetic trees (**Figure S6**). Detailed information of cluster numbers for each tumor was listed in **Table S4**, with a median of 6 clusters per tumor (range, 1 to 13). Our study showed that patients with LCC possessed significantly more cluster numbers than patients with both RCC (median number, 7.5 vs. 6; P=0.028) and RC (median number, 7.5 vs. 5.5; P=0.025) (**Figure S7**), which potentially reflected that LCC patients were structurally more complex than RCC patients in evolutionary perspective.

**Fig. 2.**
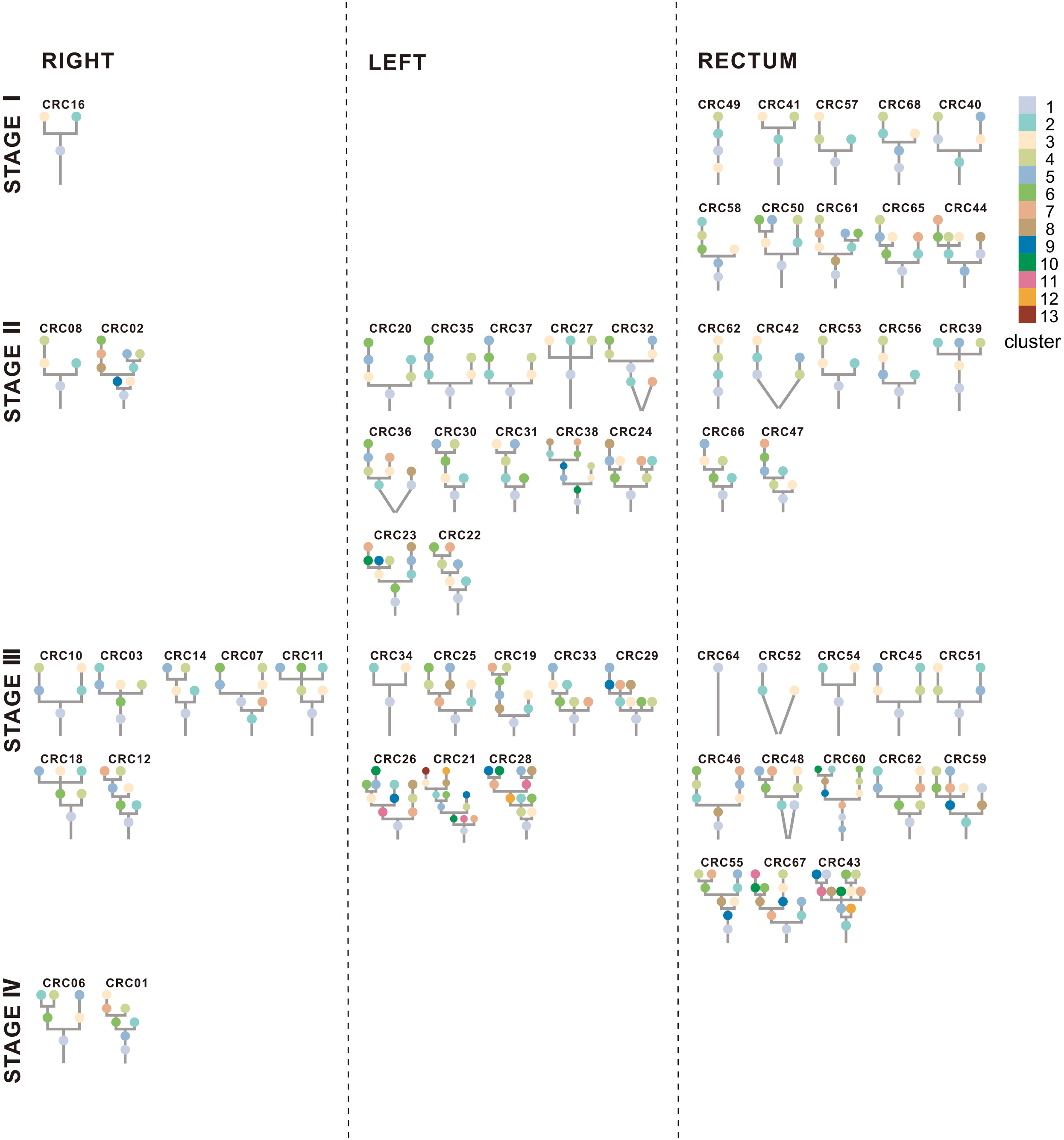
Phylogenetic trees. Phylogenetic trees for each CRC tumor were shown. The trees were ordered by overall stage (I, □, □, IV) and position (right-sided colon, left-sided colon and rectum). The cluster number corresponding to the color was displayed in the upper right corner with largest cluster labeled “1”. The lines connecting clusters does not contain any information.

### Driver event alterations in CRC evolution

Identifying cancer driver events and their clonality is highly significant to understand the driving force underlying the transformation of a benign tumor to a malignant one. Therefore, driver mutations, driver SCNAs, arm level SCNAs and their clonality were analyzed for colorectal tumors (**Fig. 3**).

**Fig. 3.**
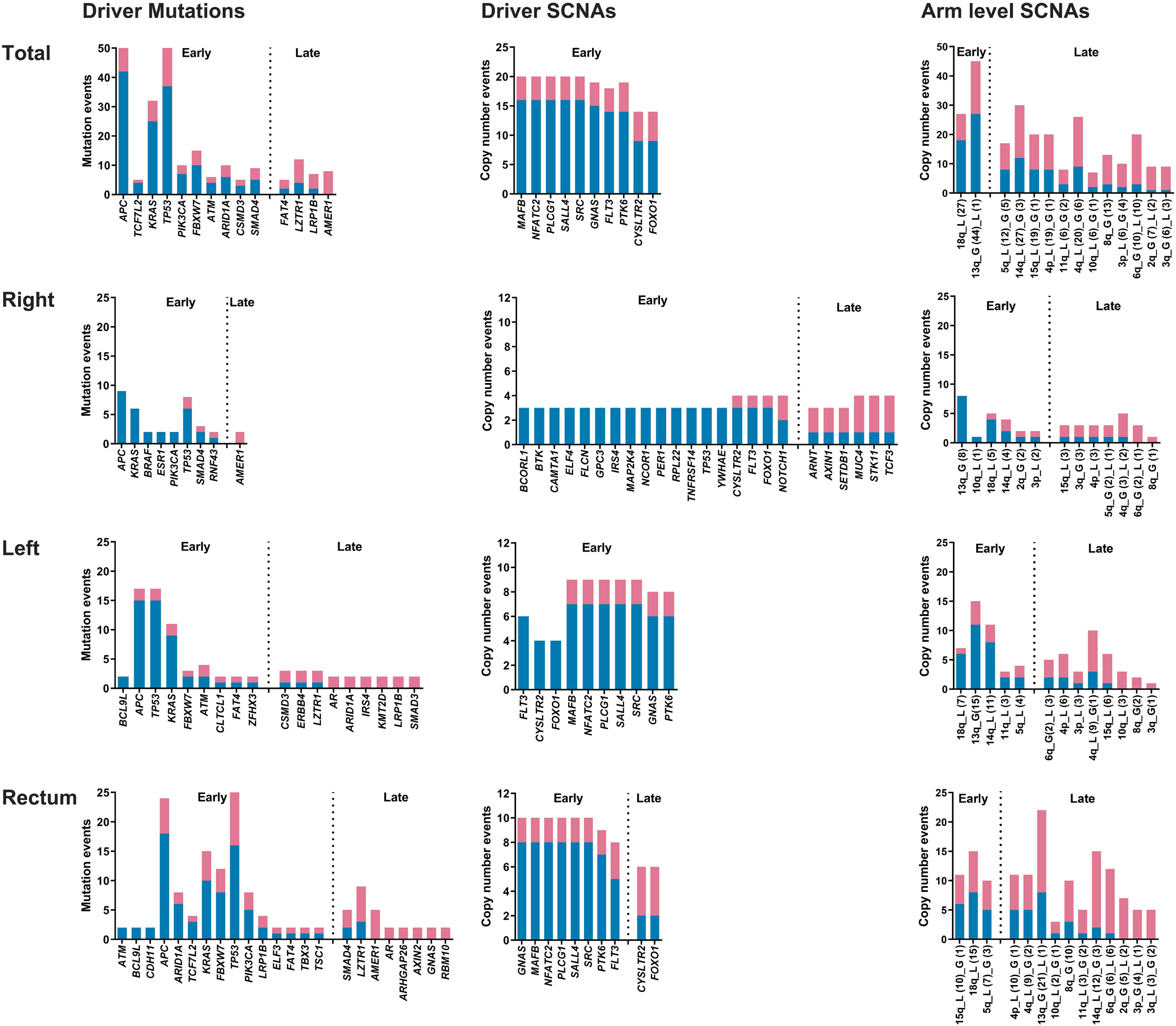
Summary of driver events in CRC evolution. Mutations and SCNAs were shown as occurrence in patients indicating whether the events are clonal (blue) or subclonal (red). Only genes that were mutated in at least five patients in total or two patients in right-sided colon/left-sided colon/rectum were shown. For SCNAs, driver SCNAs in at least 20% of the patients were shown while all the arm level SCNAs were shown. Driver events with more subclonal occurrence than clonal occurrence in tumors were late events, otherwise they were early events. In the arm level SCNAs part, “G” represented gain, “L” represented loss, and the numbers in parentheses represented the time of occurrence in tumors.

We identified 1373 driver events (405 driver mutations, 707 driver SCNAs and 261 arm level SCNAs) among 62 colorectal tumors. Among these events, 44% of driver events (605 out of 1373) were subclonal (41% of driver mutations, 40% of driver SCNAs and 60% of arm level SCNAs). Significantly lower percentage of clonal driver events were identified in RC patients than patients with both RCC (median percentage, 56% vs. 72%; P=0.031) and LCC (median percentage, 56% vs. 74%; P=0.047) (**Figures S8 and S9**). Hence, our study showed increased diversity in driver events existed in patients with RC.

In addition, no driver events were found consistently clonal among 62 patients (**Fig. 3**), suggesting high ITH status and evolutionary diversity existed among colorectal tumors, which might be the reason of low efficiency of target-based precision medicine in CRC treatment. All the driver SCNAs and most of the driver mutations were identified as “early events” while very few arm level SCNAs were identified as “early events”, suggesting that the genomic instability process occurred firstly at the driver SCNA level, then at the driver mutations level, and finally at the arm level SCNA level.

Driver mutations in *APC, TP53* and *KRAS* were mostly identified in all these 62 patients, which were predominantly clonal and identified as “early event”, suggesting their significance and key roles in tumor initiation. However, except for driver mutations in *APC, TP53* and *KRAS*, other identified driver mutations were completely different between patients with RCC and LCC (**Fig. 3**). The genes of driver SCNAs identified were the same in patients with LCC and RC while only 3 out of 24 genes of driver SCNAs (*CYSLTR2, FLT3* and *FOXO1*) were same in patients with RCC and LCC (**Fig. 3**). These huge differences in both driver mutations and driver SCNAs between the patients with RCC and LCC suggested that patients with LCC were evolutionary closer to the patients with RC than that of RCC.

### Conserved evolutionary features in CRC

In order to understand the constraints and features of CRC evolution, we analyzed conserved patterns of driver events by REVOLVER [31] (**Fig. 4**). Evolutionary trajectories were clustered by the CCF and cluster information of all the driver events in 62 patients and four clusters (cluster red, blue, green and purple) were found (**Fig. 4**). In order to understand whether conserved patterns of CRC evolution correlated to distinct clinical phenotypes, clinical and genomic metrics were shown under 4 clusters (**Fig. 4**).

**Fig. 4.**
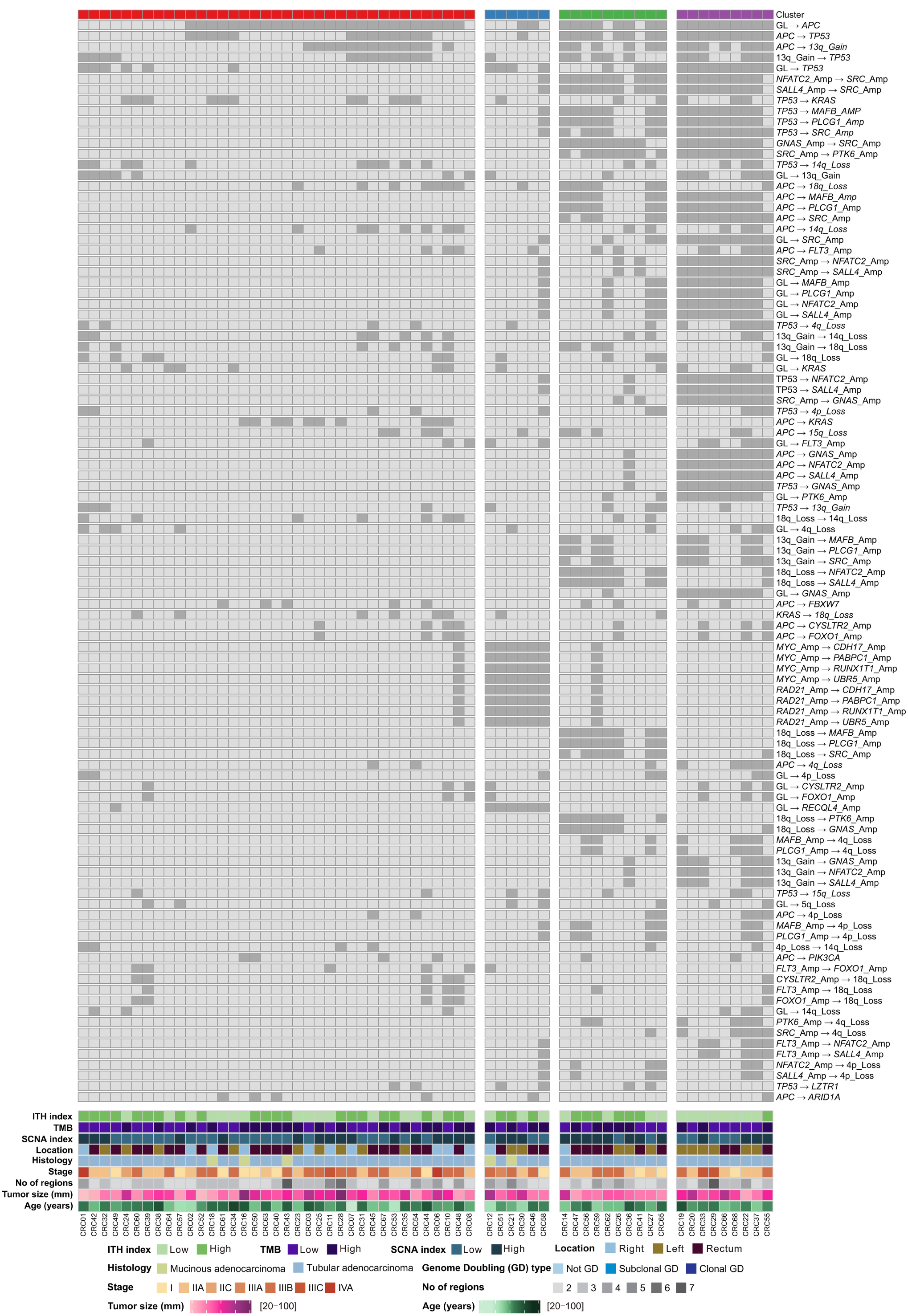
Evolutionary subtypes. Evolutionary trajectories were clustered based on CCF value and cluster information of driver mutations, driver SCNAs and arm-level SCNAs. Heat maps showed the most recurrent evolution for the most recurrent driver mutations, driver SCNAs and arm-level SCNAs. Alterations were ordered by their frequencies in CRC tumors. CRC tumors are annotated by the following parameters: ITH index (high: half of the largest ITH index value; low: the other half), TMB (high> median, low≤ median), SCNA index (high> median, low≤ median), tumor location, histology, stage, number of regions, tumor size and age.

We found that the red and blue clusters had relatively fewer driver events than green and purple clusters. There were no specific genomic or clinical features for the tumors in red cluster. The blue, green and purple clusters had similar clinical features, which were enriched in LCC and RC patients, suggesting that LCC and RC patients were functionally more divergent than RCC patients in evolutionary perspective.

### Phylogenetic distance between LN and ENTD

We analyzed 16 stage III patients to understand the phylogenetic distance and evolutionary relationship amongst primary tumor, LN and ENTD. CRC21, CRC28, CRC43 and CRC48 were identified with both LN and ENTD samples which were sequenced (**Fig. 5**). In CRC21, we identified that the clonal evolution of LN and ENTD was similar, while ENTD appeared evolutionarily later than LN (**Figure S6**). In CRC28, two ENTD samples were clustered together while LN was far away from them, which indicated that the LN and ENTD were polyclonal in origin (**Fig. 5**). In CRC 43 and CRC48, we identified that the ENTD were not clustered together with LN and evolved separately (**Figures 5** and **S6**). In tumors with more than one LN sequenced (CRC01, CRC11, CRC29 and CRC33), some LN were clustered together while some LN were not (**Fig. 5**). In tumors with two ENTD sequenced (CRC60), these two ENTD were far away from each other in the phylogenetic tree (**Fig. 5**). These findings suggested that both LN and ENTD were polyclonal in origin.

**Fig. 5.**
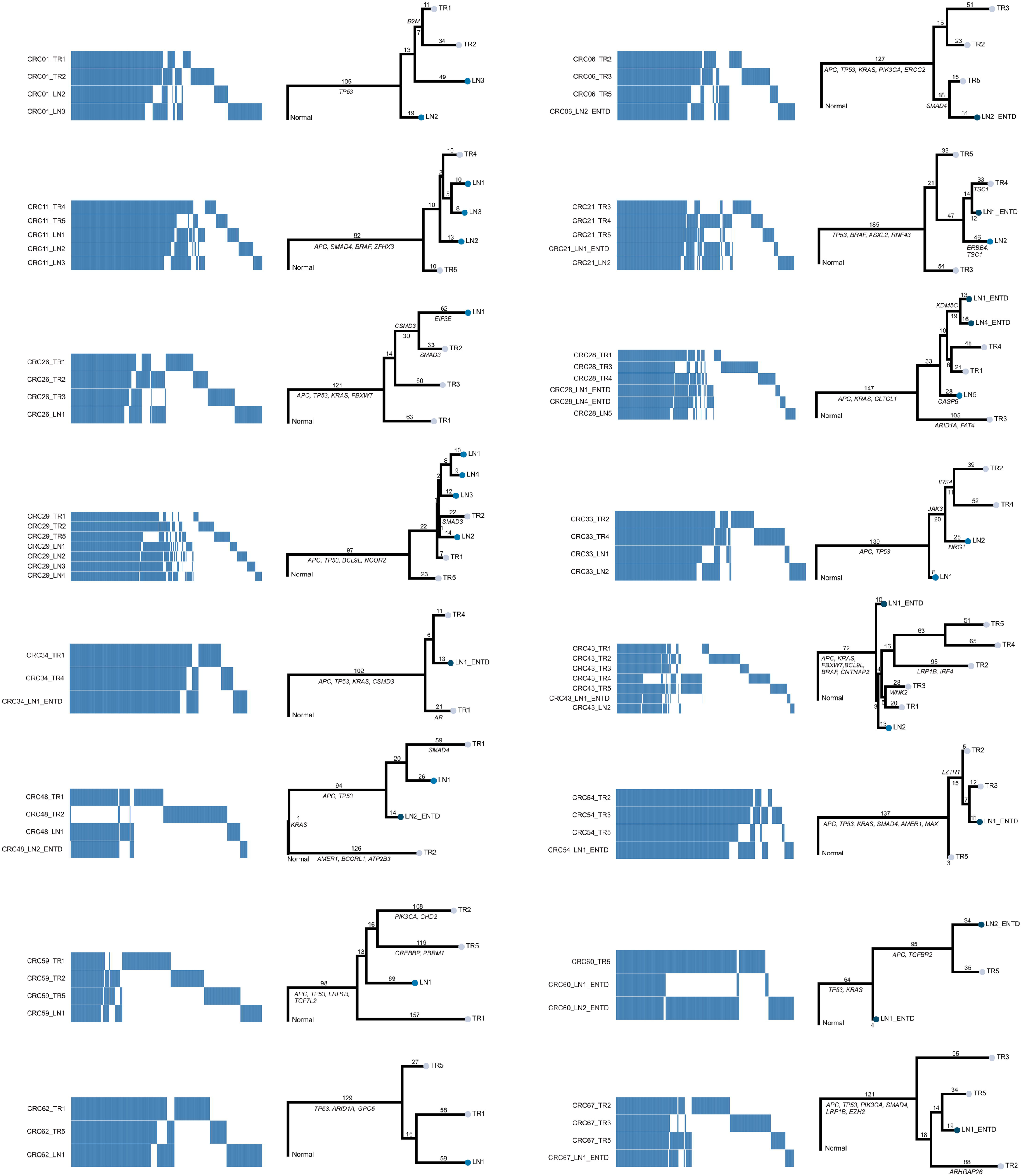
Phylogenetic distance between primary tumor, LN and ENTD. Heatmap showed the presence (blue) and absence (white) of all the mutations (SNVs and INDELs) among different tumor regions of the patients with lymph node metastasis or ENTD. Phylogeny reconstruction using maximum parsimony based on mutational presence or absence of all the mutations were shown beside heatmap. Driver genes were labeled in the phylogenetic trees.

### Evolutionary process at mutational level

#### Convergent features and parallel evolution in CRC

Evidence of convergent mutations in tumor driver genes may shed light on evolutionary selection, which may provide therapeutic targets for treatment. *APC, TP53* and *KRAS* were the most frequently mutated driver genes identified in our study, with mutation frequency of 80.6 % (50/62), 80.6 % (50/62) and 51.6 % (32/62) respectively (**Figure S10**). Among these three genes, *APC* was the most frequent mutated gene in tumor samples. Among these 50 patients with *APC* mutations, 19 (38%) had 2 mutations, consistent with the two-hit hypothesis of *APC* genes in CRC tumorigenesis [32] (**Figure S11**).

Evolutionary selection was also exemplified by parallel evolution of driver mutations, in which different driver mutations in same gene occurred among distinct regions of the same tumor. In CRC36 (LCC patient), two different nonsynonymous mutations in *TP53* were identified in tumor region 3 while another nonsynonymous mutation of *TP53* was detected in tumor region 1 and 4, indicating parallel evolution of *TP53*.

#### Mutation signature

We analyzed mutational processes based on previously published mutational signatures [33]. We found that the age-related signature 1 was the predominant mutational process for all these 62 patients, with a median percentage of age-related mutations of 70% (**Figure S12**).

The median percentage of age-related signature 1 for clonal mutations was 73%, while it dropped to 53% for subclonal mutations (**Figure S12**). This finding suggested that except for age, other mutational processes played more important roles in subclonal than clonal mutations in tumors, which accounted for ITH of CRC. Except for age, other main mutational processes were defective DNA mismatch repair-related signature 6, 15 and defective DNA double-strand break-repair-related signature 3, suggesting that the main mutational process for ITH of CRC were age and defective of DNA repair system.

### Evolutionary process at copy number alteration level

#### Chromosome instability

Previously, we analyzed the length and clonality of SCNAs (**Fig. 1A**), we then measured the SCNAs frequency pattern in RCC, LCC and RC patients. The SCNA frequency pattern in patients with LCC and RC were similar with each other, while RCC patients were very different (**Figure S13**). As shown in **Figure S13**, RCC patients had more 9p gain, 3q gain, 19p loss and less 20q gain, 18p loss, 8p loss than both LCC and RC patients. These results indicated that the SCNAs frequency pattern in CRC patients could be a potential biomarker to distinguish between RCC and LCC/RC patients.

#### Mirrored subclonal allelic imbalance

Recent studies identified parallel evolution of SCNAs in NSCLC and renal cancer through mirrored subclonal allelic imbalance (MSAI) [14, 15]. We identified MSAI events in 23 of 62 patients (37%, found in 5 RCC patients, 6 LCC patients and 12 RC patients) (**Figure S14)**. MSAI parallel gain or loss events found in this study were summarized (**Fig. 6A**). Interestingly, RCC patients had 42% MSAI events, more than both LCC (30%) and RC (40%) patients. We also analyzed parallel evolution of driver SCNAs, 5 tumors (4 tumors with parallel amplification and 1 tumor with parallel deletion) were found to have driver SCNAs which overlapped with MSAI events (**Figs. 6B and C**). Interestingly, 2 of 5 patients (CRC12 and CRC59) were identified with parallel amplification of *FLT3* gene in chromosome 13 (**Fig. 6C**).

**Fig. 6.**
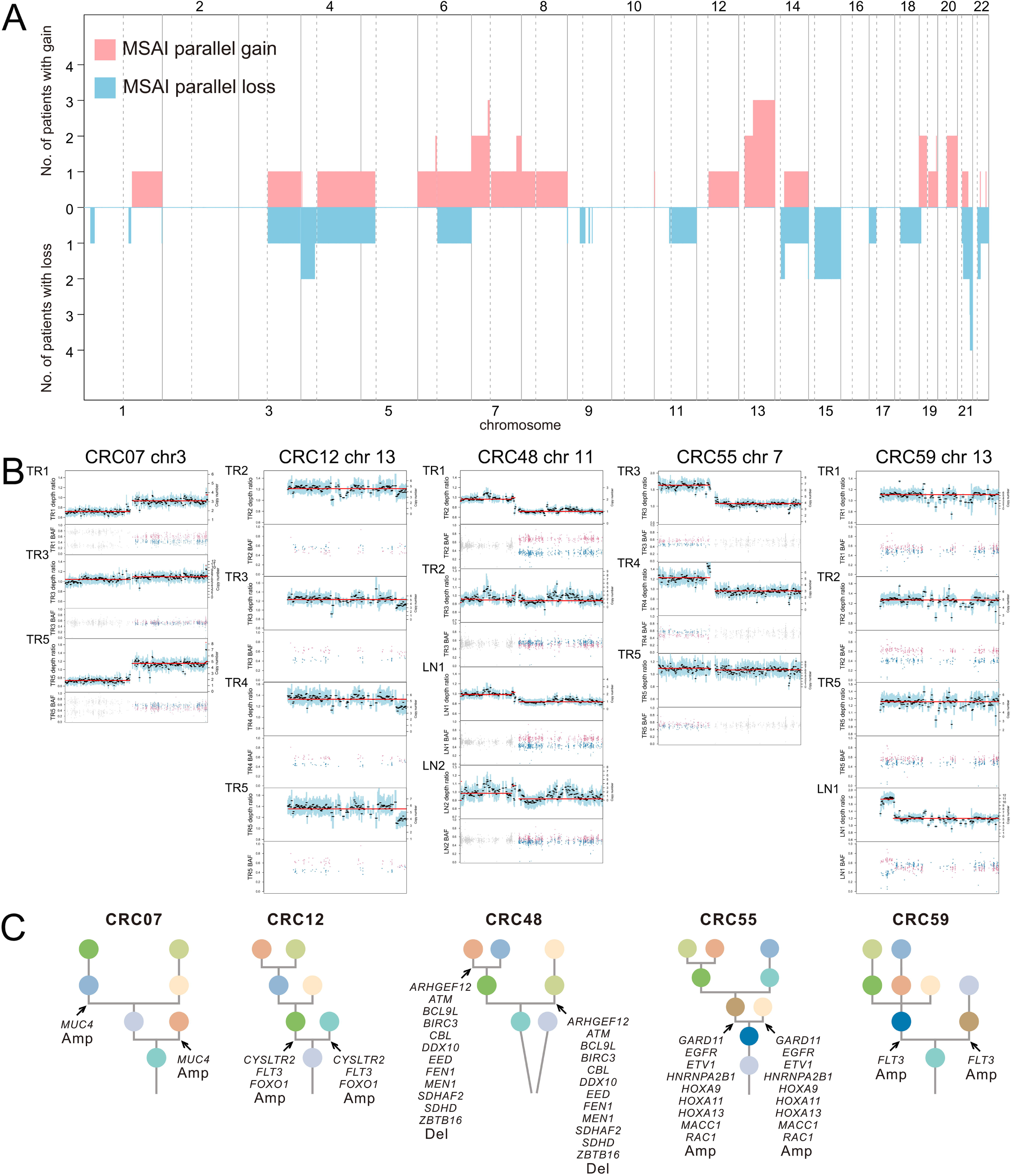
Parallel evolution. **a** Genomic position and size of all mirrored subclonal allelic imbalance (MSAI) parallel gain or loss events found in this study. This included genome-wide copy number gains and losses which was subjected to MSAI events and their occurrence in CRC tumors. **b** Parallel evolution of driver SCNAs observed in 5 CRC tumors, indicted by the depth ratio and B-allele frequency values of the same chromosome on which the driver SCNAs were located. **c** Phylogenetic trees that indicated parallel evolution of driver amplifications (Amp) or deletions (Del) (Driver SCNAs) detected through the observation of MSAI (arrows).

### Evolution landscape of hypermutated CRC tumors

All 6 (CRC04, CRC05, CRC09, CRC13, CRC15 and CRC17) hypermutated CRC patients were identified with RCC, of these two patients (CRC09 and CRC13) were with MSS and remaining four patients (CRC04, CRC05, CRC15, CRC17) were with MSI tumors (**Figure S15A**). All the 6 hypermutated patients had mutations in mismatch-repair genes, or in *POLE* or *POLD* gene family (**Figure S15A**). CRC09 had one missense mutation and one nonsense mutation of *POLE*. CRC13 had one missense mutation of *POLE* (**Figure S15A**). These findings were consistent with the predominant mutational process in these two patients with MSS tumors was *POLE*-related signature 10 (**Figure S15B**). Defective DNA mismatch repair-related signature 6, 15, or 26 contributed to the mutational process of 4 patients with MSI tumors (**Figure S15B)**. We also analyzed the evolution landscape of hypermutated tumors in SCNA level. The absolute SCNAs of hypermutated CRC patients occurred less (**Figure S15C**), which suggested that these hypermutated CRC patients were mainly having mutation driven tumors. Interestingly, CRC04 had MSAI events in X-chromosome (**Figure S16**).

## Discussion and conclusions

In this present study, we performed high-depth WES and analyzed 206 multi-region tumor samples from 68 patients with CRC. Our result showed that the LCC patients were structurally and functionally more complex and divergent than RCC patients in terms of evolutionary perspective. Our result showed ENTD were later events in the evolution of the tumor than LN. In addition, all the CRC patients followed the Darwinian pattern of evolution.

### RCC, LCC and RC patients: In the light of clonal evolution

Previous studies have shown remarkable differences among RCC, LCC and RC based on genetic mutations, genomic expression profiles, immunological composition and bacterial population in tumor microenvironment [20-24]. However, almost no research has been done till date for understanding the differences between different locations of CRC from evolutionary perspective, which is the key to explore the differences among RCC, LCC and RC in tumor initiation and progression. Our study demonstrated that ITH and evolution among LCC, RCC and RC patients were different in the following aspects: mutations, SCNAs, structure of polygenetic tree and driver events. Firstly, RC patients had shown fewer clonal mutations than RCC patients, indicating higher ITH in RC patients at mutational level. Secondly, the SCNAs frequency pattern in RCC patients were different from LCC and RC patients, which addressed the evolutionary difference between them at SCNAs level. Thirdly, the structure of phylogenetic trees in LCC and RC patients were more complicated and branched than that of the RCC patients. Specifically, LCC patients were identified with the most complicated structure of the phylogenetic tree, reflected by more cluster numbers. In addition, only LCC and RC patients were polyclonal in origin. Fourthly, LCC and RC patients were enriched in clusters (blue and purple clusters) which had more driver events, indicated that LCC and RC patients showed more functional diversity in evolution. Moreover, RC patients were identified with less percentage of clonal driver events than both LCC and RCC patients, suggested that more functional diversity occurred in the process of evolution of RC patients. In conclusion, our data showed that LCC and RC patients were more divergent and complicated in terms of evolution than RCC patients, not only structurally but also functionally, which indicated that the evolutionary diversity might play an important role in the initiation and progression of CRC among LCC and RC patients. Furthermore, the SCNA frequency pattern could be a potential and significant biomarker to distinguish between RCC, LCC and RC patients.

### Primary tumor, LN and ENTD: In evolutionary perspective

To date, no systematic research studies have been done to understand the similarities and differences between ENTD and LN. In this study, we found that ENTD were later events in the evolution of the tumor than LN according to the clonal evolution history in CRC21. LN and ENTD could not be clustered together in the polygenetic tree according to the occurrence of mutations. Unlike in previous studies [10, 12], different LN or ENTD in the same tumor did not cluster together in all cases, indicating their polyclonal origin. In conclusion, ENTD was a distinct entity from LN and evolved later.

### Evolution pattern: Darwinian pattern of evolution and neutral evolution

In this present study, we found predominantly Darwinian pattern of evolution (59 out of 62 patients) as well as linear evolution (3 out of 62 tumors). Previous studies proposed neutral evolution model for colorectal cancers [6, 34, 35], whilst our conclusion was different from them, based on three reasons. Firstly, clonal events of both mutations (SNVs and INDELs) and SCNAs were widespread, with a median percentage of 47% and 43% respectively. Secondly, 59% of driver mutations were clonal while only 41% of non-driver mutations were clonal, which indicated the enrichment of clonal driver mutations in course of evolution. Lastly, convergent and parallel events were present for driver genes in both mutational and SCNA level, especially for genes *APC, TP53* and *KRAS*. Previous studies also showed Darwinian pattern of evolution for the patients with colorectal cancer followed by neutral evolution [8, 9]. In our study, we identified that 28% of subclonal mutations were shared by tumor regions (either branch or trunk mutations), which suggested the importance of branches in phylogenetic trees.

## Methods

### Patient recruitment, sample collection and sample processing

The study was approved by the Ethics committee of the Affiliated Hospital of Qingdao University. All the samples were collected after obtaining written informed consent from the patients.

Detailed process of sample collection and sample processing has been given in Supplemental methods. The filtering pipeline is schematically presented in the CONSORT diagram (CONSORT flowchart, **Figure S1**). The workflow summarizing experiments and data analysis in our study was shown in **Figure S2**.

### Pathology diagnoses and review

Detailed process of pathological diagnoses and review has been given in Supplemental methods. Clinical details of 68 CRC patients were summarized in **Table S1**.

### WES and quality control

WES was performed for tumor tissues and matched germline tissues. Detailed process of WES and quality control has been given in the Supplemental methods.

### Somatic mutation detection and filtering

All mutations used in the analysis can be found in **Table S2**. Detailed process of somatic mutation detection and filtering has been given in the Supplemental methods.

### Driver mutation identification and copy number analysis

Detailed process of identification of driver mutations and copy number analysis has been given in the Supplemental methods. Somatic copy number alterations (SCNAs) were identified and all segmented copy number data has been given in **Table S3**.

### Sub-clonal deconstruction and phylogenetic tree construction

Sub-clonal deconstruction and phylogenetic trees were constructed. Clusters for phylogenetic tree construction were summarized in **Table S4**. Detailed process of sub-clonal deconstruction and phylogenetic tree construction has been given in the Supplemental methods.

### Analysis of evolution subtype and phylogenetic analysis

Evolutionary subtypes were clustered and visualized. Phylogenetic distance between primary tumor, LN and ENTD were analyzed. Detailed process of evolution subtype and phylogenetic analysis has been given in the Supplemental methods.

### Mutation signature analysis

Mutation signatures were estimated. Detailed process of mutation signature analysis has been given in the Supplemental methods.

### Mirrored sub-clonal allelic imbalance and statistical analysis

Mirrored sub-clonal allelic imbalance and statistical analysis were performed. All statistical analyses were performed in R statistical environment version >= 3.5.0. Detailed process of analysis of mirrored sub-clonal allelic imbalance and statistical analysis has been given in the Supplemental methods.

## Acknowledgements

We are thankful to the proband and all the family members for participating in our study and we are thankful to the China National GeneBank.

## Authors’ contributions

Conception and design: Santasree Banerjee, Shan Kuang, Junnian Liu, Yun Lu, Xin Liu; Acquisition of data (provided animals, acquired and managed patients, provided facilities, etc.): Xianxiang Zhang, Qingyao Wu, Shujian Yang, Jigang Wang, Xiaobin Ji, Peng Han, Yong Li, Xiaofen Tian, Zhiwei Wang; Analysis and interpretation of data (e.g., statistical analysis, biostatistics, computational analysis): Santasree Banerjee, Shan Kuang, Lei Li, Shui Shun, Li Deng, Yue Zhang; Writing, review, and/or revision of the manuscript: Santasree Banerjee, Shan Kuang, Lei Li, Xianxiang Zhang, Jigang Wang; Administrative, technical, or material support (i.e., reporting or organizing data, constructing databases): Huanming Yang, Lars Bolund, Yonglun Luo, Kui Wu, Shida Zhu, Guangyi Fan, Xun Xu; Study supervision: Santasree Banerjee, Shan Kuang, Junnian Liu, Yun Lu, Xin Liu.

## Availability of data and materials

The sequencing data has been deposited at the CNGB Nucleotide Sequence Archive (CNSA: https://db.cngb.org/cnsa), under accession number CNP0000594.

