## Supplementary Data for "Comparative analysis of clonal evolution among patients with right-sided colon cancer, left-sided colon cancer and rectal cancer"

**Supplemental methods**

**Patient recruitment**

The study was approved by the Ethics committee of the Affiliated Hospital of Qingdao University. All the samples were collected after obtaining written informed consent from the patients. Patients were recruited based on the following criteria. (i) age over 18 years, (ii) patients clinically diagnosed with CRC by enteroscopy, imaging, biopsy and followed by surgery, and histopathology performed with the resected tumor tissues. Patients with sufficient tissue were available for the study.

**Sample collection**

A pathologist performed macroscopic examination of all surgically resected specimens to guide the multi-region sampling in this study. Firstly, the pathologist performed routine pathological sampling for clinical diagnosis, and then multi-region sampling was performed by using the remaining samples. At least 2 regions of each tumor, which were at least 3 mm apart, were collected. Areas with significant necrosis, fibrosis, or hemorrhage were avoided to maximize the viability of tumor cells. Normal colorectal mucosa tissues were also sampled from areas remote from the primary tumor (at least 2 cm distant from the tumor edge). Peri-intestinal nodules including lymph nodes present in the resected specimen were sampled. If there was malignancy appearance (the cut section appeared tan-gray and hard), after confirming the malignancy, a portion of the lymph nodes was sampled for diagnostic requirements. The remaining part was taken for this study. Each selected tissue block was split into two for snap freezing and formalin fixing respectively (mirrored FFPE sample). Fresh samples were placed in a 2 ml cryotube, and snap frozen with immediate immersion into liquid nitrogen before transferred to -80°C freezer for storage. Peripheral blood was collected and processed into EDTA anticoagulation tube. The tumor tissue samples from 68 patients were sequenced and analyzed after filtering according to the filtering pipeline, schematically presented in the CONSORT diagram (CONSORT flowchart, Supplementary Fig. S1). The workflow summarizing experiments and data analysis in our study was shown in Supplementary Fig. S2.

**Sample processing**

Approximately 50 mm^3^ of tumor tissue from each region was used for genomic DNA extraction using the QIAamp DNA Mini Kit (Qiagen, Germany) according to the manufacturer’s instructions. 2 ml of peripheral blood was used for germline DNA extraction using the QIAamp DNA Blood midi kit (Qiagen, Germany) according to the manufacturer’s instructions. DNA was quantified by the Qubit Fluorometric Quantitation (Thermo Fisher Scientific, USA) and the quality of DNA was assessed by agarose gel electrophoresis.

**Pathology diagnoses and review**

Pathological diagnoses were established according to the WHO classification and independently reviewed by two pathologists. Clinical details were summarized in Supplementary Table S1. Hematoxylin- eosin sections of mirrored FFPE samples for each region in every case (387 sections from 70 patients) were evaluated. Only primary tumor regions with more than 30% tumor component and pathological heterogeneity were considered for sequencing. In addition, pathologist distinguished LN and ENTD by reviewing hematoxylin-eosin sections of their mirrored FFPE samples in this study were also sent for sequencing.

**Whole exome library construction and sequencing**

Tumor tissues and matched germline tissues were subjected to whole exome sequencing. Exome capture was performed on 1 μg of genomic DNA. Covaris (LE220) was used to randomly fragmented DNA into 150-250 bp. These fragments were purified and connected through a PE Index Adaptor designed by BGI, and then captured by using the the MGIeasy Exome Capture V4 probe set (~ 59 Mb; MGI Tech Co., Ltd, China). All constructed libraries were loaded onto BGISEQ-500 (MGI Tech Co., Ltd, China) and the sequences were generated as 100-bp paired-end reads.

Sequencing reads containing sequencing adapters, more than 10% of unknown bases and low-quality bases (> 50% bases with quality <5) were removed by SOAPnuke (v1.5.6) [1]. The processed sequencing reads were then aligned to UCSC human reference genome (hg19) using BWA-MEM (v0.7.12) [2]. Picard (v1.137) (https://broadinstitute.github.io/picard/) was used to generate chromosomal coordinate-sorted bam files to remove PCR duplicates. Then, the median sequencing depth of the generated data for the tumor area were reached 391 (range 179-537), and the matched germline tissues were reached 414.5 (range 243-596). We then used the Genomic Analysis Toolkit (GATK v3.8.0) [3] to perform base quality score recalibration and local realignment of the aligned reads to improve alignment accuracy.

**Quality control to prevent contamination, inter-patient sample swaps and removal of regions with extremely low mutation occurrence**

ContEst [4], a GATK module, was used to estimate the cross-individual contamination level. Samples with contamination level more than 1% were deleted (3 samples failed the QC due to contamination as shown in Supplementary Fig. S1. In order to avoid sample swaps between patients, we used BAM-matcher [5].

The number of mutations in each tumor region was called independently. The median number of mutations across all regions for each tumor was calculated. A region in one tumor was removed if less than 20% of the median mutation count of that tumor was identified in that region.

**Somatic mutation detection and filtering**

After processed the sequencing data, SAMtools (v1.2) [6] mpileup was used to locate non-reference locations in tumor and germline samples. Bases with phred scores less than 20 or reads with mapping quality (MAPQ) values less than 20 were deleted. Base-alignment quality (BAQ) computation was disabled with adjust mapping quality coefficient set of 50. Both VarScan 2 (v2.4.3) [7] and MuTect (v1.1.7) [8] were used to call somatic mutations. The somatic variants called by VarScan 2 were filtered and the minimum coverage of the germline sample was set to 10, the minimum variant frequency was changed to 0.01, and tumor purity was set to 0.5. We further filtered the resulting single nucleotide variant (SNV) calls for false positives using Varscan 2 associated fpfilter.pl script. We used bam-readcount (v0.8.0) (https://github.com/genome/bam-readcount) to prepare input files for fpfilter and min-var-freq was set to 0.02. All insertions/deletions (INDELs) called in reads that VarScan 2 processSomatic classified as "high confidence" were recorded for further downstream filtering. MuTect was used to detect SNVs using annotation files contained in GATK bundle (v2.8) and variants were filtered according to the filter parameter ‘PASS’.

Additional filtering was performed to reduce false positive mutation calls. If the variant allele frequency (VAF) is greater than 2%, and both VarScan 2 (with a somatic p-value <= 0.01) and MuTect called the mutation, then a SNV was considered as truly positive. Alternatively, if a SNV was called only in VarScan 2 with a somatic p-value <=0.01, a frequency of 5% was required. In addition, the sequencing depth supporting the variant call in each region required >= 30, and the sequence reads required >= 5. In contrast, the VAF value of the variant in the germline should be <= 1%. We filtered the INDEL using the same parameters as above, except that reads >= 10 were required to support mutation calls, somatic p-values <= 0.001 and sequencing depth >= 50.

ANNOVAR [9] was used to annotate mutations with COSMIC (v88) [10], SIFT [11], PolyPhen-2 [12] and MutationTaster [13] databases. All mutations used in the analysis can be found in Supplementary Table S2. Mutations were classified as clonal or subclonal using PyClone (v0.13.1) [14]. PyClone CCF (cancer cell fraction) value were calculated as described in the subclonal deconstruction section. Mutations with CCF>0.9 across all regions of a tumor were considered as clonal mutations, otherwise they were considered as subclonal mutations.

**Driver mutation identification**

All variants were compared with all genes identified and enlisted in the COSMIC Cancer Gene Census (v88) [10]. Then, three types of mutations were classified as a driver mutation according to the following criteria. Firstly, if the gene was annotated as TSG (tumor suppressor gene) by COSMIC, and the non-silent variant was considered deleterious: either *loss of function* (stop-gain/stop-loss, frameshift deletion/insertion or non-frameshift insertion/deletion) or predicted deleterious in two of these three computational approaches applied – SIFT [11], PolyPhen-2 [12] and MutationTaster [13], then the specific variant would be classified as a driver mutation. Secondly, if the variant was annotated as oncogene by COSMIC, then we tried to identify exact matches to non-silent variants in COSMIC. If an exact match was found ≥ 3 times, the variant was categorized as a driver mutation. Thirdly, if the gene was annotated as TSG by COSMIC, and the variant is located at the canonical splice site, then the specific variant would be classified as a driver mutation. Finally, we compared all these three types of driver mutations to the CpG island location file on UCSC Genome Bioinformatics website (http://genome.ucsc.edu). We then deleted all mutations that occurred on the CpG island and finally got all driver mutations.

**Copy number analysis**

Sequenza (v3.0.0) [15] was used to detect the somatic copy number alterations (SCNAs) and evaluate the purity and ploidy of tumor cells as follows. Firstly, we used SAMtools (v1.2) [6] mpileup to convert the Bam file to Pileup format. Secondly, paired tumors and normal Pileup files were processed by sequenza-utils to extract the sequencing depth, determine the homozygous and heterozygous positions of variants in normal samples, and calculate the variant alleles and allelic frequencies from tumor samples. The sequenza-utils output was further processed by using Sequenza R package to provide segmented copy number data, cellularity and estimated ploidy for each sample. All segmented copy number data has been given in Supplementary Table S3. Heatmap of genome-wide SCNAs is visualized by R package Copynumber (v1.24.0) [16].

The driver gene copy number variations (driver SCNAs) of all genes enlisted in the COSMIC cancer gene census were analyzed as follows. Firstly, if the gene was annotated as oncogene by COSMIC, gene level amplification was called if gene copy number >2 × ploidy of that sample. Secondly, if the gene was annotated as TSG by COSMIC, gene level deletion was called if gene copy number = 0. To determine the ITH status of driver SCNAs, we called driver SCNAs across all regions from each tumor. If at least one region showed an amplified SCNA, we called a gene as clonal amplification if all other regions of this gene showed copy number > ploidy + 1. If at least one region showed a deleted SCNA, we called a gene as clonal deletion if all other regions of this gene showed copy number < ploidy -1. All other driver SCNAs were defined as subclonal amplification or deletion. In 8 polyclonally originated tumors (CRC32, CRC36, CRC42, CRC48, CRC49, CRC51, CRC52 and CRC60) without founder clusters (cluster with CCF > 0.9 across all regions of a tumor), all their driver SCNAs were subclonal. To correlate driver SCNAs with specific mutation clusters of PyClone, we first identified all clusters where >= 50% CCF was present in each tumor region. We then identified all the clusters present in the same regions as a given driver SCNAs. We called a gene as clonally amplified if all the regions of this gene showed copy number >2 × ploidy while we called a gene as clonally deleted if all the regions of this gene showed copy number = 0. Then we repeated the association test above. If an SCNA still could not be associated with a mutant cluster, it was annotated as a subclone associated with no known cluster (NA cluster).

To determine the ITH status of global SCNA, all parts of the genome were considered independently and divided into the smallest contiguous segments that overlap in all the regions within each tumor. The gains and losses of segment were determined as follows. Firstly, copy number data for each segment was divided by the sample mean ploidy and then converted to log_2_. Secondly, gain and loss were defined as log_2_ (2.5/2) and log_2_ (1.5/2), respectively. Thirdly, any segment of gain or loss that spanned across all the regions was defined as clonal and all other segments of SCNA were defined as subclonal. Within each tumor, we summarized the length of the genome that subjected to SCNA in any region (total SCNA), the length of the genome that subjected to clonal SCNA (clonal gain or clonal loss), and the length of the genome that subjected to subclonal SCNA (subclonal gain, subclonal loss or subclonal undetermined). The proportion of subclonal SCNAs were then defined as the percentage of genomes subjected to subclonal SCNA divided by the percentage of genomes subjected to total SCNAs.

Chromosomal arm level SCNAs were determined if at least one region has shown an increase or decrease of at least 97% in chromosomal arm. To determine the ITH status of chromosome arm gain and loss, we called clonal arm gain or loss if the same chromosomal arm showed at least 75% gain or loss in all the remaining regions. While we called subclonal arm gain or loss if at least one of the remaining regions showed less than 75% gain or loss. In 8 polyclonally originated tumors, all their arm level SCNAs were subclonal. As previously described in the driver SCNAs part, we correlated arm level SCNAs with specific mutation clusters of PyClone in the same way.

**Sub-clonal deconstruction**

In order to estimate whether mutations were clonal or subclonal, and the phylogenetic trees of each tumor, the following formula were used [17, 18]:

$$vaf=\frac{{CN}_{mut}\times CCF\times p}{{CN}_{n}\times\left( 1-p \right)+{CN}_{t}\times p}$$

Where *vaf* is the mutated allele frequency of the mutated base; *p* is the estimated tumor purity; *CNt* is tumor locus specific copy number; *CNn* is normal locus specific copy number, assuming 2 for autosomal chromosomes; *CCF* is the fraction of tumor cells carrying mutations. Considering that *CNmut* is the copy number of the chromosome harboring the mutation, the possible *CNmut* range is from 1 to *CNt* (integer). We then assigned one of the possible values to *CCF*: 0.01, 0.02, ..., 1, together with every possible *CNmut* to find the best fit *CCF* using maximum likelihood. In detail, for point mutations with alternative reads as “*a*” and sequencing coverage as “*N*”, we used Bayesian probability theory and binomial distribution to estimate the probability of a given *CCF*:

$$P \left( CCF | \left( a | N \right) \right)\propto Binom (a|N, {vaf}_{ex}(CCF))$$

Then, the distribution of *CCF* was obtained by calculating *P (CCF)* on 100 uniform grids with *CCF* values from 0.01 to 1 and dividing by their sum.

Then, we used PyClone (v0.12.9) [14] Dirichlet process clustering to cluster all the mutations (SNVs and INDELs). For each mutation, we used the observed mutation count and set the reference count so that vaf equal to half of the CCF value calculated by maximum likelihood previously. We set the major allele copy number to 2, the minor allele copy number to 0 and the purity to 0.5 since they had been modified.

Since the vaf values of INDELs were potentially unreliable, we multiplied each estimated INDEL CCF with a region-specific correction factor, which was calculated by dividing the median mutation CCF of the ubiquitous mutations (mutations presented in all regions) in that region by the median INDEL CCF of the ubiquitous INDELs (INDELs presented in all regions) in that region. We ran PyClone with 10,000 iterations and a burn-in of 1000.

**Phylogenetic tree construction**

Phylogenetic trees were constructed using the published tool CITUP (v0.1.0) [19]. As input, CITUP requires mutation clusters and their mean cancer cell prevalence values which were collected from PyClone. All clusters with at least 5 mutations were used as input to CITUP. Clusters for phylogenetic tree construction were summarized in Supplementary Table S4. The optimal phylogenetic trees for each patient from CITUP were illustrated using MapScape (v1.8.0) [20].

**Evolution subtype analysis**

Evolutionary subtypes were clustered and visualized by REVOLVER (v0.2.0) [21]. CCF values and cluster information of driver events were processed as previously described, which were used as input to REVOLVER. REVOLVER requires a founder cluster for all the input tumors. Therefore, we artificially defined a founder cluster for 8 polyclonally originated tumors. ITH index was calculated as the numbers of subclonal driver events divided by the numbers of clonal driver events, and SCNA index was indicated by the length of total SCNA.

**Phylogenetic analysis**

Phylogenetic distance between primary tumor, LN and ENTD were analyzed by using the binary matrix of mutations present or absent in each region of tumors with LN or ENTD. Private mutations of each region were discarded from phylogenetic tree building due to lack of information. Fake outgroups with no mutations were generated for each individual as a root. Phylogenies were constructed using the PHYLIP (v3.697) [22] suite of tools. For each tumor, we used seqboot to generate 100 bootstrap replicates by resampling of the mutations with replacement.

Phylogenetic trees were then constructed for each bootstrap replicate by maximum parsimony using the Mix programme in Wagner method. The jumble = 10 option was used and the order of the input samples was randomized 10 times for each bootstrap replicate. Finally, the Consense program was used to build a consensus of all the phylogenetic trees by using the majority rule (extended) option. Phylogenetic trees were redrawn by FigTree (v1.4.4) [23] with the length of trunks and branches, proportional to the number of mutations.

**Mutation signature analysis**

Mutation signatures were estimated by using the DeconstructSigs (v1.8.0) [24] package in R. Mutational signature analysis was applied only in the presence of at least 15 mutations.

**Mirrored sub-clonal allelic imbalance analysis**

Single nucleotide polymorphisms (SNPs) were called by using Platypus (v0.8.1) [25] and only SNPs with a minimum coverage of 20× were analyzed. The B allele frequency (BAF) of each SNP was calculated as the ratio of reads of reference base to variant. Heterozygous SNPs and BAFs were used as input and mirror subclone allelic imbalances (MSAI) were analyzed and visualized by RECUR [26].

Parallel evolution events for driver SCNAs were identified as follows. Firstly, driver SCNAs were identified as described in the "copy number analysis" section. Secondly, we annotated the regions of MSAI events in each tumor to the events of driver SCNAs. If two events coincided with each other, then these driver SCNAs undergone parallel evolution.

**Statistical analysis**

All analyses were performed in R statistical environment version >= 3.5.0. All statistical comparisons of two distributions used the Wilcoxon test (wilcox.test function in R).
