## Supplementary figures for "Comparative analysis of clonal evolution among patients with right-sided colon cancer, left-sided colon cancer and rectal cancer"

#### Supplemental Figures

#### Figure S1

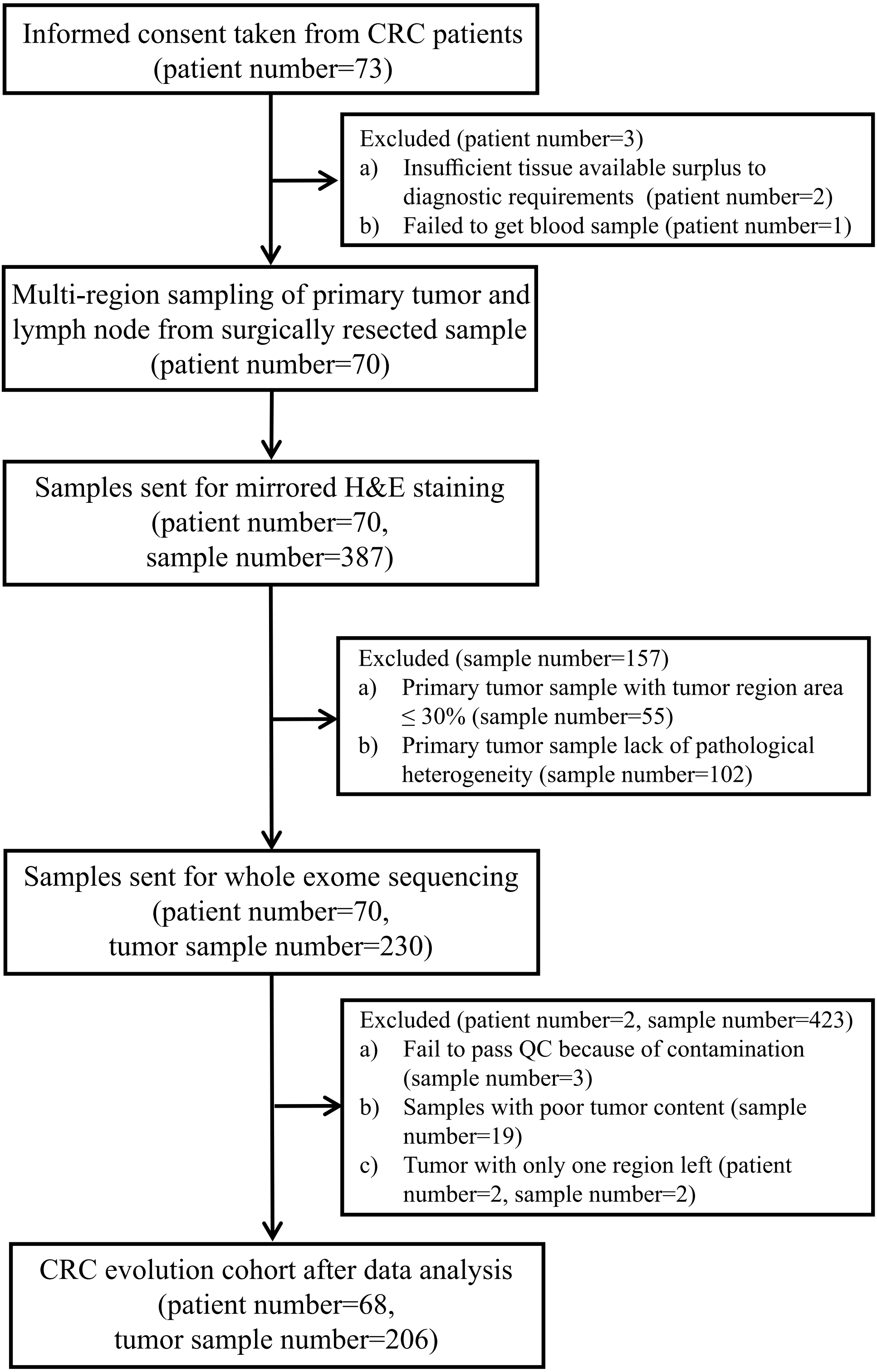

**Figure S1. CONSORT diagram for patient recruitment in this study and eventual selection of the 68 patients cohort.**

### Figure S2

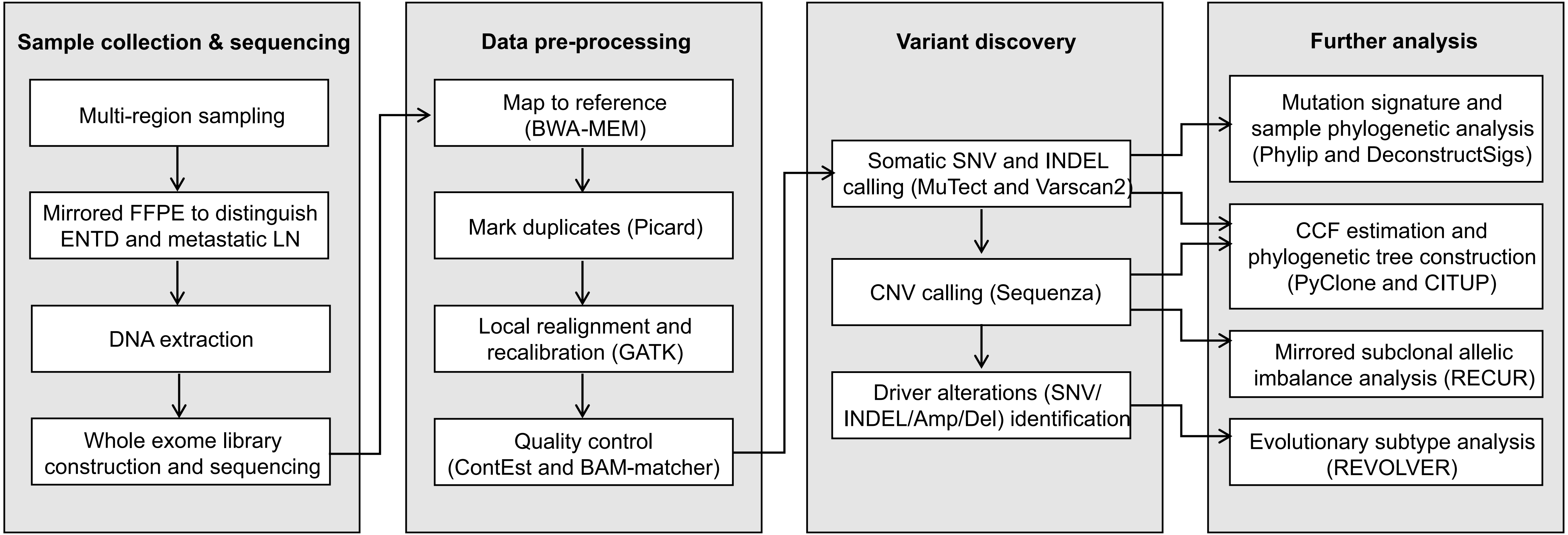

**Figure S2. Workflow summarizing experiments and data analysis.**
Overview of experiments and analysis workflow based on whole-exome sequencing of multi-region CRC tumors (primary tumors, lymph node metastasis and ENTDs) and paired normal controls. Analysis tools/methods are indicated by parentheses.

**Figure S3**

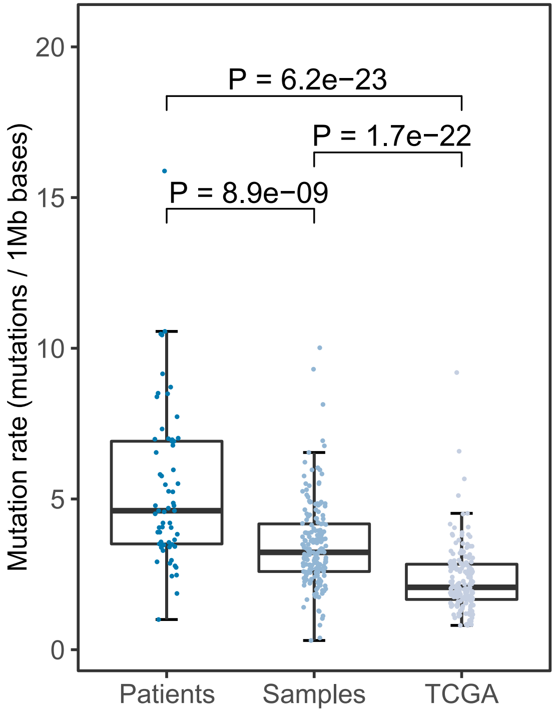

**Figure S3. Comparison of tumor mutation rate between TCGA and CRC tumors.** Box plots of mutation rate in non-hypermutated CRC patients analyzed as single samples, multi-region samples of non-hypermutated CRC patients and TCGA non-hypermutated CRC samples. The definition of hypermutated patients is all the samples in these patients have more than 10 mutations/1 Mb bases. In our CRC tumors, 6 patients are hypermutated patients and all other 62 patients are included into analysis.

### Figure S4

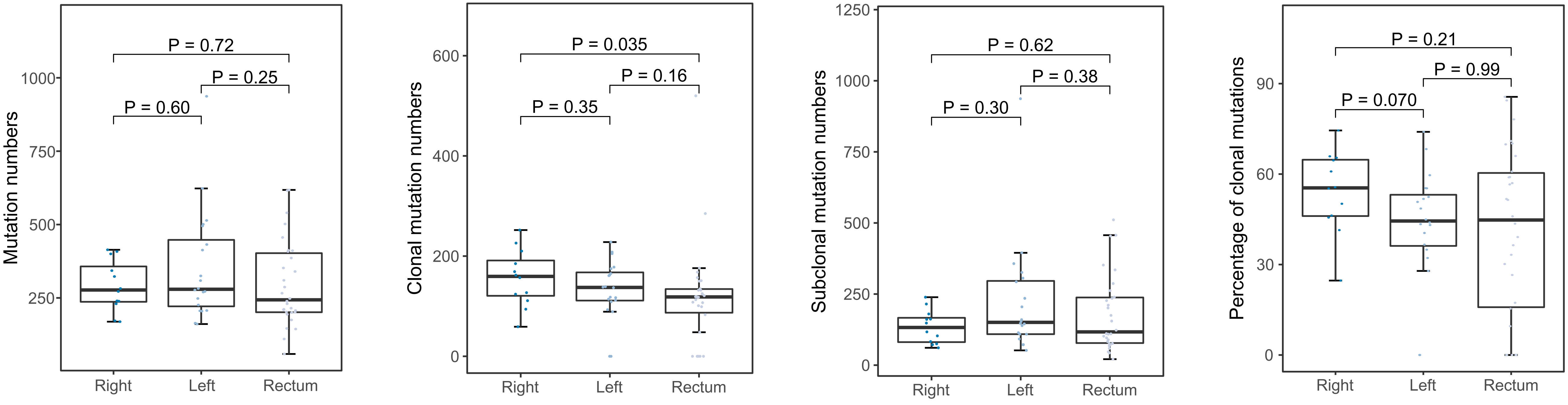

**Figure S4. Intratumor heterogeneity of mutations among right-sided colon, left-sided colon and rectal cancers.** Box plots of total, clonal, subclonal and percentage of clonal mutations by tumor position.

### Figure S5

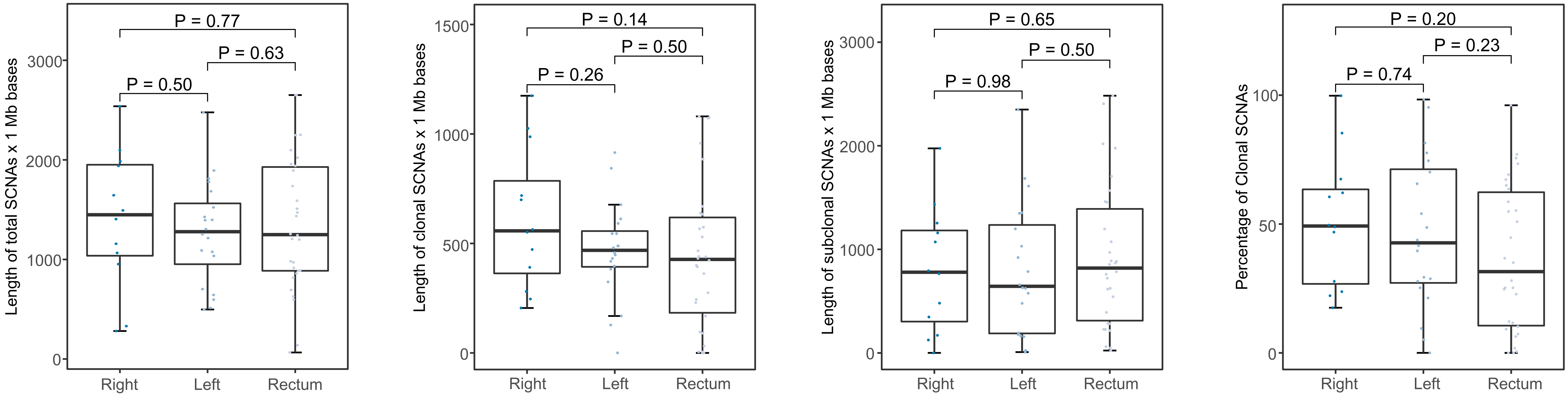

**Figure S5. Intratumor heterogeneity of somatic copy number alterations (SCNAs) between right-sided colon, left-sided colon and rectal cancers.** Box plots of total, clonal, subclonal and percentage of clonal SCNAs by tumor position.

### Figure S6

**Figure S6. Phylogenetic trees.**

Phylogenetic trees for 62 non-hypermutated CRC tumors. For each multi-panel figure, top-left panel showed cancer cell fractions (CCF) as a heatmap for all clustered mutations. CCF value for each mutation represents the mean of the mutation cluster CCF values, with darker color indicating higher CCF value. Regions were indicated below heatmap, "TR" represented primary tumor regions, "LN" represented lymph node metastasis regions and "LN_ENTD" represented extranodal tumour deposits regions. Middle top panel showed the complete phylogenetic tree as constructed based on the mutation clusters. Cancer driver genes found in the tumor, with cytoband, type [mutation (SNV and InDel)/copy number aberration (Amp and Del)] and mutation cluster were indicated on the right, with a colored bar representing individual clusters of driver genes. Below were phylogenetic trees drawn for each region. Grid of 100 representative cells were shown beneath the tree. The colors within each cell represented the mutational clusters.

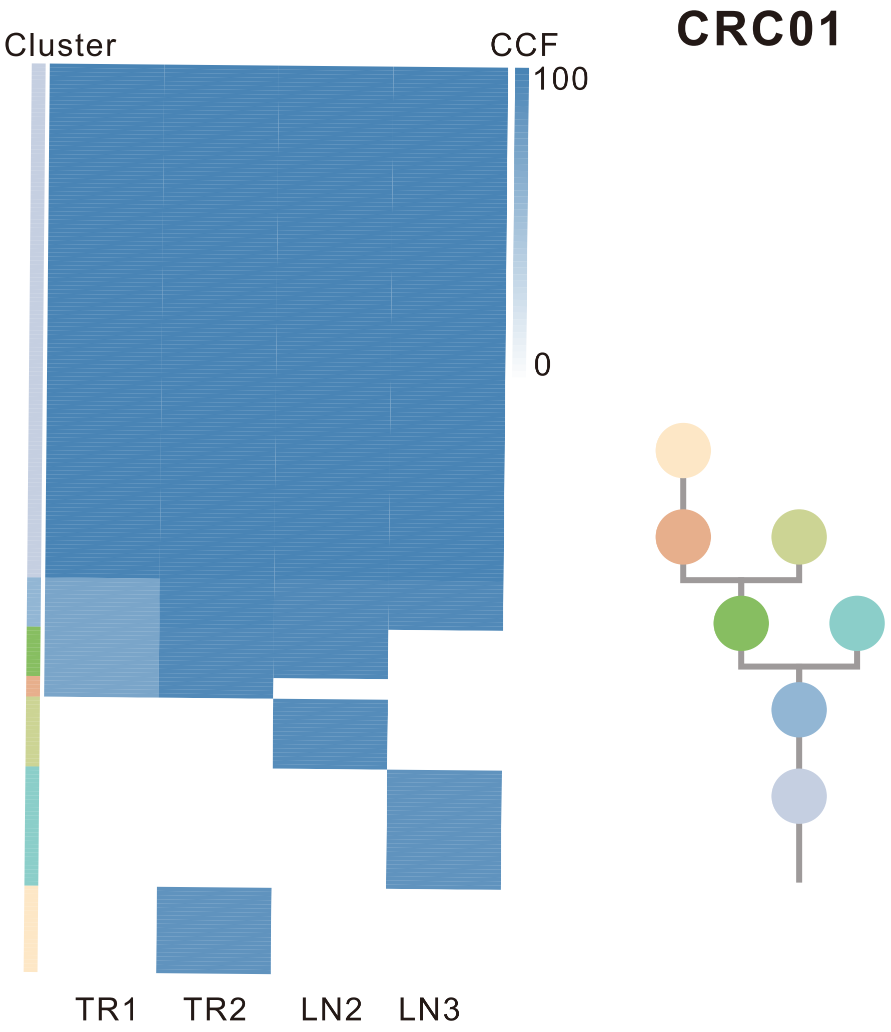

| Gene | Cytoband | Type | Cluster |
|---|---|---|---|
| *TP53* | 17p13.1 | SNV | 1 |
| *MUC4* | 3q29 | Amp | 1 |
| *BCL9* | 1q21.2 | Amp | 4 |
| *TCF3* | 19p13.3 | Del | 4 |
| *B2M* | 15q21.1 | InDel | 7 |
| *STK11* | 19p13.3 | Del | NA |

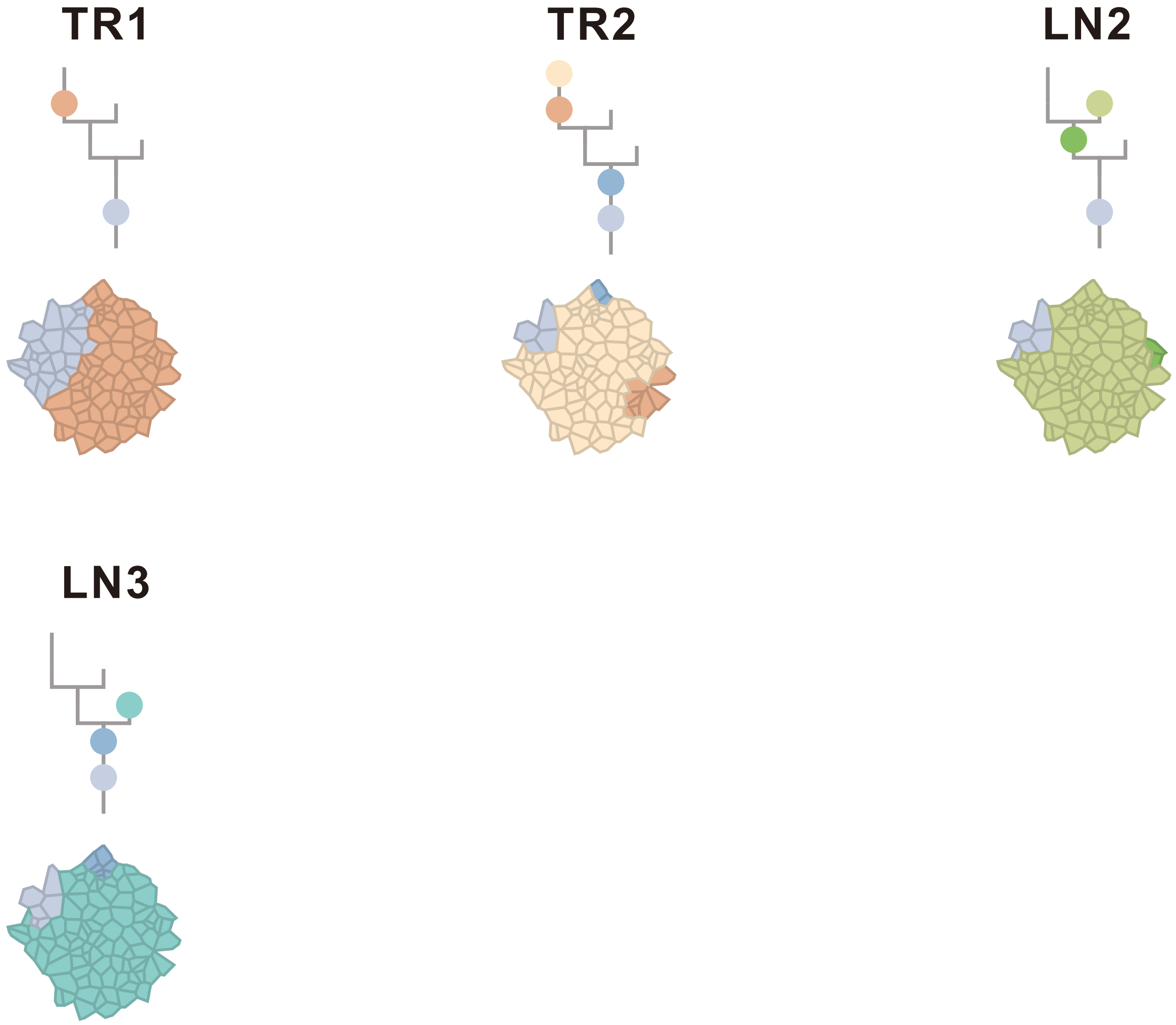

### CRC02

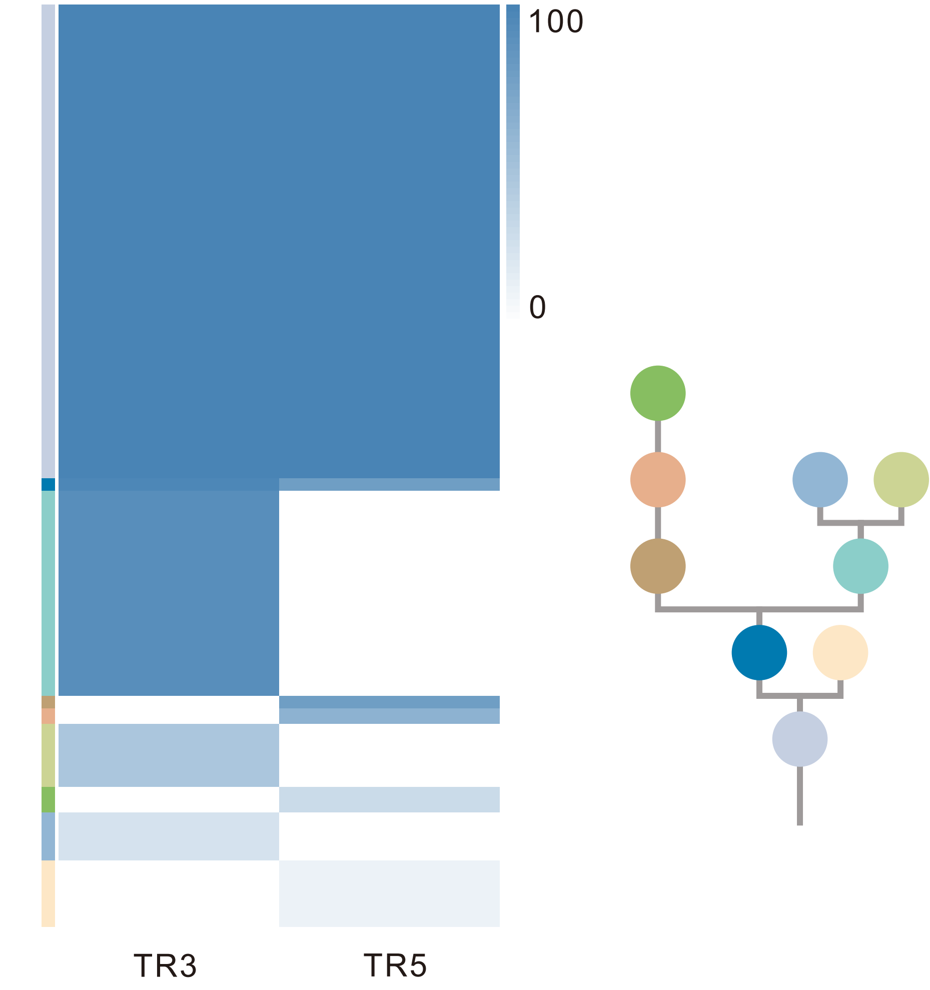

| Gene | Cytoband | Type | Cluster |
|---|---|---|---|
| * *APC* | 5q22.2 | SNV | 1 |
| *CUX1* | 7q22.1 | SNV | 1 |
| *TP53* | 17p13.1 | SNV | 1 |
| * *APC* | 5q22.2 | InDel | 1 |
| *FGFR1* | 8p11.23 | Amp | 1 |
| *CDKN2A* | 9p21.3 | Del | 1 |
| *AMER1* | Xq11.2 | SNV | NA |
| *ATRX* | Xq21.1 | SNV | NA |

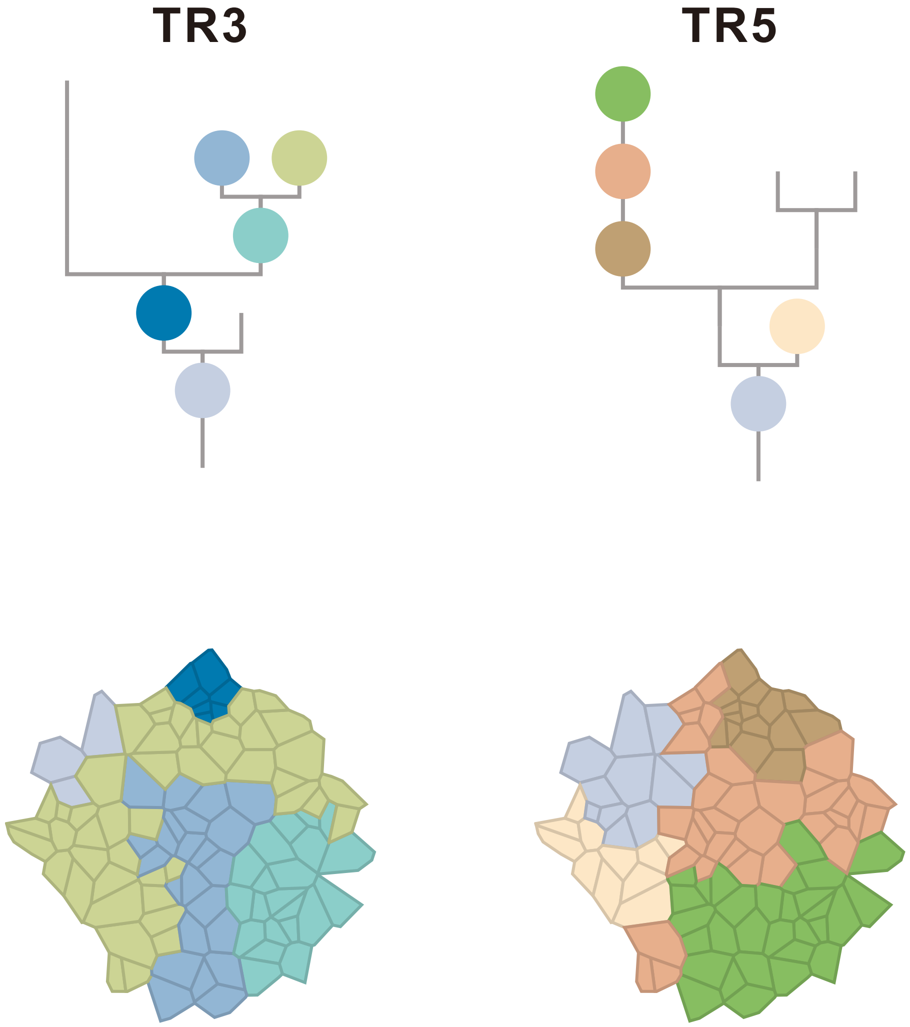

### CRC03

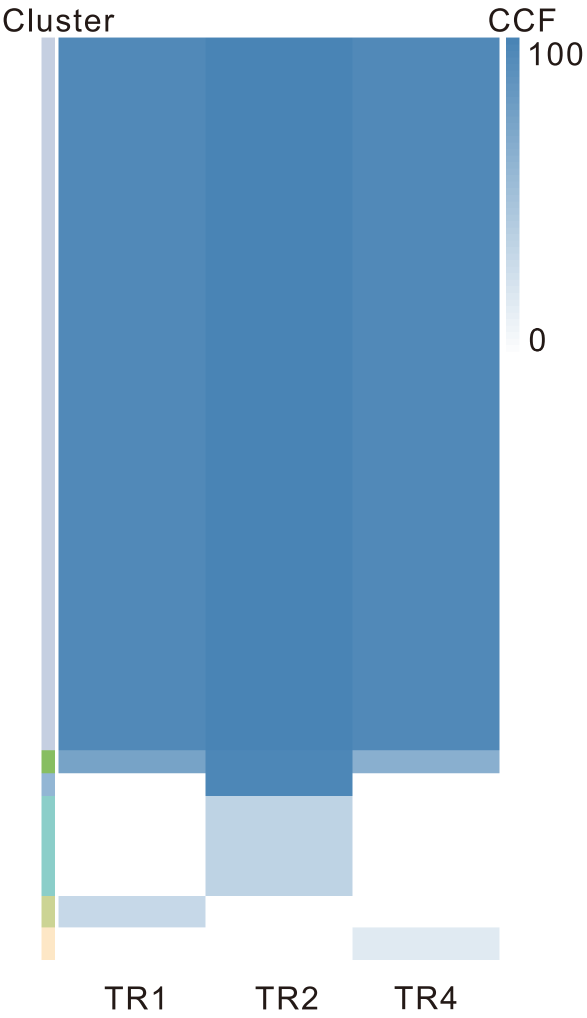

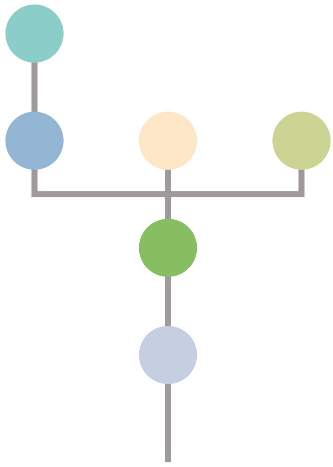

| Gene | Cytoband | Type | Cluster |
|---|---|---|---|
| *APC* | 5q22.2 | SNV | 1 |
| *KRAS* | 12p12.1 | SNV | 1 |
| *ETV6* | 12p13.2 | SNV | 1 |
| *CSF3R* | 1p34.3 | InDel | 6 |
| *AMER1* | Xq11.2 | SNV | NA |
| *PTEN* | 10q23.31 | Del | NA |

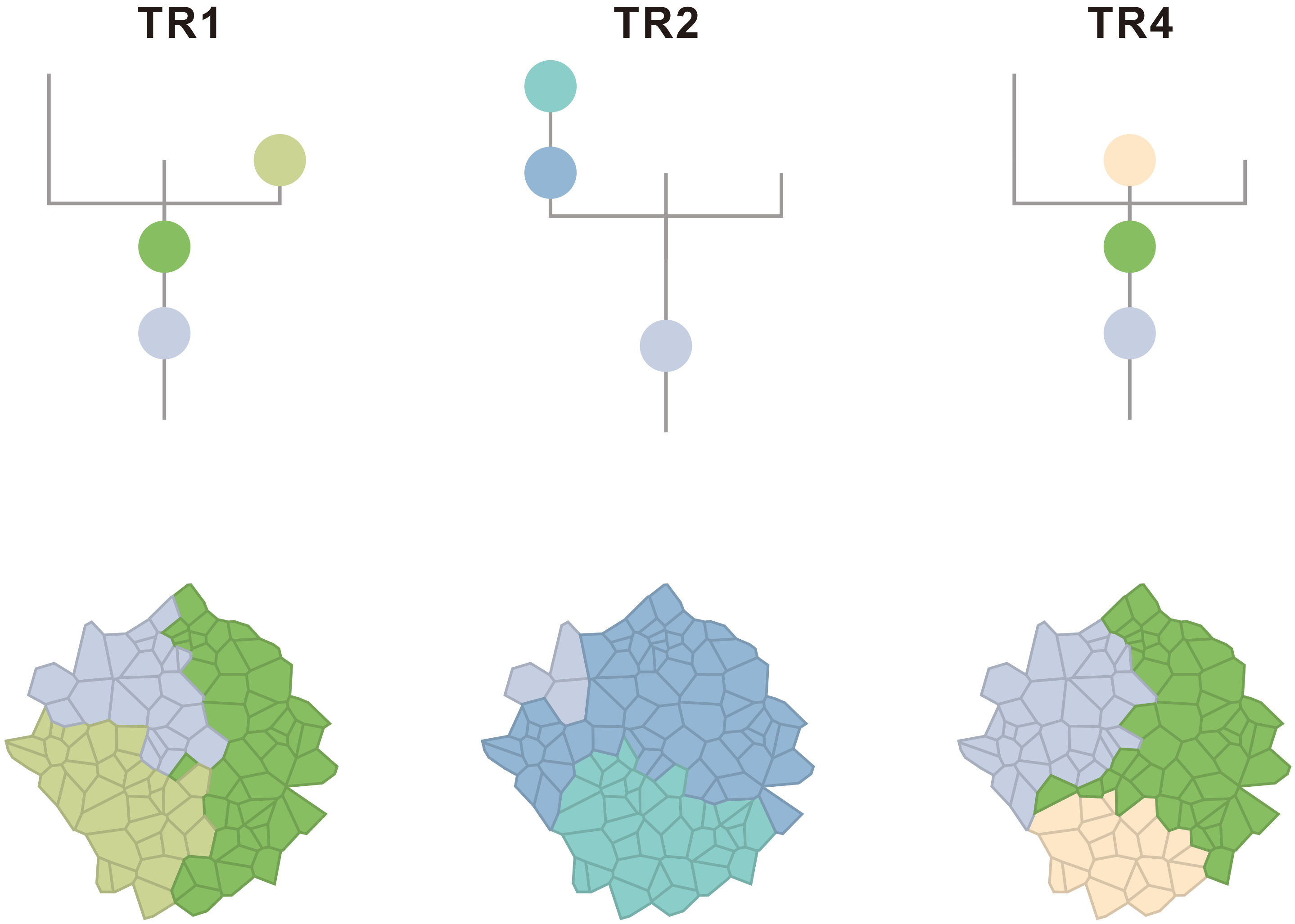

### CRC06

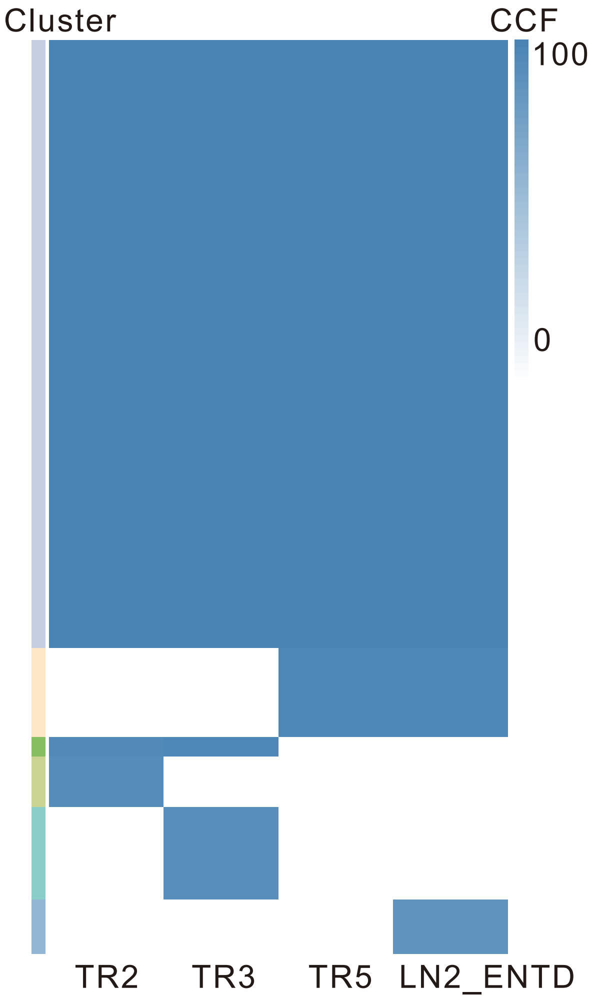

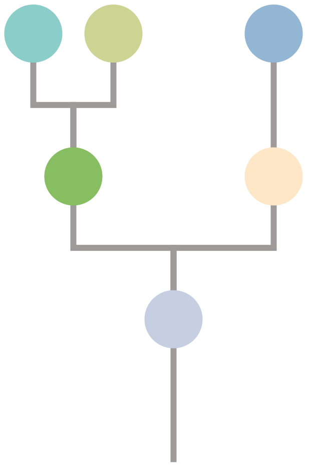

| Gene | Cytoband | Type | Cluster |
|---|---|---|---|
| * *PIK3CA* | 3q26.32 | SNV | 1 |
| * *PIK3CA* | 3q26.32 | SNV | 1 |
| *KRAS* | 12p12.1 | SNV | 1 |
| *ERCC2* | 19q13.32 | SNV | 1 |
| *APC* | 5q22.2 | InDel | 1 |
| *SSX2* | Xp11.22 | Amp | 1 |
| *ARAF* | Xp11.23 | Amp | 1 |
| *GATA1* | Xp11.23 | Amp | 1 |
| *SSX1* | Xp11.23 | Amp | 1 |
| *SSX4* | Xp11.23 | Amp | 1 |
| *TFE3* | Xp11.23 | Amp | 1 |
| *WAS* | Xp11.23 | Amp | 1 |
| *KDM6A* | Xp11.3 | Amp | 1 |
| *FOXO4* | Xq13.1 | Amp | 1 |
| *BTK* | Xq22.1 | Amp | 1 |
| *IRS4* | Xq22.3 | Amp | 1 |
| *BCORL1* | Xq26.1 | Amp | 1 |
| *ELF4* | Xq26.1 | Amp | 1 |
| *GPC3* | Xq26.2 | Amp | 1 |
| *CAMTA1* | 1p36.31 | Del | 1 |
| *RPL22* | 1p36.31 | Del | 1 |
| *TNFRSF14* | 1p36.32 | Del | 1 |
| *NOTCH1* | 9q34.3 | Del | 1 |
| *AXIN1* | 16p13.3 | Del | 1 |
| *CREBBP* | 16p13.3 | Del | 1 |
| *NTHL1* | 16p13.3 | Del | 1 |
| *TRAF7* | 16p13.3 | Del | 1 |
| *TSC2* | 16p13.3 | Del | 1 |
| *FLCN* | 17p11.2 | Del | 1 |
| *NCOR1* | 17p11.2 | Del | 1 |
| *MAP2K4* | 17p12 | Del | 1 |
| *PER1* | 17p13.1 | Del | 1 |
| *TP53* | 17p13.1 | Del | 1 |
| *YWHAE* | 17p13.3 | Del | 1 |
| *SMAD4* | 18q21.2 | InDel | 3 |
| *CARS* | 11p15.4 | Del | 3 |
| *NOTCH2* | 1p12 | Amp | 4 |
| *ARNT* | 1q21.3 | Amp | 4 |
| *SETDB1* | 1q21.3 | Amp | 4 |
| *STK11* | 19p13.3 | Del | 5 |
| *TCF3* | 19p13.3 | Del | 5 |
| *SDHA* | 5p15.33 | Del | 5 |
| *TERT* | 5p15.33 | Del | 5 |
| *TP53* | 17p13.1 | SNV | NA |

**TR2**

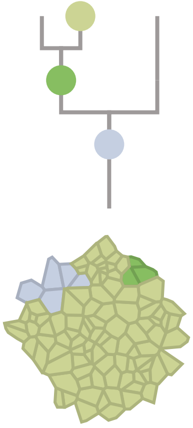

**TR3**

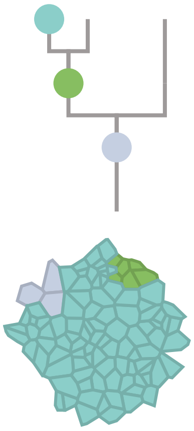

**TR5**

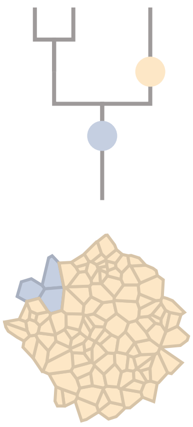

**LN2_ENTD**

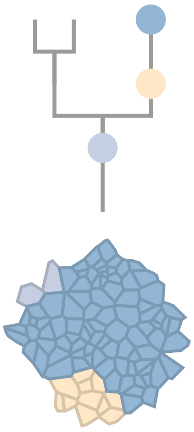

### CRC07

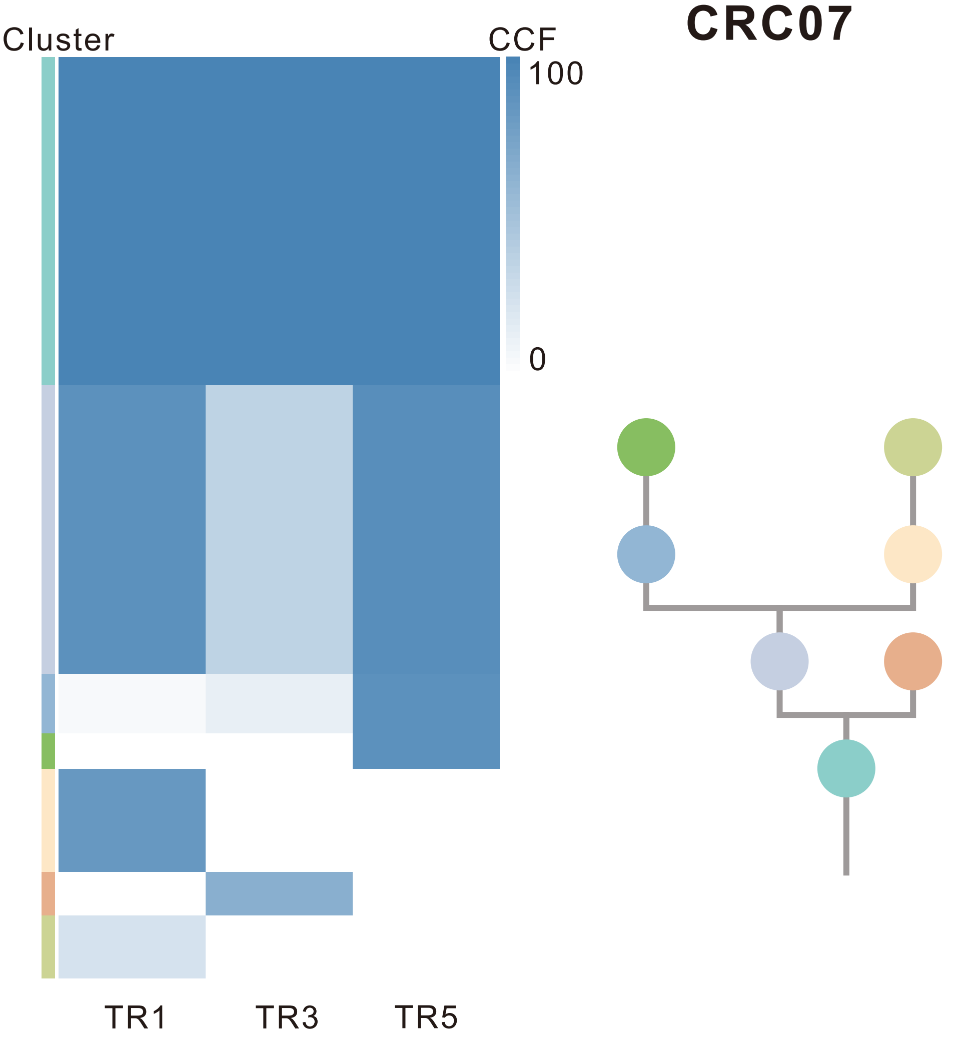

| Gene | Cytoband | Type | Cluster |
|---|---|---|---|
| * *APC* | 5q22.2 | SNV | 1 |
| *TBL1XR1* | 3q26.32 | SNV | 2 |
| *KRAS* | 12p12.1 | SNV | 2 |
| *TP53* | 17p13.1 | SNV | 2 |
| * *APC* | 5q22.2 | InDel | 2 |
| *RUNX1* | 21q22.12 | Del | 2 |
| *CLTCL1* | 22q11.21 | Del | 2 |
| *LZTR1* | 22q11.21 | Del | 2 |
| *SMARCB1* | 22q11.23 | Del | 2 |
| *CHEK2* | 22q12.1 | Del | 2 |
| *ZNRF3* | 22q12.1 | Del | 2 |
| *NF2* | 22q12.2 | Del | 2 |
| *MYH9* | 22q12.3 | Del | 2 |
| *APOBEC3B* | 22q13.1 | Del | 2 |
| *EP300* | 22q13.2 | Del | 2 |
| *MKL1* | 22q13.2 | Del | 2 |
| *FOXO4* | Xq13.1 | Del | 2 |
| *MED12* | Xq13.1 | Del | 2 |
| *ZMYM3* | Xq13.1 | Del | 2 |
| *ATRX* | Xq21.1 | Del | 2 |
| *BTK* | Xq22.1 | Del | 2 |
| *IRS4* | Xq22.3 | Del | 2 |
| *STAG2* | Xq25 | Del | 2 |
| *BCORL1* | Xq26.1 | Del | 2 |
| *ELF4* | Xq26.1 | Del | 2 |
| *GPC3* | Xq26.2 | Del | 2 |
| *PHF6* | Xq26.2 | Del | 2 |
| *ATP2B3* | Xq28 | Del | 2 |
| *RPL10* | Xq28 | Del | 2 |
| *ERBB4* | 2q34 | SNV | 3 |
| *RNF43* | 17q22 | SNV | 4 |
| *ATP1A1* | 1p13.1 | Amp | 5 |
| *MUC4* | 3q29 | Amp | 5 |
| *SIRPA* | 20p13 | Del | 7 |
| *ASXL1* | 20q11.21 | Del | 7 |
| *PTPRT* | 20q13.11 | Del | 7 |
| *PTK6* | 20q13.33 | Del | 7 |

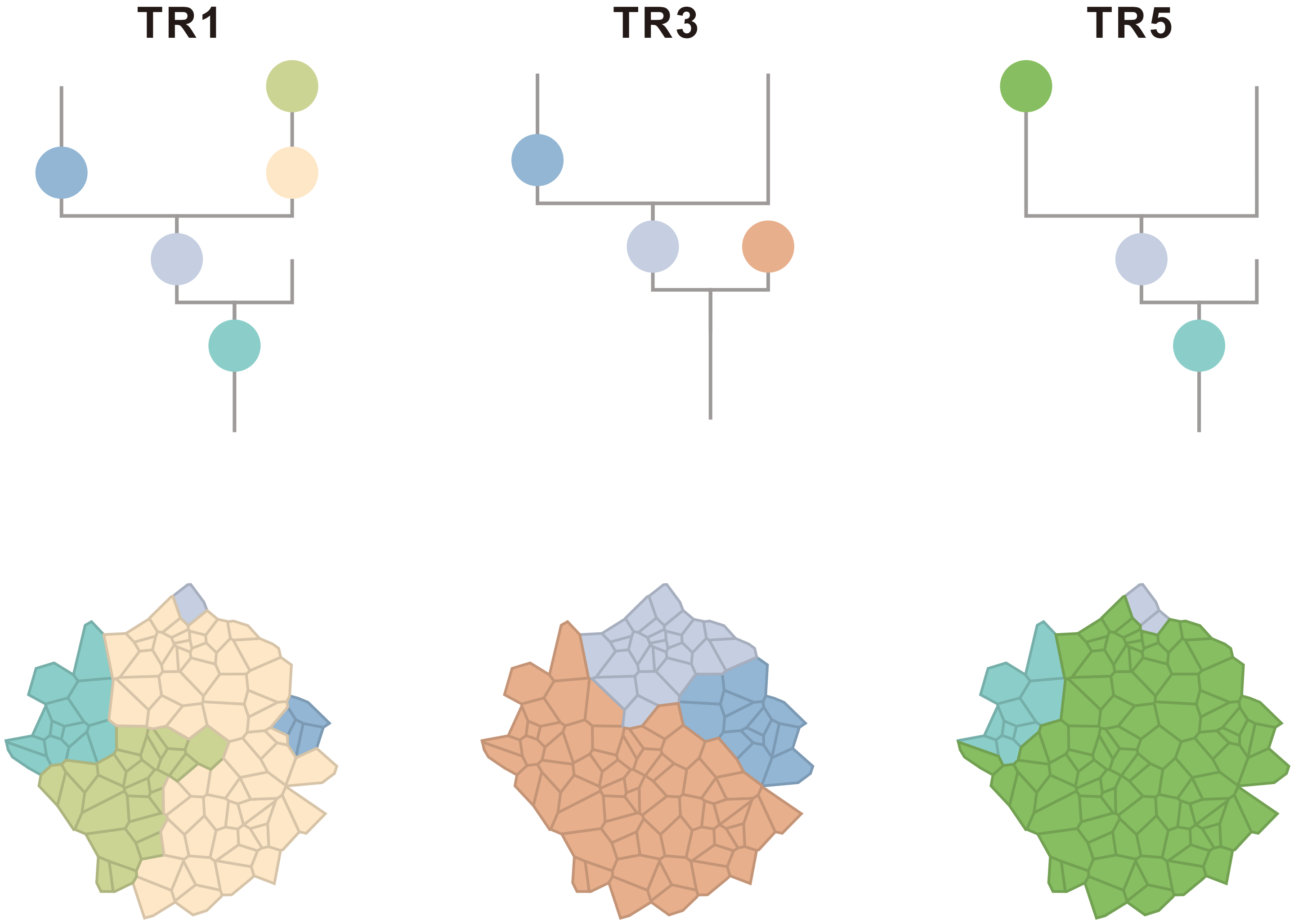

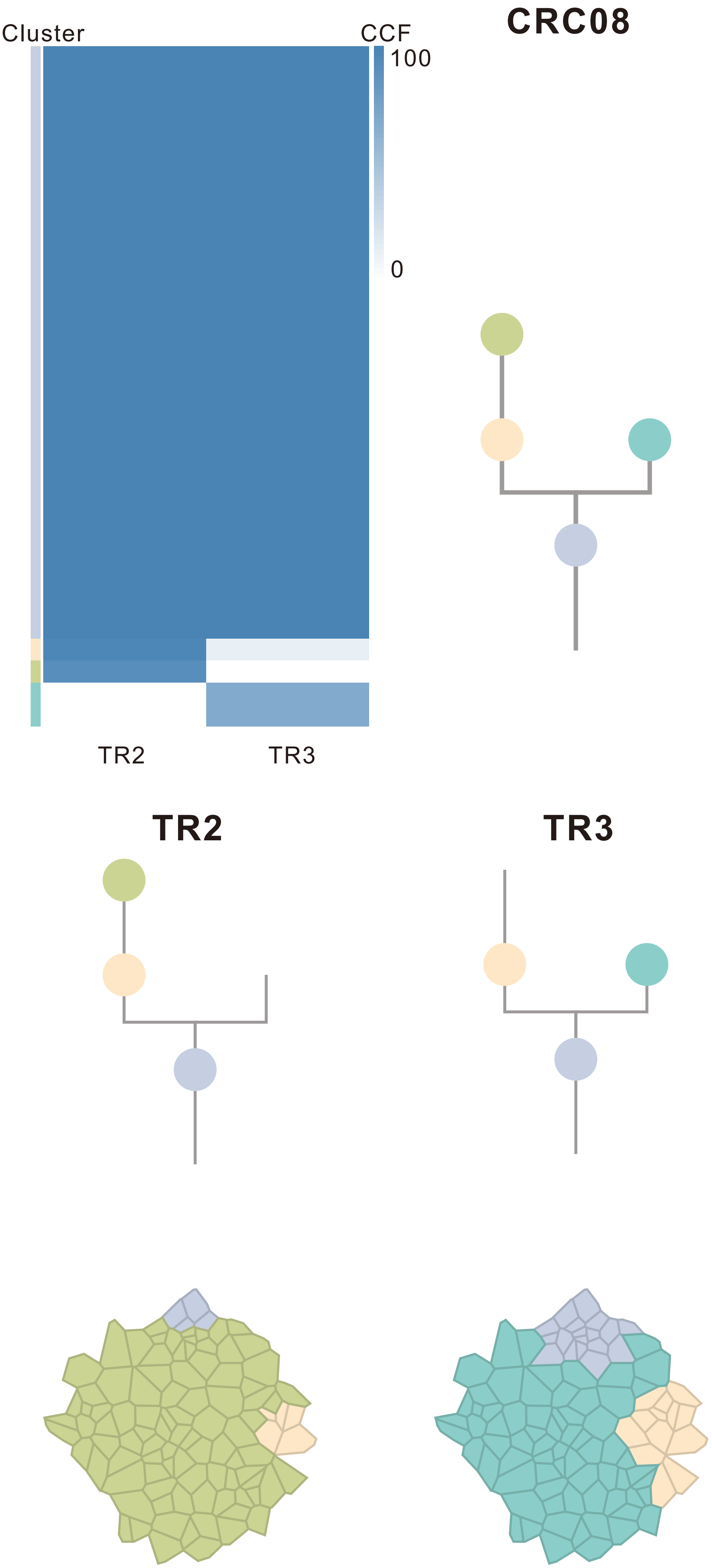

| Gene | Cytoband | Type | Cluster |
|---|---|---|---|
| *ESR1* | 6q25.1 | SNV | 1 |
| *ARHGEF10* | 8p23.3 | SNV | 1 |
| *PREX2* | 8q13.2 | SNV | 1 |
| *CSMD3* | 8q23.3 | SNV | 1 |
| *BCL9* | 1q21.2 | Amp | 1 |
| *ARNT* | 1q21.3 | Amp | 1 |
| *SETDB1* | 1q21.3 | Amp | 1 |
| *POU5F1* | 6p21.33 | Amp | 1 |
| *TRIM27* | 6p22.1 | Amp | 1 |
| *HIST1H3B* | 6p22.2 | Amp | 1 |
| *DEK* | 6p22.3 | Amp | 1 |
| *IRF4* | 6p25.3 | Amp | 1 |
| *CUX1* | 7q22.1 | Amp | 1 |
| *FLT3* | 13q12.2 | Amp | 1 |
| *FOXO1* | 13q14.11 | Amp | 1 |
| *CYSLTR2* | 13q14.2 | Amp | 1 |
| *SSX2* | Xp11.22 | Amp | 1 |
| *ARAF* | Xp11.23 | Amp | 1 |
| *GATA1* | Xp11.23 | Amp | 1 |
| *SSX1* | Xp11.23 | Amp | 1 |
| *SSX4* | Xp11.23 | Amp | 1 |
| *TFE3* | Xp11.23 | Amp | 1 |
| *WAS* | Xp11.23 | Amp | 1 |
| *KDM6A* | Xp11.3 | Amp | 1 |
| *BTK* | Xq22.1 | Amp | 1 |
| *IRS4* | Xq22.3 | Amp | 1 |
| *BCORL1* | Xq26.1 | Amp | 1 |
| *ELF4* | Xq26.1 | Amp | 1 |
| *GPC3* | Xq26.2 | Amp | 1 |
| *CASP9* | 1p36.21 | Del | 1 |
| *PRDM2* | 1p36.21 | Del | 1 |
| *SPEN* | 1p36.21 | Del | 1 |
| *CAMTA1* | 1p36.31 | Del | 1 |
| *RPL22* | 1p36.31 | Del | 1 |
| *TNFRSF14* | 1p36.32 | Del | 1 |
| *DAXX* | 6p21.32 | Del | 1 |
| *NDRG1* | 8q24.22 | Del | 1 |
| *RECQL4* | 8q24.3 | Del | 1 |
| *FEN1* | 11q12.2 | Del | 1 |
| *SDHAF2* | 11q12.2 | Del | 1 |
| *MEN1* | 11q13.1 | Del | 1 |
| *FLCN* | 17p11.2 | Del | 1 |
| *NCOR1* | 17p11.2 | Del | 1 |
| *MAP2K4* | 17p12 | Del | 1 |
| *PER1* | 17p13.1 | Del | 1 |
| *TP53* | 17p13.1 | Del | 1 |
| *YWHAE* | 17p13.3 | Del | 1 |
| *ELL* | 19p13.11 | Del | 1 |
| *DNM2* | 19p13.2 | Del | 1 |
| *KEAP1* | 19p13.2 | Del | 1 |
| *SMARCA4* | 19p13.2 | Del | 1 |
| *STK11* | 19p13.3 | Del | 1 |
| *TCF3* | 19p13.3 | Del | 1 |
| *CLTCL1* | 22q11.21 | Del | 1 |
| *LZTR1* | 22q11.21 | Del | 1 |
| *SMARCB1* | 22q11.23 | Del | 1 |
| *CHEK2* | 22q12.1 | Del | 1 |
| *ZNRF3* | 22q12.1 | Del | 1 |
| *NF2* | 22q12.2 | Del | 1 |
| *MYH9* | 22q12.3 | Del | 1 |
| *APOBEC3B* | 22q13.1 | Del | 1 |
| *EP300* | 22q13.2 | Del | 1 |
| *MKL1* | 22q13.2 | Del | 1 |
| *MUC4* | 3q29 | Amp | 2 |
| *NOTCH1* | 9q34.3 | Del | 2 |
| *TBX3* | 12q24.21 | Del | 2 |
| *HNF1A* | 12q24.31 | Del | 2 |
| *NCOR2* | 12q24.31 | Del | 2 |
| *SETD1B* | 12q24.31 | Del | 2 |
| *POLE* | 12q24.33 | Del | 2 |
| *TP53* | 17p13.1 | SNV | NA |

### CRC10

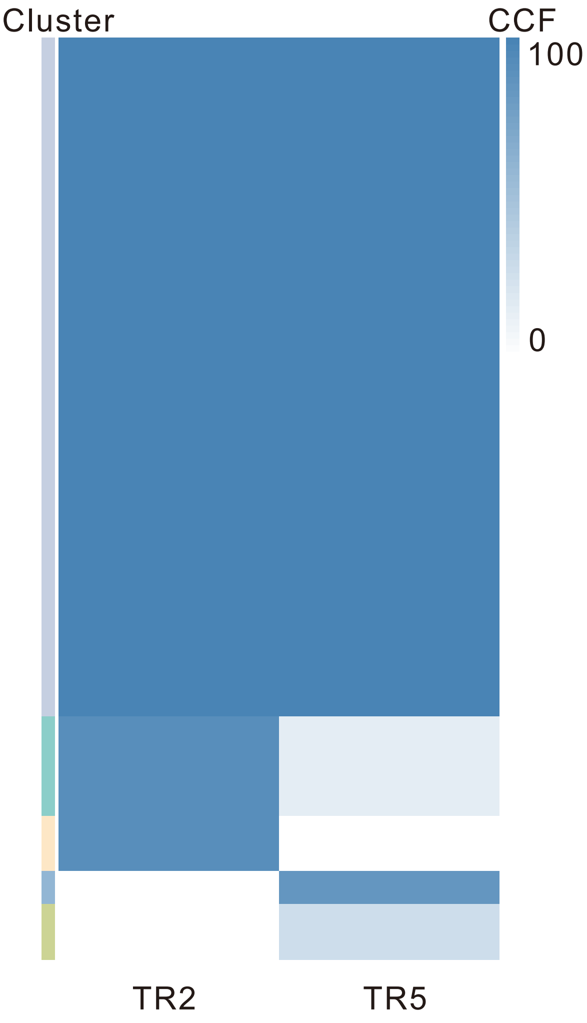

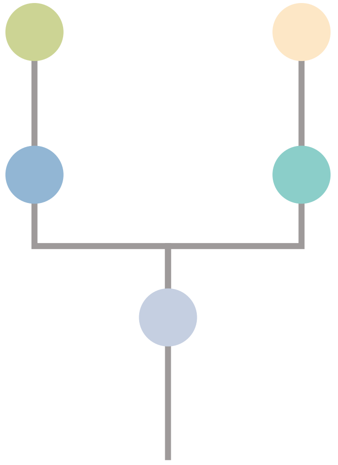

| Gene | Cytoband | Type | Cluster |
| --- | --- | --- | --- |
| *KRAS* | 12p12.1 | SNV | 1 |
| *APC* | 5q22.2 | InDel | 1 |
| *FLT3* | 13q12.2 | Amp | 1 |
| *FOXO1* | 13q14.11 | Amp | 1 |
| *CYSLTR2* | 13q14.2 | Amp | 1 |
| *CAMTA1* | 1p36.31 | Del | 1 |
| *RPL22* | 1p36.31 | Del | 1 |
| *TNFRSF14* | 1p36.32 | Del | 1 |
| *FLCN* | 17p11.2 | Del | 1 |
| *NCOR1* | 17p11.2 | Del | 1 |
| *MAP2K4* | 17p12 | Del | 1 |
| *PER1* | 17p13.1 | Del | 1 |
| *TP53* | 17p13.1 | Del | 1 |
| *YWHAE* | 17p13.3 | Del | 1 |
| *TSC1* | 9q34.13 | Del | 2 |
| *NOTCH1* | 9q34.3 | Del | 2 |
| *AXIN1* | 16p13.3 | Del | 2 |
| *STK11* | 19p13.3 | Del | 2 |
| *TCF3* | 19p13.3 | Del | 2 |

## TR2

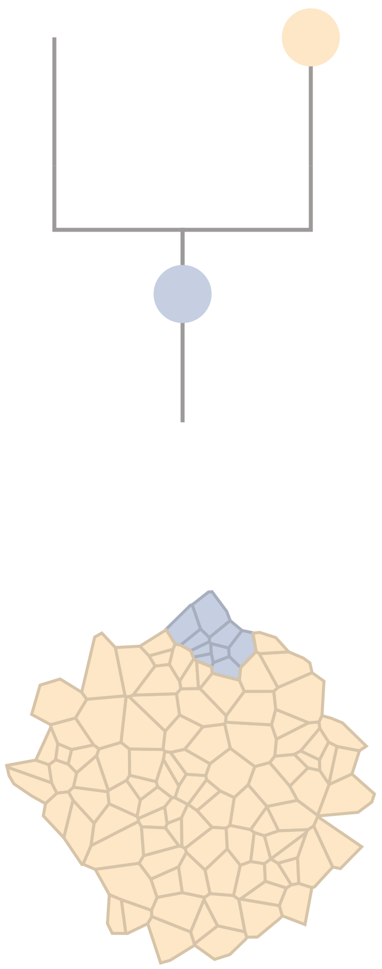

## TR5

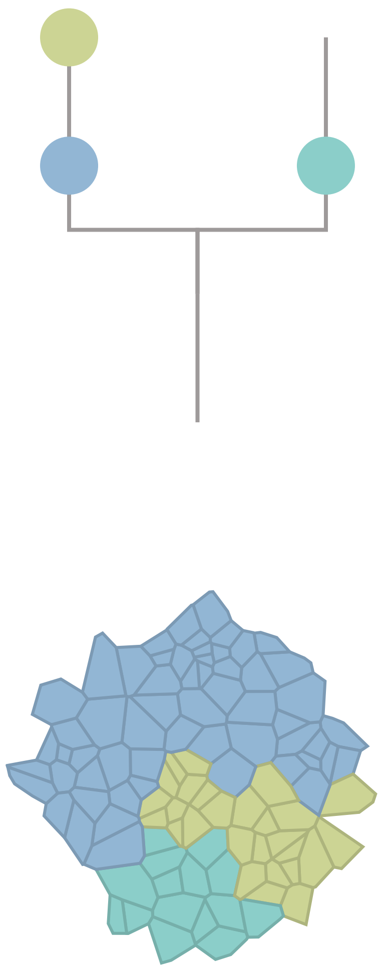

### CRC11

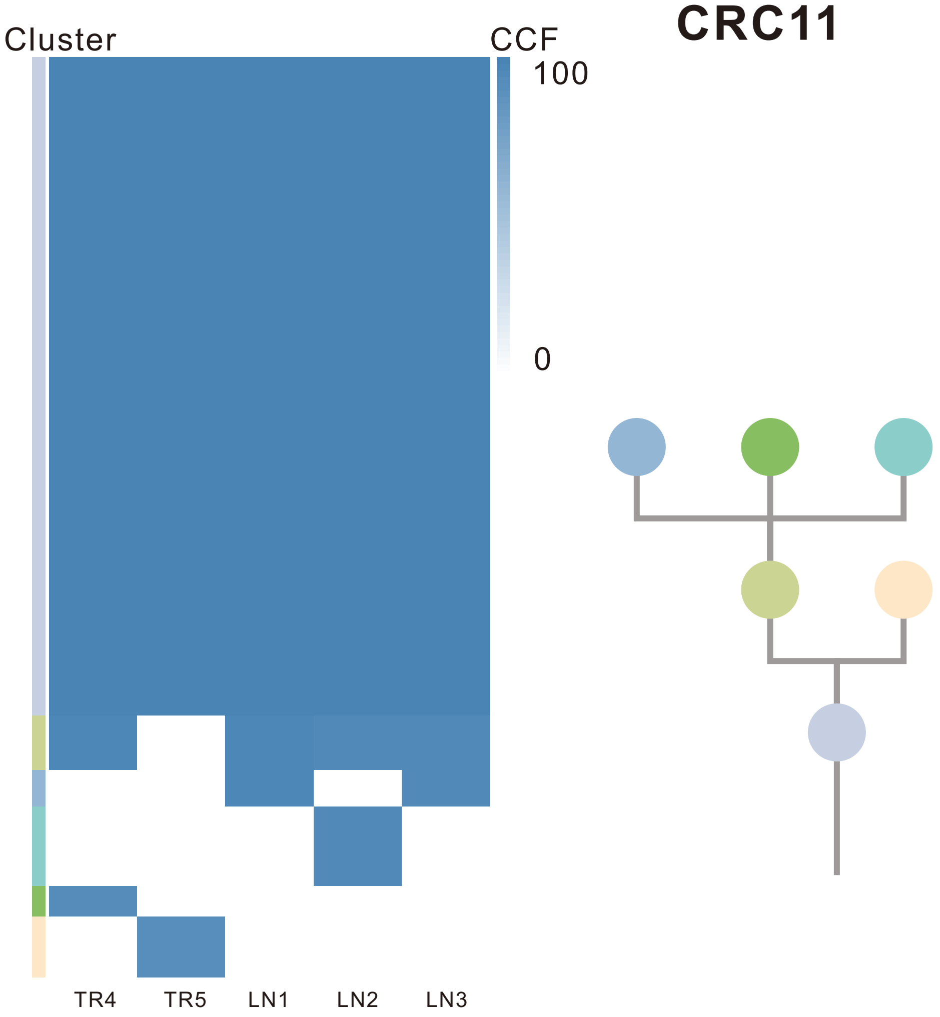

| Gene | Cytoband | Type | Cluster |
|---|---|---|---|
| * *APC* | 5q22.2 | SNV | 1 |
| *BRAF* | 7q34 | SNV | 1 |
| *ZFHX3* | 16q22.2 | SNV | 1 |
| * *SMAD4* | 18q21.2 | SNV | 1 |
| * *SMAD4* | 18q21.2 | SNV | 1 |
| * *APC* | 5q22.2 | InDel | 1 |
| *PAX5* | 9p13.2 | Amp | 1 |
| *PSIP1* | 9p22.3 | Amp | 1 |
| *JAK2* | 9p24.1 | Amp | 1 |
| *PDCD1LG2* | 9p24.1 | Amp | 1 |
| *GNAQ* | 9q21.2 | Amp | 1 |
| *SYK* | 9q22.2 | Amp | 1 |
| *NR4A3* | 9q31.1 | Amp | 1 |
| *KLF4* | 9q31.2 | Amp | 1 |
| *TAL2* | 9q31.2 | Amp | 1 |
| *TNC* | 9q33.1 | Amp | 1 |
| *SET* | 9q34.11 | Amp | 1 |
| *ABL1* | 9q34.12 | Amp | 1 |
| *BRD3* | 9q34.2 | Amp | 1 |
| *NOTCH1* | 9q34.3 | Amp | 1 |
| *FLT3* | 13q12.2 | Amp | 2 |
| *FOXO1* | 13q14.11 | Amp | 2 |
| *CYSLTR2* | 13q14.2 | Amp | 2 |
| *SDHA* | 5p15.33 | Del | 3 |
| *MUC4* | 3q29 | Amp | NA |
| *PLAG1* | 8q12.1 | Amp | NA |
| *PREX2* | 8q13.2 | Amp | NA |
| *NCOA2* | 8q13.3 | Amp | NA |
| *HEY1* | 8q21.13 | Amp | NA |
| *RUNX1T1* | 8q21.3 | Amp | NA |
| *CDH17* | 8q22.1 | Amp | NA |
| *PABPC1* | 8q22.3 | Amp | NA |
| *UBR5* | 8q22.3 | Amp | NA |
| *RAD21* | 8q24.11 | Amp | NA |
| *AXIN1* | 16p13.3 | Del | NA |

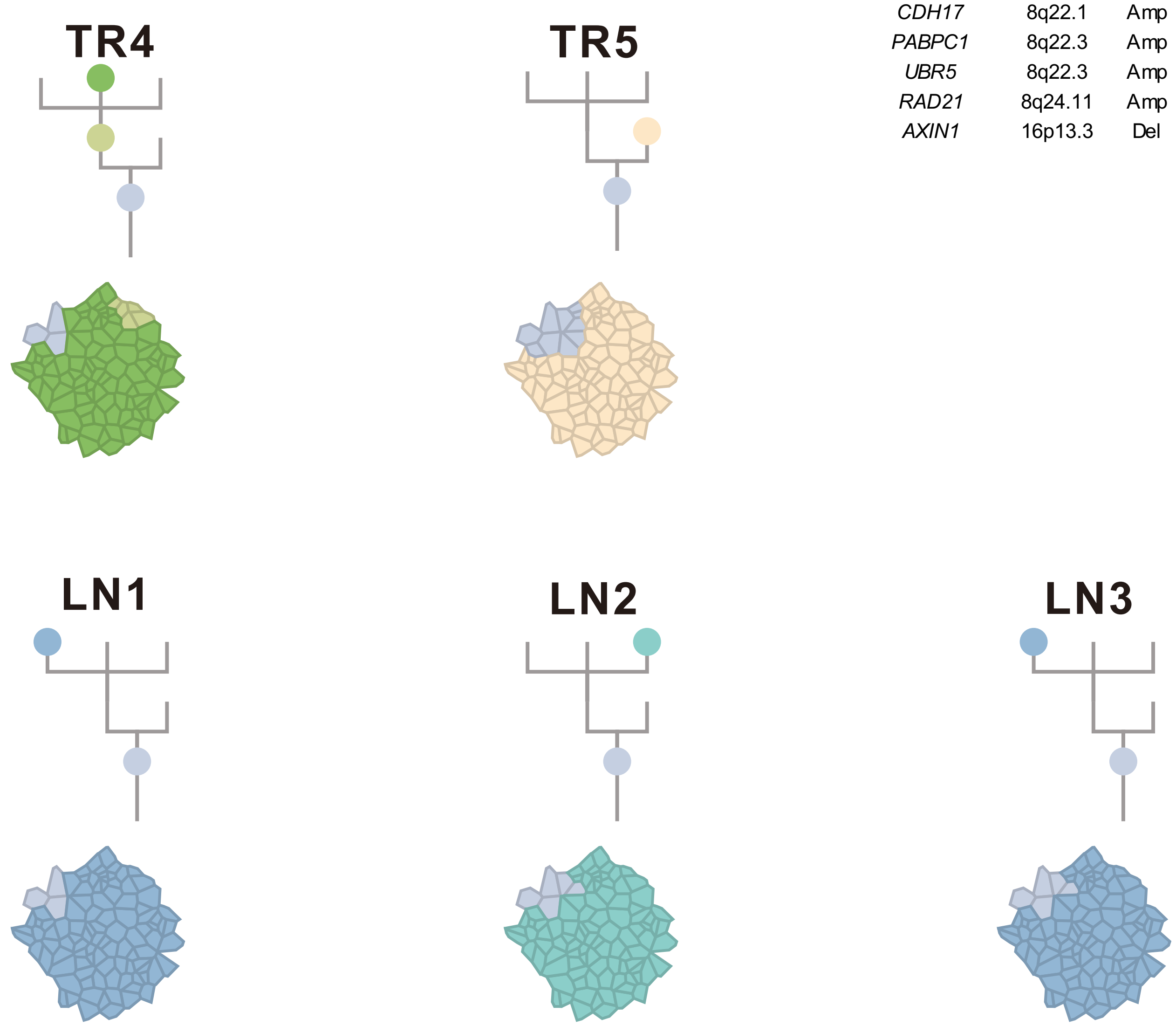

### CRC12

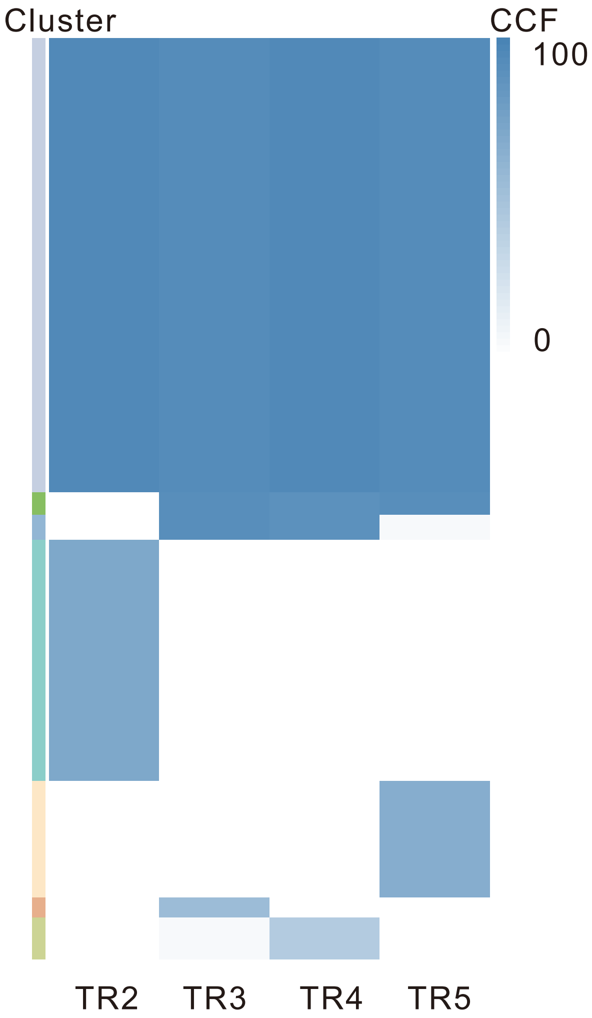

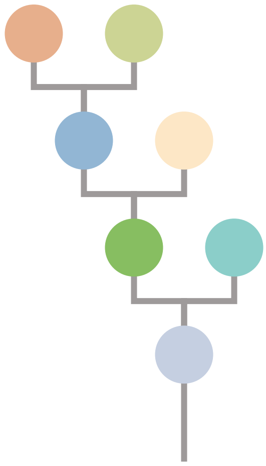

| Gene | Cytoband | Type | Cluster |
|---|---|---|---|
| *IDH1* | 2q34 | SNV | 1 |
| *BRAF* | 7q34 | SNV | 1 |
| *ARID2* | 12q12 | SNV | 1 |
| *RNF43* | 17q22 | SNV | 1 |
| *SMAD4* | 18q21.2 | SNV | 1 |
| *TP53* | 17p13.1 | InDel | 1 |
| *CREB3L2* | 7q33 | Amp | 1 |
| *TRIM24* | 7q33 | Amp | 1 |
| *BRAF* | 7q34 | Amp | 1 |
| *EZH2* | 7q36.1 | Amp | 1 |
| *RUNX1T1* | 8q21.3 | Amp | 1 |
| *CDH17* | 8q22.1 | Amp | 1 |
| *PABPC1* | 8q22.3 | Amp | 1 |
| *UBR5* | 8q22.3 | Amp | 1 |
| *RAD21* | 8q24.11 | Amp | 1 |
| *MYC* | 8q24.21 | Amp | 1 |
| *RECQL4* | 8q24.3 | Amp | 1 |
| *FLT3* | 13q12.2 | Amp | 1 |
| *FOXO1* | 13q14.11 | Amp | 1 |
| *CYSLTR2* | 13q14.2 | Amp | 1 |
| *ELK4* | 1q32.1 | Amp | 2 |
| *MDM4* | 1q32.1 | Amp | 2 |
| *ROBO2* | 3p12.3 | SNV | 3 |
| *FAT4* | 4q28.1 | SNV | 4 |
| *PRDM16* | 1p36.32 | Amp | NA |
| *ARNT* | 1q21.3 | Amp | NA |
| *SETDB1* | 1q21.3 | Amp | NA |
| *FCRL4* | 1q23.1 | Amp | NA |
| *NTRK1* | 1q23.1 | Amp | NA |
| *DDR2* | 1q23.3 | Amp | NA |
| *FCGR2B* | 1q23.3 | Amp | NA |
| *PBX1* | 1q23.3 | Amp | NA |

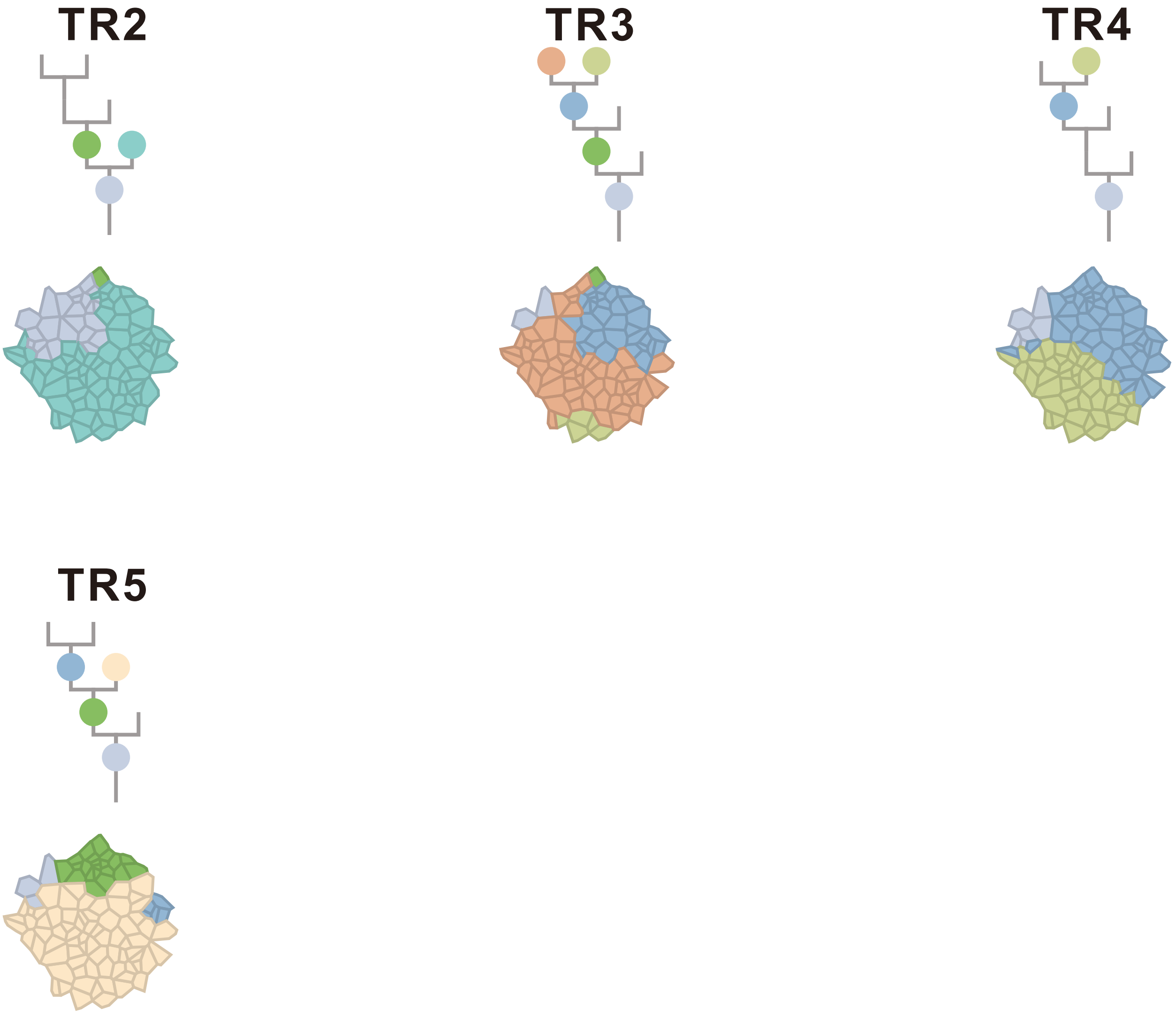

### CRC14

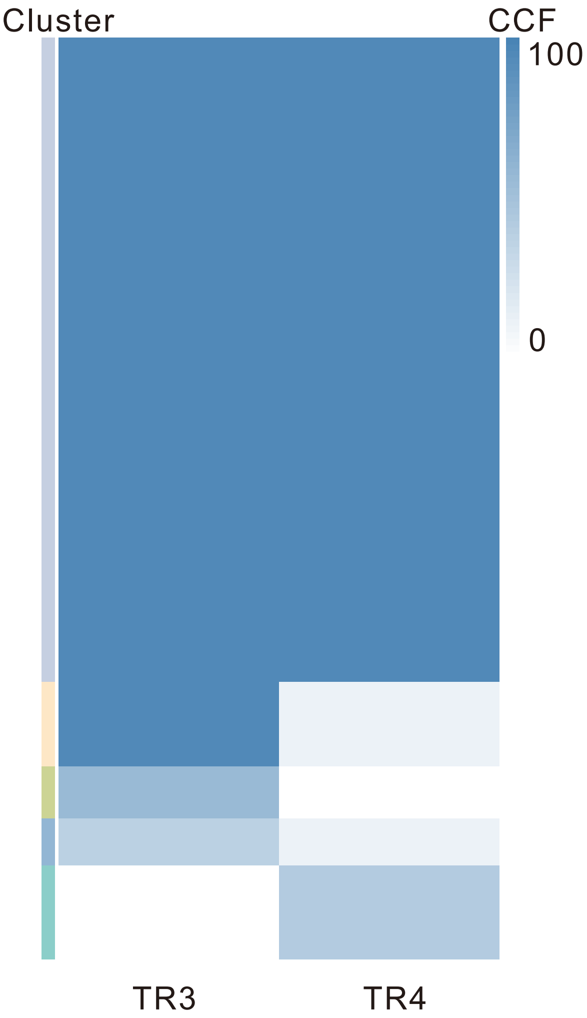

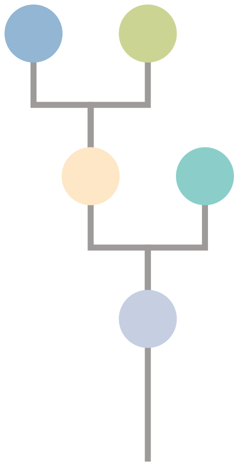

| Gene | Cytoband | Type | Cluster |
|---|---|---|---|
| *SDHB* | 1p36.13 | SNV | 1 |
| *TP63* | 3q28 | SNV | 1 |
| *APC* | 5q22.2 | SNV | 1 |
| *ESR1* | 6q25.1 | SNV | 1 |
| *DICER1* | 14q32.13 | SNV | 1 |
| *TP53* | 17p13.1 | SNV | 1 |
| *IKBKB* | 8p11.21 | Amp | 1 |
| *KAT6A* | 8p11.21 | Amp | 1 |
| *FGFR1* | 8p11.23 | Amp | 1 |
| *SRC* | 20q11.23 | Amp | 1 |
| *MAFB* | 20q12 | Amp | 1 |
| *PLCG1* | 20q12 | Amp | 1 |
| *NFATC2* | 20q13.2 | Amp | 1 |
| *SALL4* | 20q13.2 | Amp | 1 |
| *GNAS* | 20q13.32 | Amp | 1 |
| *PTK6* | 20q13.33 | Amp | 1 |

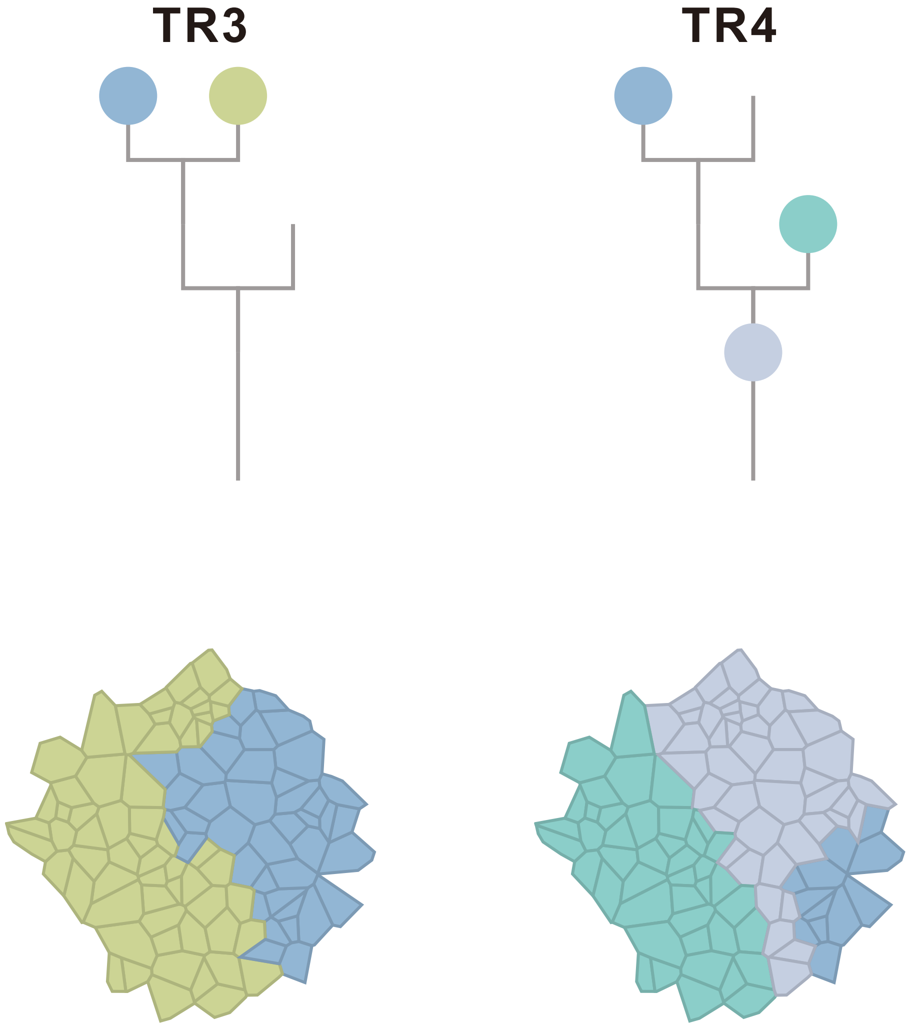

### CRC16

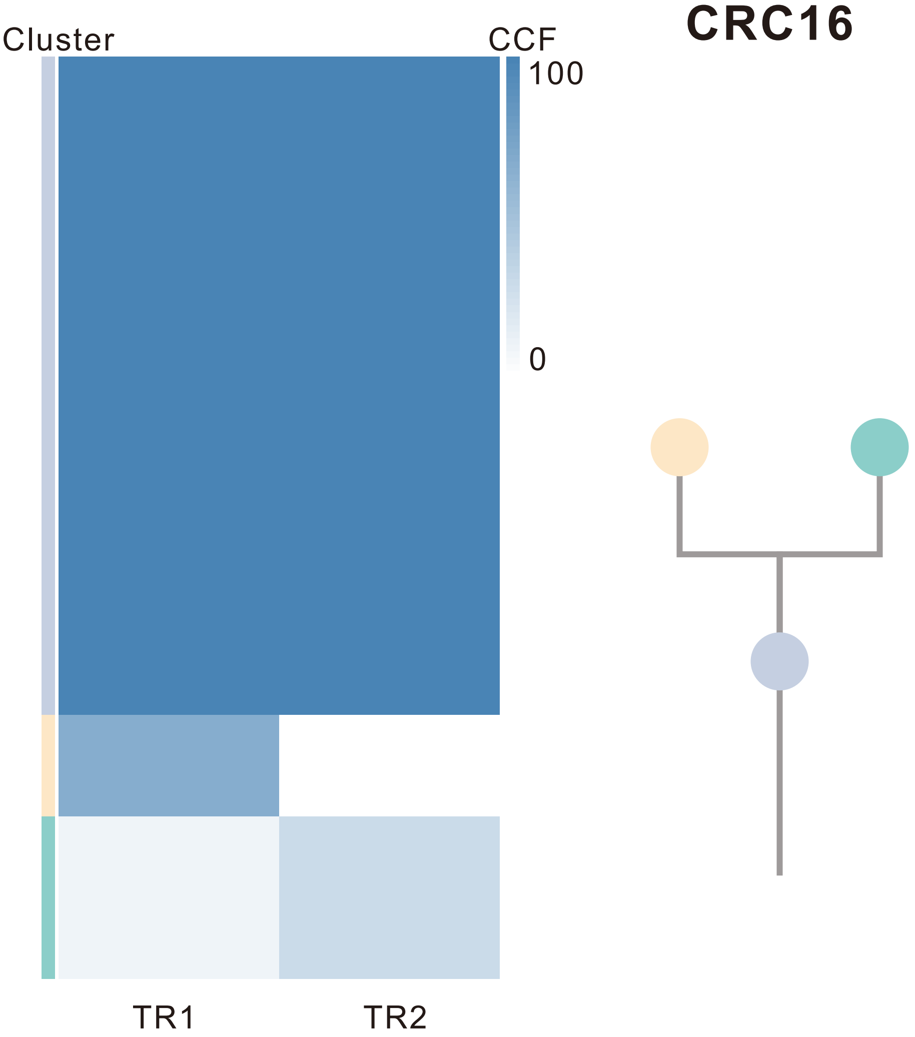

| Gene | Cytoband | Type | Cluster |
|---|---|---|---|
| *MITF* | 3p13 | SNV | 1 |
| *CTNNB1* | 3p22.1 | SNV | 1 |
| *PIK3CA* | 3q26.32 | SNV | 1 |
| *APC* | 5q22.2 | SNV | 1 |
| *KRAS* | 12p12.1 | SNV | 1 |
| *RSPO2* | 8q23.1 | SNV | 2 |

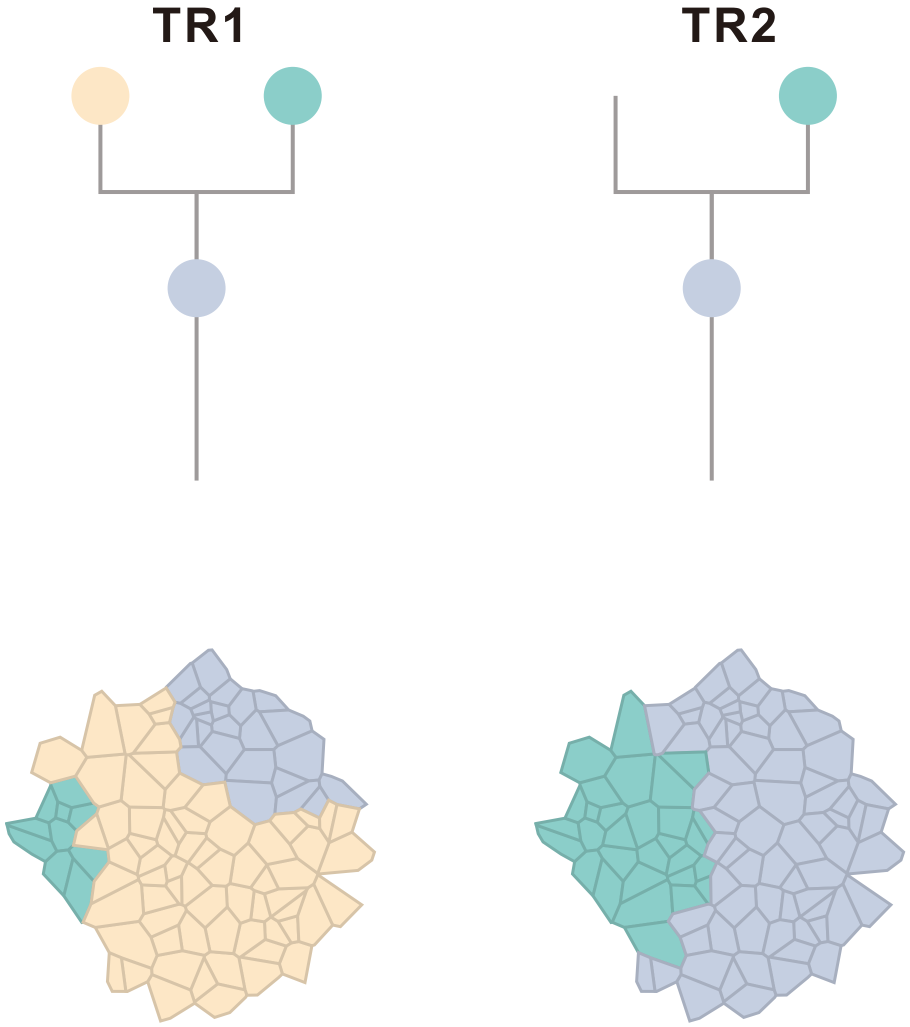

### CRC18

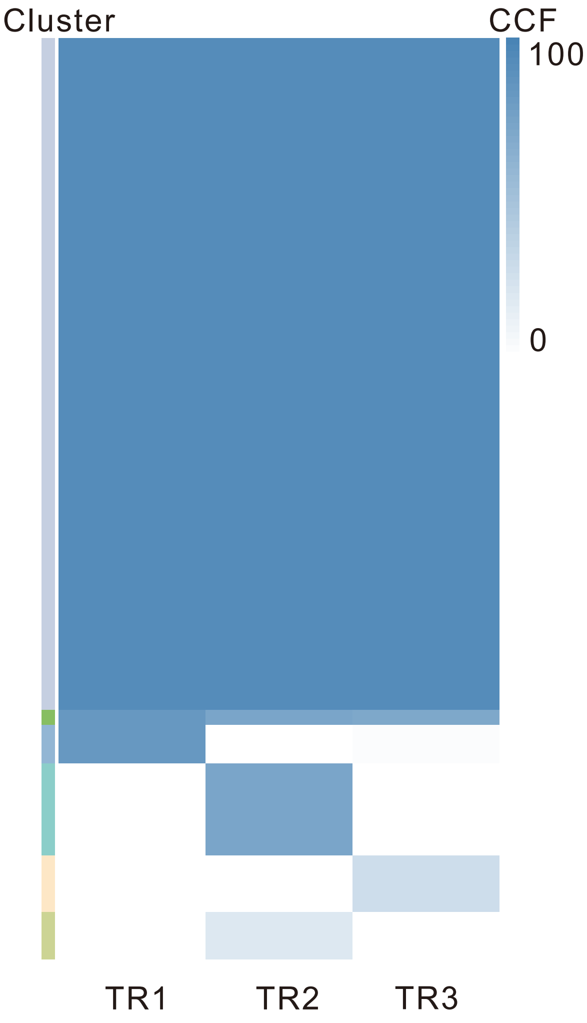

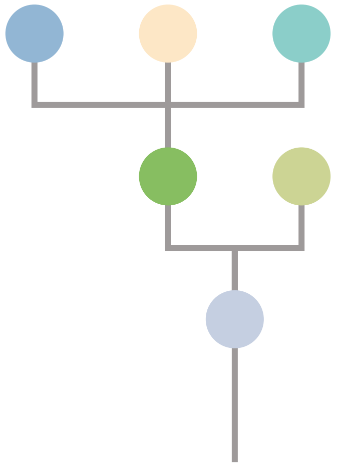

| Gene | Cytoband | Type | Cluster |
|---|---|---|---|
| *CBLB* | 3q13.11 | SNV | 1 |
| *KRAS* | 12p12.1 | SNV | 1 |
| *AKT1* | 14q32.33 | SNV | 1 |
| *TP53* | 17p13.1 | SNV | 1 |
| *APC* | 5q22.2 | InDel | 1 |
| *TCF7L2* | 10q25.3 | InDel | 1 |
| *LRP1B* | 2q22.1 | SNV | 3 |

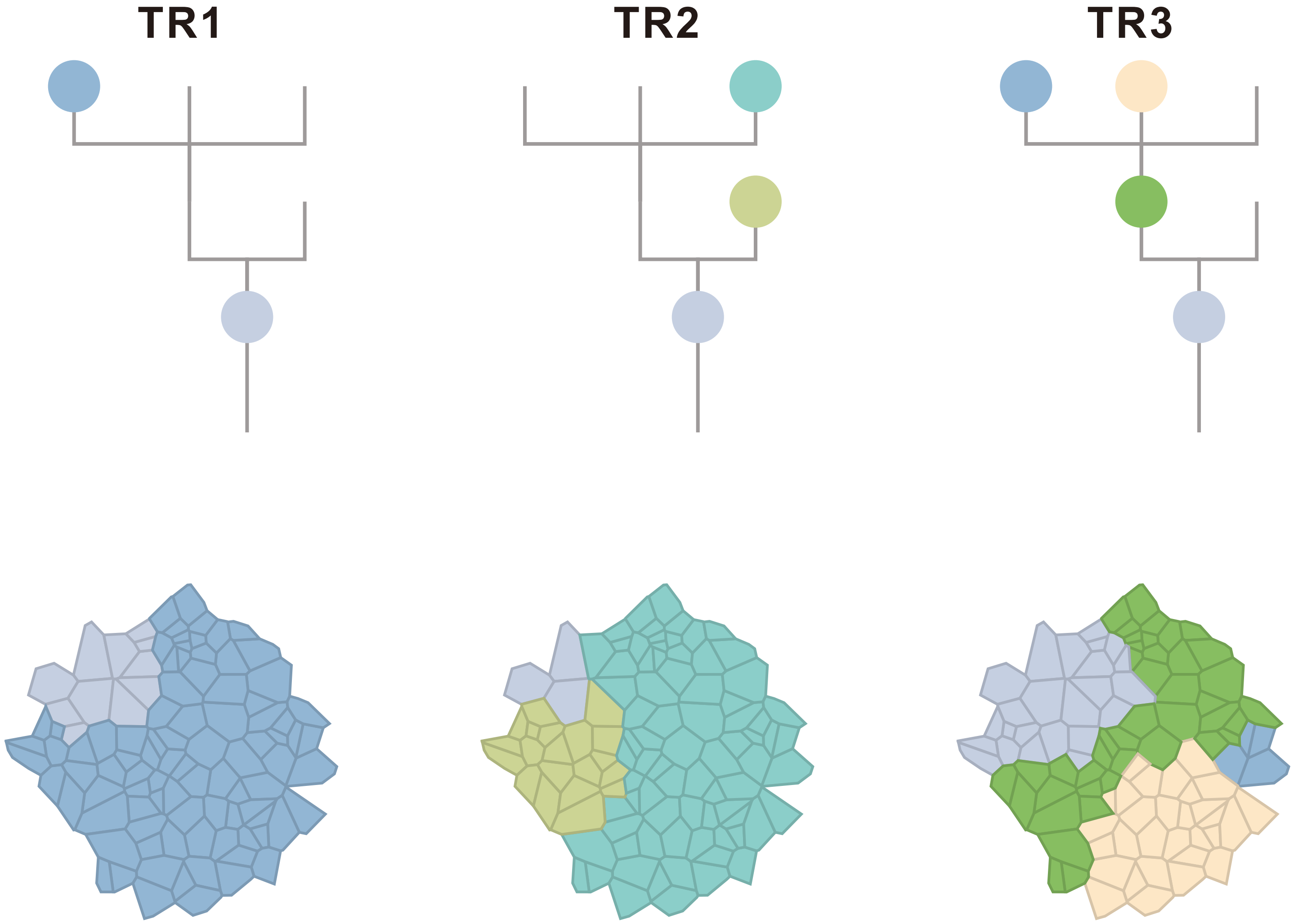

### CRC19

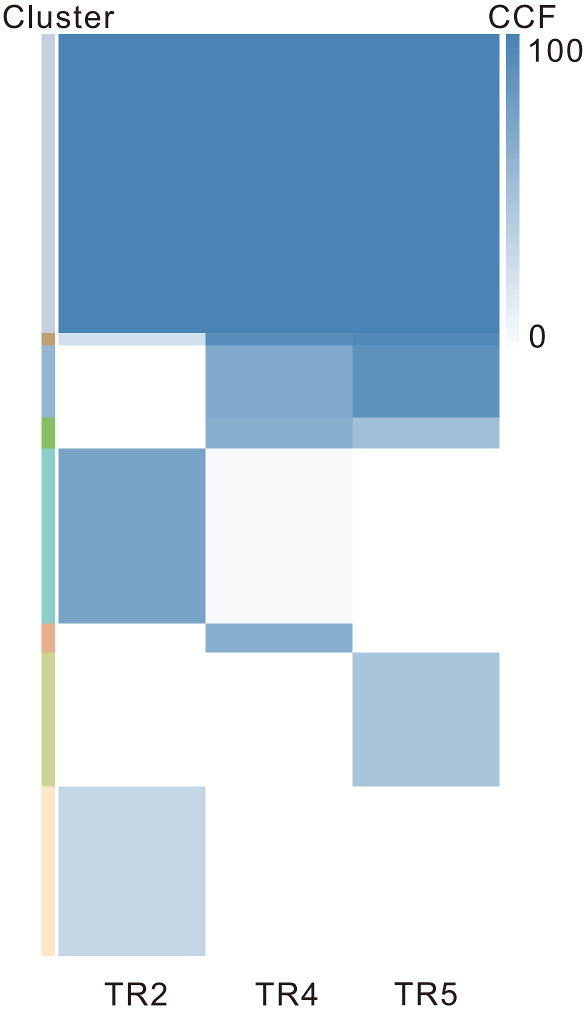

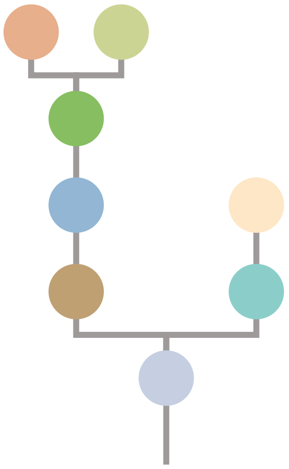

| Gene | Cytoband | Type | Cluster |
|---|---|---|---|
| *APC* | 5q22.2 | SNV | 1 |
| *KRAS* | 12p12.1 | SNV | 1 |
| *ARID2* | 12q12 | SNV | 1 |
| *FLCN* | 17p11.2 | SNV | 1 |
| *TP53* | 17p13.1 | SNV | 1 |
| *SRC* | 20q11.23 | Amp | 1 |
| *MAFB* | 20q12 | Amp | 1 |
| *PLCG1* | 20q12 | Amp | 1 |
| *NFATC2* | 20q13.2 | Amp | 1 |
| *SALL4* | 20q13.2 | Amp | 1 |
| *GNAS* | 20q13.32 | Amp | 1 |
| *PTK6* | 20q13.33 | Amp | 1 |
| *PER1* | 17p13.1 | SNV | 3 |
| *AMER1* | Xq11.2 | SNV | NA |

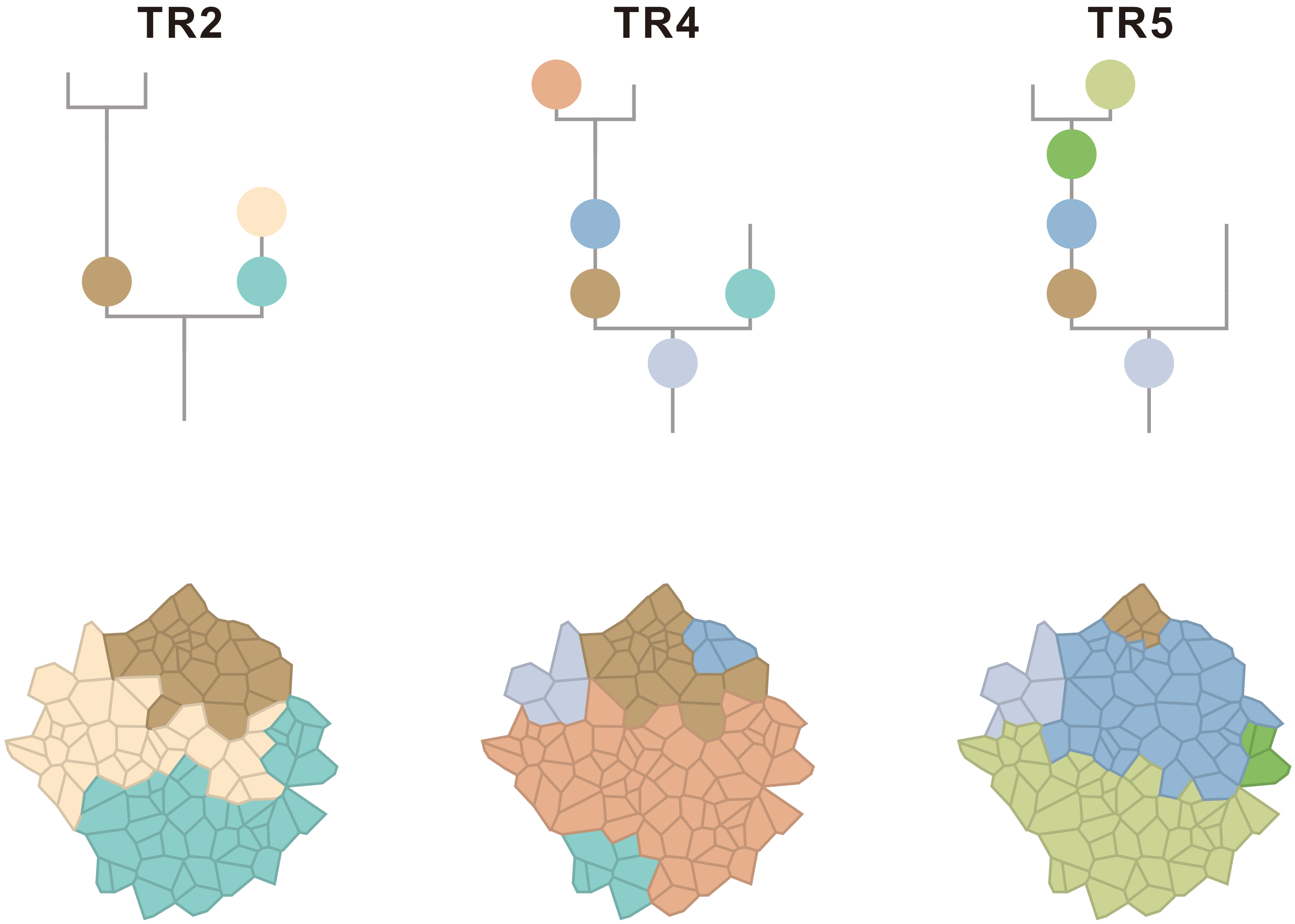

### CRC20

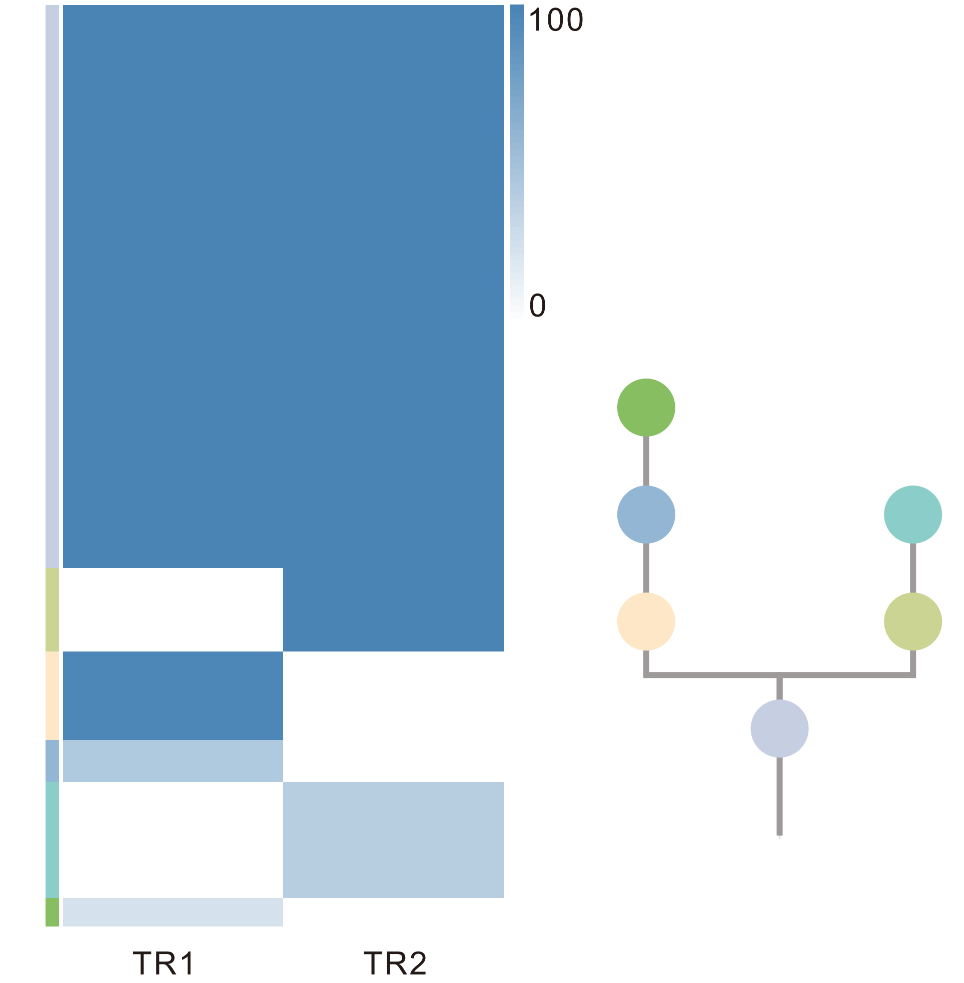

| Gene | Cytoband | Type | Cluster |
|---|---|---|---|
| *FBXW7* | 4q31.3 | SNV | 1 |
| *APC* | 5q22.2 | SNV | 1 |
| *TP53* | 17p13.1 | SNV | 1 |
| *SRC* | 20q11.23 | Amp | 1 |
| *MAFB* | 20q12 | Amp | 1 |
| *PLCG1* | 20q12 | Amp | 1 |
| *NFATC2* | 20q13.2 | Amp | 1 |
| *SALL4* | 20q13.2 | Amp | 1 |
| *GNAS* | 20q13.32 | Amp | 1 |
| *PTK6* | 20q13.33 | Amp | 1 |

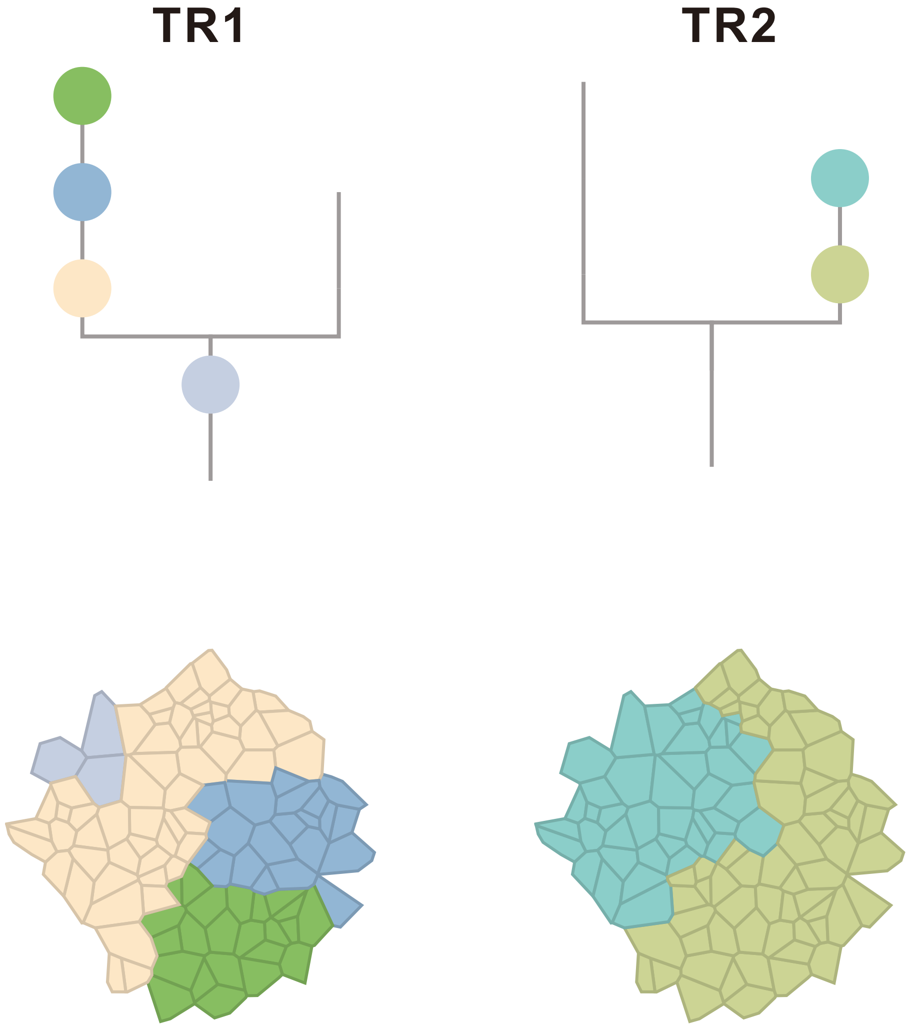

### CRC21

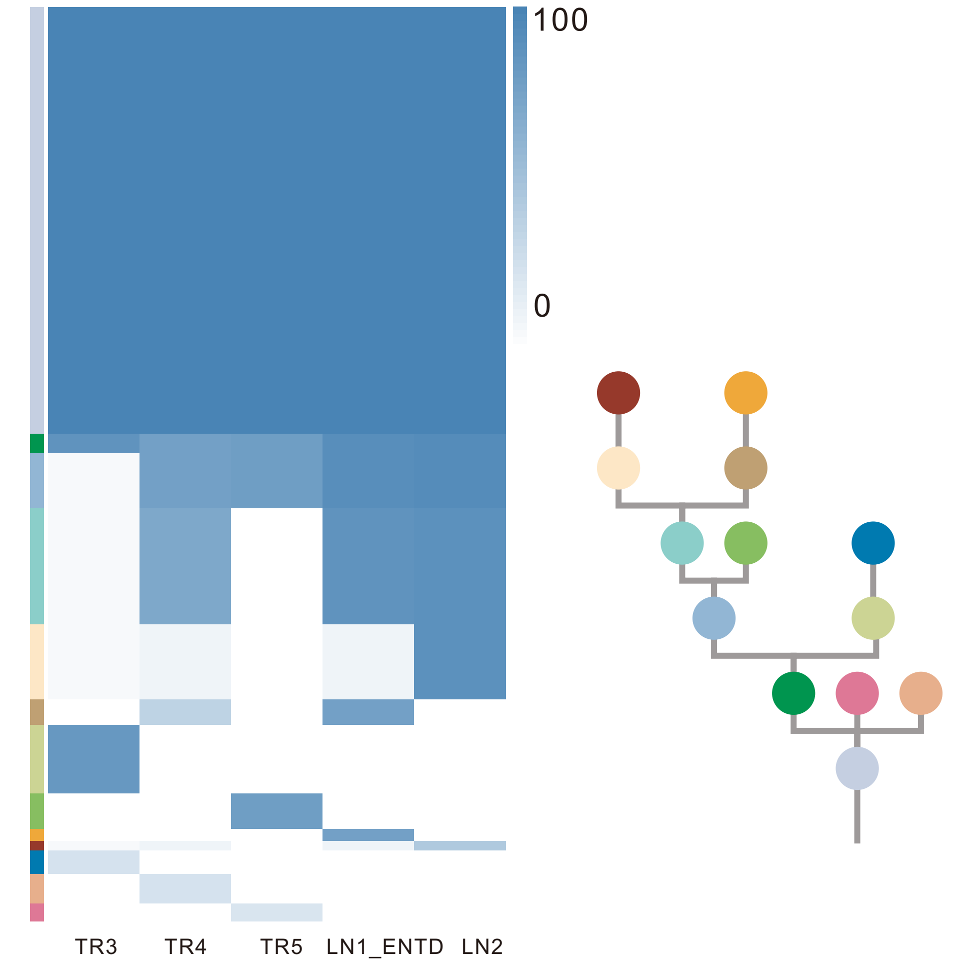

| Gene | Cytoband | Type | Cluster |
|---|---|---|---|
| * *ASXL2* | 2p23.3 | SNV | 1 |
| *BRAF* | 7q34 | SNV | 1 |
| * *ASXL2* | 2p23.3 | InDel | 1 |
| *TP53* | 17p13.1 | InDel | 1 |
| *RNF43* | 17q22 | InDel | 1 |
| *NCOA2* | 8q13.3 | Amp | 1 |
| *HEY1* | 8q21.13 | Amp | 1 |
| *RUNX1T1* | 8q21.3 | Amp | 1 |
| *CDH17* | 8q22.1 | Amp | 1 |
| *PABPC1* | 8q22.3 | Amp | 1 |
| *UBR5* | 8q22.3 | Amp | 1 |
| *RAD21* | 8q24.11 | Amp | 1 |
| *MYC* | 8q24.21 | Amp | 1 |
| *RECQL4* | 8q24.3 | Amp | 1 |
| *CCND2* | 12p13.32 | Amp | 2 |
| *KDM5A* | 12p13.33 | Amp | 2 |
| *ERBB4* | 2q34 | SNV | 3 |
| *TSC1* | 9q34.13 | SNV | 3 |

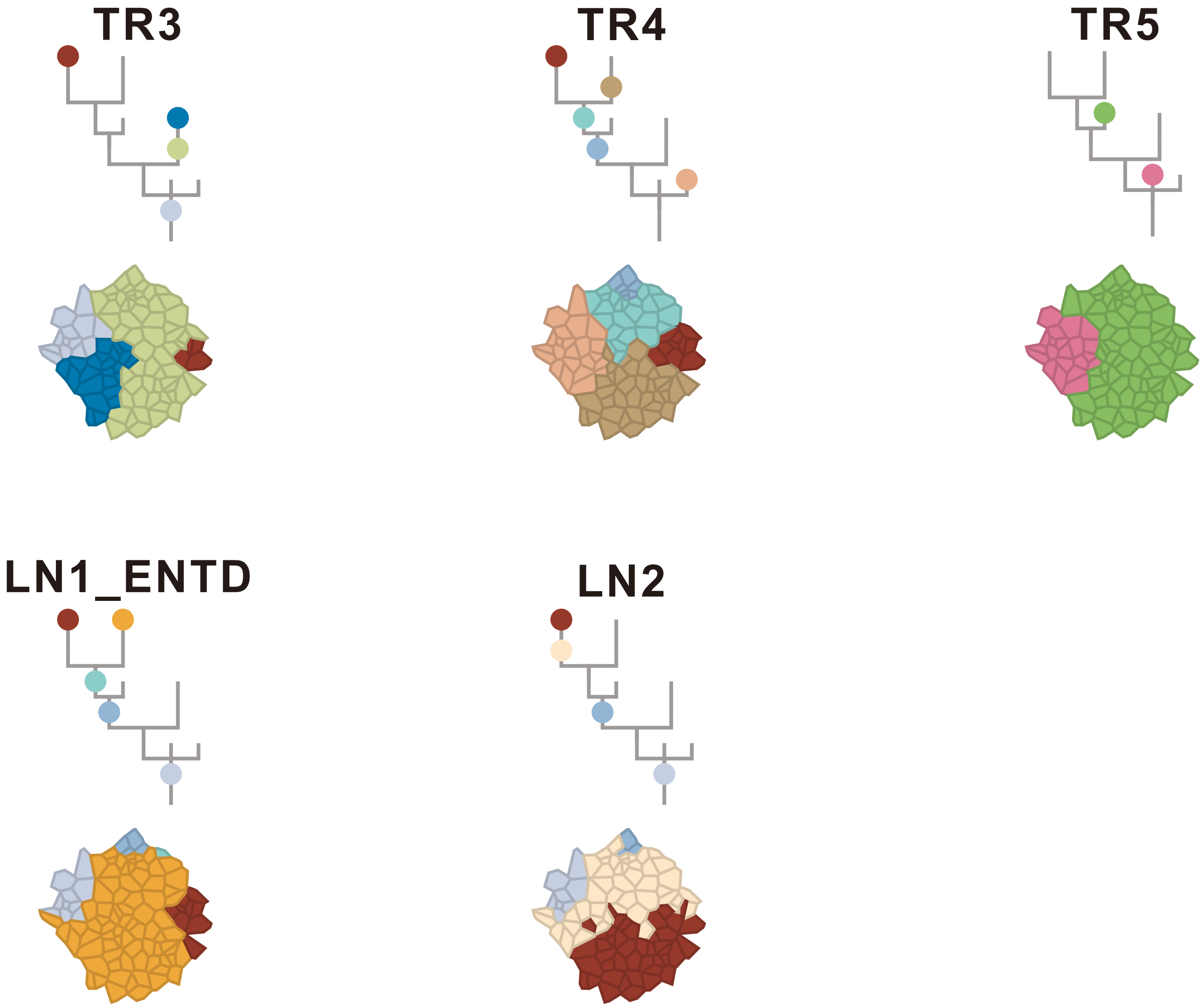

### CRC22

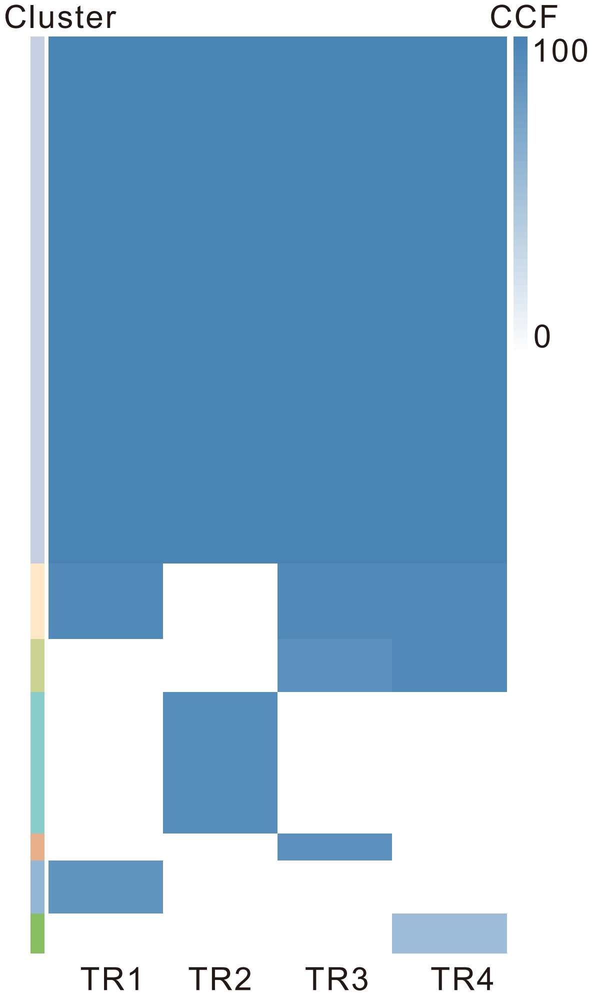

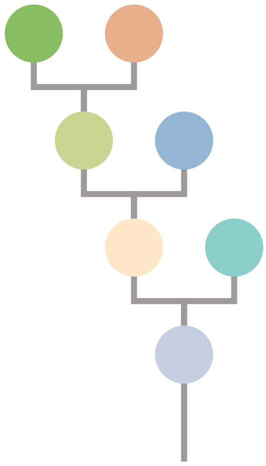

| Gene | Cytoband | Type | Cluster |
|---|---|---|---|
| *APC* | 5q22.2 | SNV | 1 |
| *KRAS* | 12p12.1 | SNV | 1 |
| *TP53* | 17p13.1 | SNV | 1 |
| *FLT3* | 13q12.2 | Amp | 1 |
| *FOXO1* | 13q14.11 | Amp | 1 |
| *CYSLTR2* | 13q14.2 | Amp | 1 |
| *SRC* | 20q11.23 | Amp | 1 |
| *MAFB* | 20q12 | Amp | 1 |
| *PLCG1* | 20q12 | Amp | 1 |
| *NFATC2* | 20q13.2 | Amp | 1 |
| *SALL4* | 20q13.2 | Amp | 1 |
| *GNAS* | 20q13.32 | Amp | 1 |
| *PTK6* | 20q13.33 | Amp | 1 |
| *KMT2D* | 12q13.12 | SNV | 2 |
| *AR* | Xq12 | SNV | NA |

## TR1

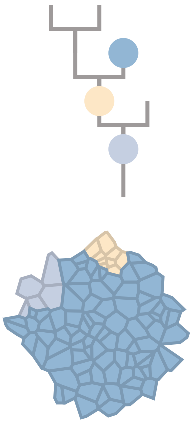

## TR2

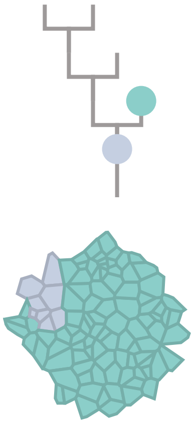

## TR3

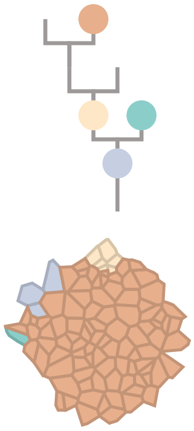

## TR4

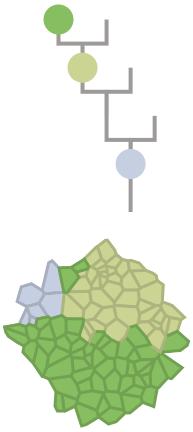

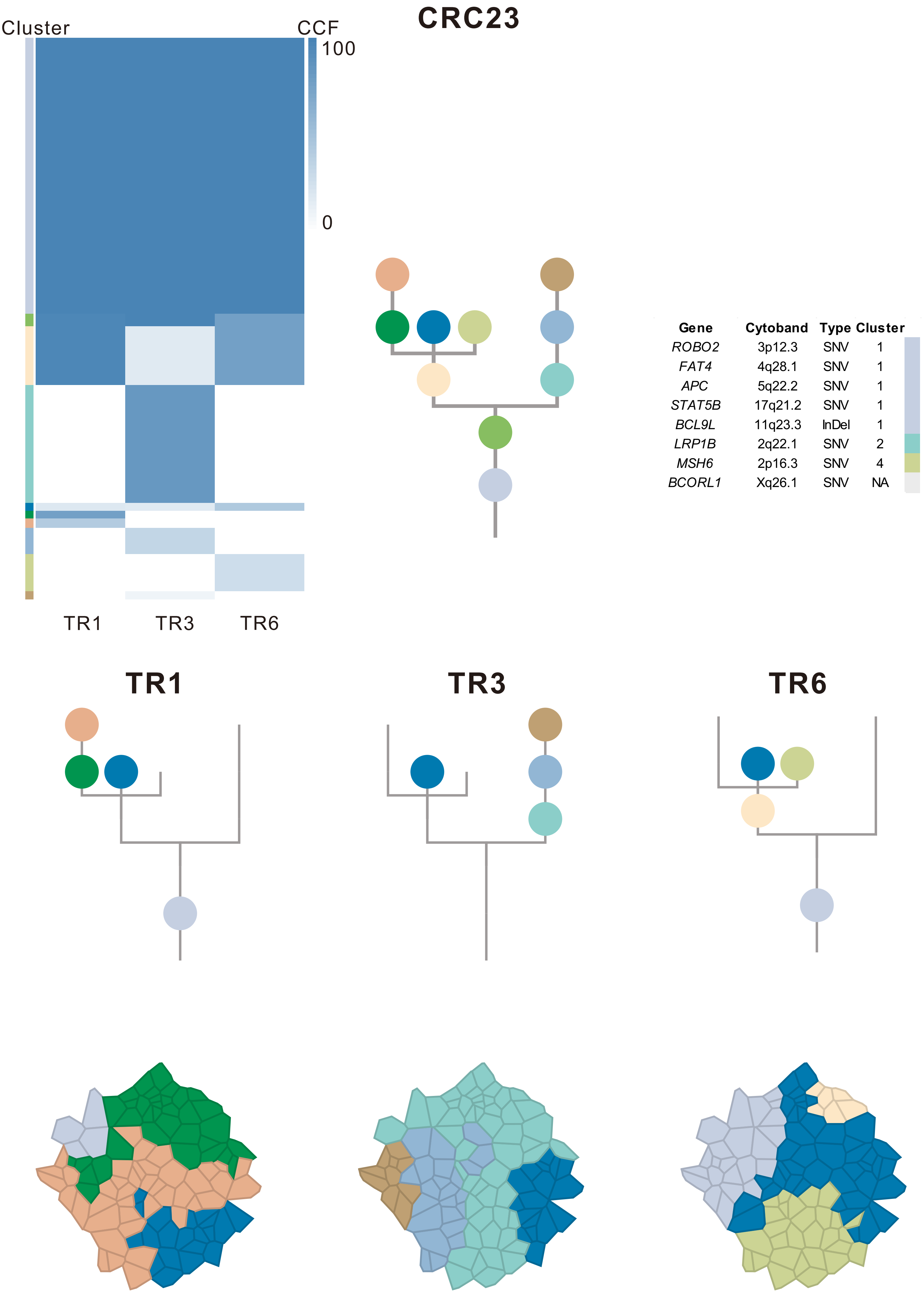
CRC23
Cluster
CCF
100
0
TR1
TR3
TR6
Gene Cytoband Type Cluster
ROBO2 3p12.3 SNV 1
FAT4 4q28.1 SNV 1
APC 5q22.2 SNV 1
STAT5B 17q21.2 SNV 1
BCL9L 11q23.3 InDel 1
LRP1B 2q22.1 SNV 2
MSH6 2p16.3 SNV 4
BCORL1 Xq26.1 SNV NA
TR1
TR3
TR6

### CRC24

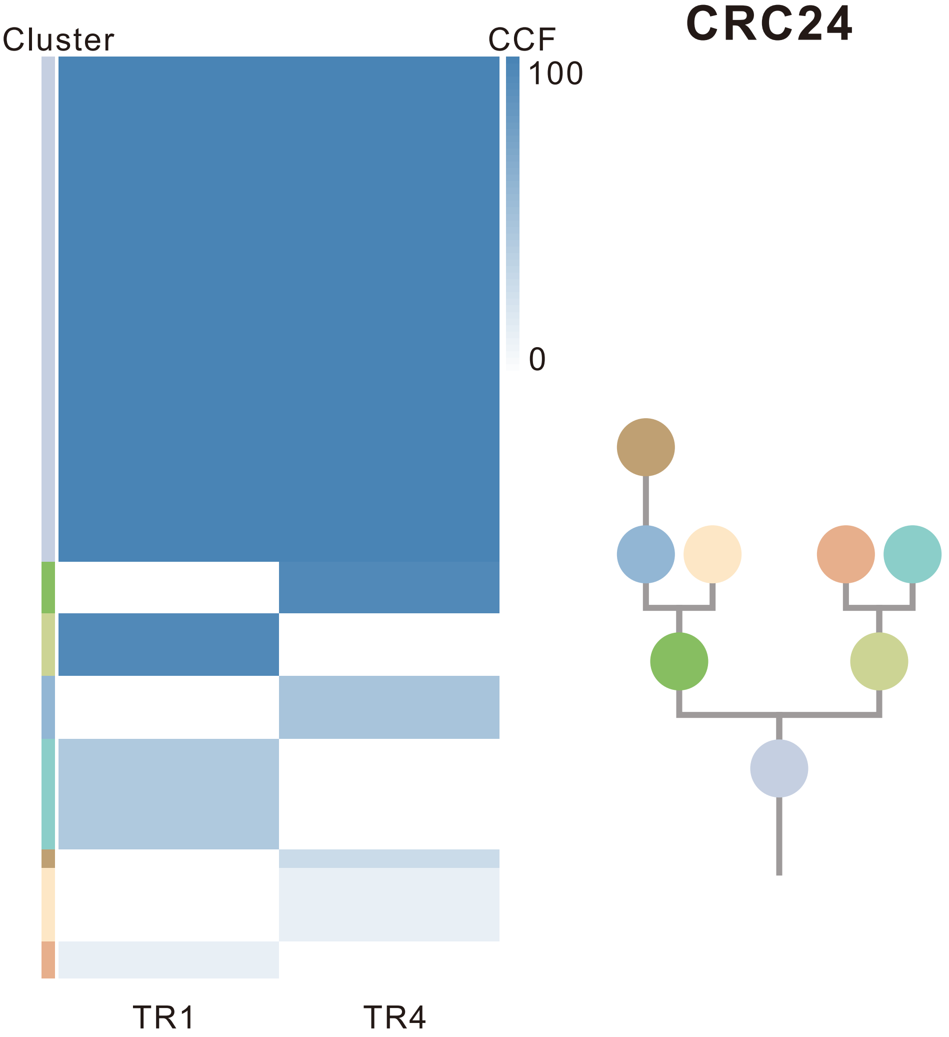

| Gene | Cytoband | Type | Cluster |
| --- | --- | --- | --- |
| *BMPR1A* | 10q23.2 | SNV | 1 |
| *KRAS* | 12p12.1 | SNV | 1 |
| *TP53* | 17p13.1 | SNV | 1 |
| *GRIN2A* | 16p13.2 | SNV | 3 |

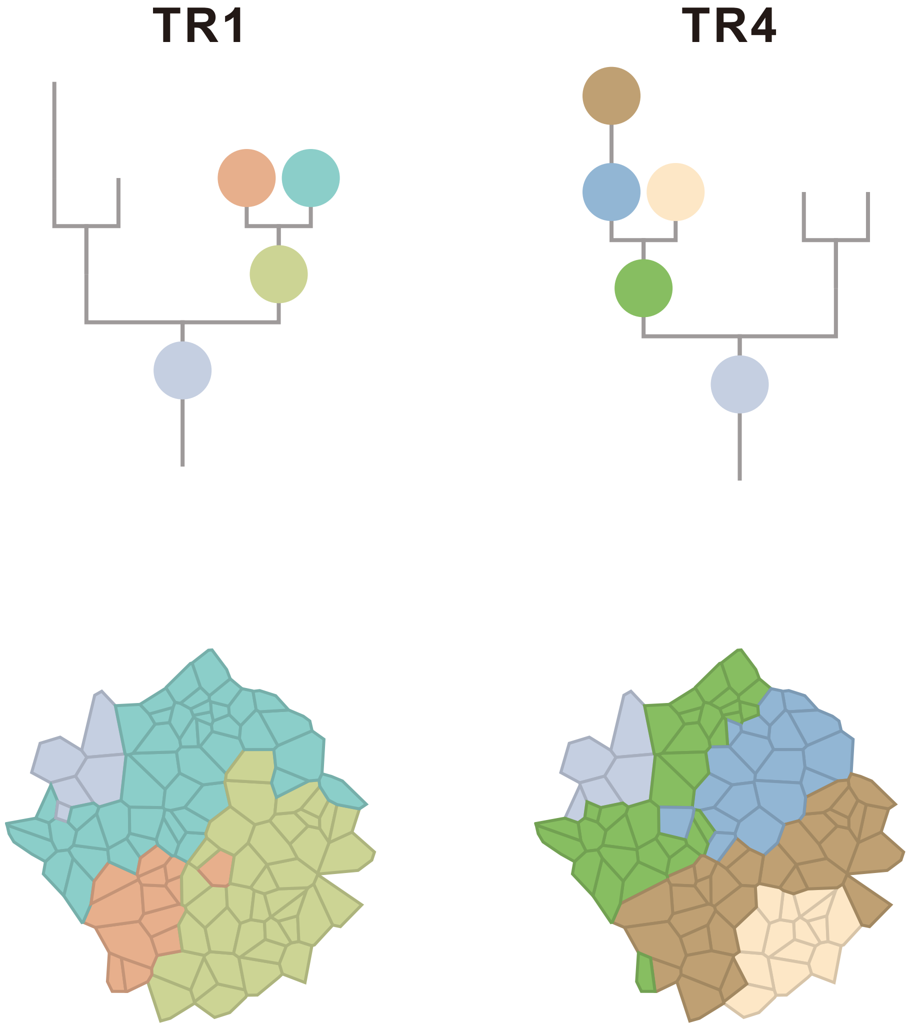

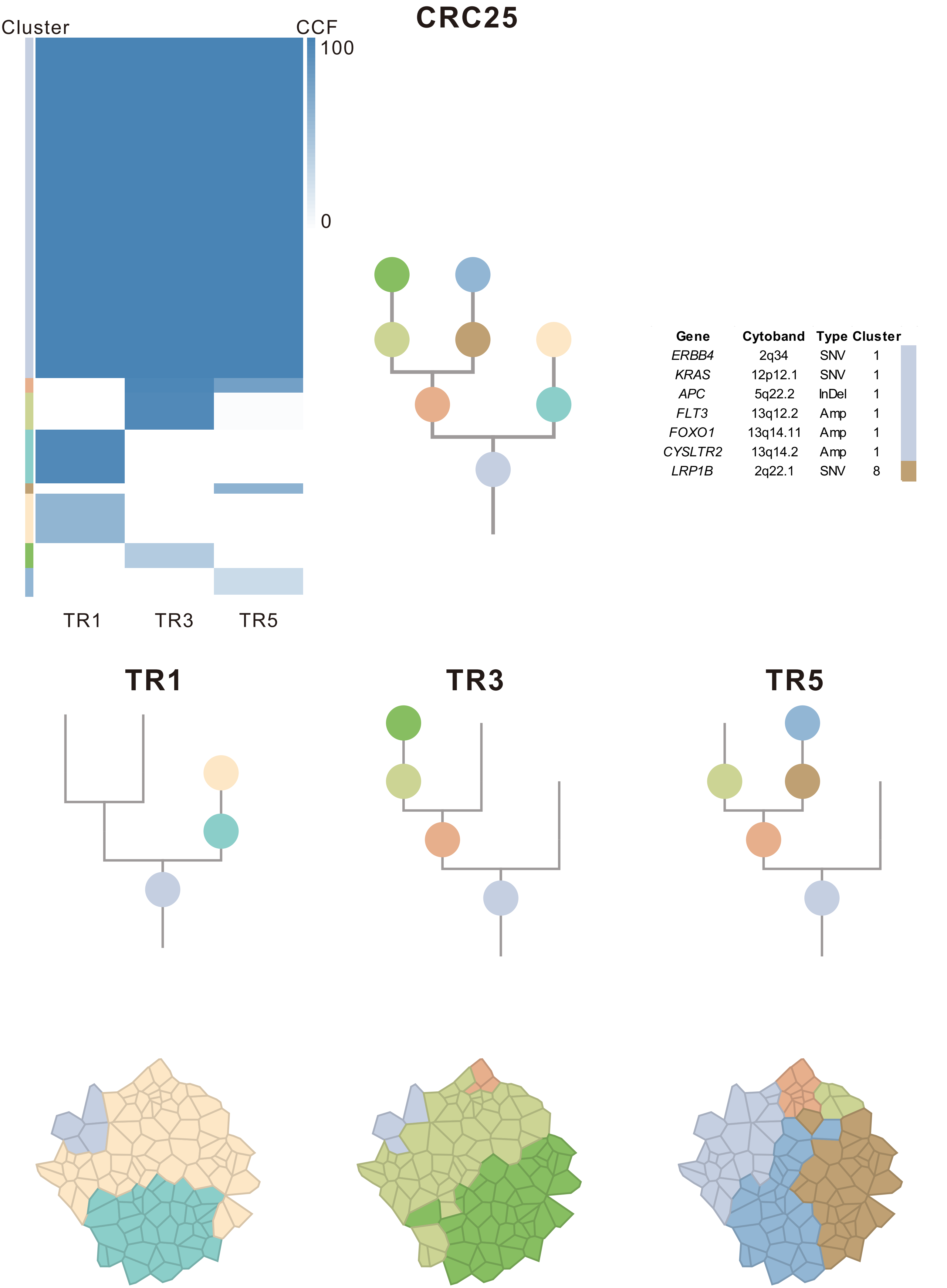

| Gene | Cytoband | Type | Cluster |
|---|---|---|---|
| *ERBB4* | 2q34 | SNV | 1 |
| *KRAS* | 12p12.1 | SNV | 1 |
| *APC* | 5q22.2 | InDel | 1 |
| *FLT3* | 13q12.2 | Amp | 1 |
| *FOXO1* | 13q14.11 | Amp | 1 |
| *CYSLTR2* | 13q14.2 | Amp | 1 |
| *LRP1B* | 2q22.1 | SNV | 8 |

### CRC26

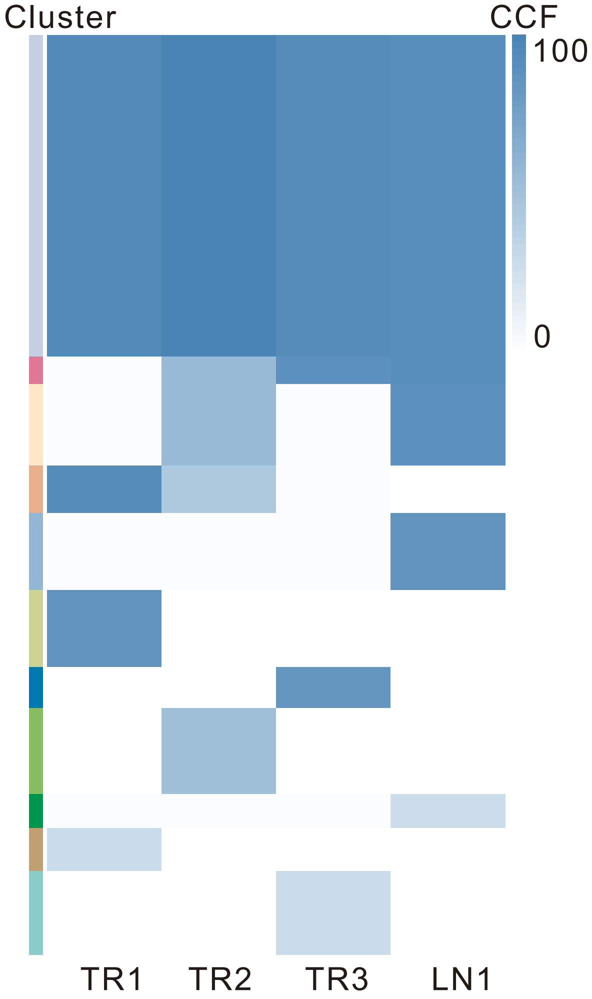

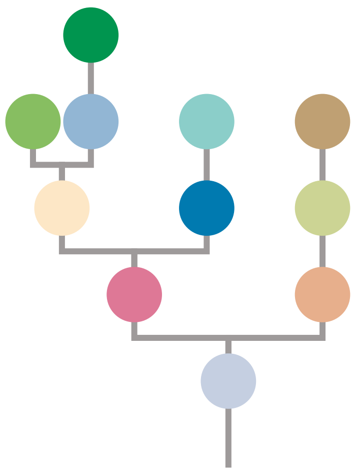

| Gene | Cytoband | Type | Cluster |
|---|---|---|---|
| *LEF1* | 4q25 | SNV | 1 |
| *FBXW7* | 4q31.3 | SNV | 1 |
| * *APC* | 5q22.2 | SNV | 1 |
| *KRAS* | 12p12.1 | SNV | 1 |
| * *APC* | 5q22.2 | InDel | 1 |
| *TP53* | 17p13.1 | InDel | 1 |
| *FLT3* | 13q12.2 | Amp | 1 |
| *CYSLTR2* | 13q14.2 | Amp | 1 |
| *FOXO1* | 13q14.11 | Amp | 1 |
| *CSMD3* | 8q23.3 | SNV | 3 |
| *EIF3E* | 8q23.1 | SNV | 5 |
| *SMAD3* | 15q22.33 | InDel | 6 |
| *MUC4* | 3q29 | Amp | 6 |
| *CREB3L2* | 7q33 | Amp | 6 |
| *TRIM24* | 7q33 | Amp | 6 |
| *BRAF* | 7q34 | Amp | 6 |
| *EZH2* | 7q36.1 | Amp | 6 |
| *SRC* | 20q11.23 | Amp | 6 |
| *MAFB* | 20q12 | Amp | 6 |
| *PLCG1* | 20q12 | Amp | 6 |
| *NFATC2* | 20q13.2 | Amp | 6 |
| *SALL4* | 20q13.2 | Amp | 6 |
| *GNAS* | 20q13.32 | Amp | 6 |
| *PTK6* | 20q13.33 | Amp | 6 |
| *AXIN1* | 16p13.3 | Del | 7 |
| *FGFR1* | 8p11.23 | Amp | NA |

## TR1

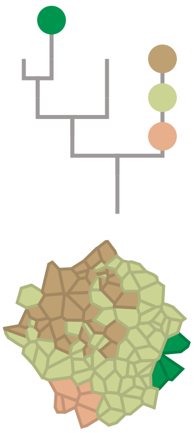

## TR2

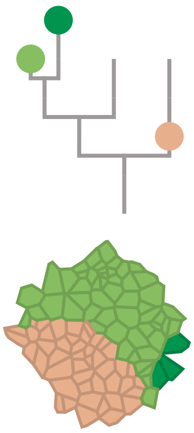

## TR3

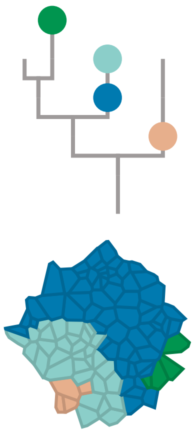

## LN1

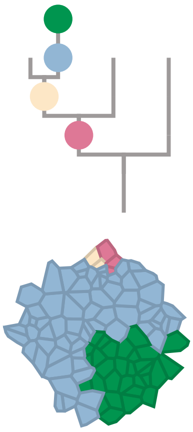

### CRC27

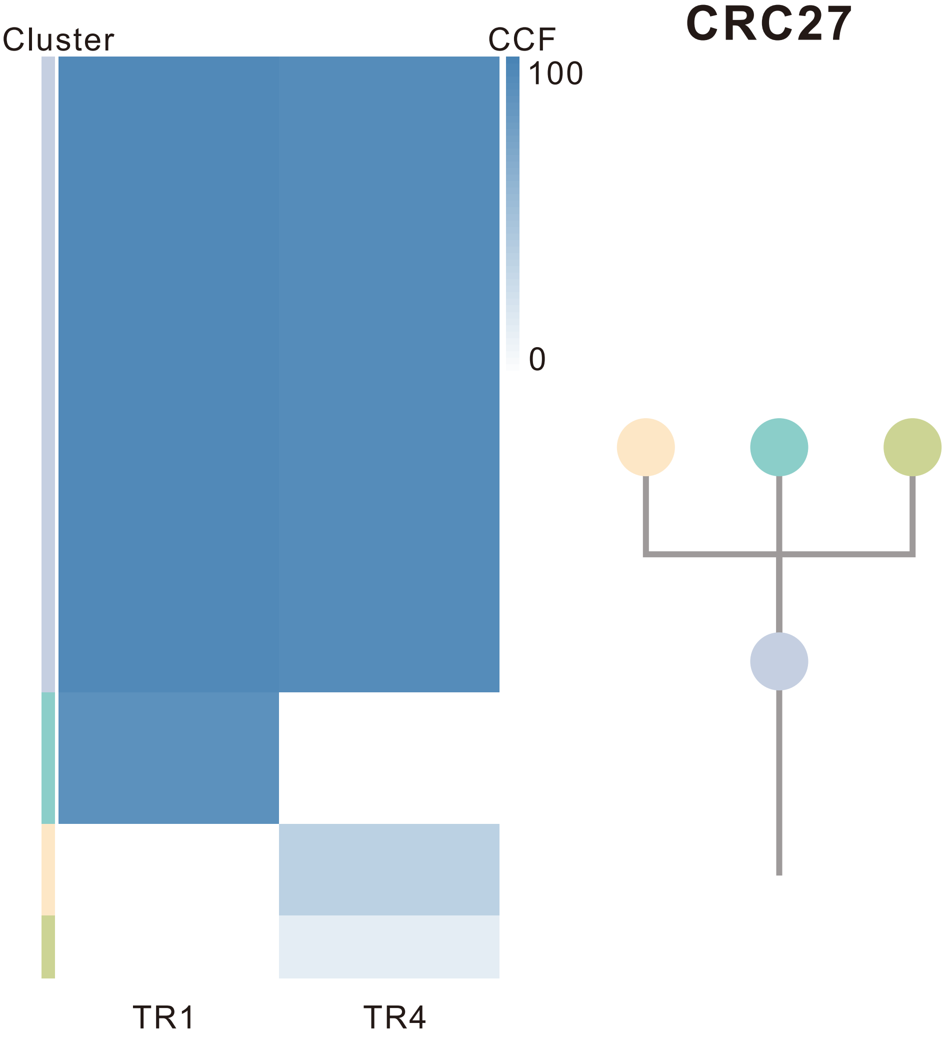

| Gene | Cytoband | Type | Cluster |
|---|---|---|---|
| * *APC* | 5q22.2 | SNV | 1 |
| * *TP53* | 17p13.1 | SNV | 1 |
| * *TP53* | 17p13.1 | SNV | 1 |
| * *APC* | 5q22.2 | InDel | 1 |
| *LZTR1* | 22q11.21 | InDel | 1 |
| *MUC4* | 3q29 | Amp | 1 |
| *SRC* | 20q11.23 | Amp | 1 |
| *MAFB* | 20q12 | Amp | 1 |
| *PLCG1* | 20q12 | Amp | 1 |
| *NFATC2* | 20q13.2 | Amp | 1 |
| *SALL4* | 20q13.2 | Amp | 1 |
| *SSX2* | Xp11.22 | Amp | 1 |
| *ARAF* | Xp11.23 | Amp | 1 |
| *GATA1* | Xp11.23 | Amp | 1 |
| *SSX1* | Xp11.23 | Amp | 1 |
| *SSX4* | Xp11.23 | Amp | 1 |
| *TFE3* | Xp11.23 | Amp | 1 |
| *WAS* | Xp11.23 | Amp | 1 |
| *KDM6A* | Xp11.3 | Amp | 1 |
| *IRS4* | Xq22.3 | Del | 1 |
| *STAG2* | Xq25 | Del | 1 |
| *BCORL1* | Xq26.1 | Del | 1 |
| *ELF4* | Xq26.1 | Del | 1 |
| *GPC3* | Xq26.2 | Del | 1 |
| *PHF6* | Xq26.2 | Del | 1 |
| *ATP2B3* | Xq28 | Del | 1 |
| *RPL10* | Xq28 | Del | 1 |
| *NOTCH2* | 1p12 | Amp | NA |
| *BCL9* | 1q21.2 | Amp | NA |

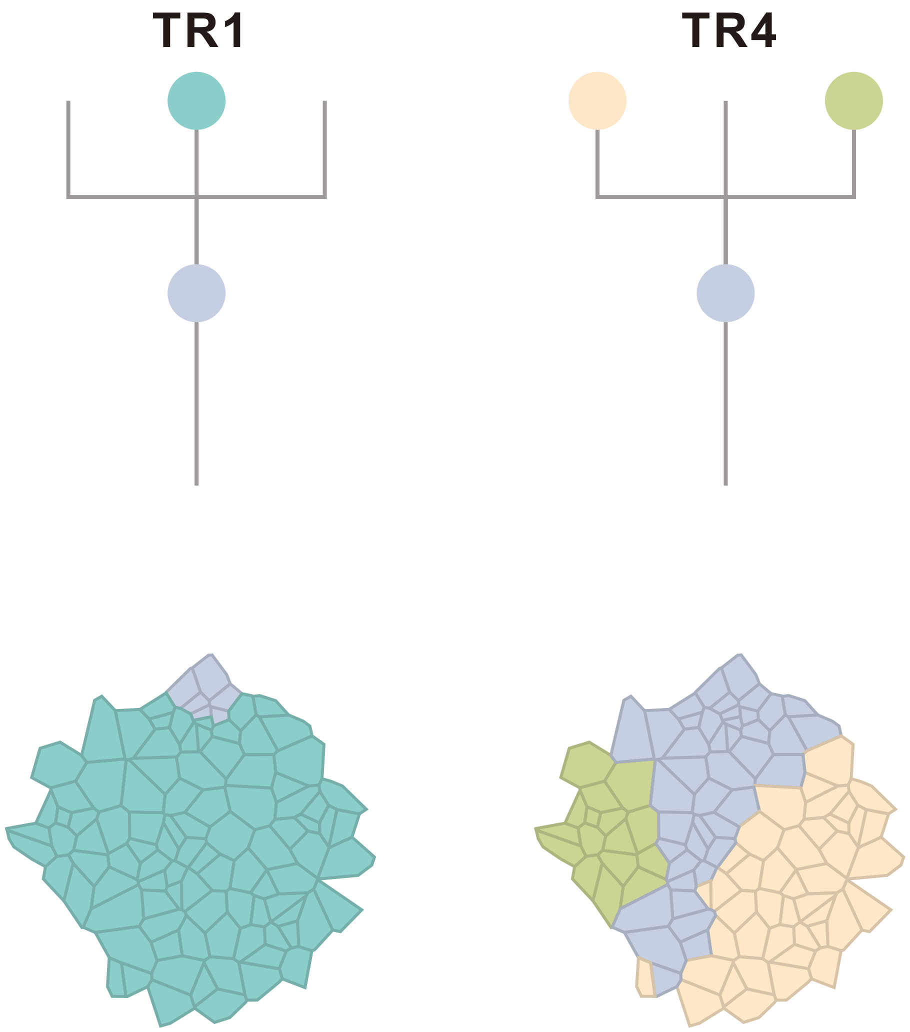

### CRC28

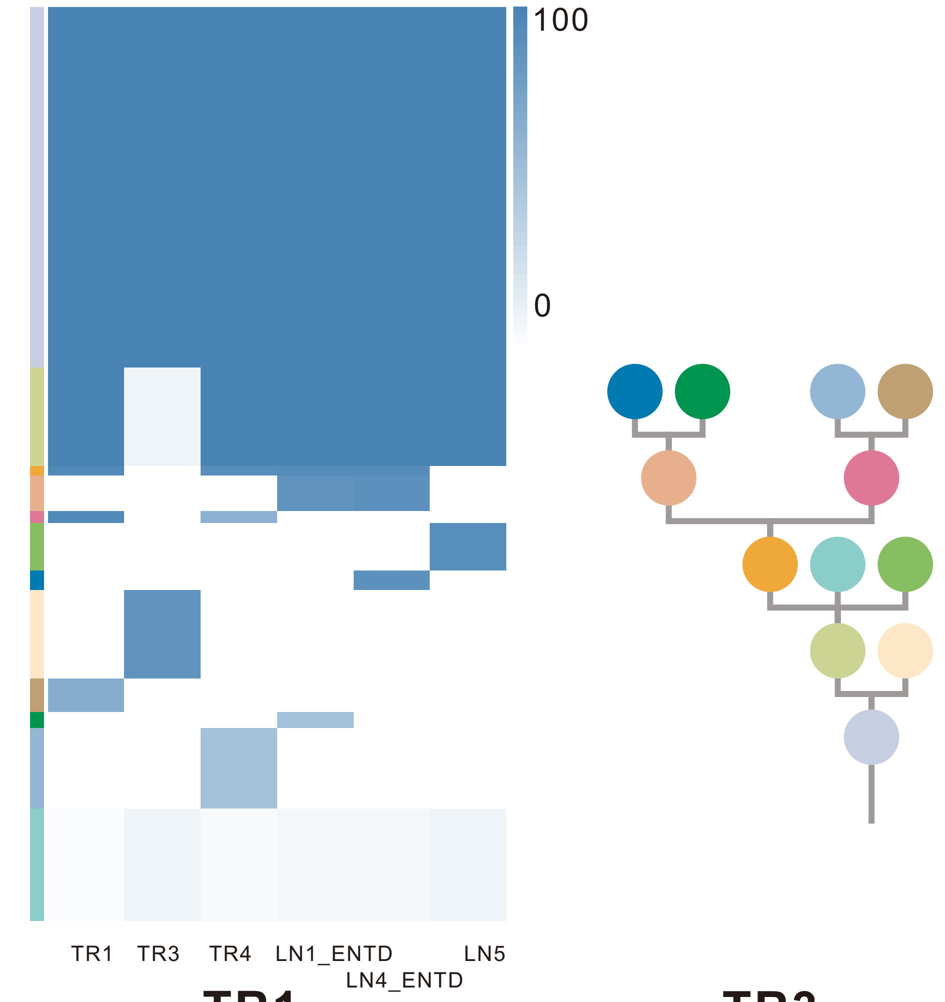

| Gene | Cytoband | Type | Cluster |
|---|---|---|---|
| * *APC* | 5q22.2 | SNV | 1 |
| * *APC* | 5q22.2 | SNV | 1 |
| *KRAS* | 12p12.1 | SNV | 1 |
| *CLTCL1* | 22q11.21 | SNV | 1 |
| *KRAS* | 12p12.1 | Amp | 1 |
| *CHD4* | 12p13.31 | Amp | 1 |
| *CCND2* | 12p13.32 | Amp | 1 |
| *KDM5A* | 12p13.33 | Amp | 1 |
| *IRS4* | Xq22.3 | Amp | 1 |
| *BCORL1* | Xq26.1 | Amp | 1 |
| *ELF4* | Xq26.1 | Amp | 1 |
| *GPC3* | Xq26.2 | Amp | 1 |
| *FAT4* | 4q28.1 | SNV | 2 |
| *ARID1A* | 1p36.11 | SNV | 3 |
| *ID3* | 1p36.12 | Del | 4 |
| *ARHGEF10L* | 1p36.13 | Del | 4 |
| *SDHB* | 1p36.13 | Del | 4 |
| *CAMTA1* | 1p36.31 | Del | 4 |
| *RPL22* | 1p36.31 | Del | 4 |
| *TNFRSF14* | 1p36.32 | Del | 4 |
| *CASP8* | 2q33.1 | SNV | 6 |
| *JAK2* | 9p24.1 | Amp | 6 |
| *PDCD1LG2* | 9p24.1 | Amp | 6 |
| *KDM5C* | Xp11.22 | SNV | NA |

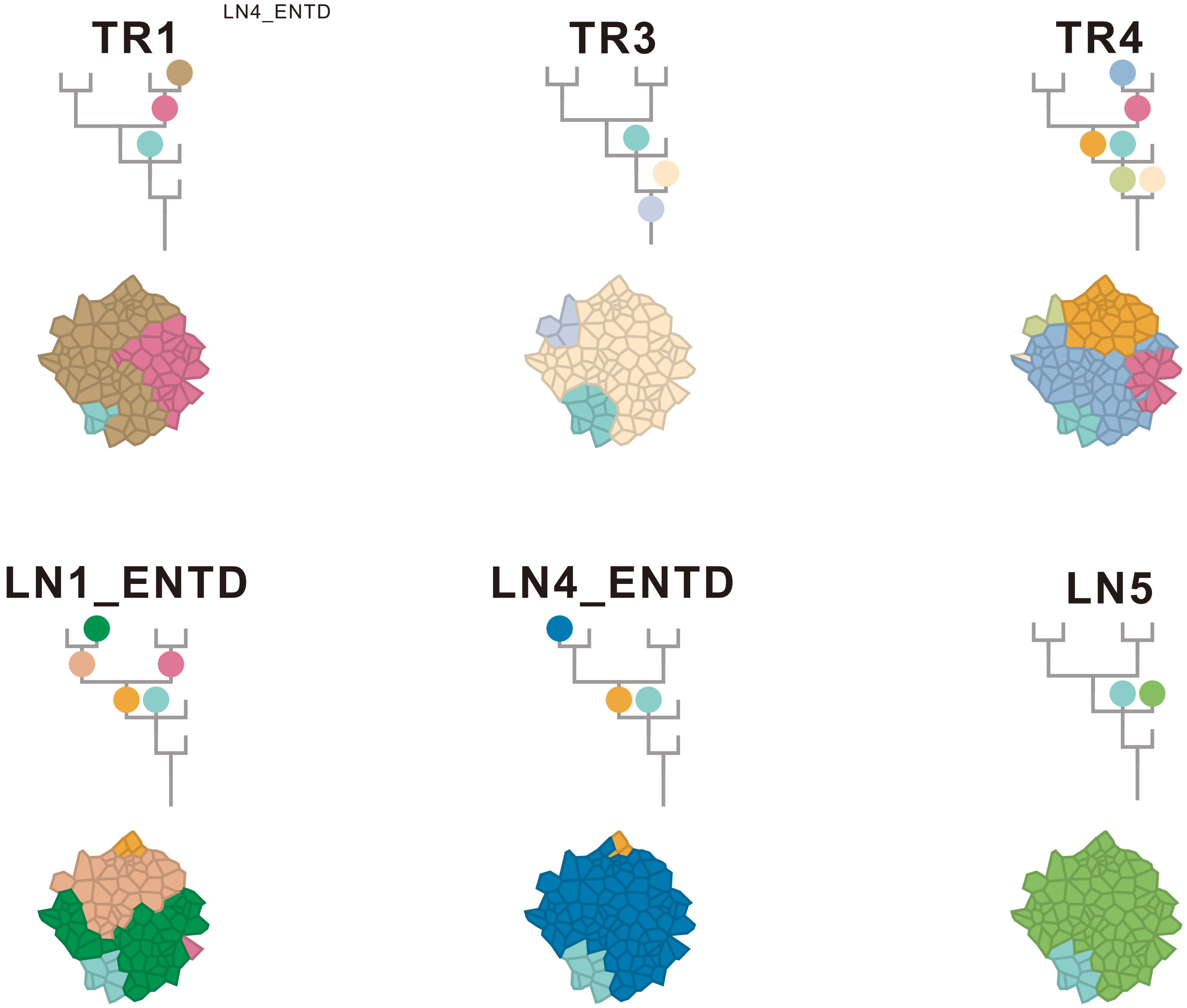

### CRC29

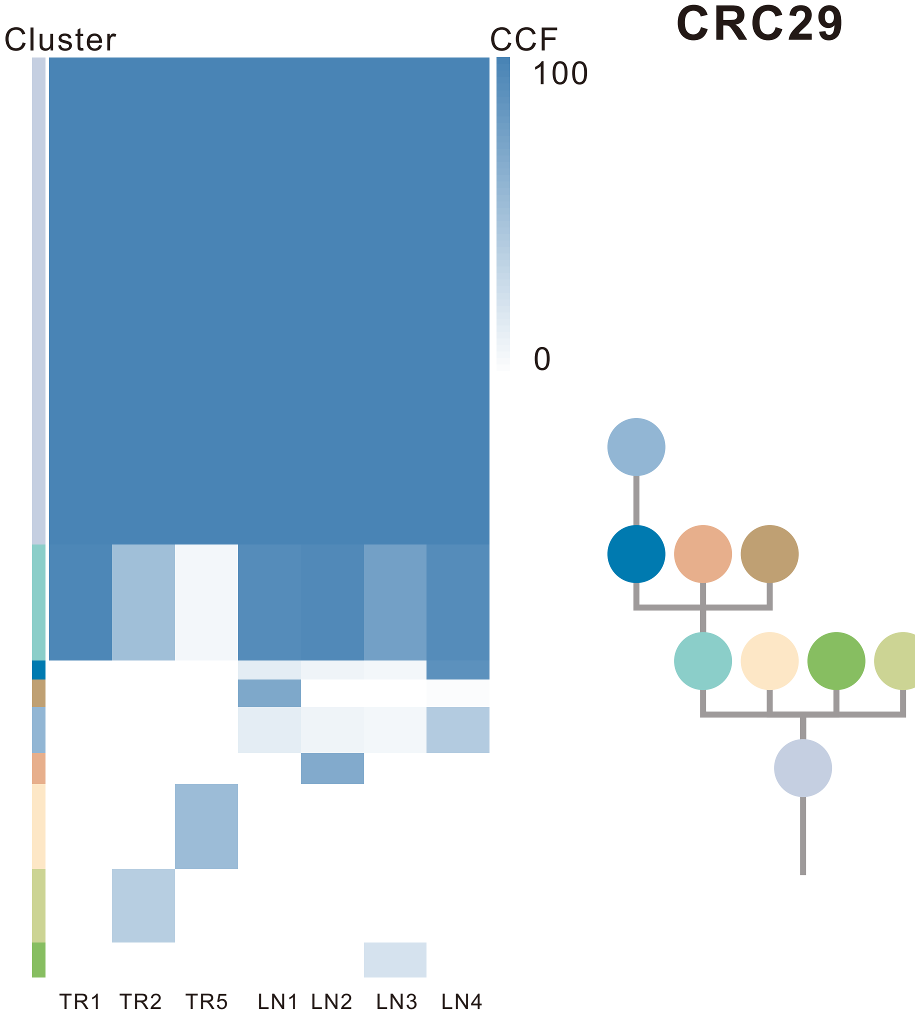

| Gene | Cytoband | Type | Cluster |
|---|---|---|---|
| *APC* | 5q22.2 | SNV | 1 |
| *BCL9L* | 11q23.3 | SNV | 1 |
| *NCOR2* | 12q24.31 | SNV | 1 |
| *TP53* | 17p13.1 | SNV | 1 |
| *FLT3* | 13q12.2 | Amp | 1 |
| *SRC* | 20q11.23 | Amp | 1 |
| *MAFB* | 20q12 | Amp | 1 |
| *PLCG1* | 20q12 | Amp | 1 |
| *NFATC2* | 20q13.2 | Amp | 1 |
| *SALL4* | 20q13.2 | Amp | 1 |
| *GNAS* | 20q13.32 | Amp | 1 |
| *PTK6* | 20q13.33 | Amp | 1 |
| *DDX5* | 17q23.3 | Amp | 2 |
| *SMAD3* | 15q22.33 | SNV | 4 |
| *LZTR1* | 22q11.21 | InDel | NA |
| *MUC4* | 3q29 | Amp | NA |
| *EGFR* | 7p11.2 | Amp | NA |
| *PLAG1* | 8q12.1 | Amp | NA |
| *PREX2* | 8q13.2 | Amp | NA |
| *NCOA2* | 8q13.3 | Amp | NA |
| *HEY1* | 8q21.13 | Amp | NA |
| *RUNX1T1* | 8q21.3 | Amp | NA |
| *CDH17* | 8q22.1 | Amp | NA |
| *PABPC1* | 8q22.3 | Amp | NA |
| *UBR5* | 8q22.3 | Amp | NA |
| *RAD21* | 8q24.11 | Amp | NA |
| *MYC* | 8q24.21 | Amp | NA |
| *RECQL4* | 8q24.3 | Amp | NA |
| *PPM1D* | 17q23.2 | Amp | NA |
| *CD79B* | 17q23.3 | Amp | NA |
| *AXIN1* | 16p13.3 | Del | NA |
| *PRKAR1A* | 17q24.2 | Del | NA |

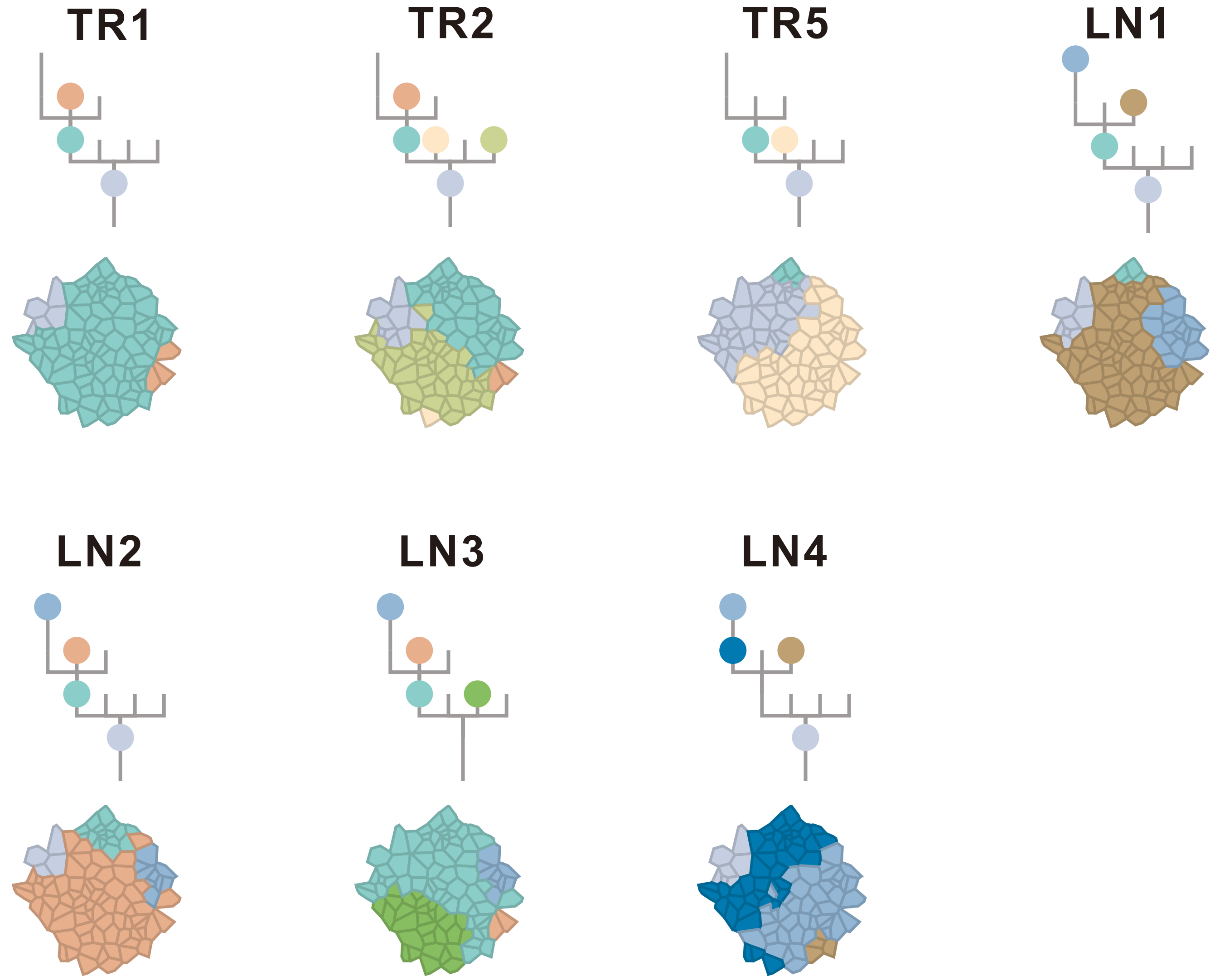

### CRC30

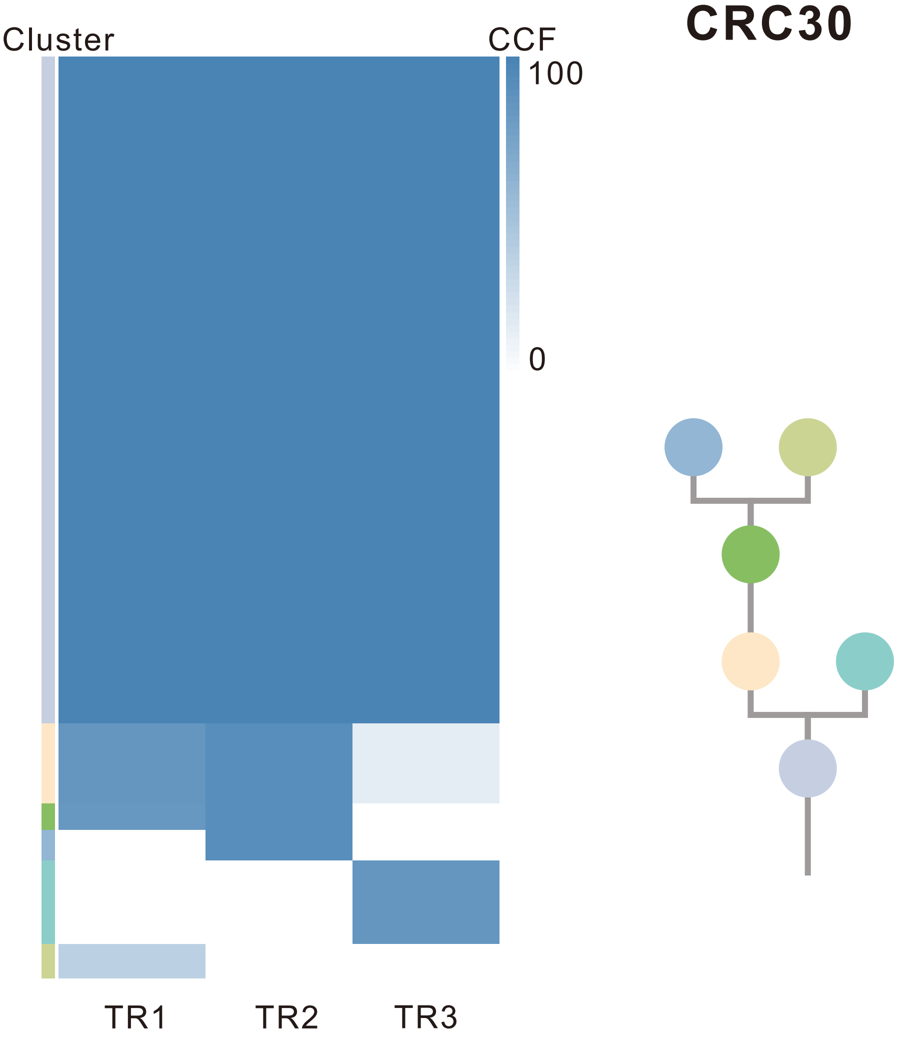

| Gene | Cytoband | Type | Cluster |
|---|---|---|---|
| *APC* | 5q22.2 | SNV | 1 |
| *PTPRB* | 12q15 | SNV | 1 |
| *TP53* | 17p13.1 | SNV | 1 |
| *IL7R* | 5p13.2 | Amp | 1 |
| *IKBKB* | 8p11.21 | Amp | 1 |
| *KAT6A* | 8p11.21 | Amp | 1 |
| *PLAG1* | 8q12.1 | Amp | 1 |
| *PREX2* | 8q13.2 | Amp | 1 |
| *NCOA2* | 8q13.3 | Amp | 1 |
| *HEY1* | 8q21.13 | Amp | 1 |
| *RUNX1T1* | 8q21.3 | Amp | 1 |
| *CDH17* | 8q22.1 | Amp | 1 |
| *PABPC1* | 8q22.3 | Amp | 1 |
| *UBR5* | 8q22.3 | Amp | 1 |
| *RAD21* | 8q24.11 | Amp | 1 |
| *MYC* | 8q24.21 | Amp | 1 |
| *RECQL4* | 8q24.3 | Amp | 1 |

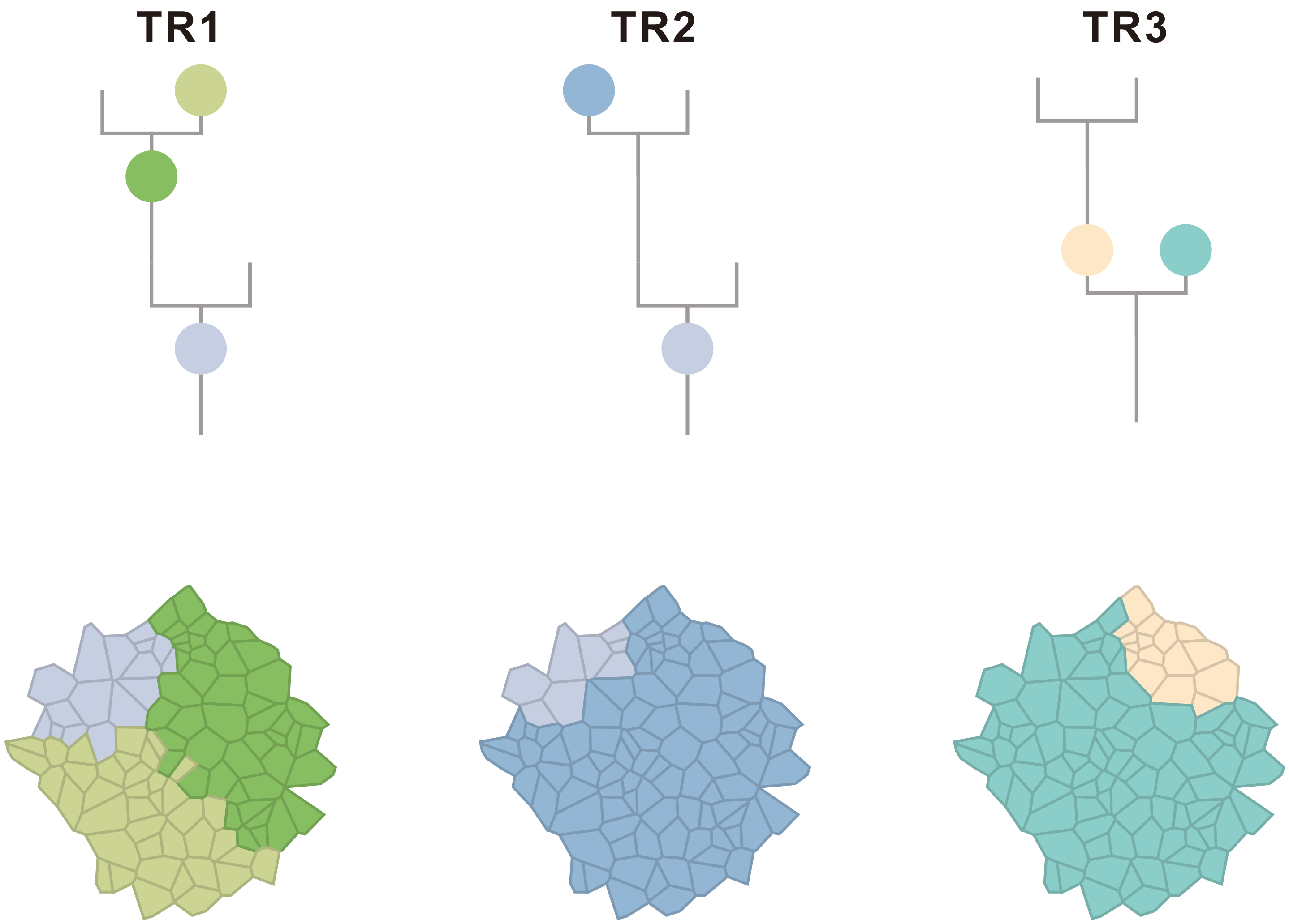

### CRC31

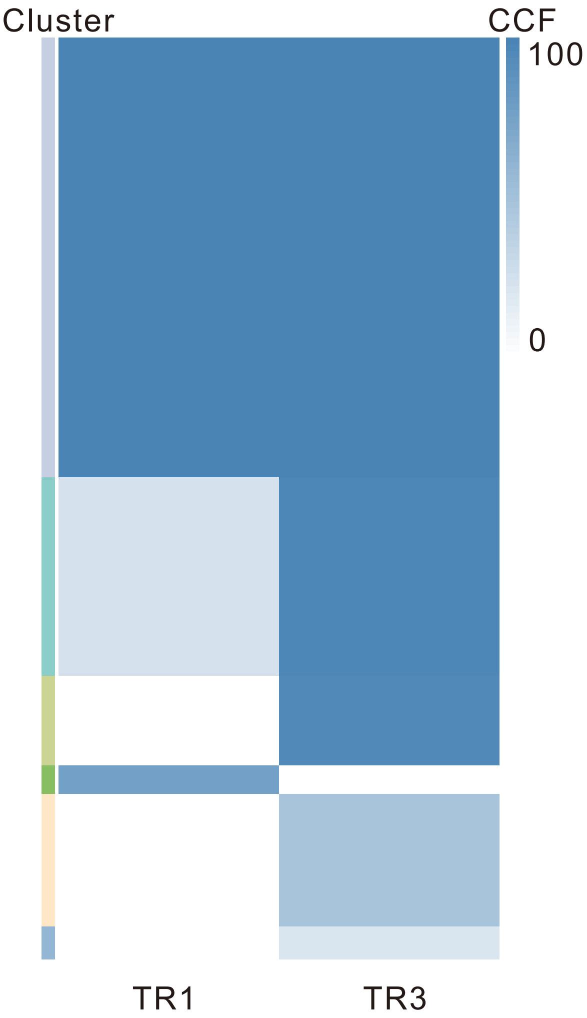

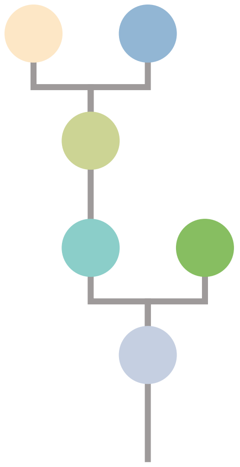

| Gene | Cytoband | Type | Cluster |
|---|---|---|---|
| *APC* | 5q22.2 | SNV | 1 |
| *KRAS* | 12p12.1 | SNV | 1 |
| *TP53* | 17p13.1 | SNV | 1 |
| *ARID1A* | 1p36.11 | SNV | 2 |
| *CLTCL1* | 22q11.21 | SNV | 3 |

## TR1

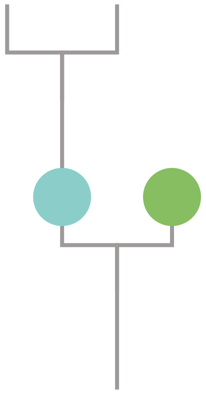

## TR3

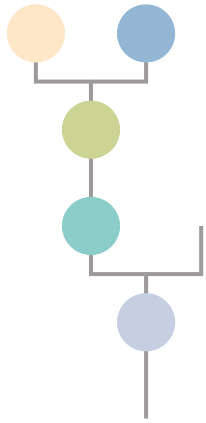

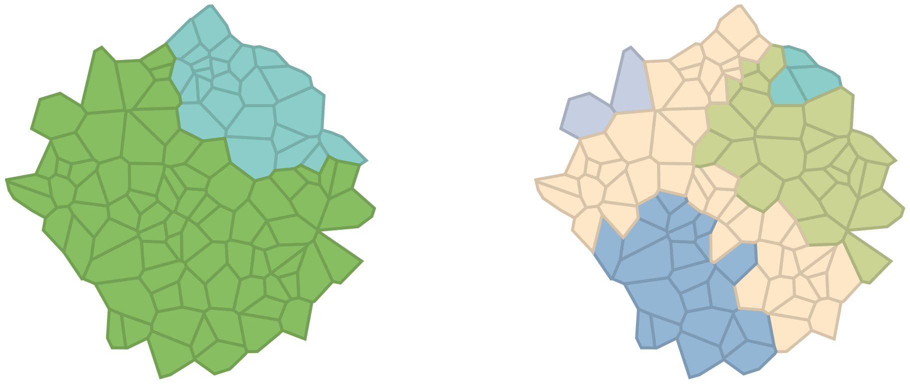

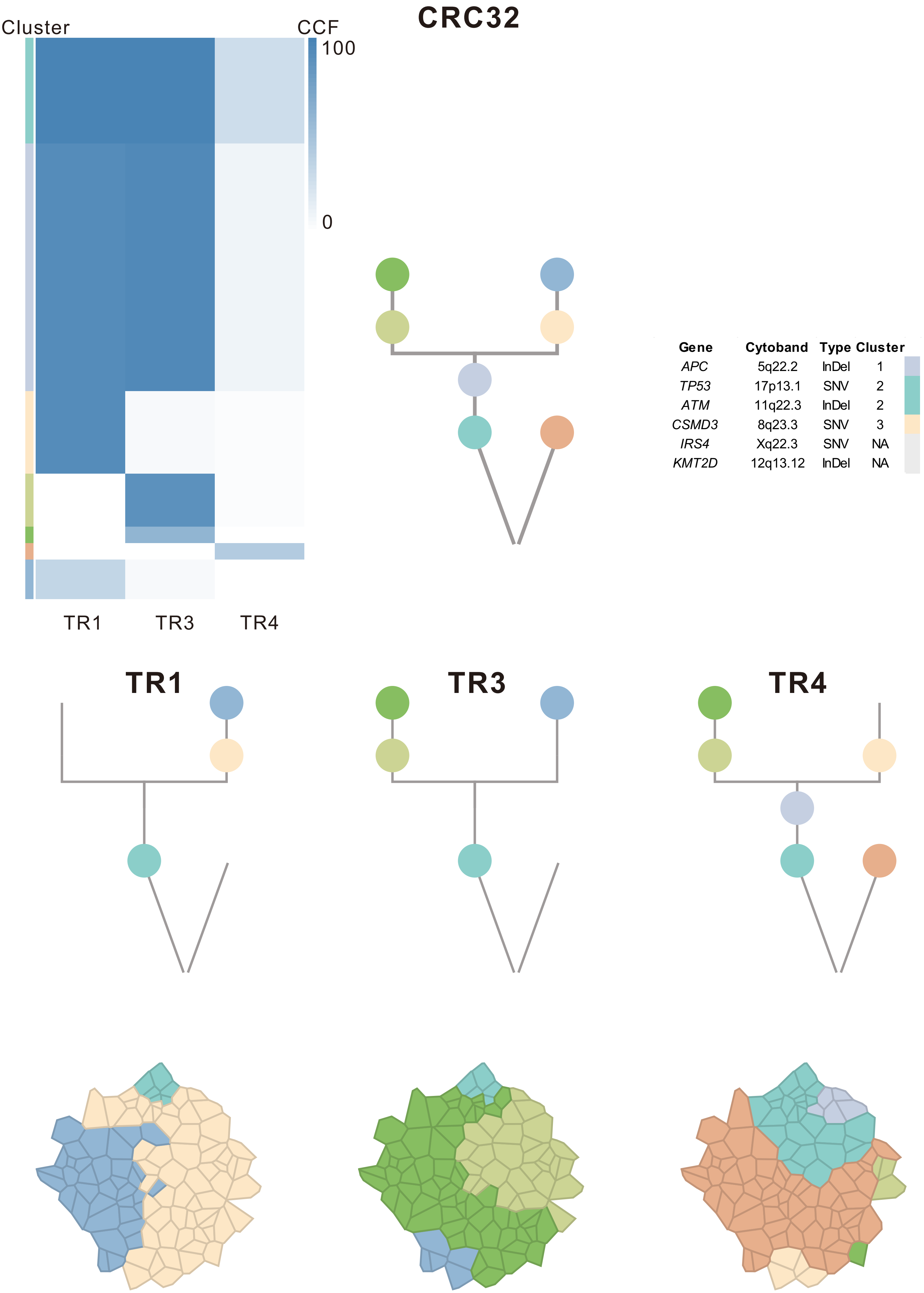
CRC32
Cluster
CCF
100
0
TR1
TR3
TR4
Gene Cytoband Type Cluster
APC 5q22.2 InDel 1
TP53 17p13.1 SNV 2
ATM 11q22.3 InDel 2
CSMD3 8q23.3 SNV 3
IRS4 Xq22.3 SNV NA
KMT2D 12q13.12 InDel NA
TR1
TR3
TR4

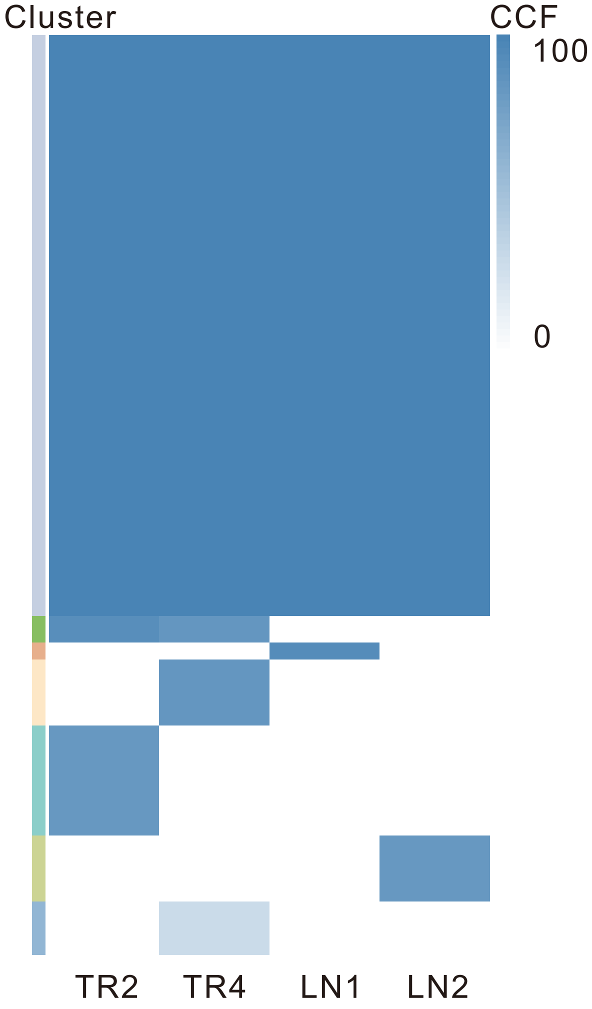

### CRC33

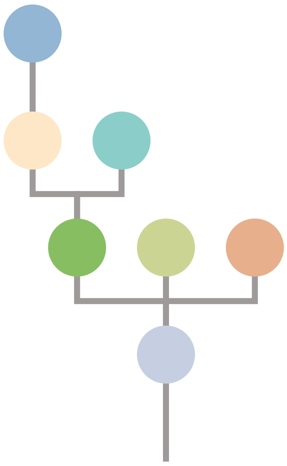

| Gene | Cytoband | Type | Cluster |
|---|---|---|---|
| *TP53* | 17p13.1 | SNV | 1 |
| *JAK3* | 19p13.11 | SNV | 1 |
| *APC* | 5q22.2 | InDel | 1 |
| *FLT3* | 13q12.2 | Amp | 1 |
| *FOXO1* | 13q14.11 | Amp | 1 |
| *CYSLTR2* | 13q14.2 | Amp | 1 |
| *SRC* | 20q11.23 | Amp | 1 |
| *MAFB* | 20q12 | Amp | 1 |
| *PLCG1* | 20q12 | Amp | 1 |
| *NFATC2* | 20q13.2 | Amp | 1 |
| *SALL4* | 20q13.2 | Amp | 1 |
| *GNAS* | 20q13.32 | Amp | 1 |
| *PTK6* | 20q13.33 | Amp | 1 |
| *NRG1* | 8p12 | SNV | 4 |
| *EGFR* | 7p11.2 | Amp | 4 |
| *IRS4* | Xq22.3 | InDel | NA |

## TR2

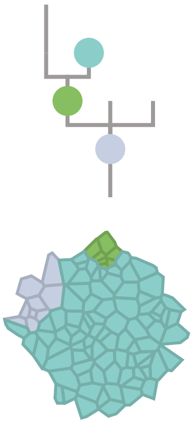

## TR4

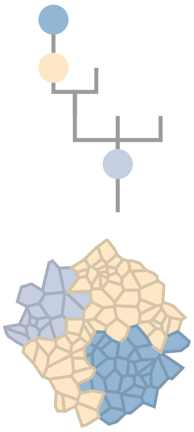

## LN1

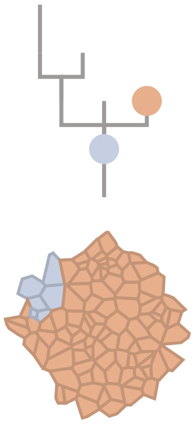

## LN2

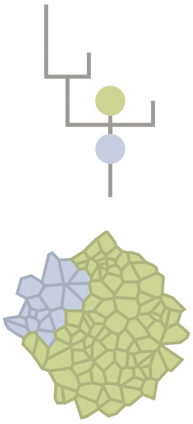

### CRC34

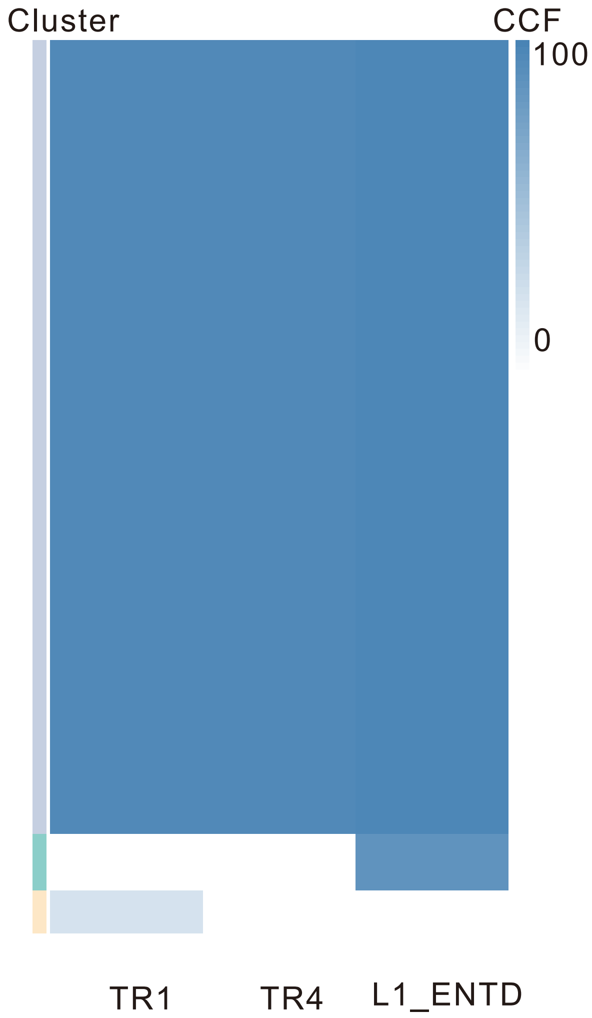

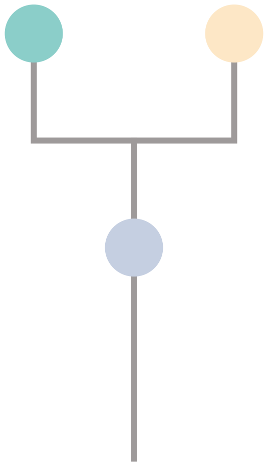

| Gene | Cytoband | Type | Cluster |
|---|---|---|---|
| * *APC* | 5q22.2 | SNV | 1 |
| * *APC* | 5q22.2 | SNV | 1 |
| *KRAS* | 12p12.1 | SNV | 1 |
| *TP53* | 17p13.1 | SNV | 1 |
| *CSMD3* | 8q23.3 | InDel | 1 |
| *AR* | Xq12 | SNV | NA |

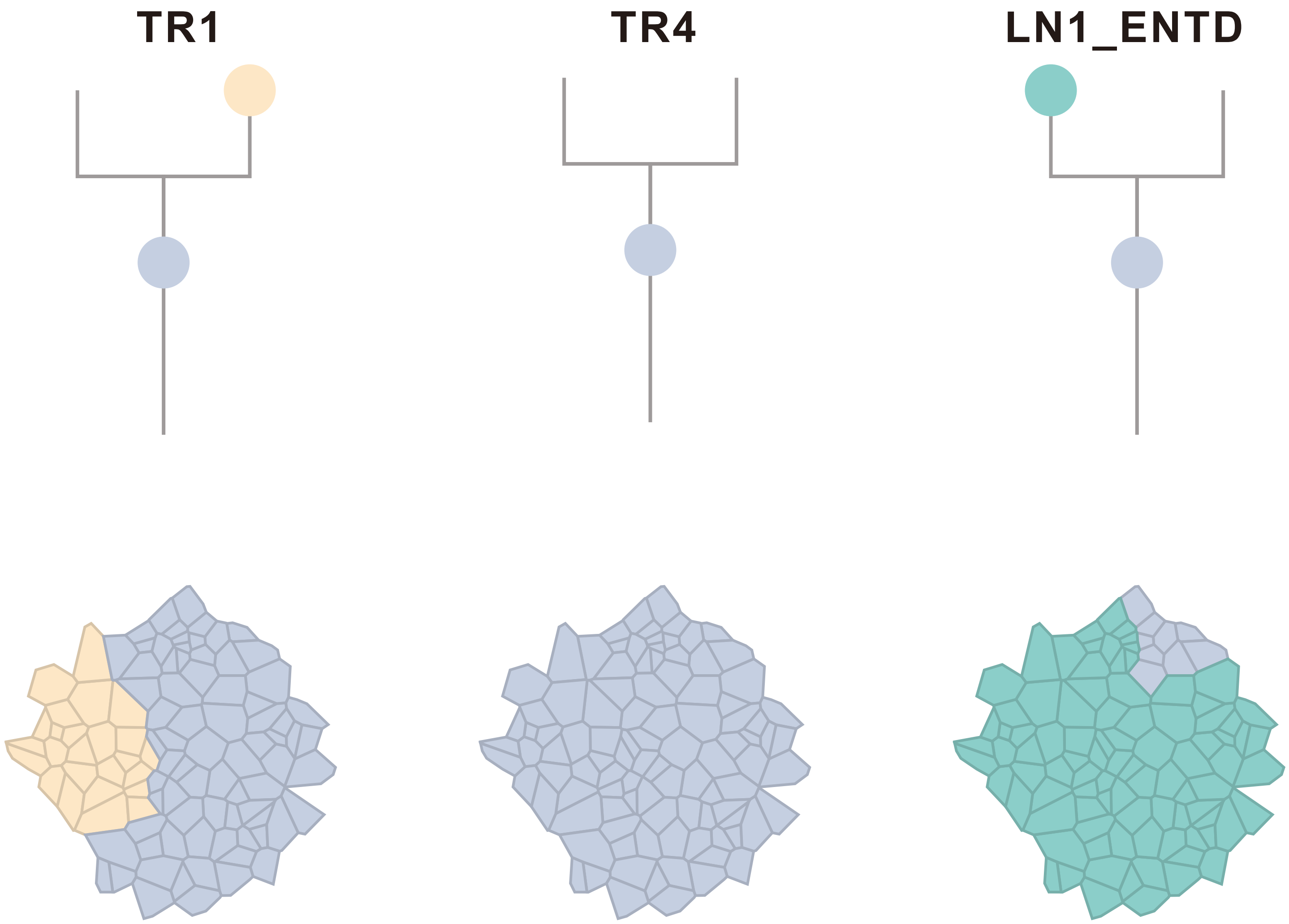

### CRC35

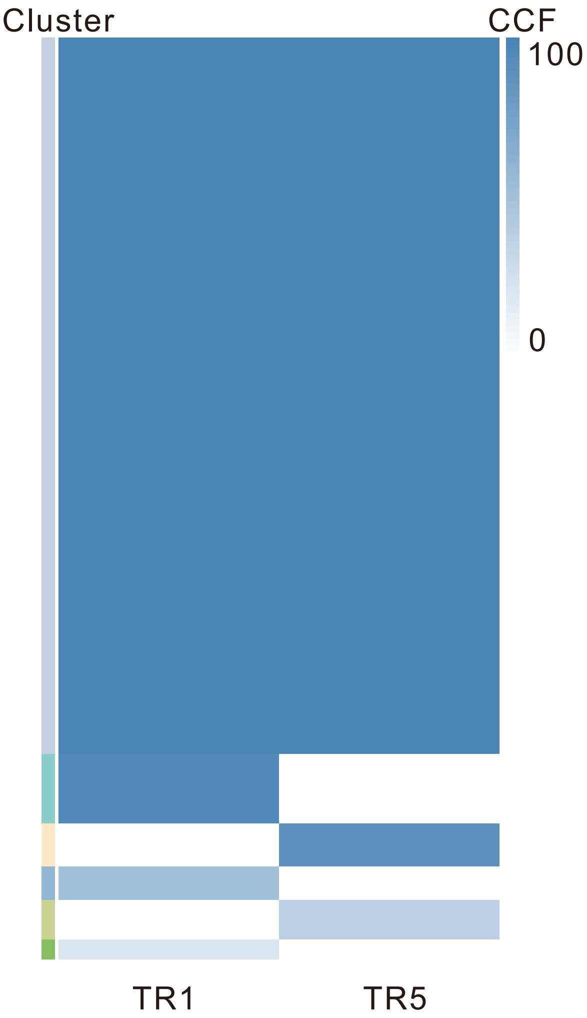

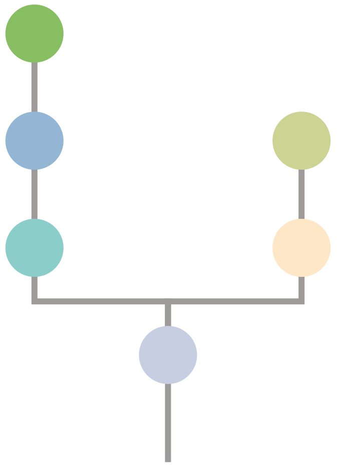

| Gene | Cytoband | Type | Cluster |
|---|---|---|---|
| *APC* | 5q22.2 | SNV | 1 |
| *ATM* | 11q22.3 | SNV | 1 |
| *KRAS* | 12p12.1 | SNV | 1 |
| *POLG* | 15q26.1 | SNV | 1 |
| *TP53* | 17p13.1 | SNV | 1 |
| *IL7R* | 5p13.2 | Amp | 1 |
| *CTNND2* | 5p15.2 | Amp | 1 |
| *TERT* | 5p15.33 | Amp | 1 |
| *SSX2* | Xp11.22 | Amp | 1 |
| *ARAF* | Xp11.23 | Amp | 1 |
| *GATA1* | Xp11.23 | Amp | 1 |
| *SSX1* | Xp11.23 | Amp | 1 |
| *SSX4* | Xp11.23 | Amp | 1 |
| *TFE3* | Xp11.23 | Amp | 1 |
| *WAS* | Xp11.23 | Amp | 1 |
| *KDM6A* | Xp11.3 | Amp | 1 |

## TR1

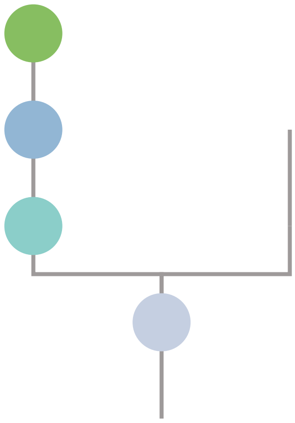

## TR5

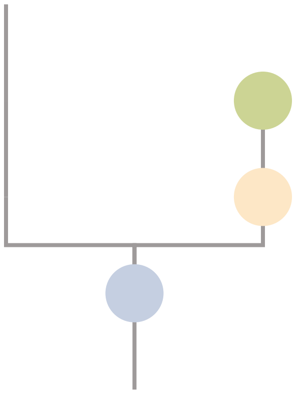

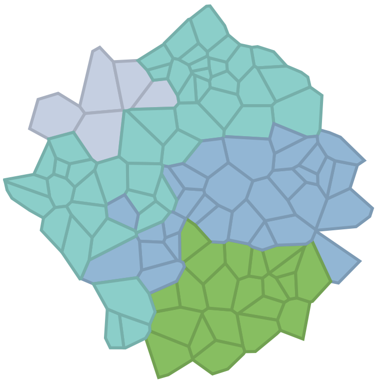

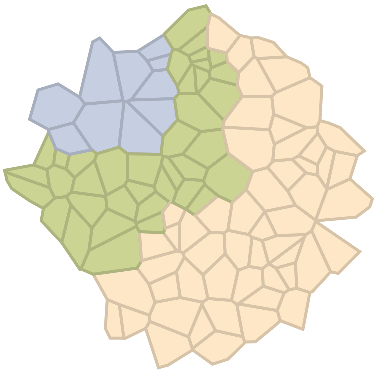

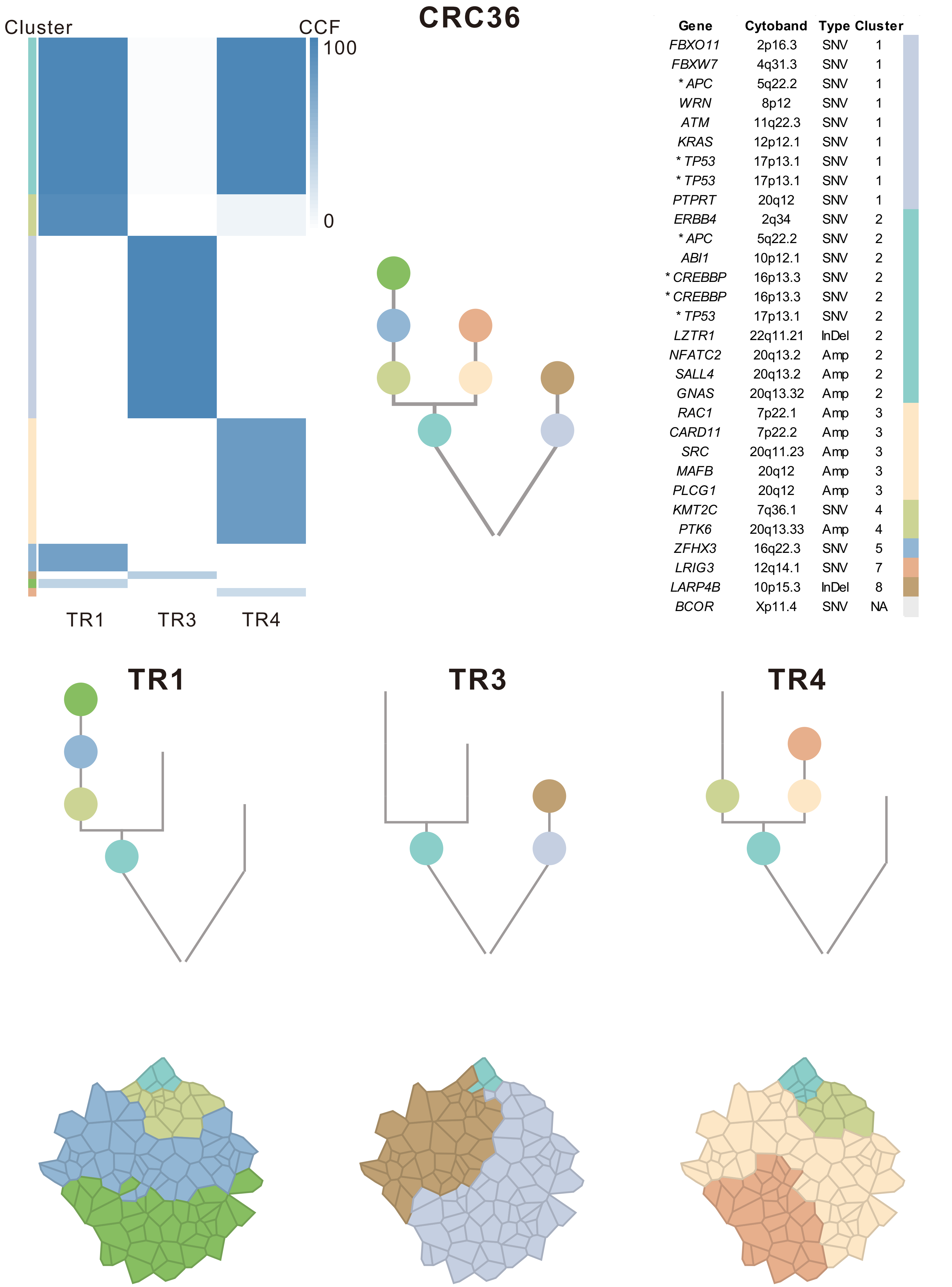
CRC36
Cluster
CCF
100
0
TR1
TR3
TR4
Gene Cytoband Type Cluster
FBXO11 2p16.3 SNV 1
FBXW7 4q31.3 SNV 1
* APC 5q22.2 SNV 1
WRN 8p12 SNV 1
ATM 11q22.3 SNV 1
KRAS 12p12.1 SNV 1
* TP53 17p13.1 SNV 1
* TP53 17p13.1 SNV 1
PTPRT 20q12 SNV 1
ERBB4 2q34 SNV 2
* APC 5q22.2 SNV 2
ABI1 10p12.1 SNV 2
* CREBBP 16p13.3 SNV 2
* CREBBP 16p13.3 SNV 2
* TP53 17p13.1 SNV 2
LZTR1 22q11.21 InDel 2
NFATC2 20q13.2 Amp 2
SALL4 20q13.2 Amp 2
GNAS 20q13.32 Amp 2
RAC1 7p22.1 Amp 3
CARD11 7p22.2 Amp 3
SRC 20q11.23 Amp 3
MAFB 20q12 Amp 3
PLCG1 20q12 Amp 3
KMT2C 7q36.1 SNV 4
PTK6 20q13.33 Amp 4
ZFHX3 16q22.3 SNV 5
LRIG3 12q14.1 SNV 7
LARP4B 10p15.3 InDel 8
BCOR Xp11.4 SNV NA
TR1
TR3
TR4

### CRC37

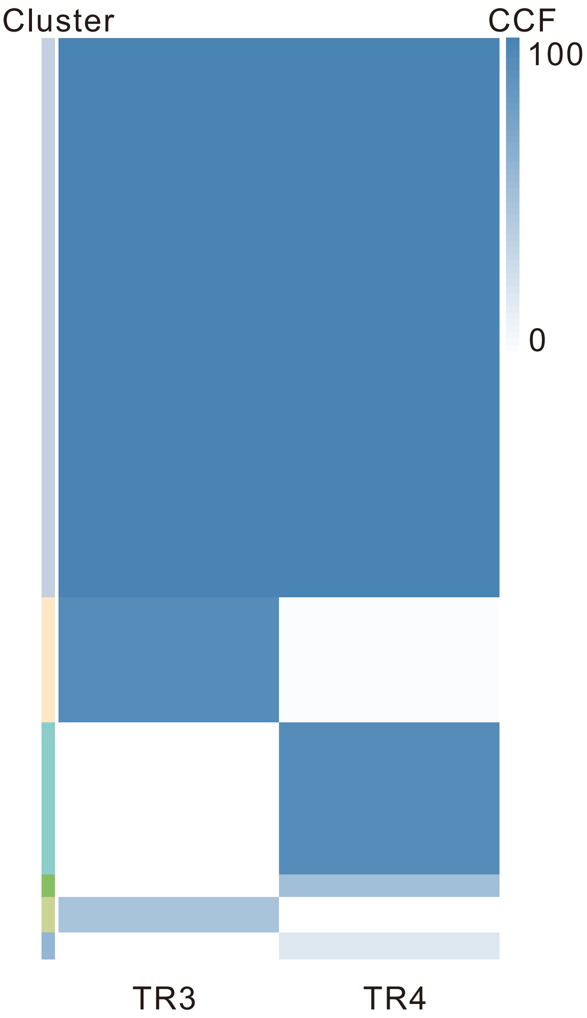

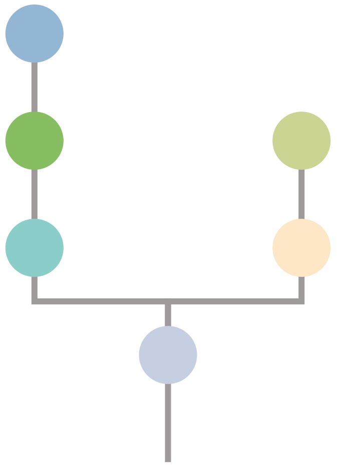

| Gene | Cytoband | Type | Cluster |
|---|---|---|---|
| * *TP53* | 17p13.1 | SNV | 1 |
| * *TP53* | 17p13.1 | SNV | 1 |
| * *APC* | 5q22.2 | InDel | 1 |
| *FLT3* | 13q12.2 | Amp | 1 |
| *SRC* | 20q11.23 | Amp | 1 |
| *MAFB* | 20q12 | Amp | 1 |
| *PLCG1* | 20q12 | Amp | 1 |
| *NFATC2* | 20q13.2 | Amp | 1 |
| *SALL4* | 20q13.2 | Amp | 1 |
| *GNAS* | 20q13.32 | Amp | 1 |
| *PTK6* | 20q13.33 | Amp | 1 |
| * *APC* | 5q22.2 | InDel | NA |

## TR3

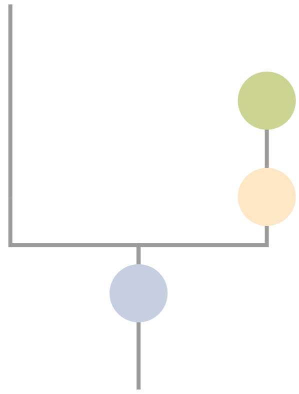

## TR4

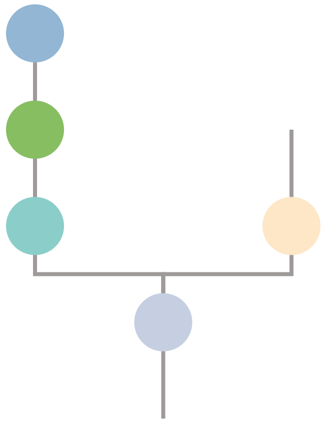

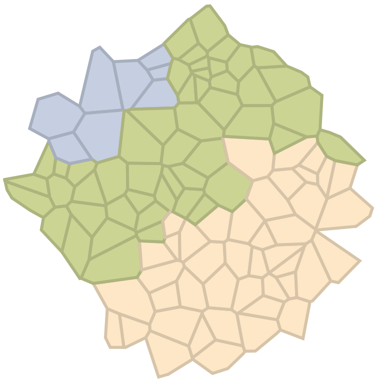

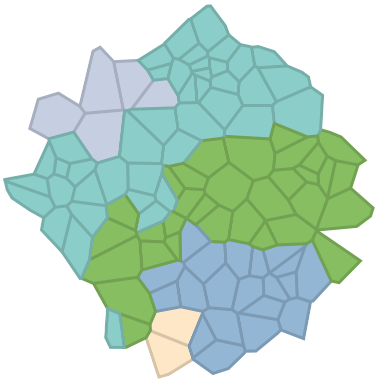

### CRC38

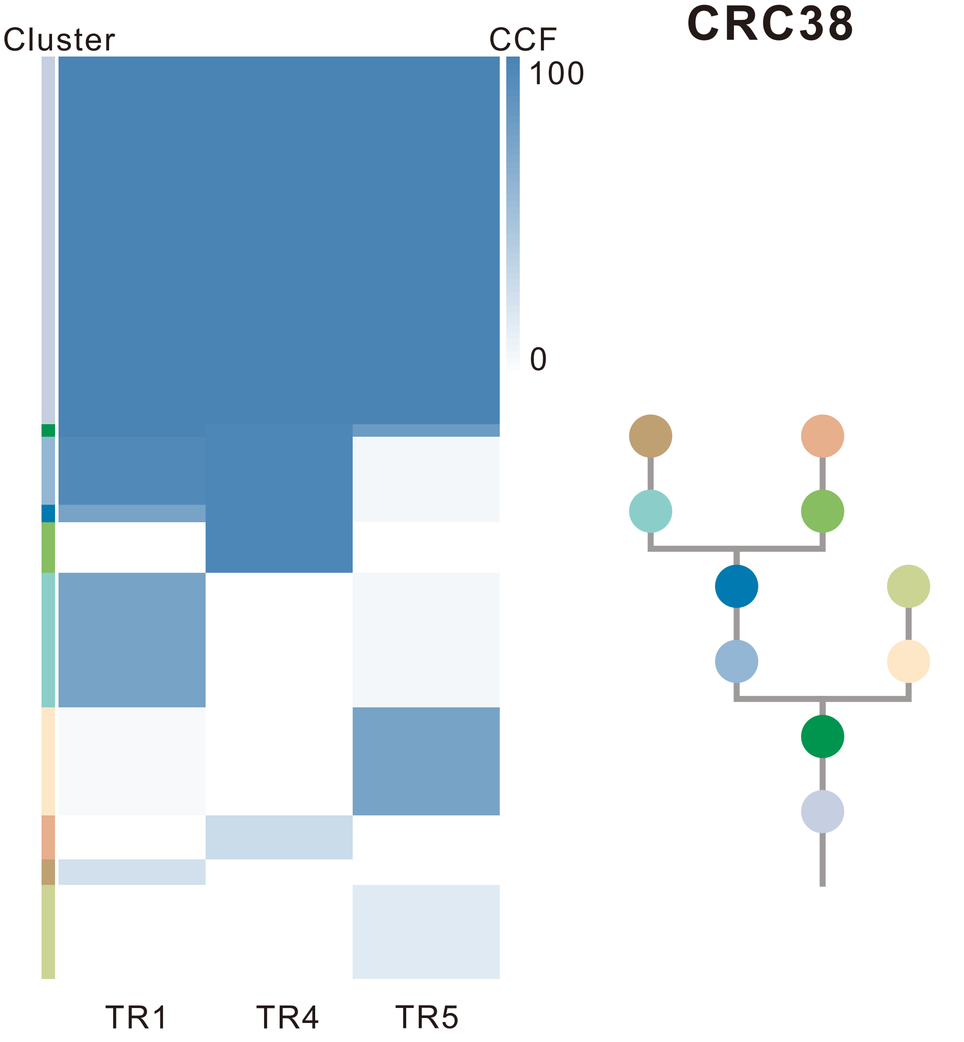

| Gene | Cytoband | Type | Cluster |
|---|---|---|---|
| *ATM* | 11q22.3 | SNV | 1 |
| *ZFHX3* | 16q22.3 | SNV | 1 |
| *TP53* | 17p13.1 | SNV | 1 |
| *SMAD4* | 18q21.2 | SNV | 1 |
| *CCND3* | 6p21.1 | Amp | 1 |
| *TFEB* | 6p21.1 | Amp | 1 |
| *IKBKB* | 8p11.21 | Amp | 1 |
| *KAT6A* | 8p11.21 | Amp | 1 |
| *RAD51B* | 14q24.1 | SNV | 2 |
| *NCOR1* | 17p11.2 | SNV | 4 |
| *KRAS* | 12p12.1 | SNV | 10 |

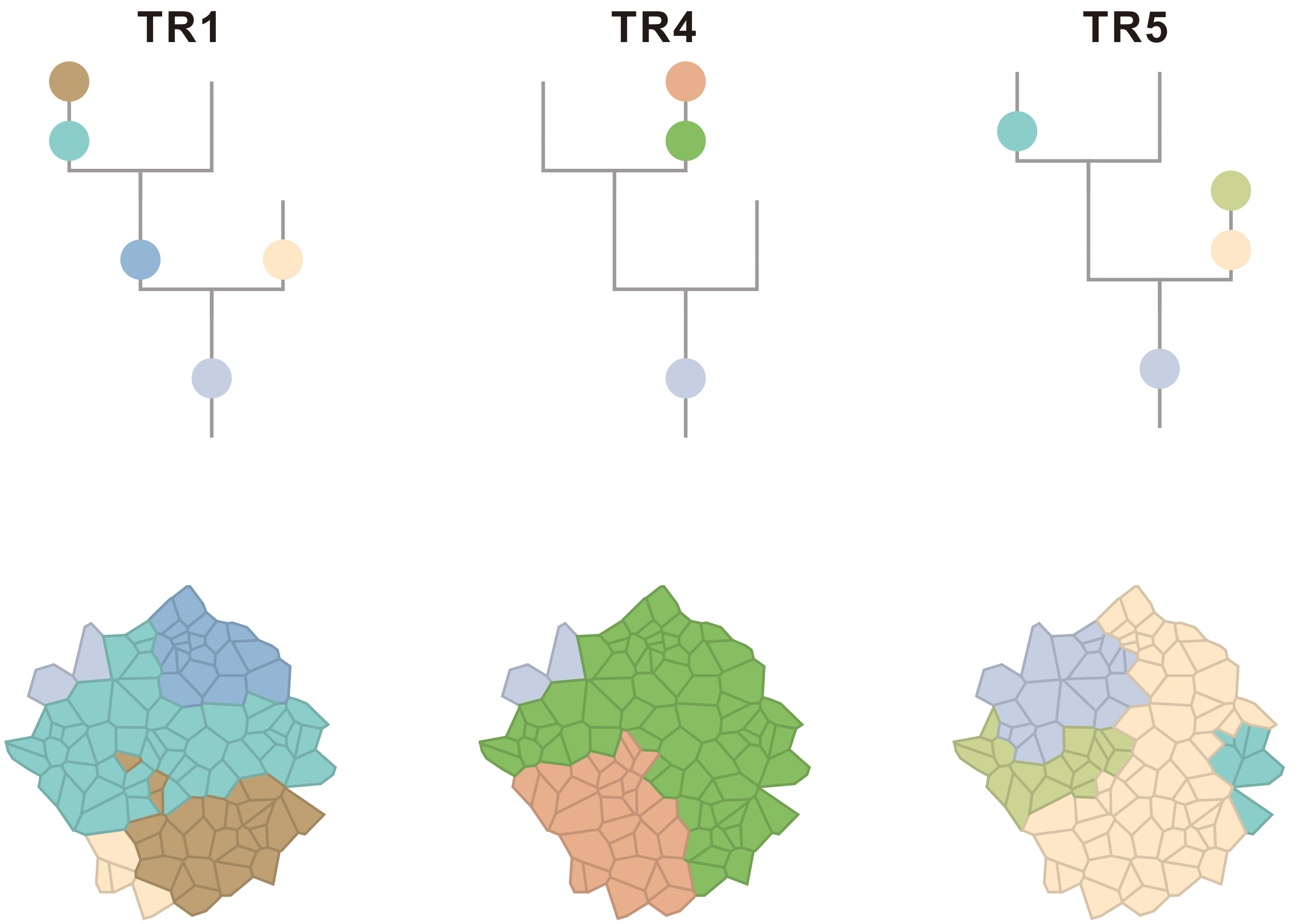

### CRC39

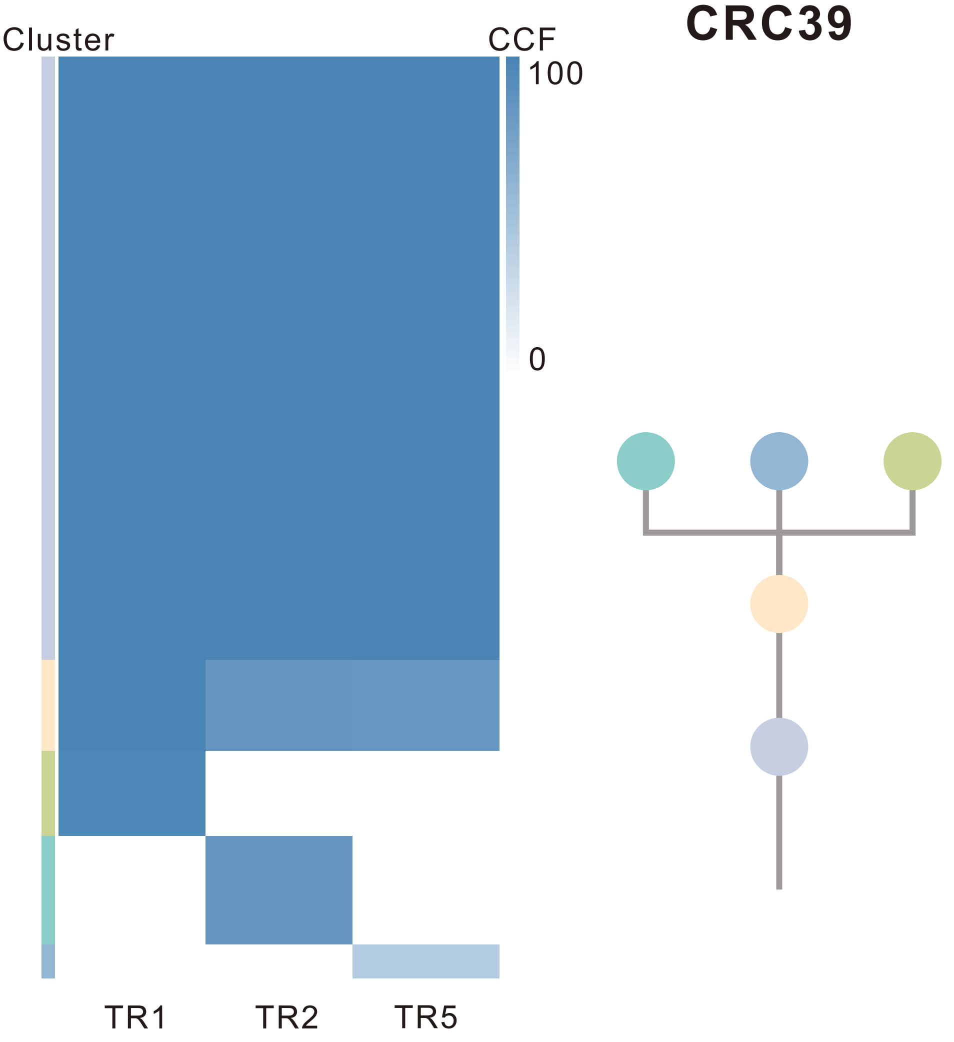

| Gene | Cytoband | Type | Cluster |
|---|---|---|---|
| *TP53* | 17p13.1 | SNV | 1 |
| *MAP3K1* | 5q11.2 | InDel | 1 |
| *ATM* | 11q22.3 | InDel | 1 |
| *FLT3* | 13q12.2 | Amp | 1 |
| *FOXO1* | 13q14.11 | Amp | 1 |
| *CYSLTR2* | 13q14.2 | Amp | 1 |
| *KRAS* | 12p12.1 | SNV | 3 |
| *FBXW7* | 4q31.3 | SNV | NA |
| *IRS4* | Xq22.3 | Amp | NA |
| *BCORL1* | Xq26.1 | Amp | NA |
| *ELF4* | Xq26.1 | Amp | NA |
| *GPC3* | Xq26.2 | Amp | NA |

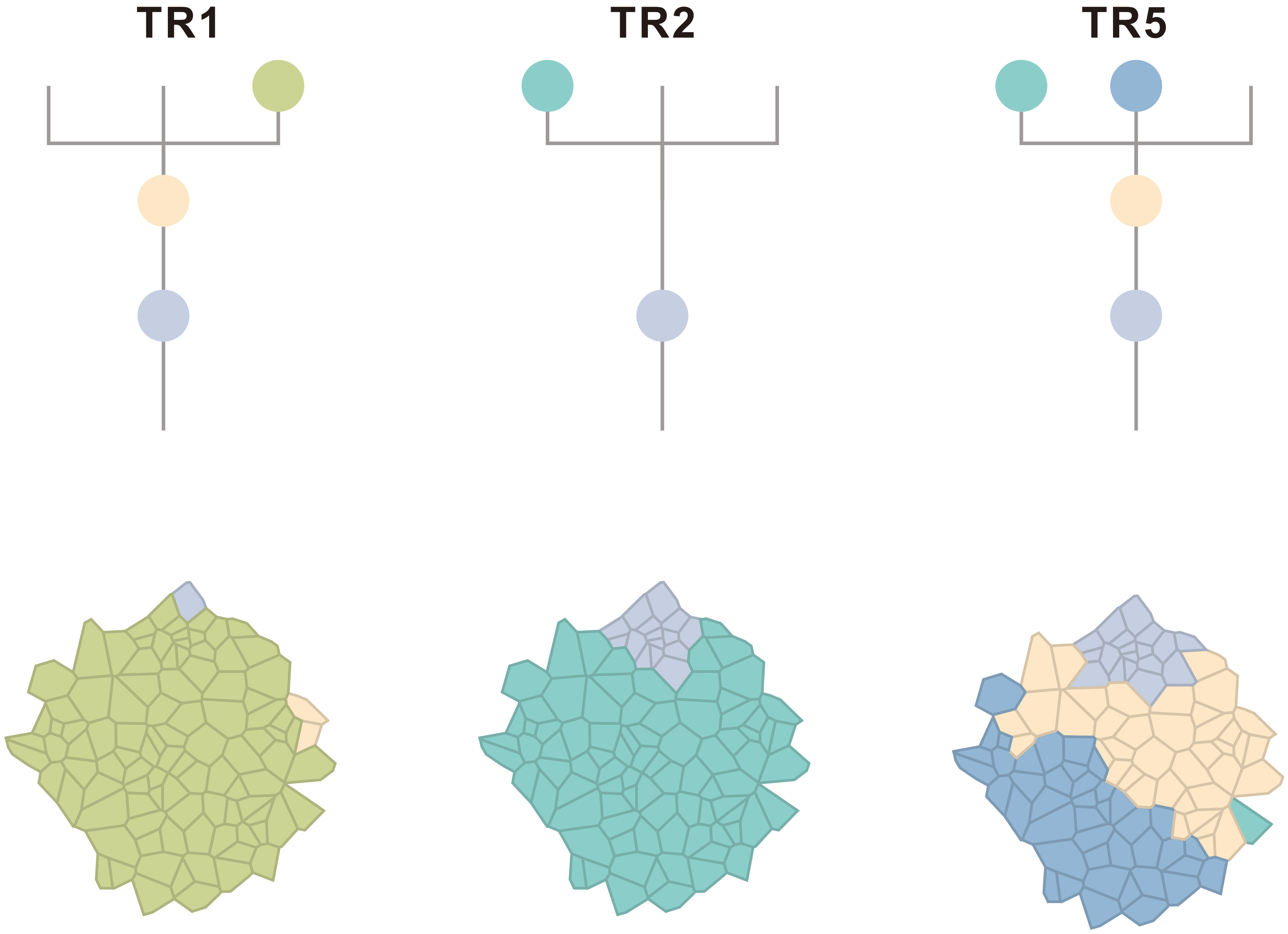

### CRC40

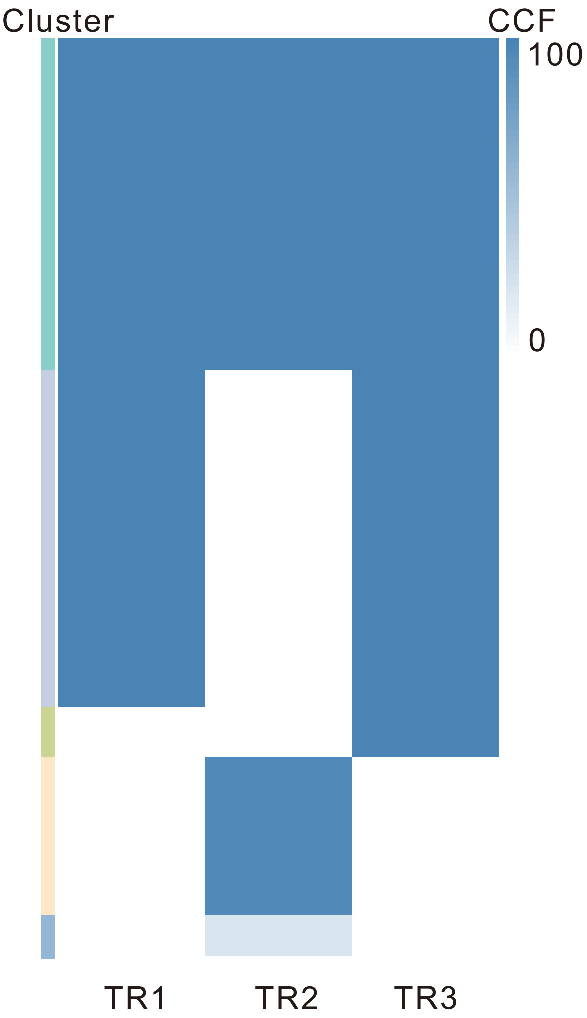

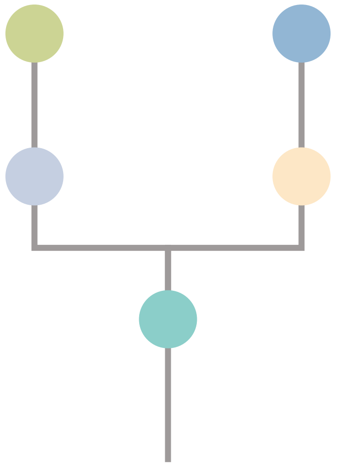

| Gene | Cytoband | Type | Cluster |
|---|---|---|---|
| *PIK3CA* | 3q26.32 | SNV | 1 |
| *TCF7L2* | 10q25.3 | SNV | 1 |
| *LZTR1* | 22q11.21 | InDel | 1 |
| * *APC* | 5q22.2 | SNV | 2 |
| * *APC* | 5q22.2 | SNV | 2 |
| *KRAS* | 12p12.1 | SNV | 2 |
| *ARID1A* | 1p36.11 | InDel | 2 |
| *ELF3* | 1q32.1 | InDel | 2 |
| *KMT2C* | 7q36.1 | SNV | 3 |
| *NDRG1* | 8q24.22 | SNV | 3 |
| *GNAS* | 20q13.32 | SNV | 3 |
| *MED12* | Xq13.1 | SNV | NA |
| *RECQL4* | 8q24.3 | Amp | NA |
| *RUNX1* | 21q22.12 | Del | NA |
| *SMARCB1* | 22q11.23 | Del | NA |
| *CHEK2* | 22q12.1 | Del | NA |
| *ZNRF3* | 22q12.1 | Del | NA |
| *NF2* | 22q12.2 | Del | NA |
| *MYH9* | 22q12.3 | Del | NA |
| *APOBEC3B* | 22q13.1 | Del | NA |
| *EP300* | 22q13.2 | Del | NA |
| *MKL1* | 22q13.2 | Del | NA |

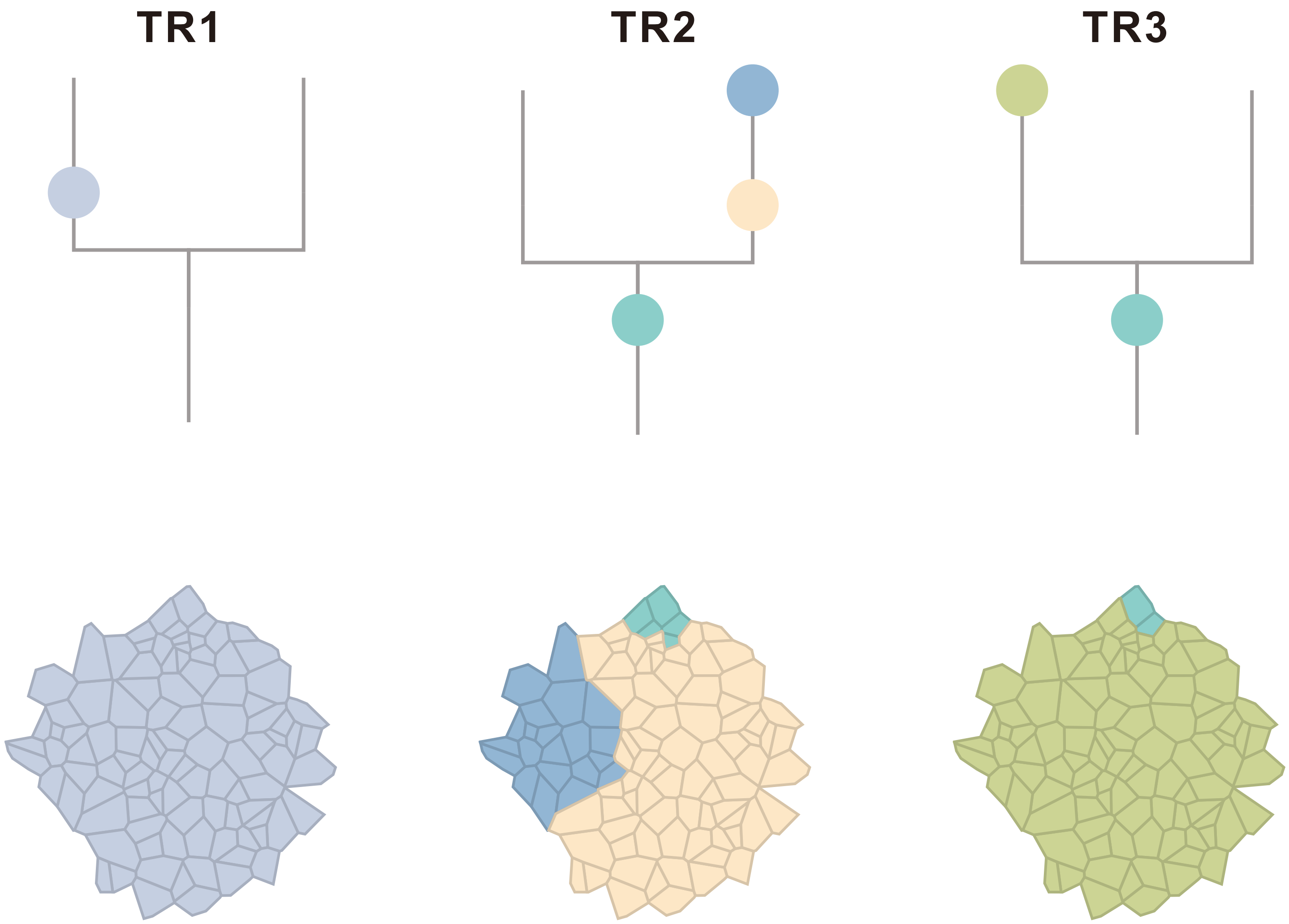

### CRC41

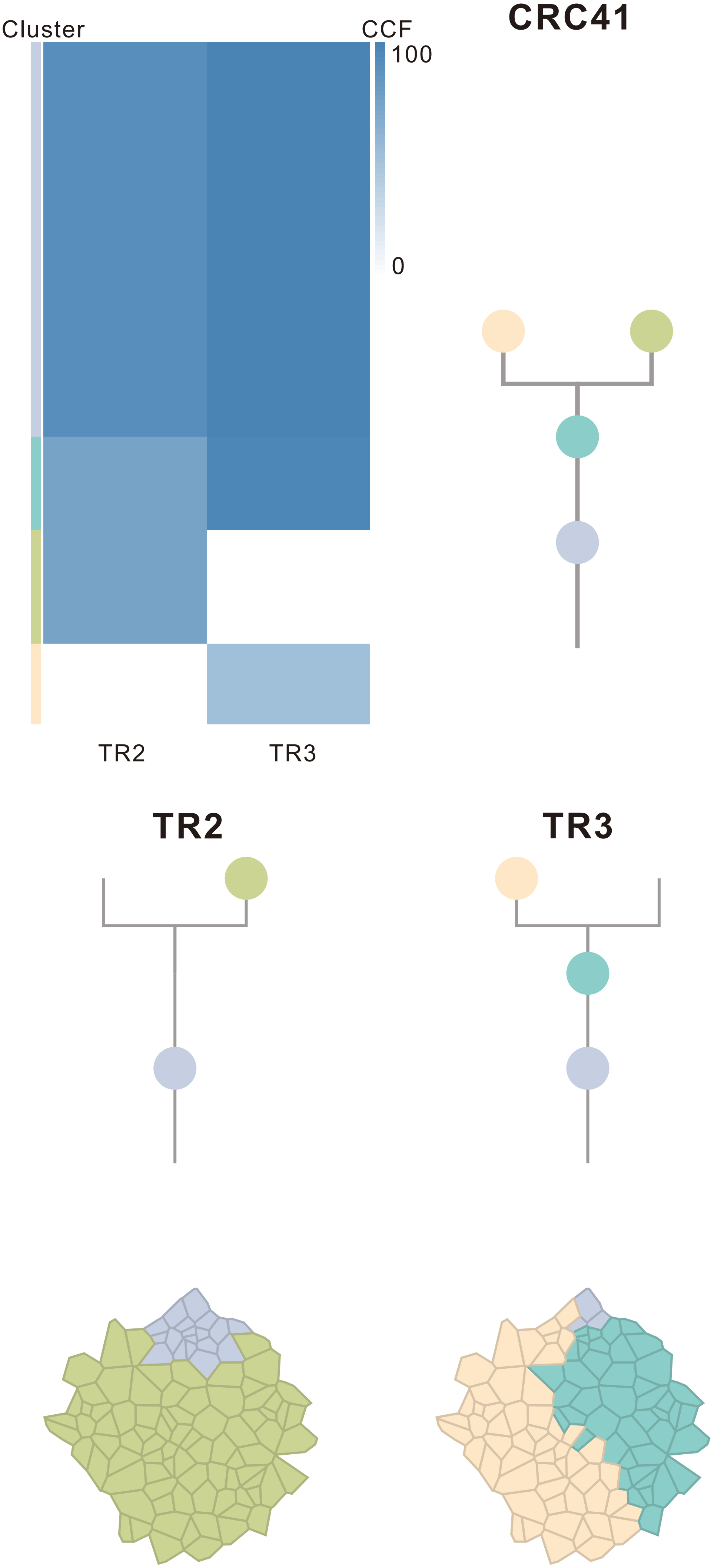

| Gene | Cytoband | Type | Cluster |
|---|---|---|---|
| *TCF7L2* | 10q25.3 | SNV | 1 |
| *TP53* | 17p13.1 | SNV | 1 |
| *FBXW7* | 4q31.3 | InDel | 1 |
| *LZTR1* | 22q11.21 | InDel | 1 |
| *IKBKB* | 8p11.21 | Amp | 1 |
| *KAT6A* | 8p11.21 | Amp | 1 |
| *FGFR1* | 8p11.23 | Amp | 1 |
| *ARID1A* | 1p36.11 | SNV | 2 |
| *SRC* | 20q11.23 | Amp | 3 |
| *MAFB* | 20q12 | Amp | 3 |
| *PLCG1* | 20q12 | Amp | 3 |
| *NFATC2* | 20q13.2 | Amp | 3 |
| *SALL4* | 20q13.2 | Amp | 3 |
| *GNAS* | 20q13.32 | Amp | 3 |
| *PTK6* | 20q13.33 | Amp | 3 |
| *RUNX1* | 21q22.12 | Del | 4 |
| *SMARCB1* | 22q11.23 | Del | 4 |
| *CHEK2* | 22q12.1 | Del | 4 |
| *ZNRF3* | 22q12.1 | Del | 4 |
| *NF2* | 22q12.2 | Del | 4 |
| *MYH9* | 22q12.3 | Del | 4 |
| *APOBEC3B* | 22q13.1 | Del | 4 |
| *EP300* | 22q13.2 | Del | 4 |
| *MKL1* | 22q13.2 | Del | 4 |
| *KDM5C* | Xp11.22 | Del | 4 |
| *SMC1A* | Xp11.22 | Del | 4 |
| *GATA1* | Xp11.23 | Del | 4 |
| *RBM10* | Xp11.23 | Del | 4 |
| *KDM6A* | Xp11.3 | Del | 4 |
| *BCOR* | Xp11.4 | Del | 4 |
| *DDX3X* | Xp11.4 | Del | 4 |
| *ZRSR2* | Xp22.2 | Del | 4 |
| *FOXO4* | Xq13.1 | Del | 4 |
| *MED12* | Xq13.1 | Del | 4 |
| *ZMYM3* | Xq13.1 | Del | 4 |
| *ATRX* | Xq21.1 | Del | 4 |
| *BTK* | Xq22.1 | Del | 4 |
| *IRS4* | Xq22.3 | Del | 4 |
| *STAG2* | Xq25 | Del | 4 |
| *BCORL1* | Xq26.1 | Del | 4 |
| *ELF4* | Xq26.1 | Del | 4 |
| *GPC3* | Xq26.2 | Del | 4 |
| *PHF6* | Xq26.2 | Del | 4 |
| *ATP2B3* | Xq28 | Del | 4 |
| *RPL10* | Xq28 | Del | 4 |
| *AR* | Xq12 | SNV | NA |
| *FBXW7* | 4q31.3 | InDel | NA |

### CRC42

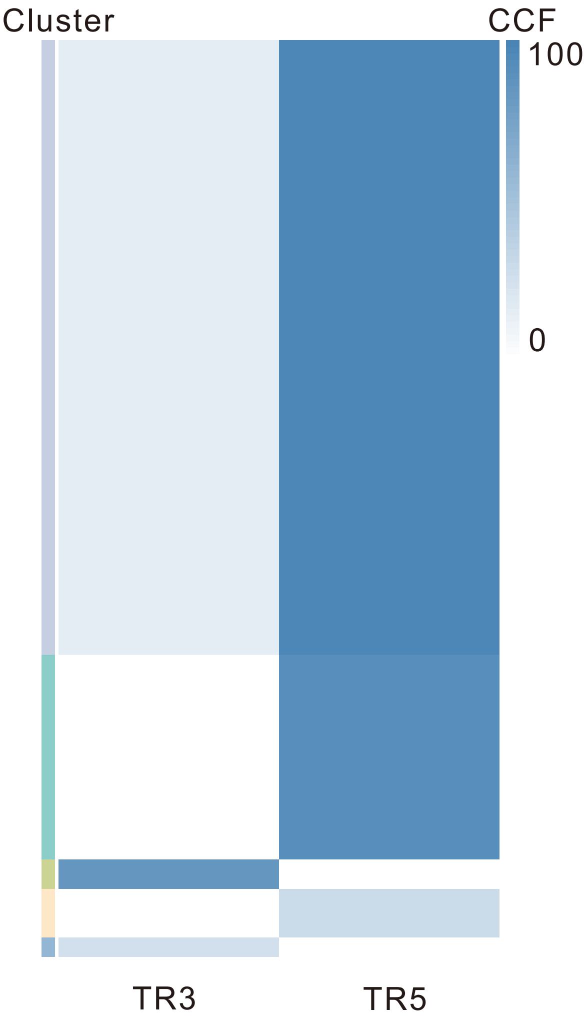

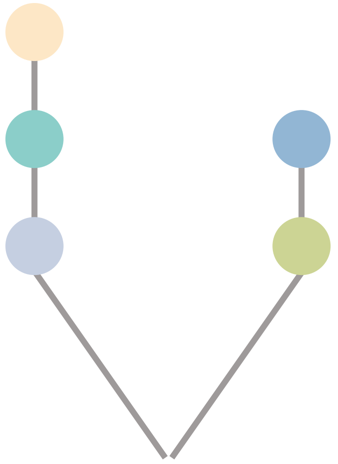

| Gene | Cytoband | Type | Cluster |
|---|---|---|---|
| *TP53* | 17p13.1 | SNV | 1 |
| *DNM2* | 19p13.2 | SNV | 1 |
| *PTPRT* | 20q12 | SNV | 1 |
| *PABPC1* | 8q22.3 | InDel | 1 |
| *FAT1* | 4q35.2 | Del | 1 |
| *EXT2* | 11p11.2 | SNV | 3 |

## TR3

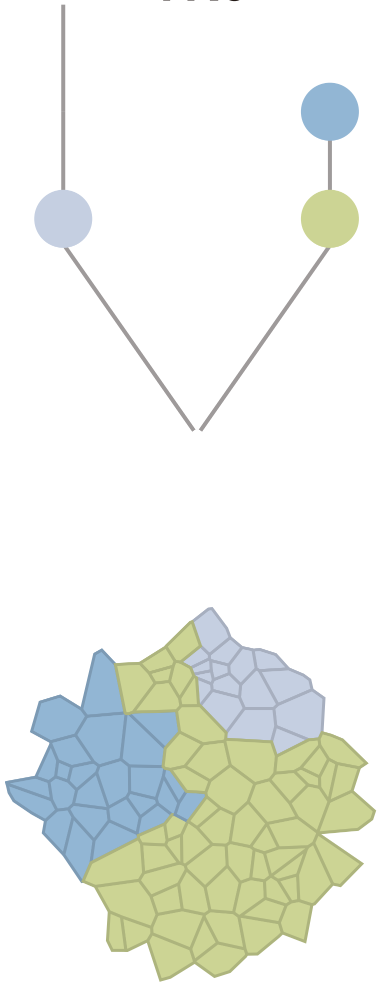

## TR5

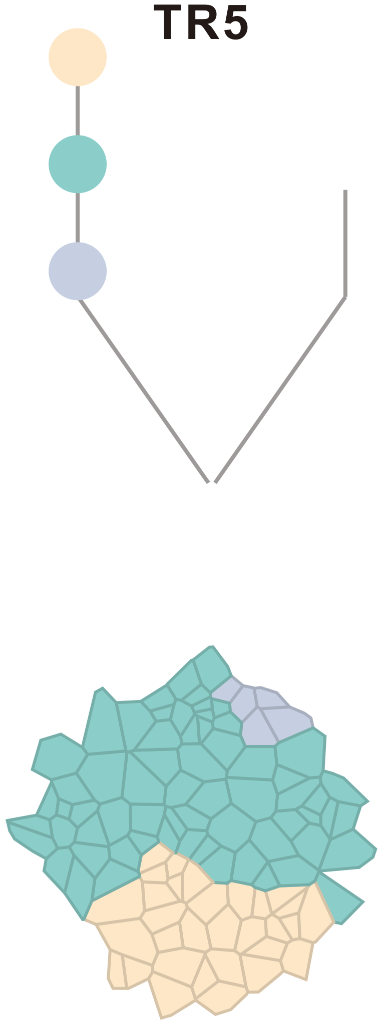

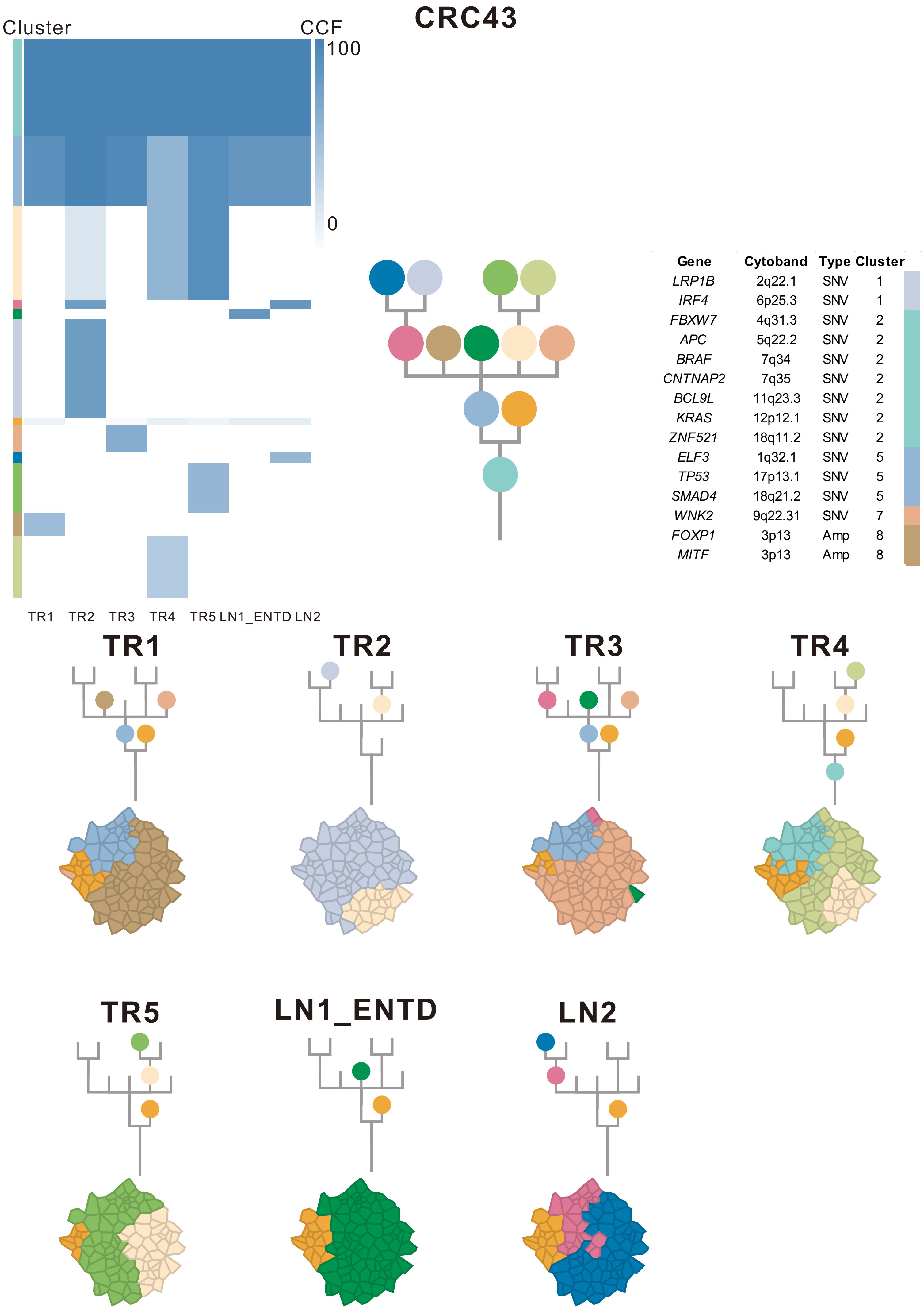

| Gene | Cytoband | Type | Cluster |
|---|---|---|---|
| *LRP1B* | 2q22.1 | SNV | 1 |
| *IRF4* | 6p25.3 | SNV | 1 |
| *FBXW7* | 4q31.3 | SNV | 2 |
| *APC* | 5q22.2 | SNV | 2 |
| *BRAF* | 7q34 | SNV | 2 |
| *CNTNAP2* | 7q35 | SNV | 2 |
| *BCL9L* | 11q23.3 | SNV | 2 |
| *KRAS* | 12p12.1 | SNV | 2 |
| *ZNF521* | 18q11.2 | SNV | 2 |
| *ELF3* | 1q32.1 | SNV | 5 |
| *TP53* | 17p13.1 | SNV | 5 |
| *SMAD4* | 18q21.2 | SNV | 5 |
| *WNK2* | 9q22.31 | SNV | 7 |
| *FOXP1* | 3p13 | Amp | 8 |
| *MITF* | 3p13 | Amp | 8 |

### CRC44

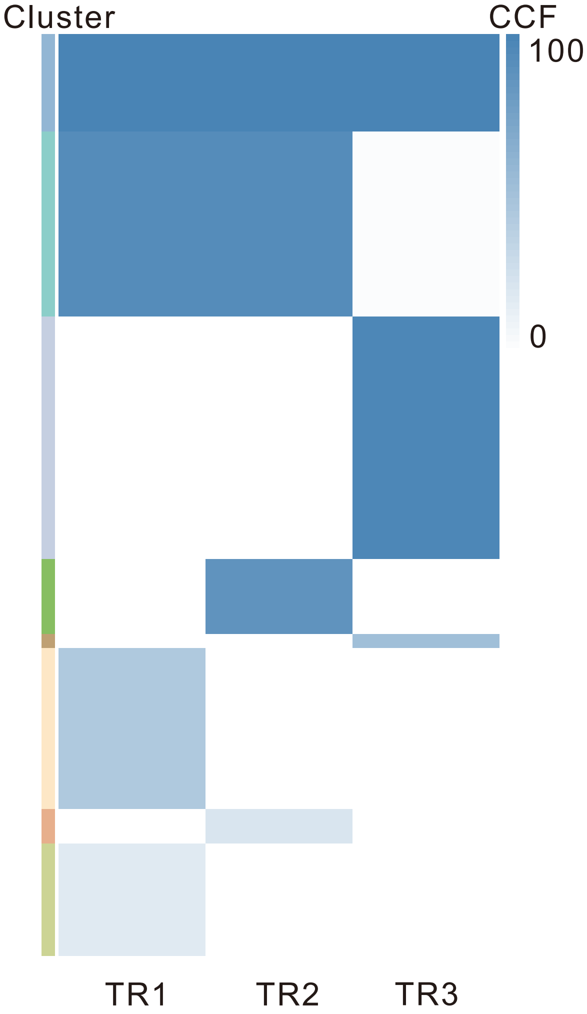

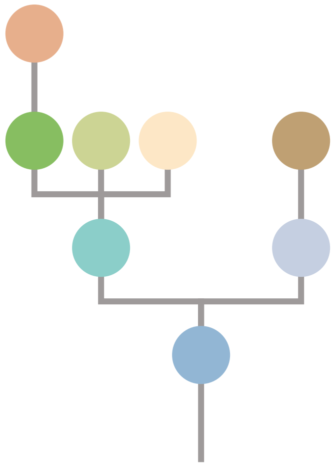

| Gene | Cytoband | Type | Cluster |
|---|---|---|---|
| * *FBXW7* | 4q31.3 | SNV | 1 |
| *TP53* | 17p13.1 | SNV | 1 |
| *FLT3* | 13q12.2 | Amp | 1 |
| *FOXO1* | 13q14.11 | Amp | 1 |
| *CYSLTR2* | 13q14.2 | Amp | 1 |
| *ARID1A* | 1p36.11 | SNV | 2 |
| * *FBXW7* | 4q31.3 | SNV | 2 |
| *KRAS* | 12p12.1 | SNV | 2 |
| *GNAS* | 20q13.32 | SNV | 4 |
| *MUTYH* | 1p34.1 | SNV | 5 |
| *APC* | 5q22.2 | SNV | 5 |
| *ACVR2A* | 2q22.3 | SNV | 7 |
| *LZTR1* | 22q11.21 | InDel | NA |
| *AMER1* | Xq11.2 | SNV | NA |

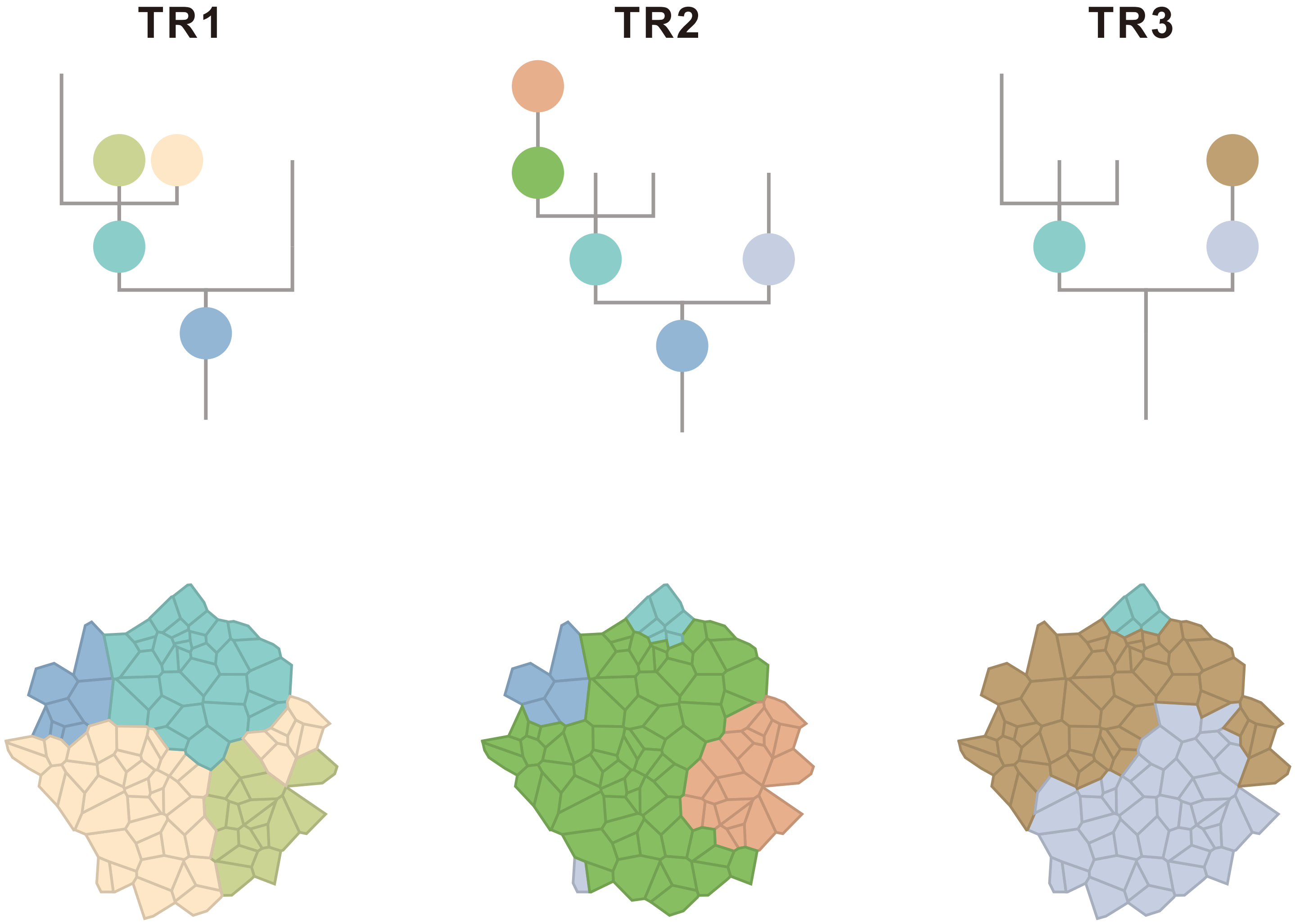

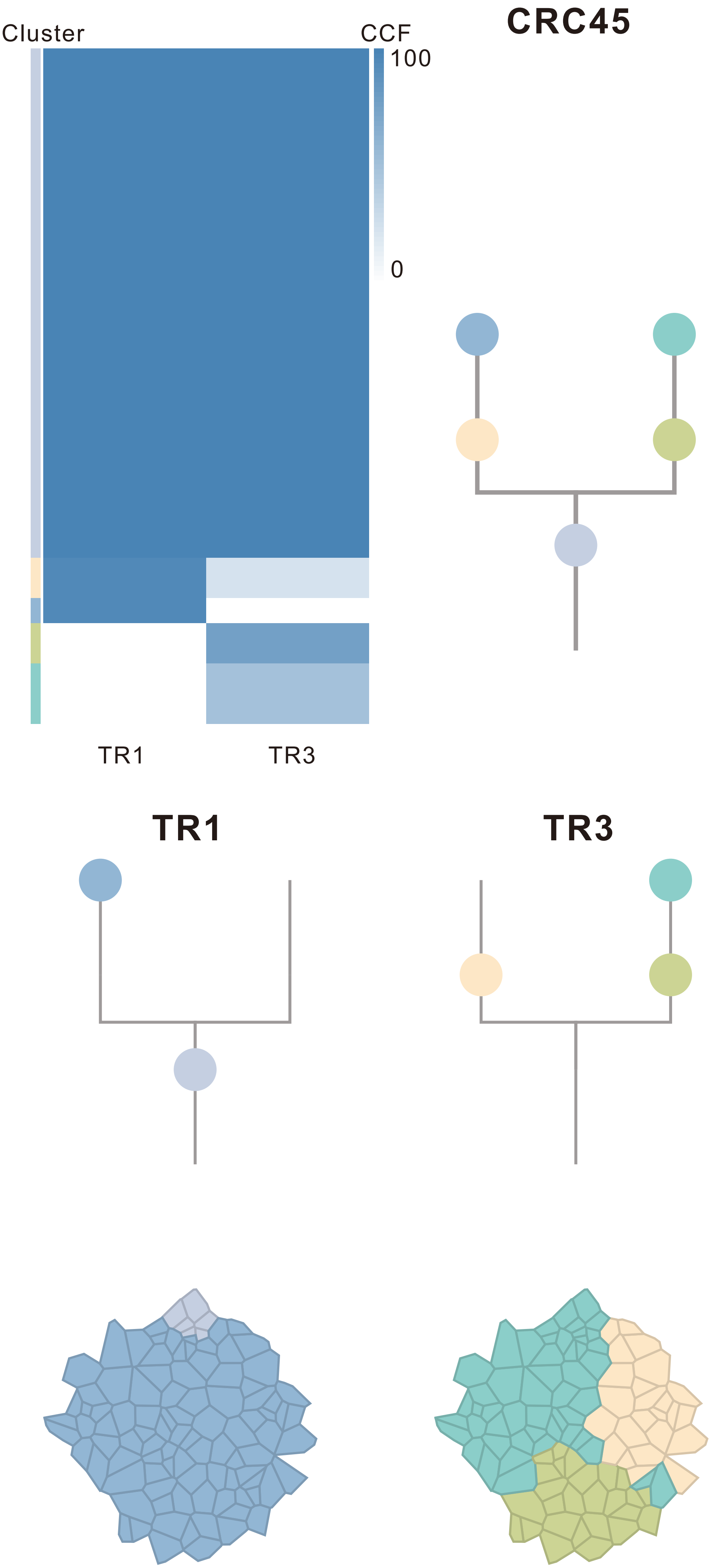

| Gene | Cytoband | Type | Cluster |
|---|---|---|---|
| *APC* | 5q22.2 | SNV | 1 |
| *POT1* | 7q31.33 | SNV | 1 |
| *TP53* | 17p13.1 | SNV | 1 |
| *FGFR1* | 8p11.23 | Amp | 4 |

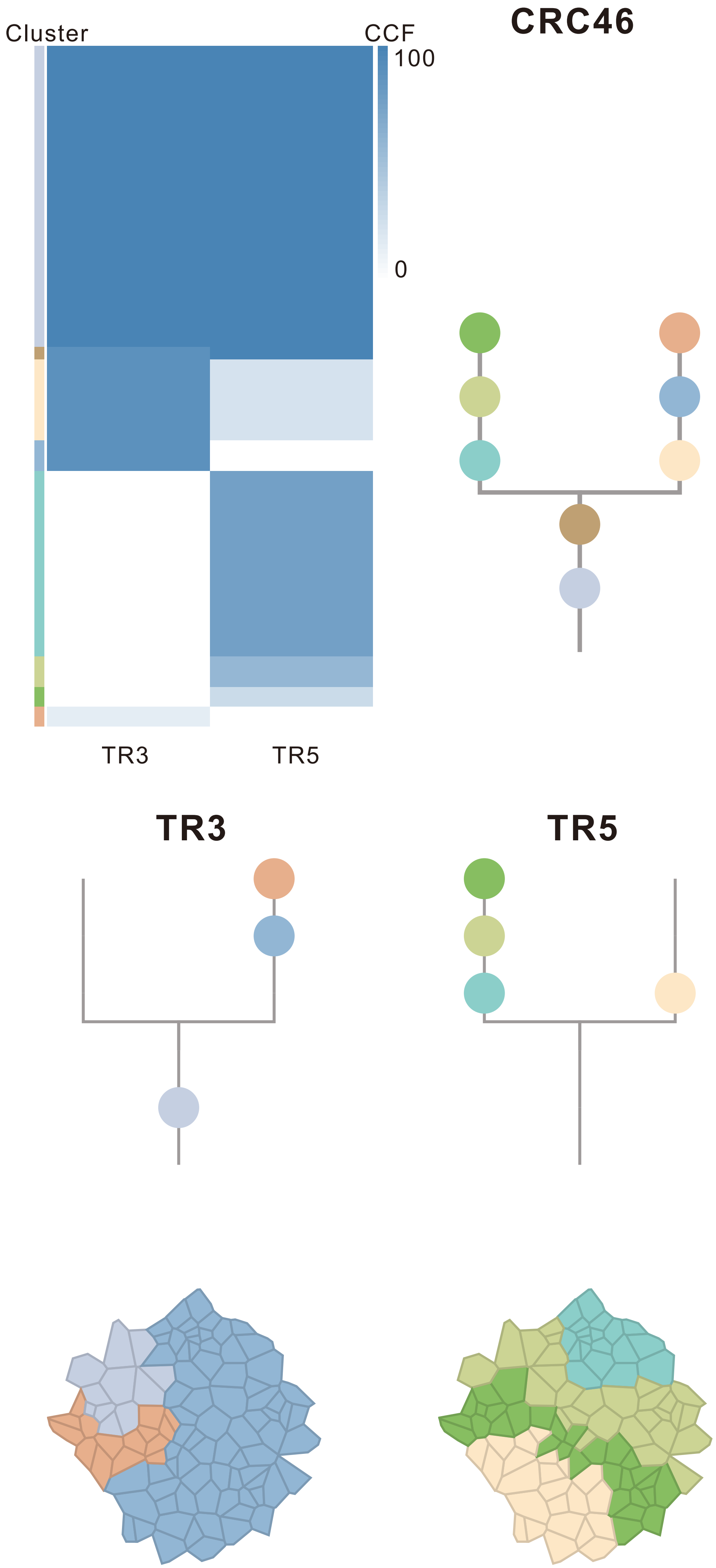

| Gene | Cytoband | Type | Cluster |
|---|---|---|---|
| *APC* | 5q22.2 | SNV | 1 |
| * *ATM* | 11q22.3 | SNV | 1 |
| *TBX3* | 12q24.21 | SNV | 1 |
| *ARID1A* | 1p36.11 | InDel | 1 |
| *NFE2L2* | 2q31.2 | InDel | 1 |
| *DDB2* | 11p11.2 | InDel | 1 |
| * *ATM* | 11q22.3 | InDel | 1 |
| *PLAG1* | 8q12.1 | Amp | 1 |
| *PREX2* | 8q13.2 | Amp | 1 |
| *NCOA2* | 8q13.3 | Amp | 1 |
| *HEY1* | 8q21.13 | Amp | 1 |
| *RUNX1T1* | 8q21.3 | Amp | 1 |
| *CDH17* | 8q22.1 | Amp | 1 |
| *PABPC1* | 8q22.3 | Amp | 1 |
| *UBR5* | 8q22.3 | Amp | 1 |
| *RAD21* | 8q24.11 | Amp | 1 |
| *MYC* | 8q24.21 | Amp | 1 |
| *RECQL4* | 8q24.3 | Amp | 1 |
| *FLT3* | 13q12.2 | Amp | 1 |
| *IRS4* | Xq22.3 | Amp | 1 |
| *BCORL1* | Xq26.1 | Amp | 1 |
| *ELF4* | Xq26.1 | Amp | 1 |
| *GPC3* | Xq26.2 | Amp | 1 |
| *EGFR* | 7p11.2 | Amp | 2 |
| *HNRNPA2B1* | 7p15.2 | Amp | 2 |
| *HOXA11* | 7p15.2 | Amp | 2 |
| *HOXA13* | 7p15.2 | Amp | 2 |
| *HOXA9* | 7p15.2 | Amp | 2 |
| *MACC1* | 7p21.1 | Amp | 2 |
| *ETV1* | 7p21.2 | Amp | 2 |
| *RAC1* | 7p22.1 | Amp | 2 |
| *CARD11* | 7p22.2 | Amp | 2 |
| *SSX2* | Xp11.22 | Amp | 2 |
| *ARAF* | Xp11.23 | Amp | 2 |
| *GATA1* | Xp11.23 | Amp | 2 |
| *SSX1* | Xp11.23 | Amp | 2 |
| *SSX4* | Xp11.23 | Amp | 2 |
| *TFE3* | Xp11.23 | Amp | 2 |
| *WAS* | Xp11.23 | Amp | 2 |
| *KDM6A* | Xp11.3 | Amp | 2 |
| *LRP1B* | 2q22.1 | SNV | 3 |
| *FOXO1* | 13q14.11 | Amp | 3 |
| *NTRK3* | 15q25.3 | SNV | 8 |
| *CYSLTR2* | 13q14.2 | Amp | NA |

### CRC47

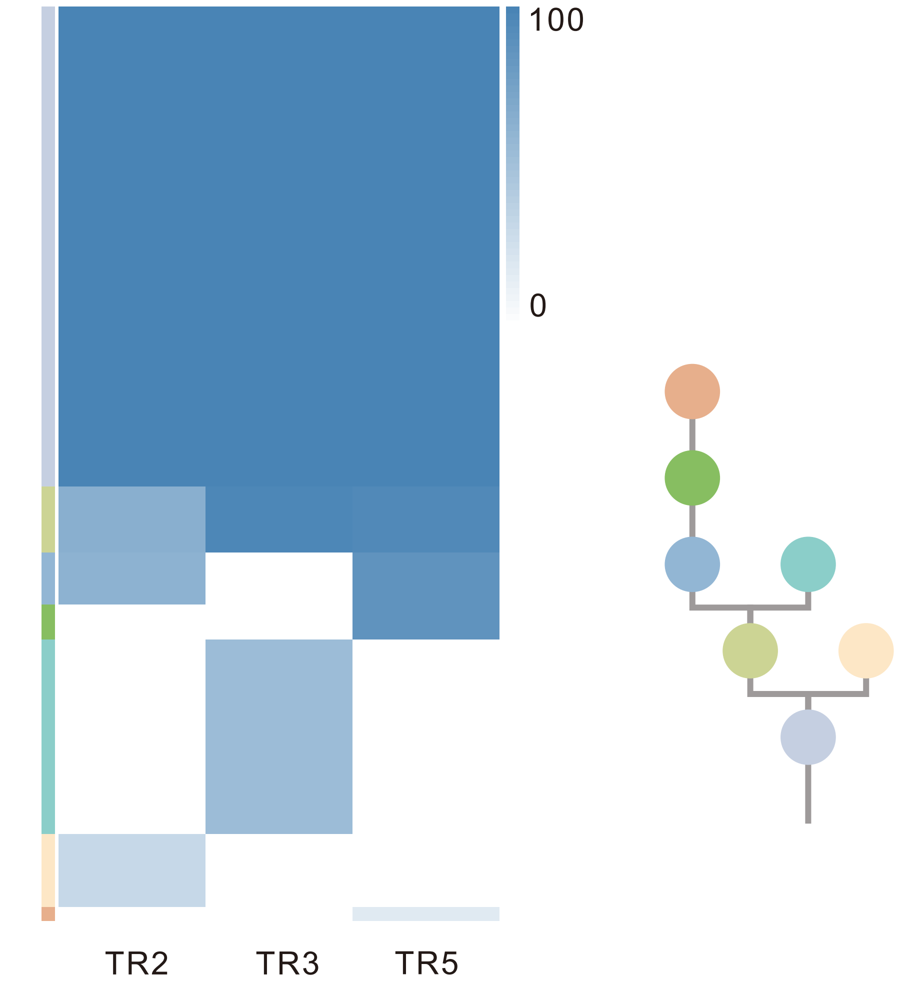

| Gene | Cytoband | Type | Cluster |
|---|---|---|---|
| *APC* | 5q22.2 | SNV | 1 |
| *TP53* | 17p13.1 | SNV | 1 |
| *MAFB* | 20q12 | Amp | 1 |
| *PLCG1* | 20q12 | Amp | 1 |
| *NFATC2* | 20q13.2 | Amp | 1 |
| *SALL4* | 20q13.2 | Amp | 1 |
| *GNAS* | 20q13.32 | Amp | 1 |
| *PTK6* | 20q13.33 | Amp | 1 |
| *SRC* | 20q11.23 | Amp | 2 |
| *PIK3CA* | 3q26.32 | SNV | 5 |
| *RBM10* | Xp11.23 | InDel | NA |

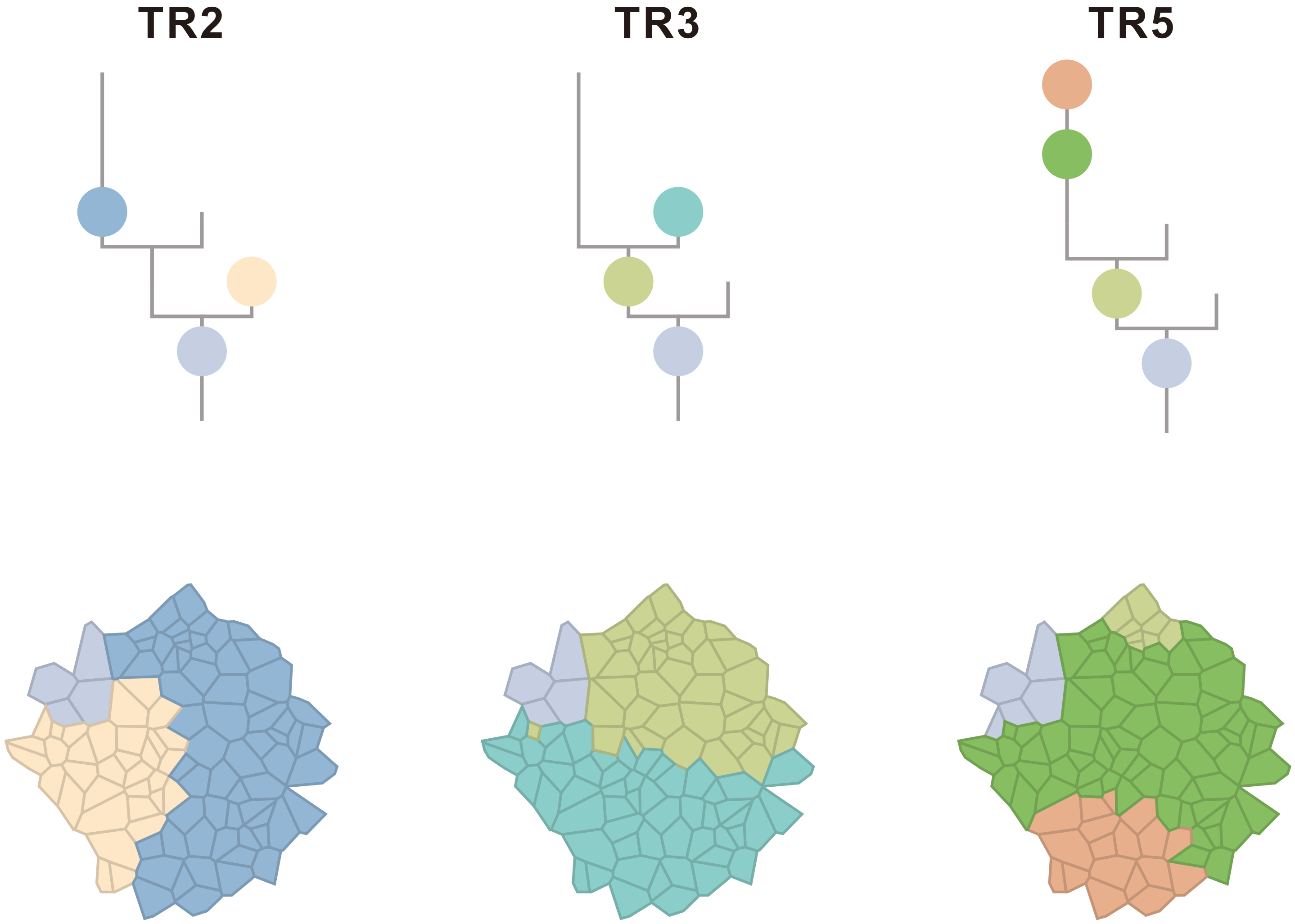

### CRC48

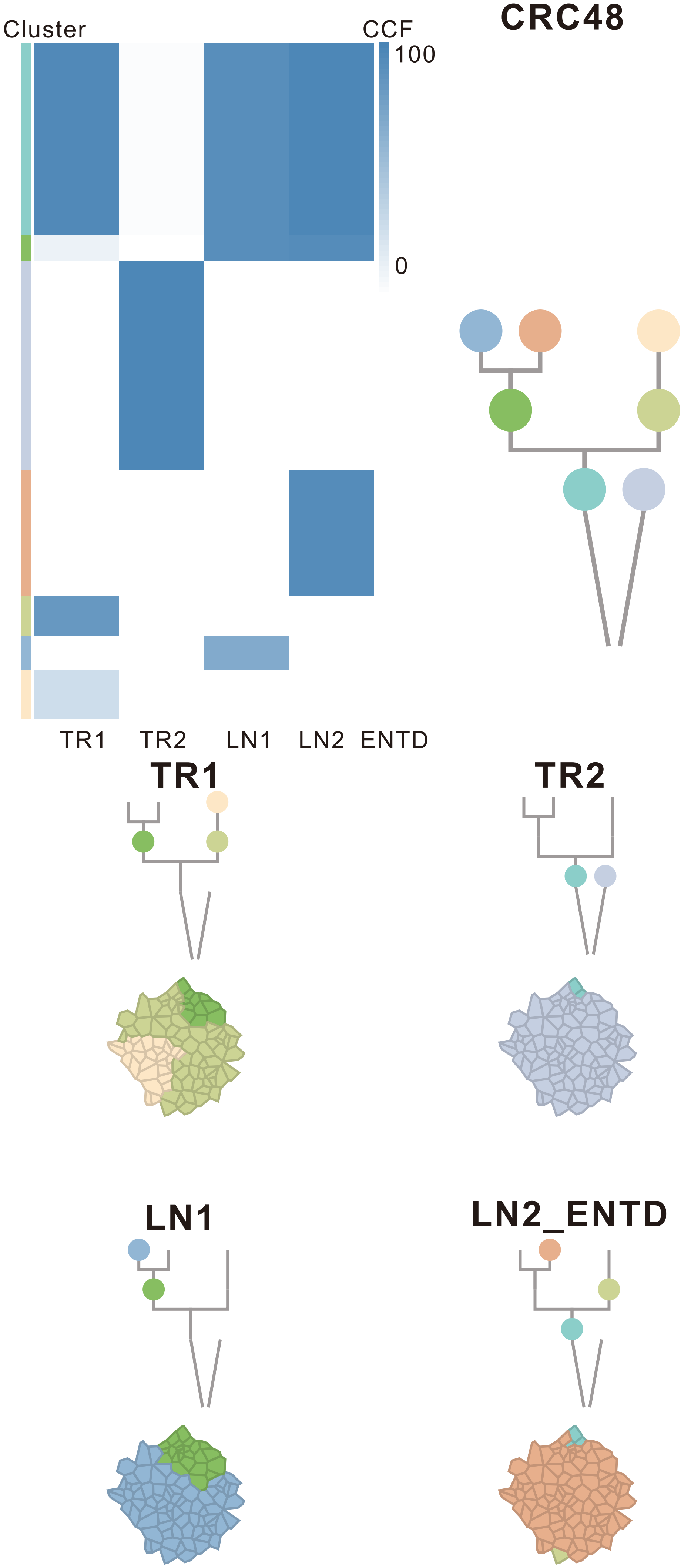

| Gene | Cytoband | Type | Cluster |
|---|---|---|---|
| * *APC* | 5q22.2 | InDel | 2 |
| * *APC* | 5q22.2 | InDel | 2 |
| *FLT3* | 13q12.2 | Amp | 2 |
| *FOXO1* | 13q14.11 | Amp | 2 |
| *CYSLTR2* | 13q14.2 | Amp | 2 |
| *PTPRD* | 9p24.1 | SNV | 6 |
| *RAD21* | 8q24.11 | Amp | 6 |
| *MYC* | 8q24.21 | Amp | 6 |
| *MUC4* | 3q29 | Amp | 7 |
| *PLAG1* | 8q12.1 | Amp | 7 |
| *PREX2* | 8q13.2 | Amp | 7 |
| *NCOA2* | 8q13.3 | Amp | 7 |
| *RUNX1T1* | 8q21.3 | Amp | 7 |
| *HEY1* | 8q21.13 | Amp | 7 |
| *CDH17* | 8q22.1 | Amp | 7 |
| *PABPC1* | 8q22.3 | Amp | 7 |
| *UBR5* | 8q22.3 | Amp | 7 |
| *CASP9* | 1p36.21 | Del | 7 |
| *PRDM2* | 1p36.21 | Del | 7 |
| *SPEN* | 1p36.21 | Del | 7 |
| *CAMTA1* | 1p36.31 | Del | 7 |
| *RPL22* | 1p36.31 | Del | 7 |
| *TNFRSF14* | 1p36.32 | Del | 7 |
| *SDHA* | 5p15.33 | Del | 7 |
| *FEN1* | 11q12.2 | Del | 7 |
| *SDHAF2* | 11q12.2 | Del | 7 |
| *MEN1* | 11q13.1 | Del | 7 |
| *EED* | 11q14.2 | Del | 7 |
| *BIRC3* | 11q22.2 | Del | 7 |
| *ATM* | 11q22.3 | Del | 7 |
| *DDX10* | 11q22.3 | Del | 7 |
| *SDHD* | 11q23.1 | Del | 7 |
| *ZBTB16* | 11q23.2 | Del | 7 |
| *ARHGEF12* | 11q23.3 | Del | 7 |
| *BCL9L* | 11q23.3 | Del | 7 |
| *CBL* | 11q23.3 | Del | 7 |
| *CCNB1IP1* | 14q11.2 | Del | 7 |
| *BAZ1A* | 14q13.2 | Del | 7 |
| *NKX2-1* | 14q13.3 | Del | 7 |
| *MAX* | 14q23.3 | Del | 7 |
| *RAD51B* | 14q24.1 | Del | 7 |
| *BCL11B* | 14q32.2 | Del | 7 |
| *DICER1* | 14q32.13 | Del | 7 |
| *AXIN1* | 16p13.3 | Del | 7 |
| *NTHL1* | 16p13.3 | Del | 7 |
| *TRAF7* | 16p13.3 | Del | 7 |
| *TSC2* | 16p13.3 | Del | 7 |
| *FLCN* | 17p11.2 | Del | 7 |
| *NCOR1* | 17p11.2 | Del | 7 |
| *MAP2K4* | 17p12 | Del | 7 |
| *PER1* | 17p13.1 | Del | 7 |
| *TP53* | 17p13.1 | Del | 7 |
| *YWHAE* | 17p13.3 | Del | 7 |
| *DNM2* | 19p13.2 | Del | 7 |
| *KEAP1* | 19p13.2 | Del | 7 |
| *SMARCA4* | 19p13.2 | Del | 7 |
| *STK11* | 19p13.3 | Del | 7 |
| *TCF3* | 19p13.3 | Del | 7 |
| *ELL* | 19p13.11 | Del | 7 |
| *SIRPA* | 20p13 | Del | 7 |
| *KRAS* | 12p12.1 | SNV | NA |
| *TP53* | 17p13.1 | SNV | NA |
| *SMAD4* | 18q21.2 | SNV | NA |
| *AMER1* | Xq11.2 | SNV | NA |
| *ATP2B3* | Xq28 | SNV | NA |
| *LZTR1* | 22q11.21 | InDel | NA |
| *BCORL1* | Xq26.1 | InDel | NA |

### CRC49

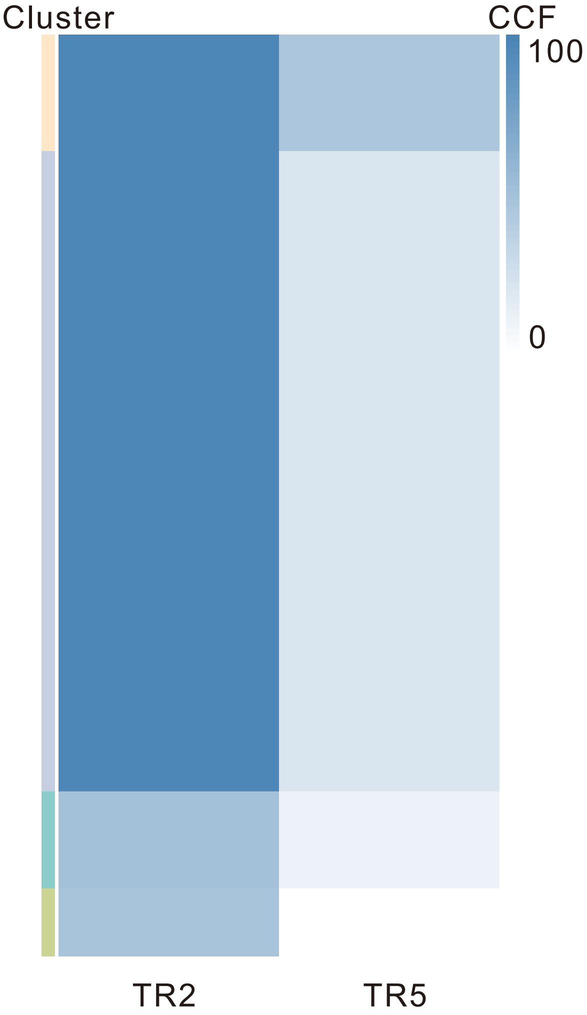

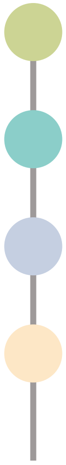

| Gene | Cytoband | Type | Cluster |
|---|---|---|---|
| * *APC* | 5q22.2 | SNV | 1 |
| * *APC* | 5q22.2 | SNV | 1 |
| *TP53* | 17p13.1 | SNV | 1 |
| *MUC4* | 3q29 | Amp | 3 |
| *RECQL4* | 8q24.3 | Amp | 3 |

## TR2

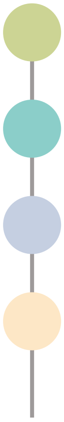

## TR5

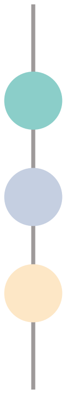

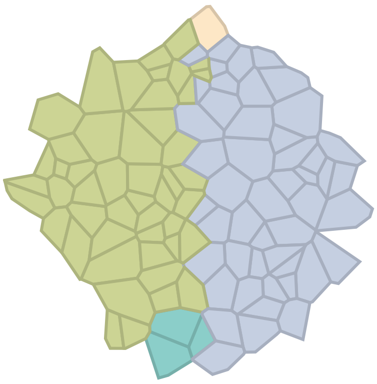

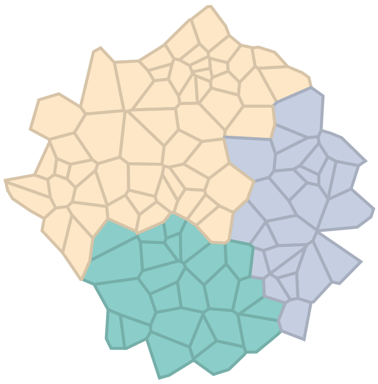

### CRC50

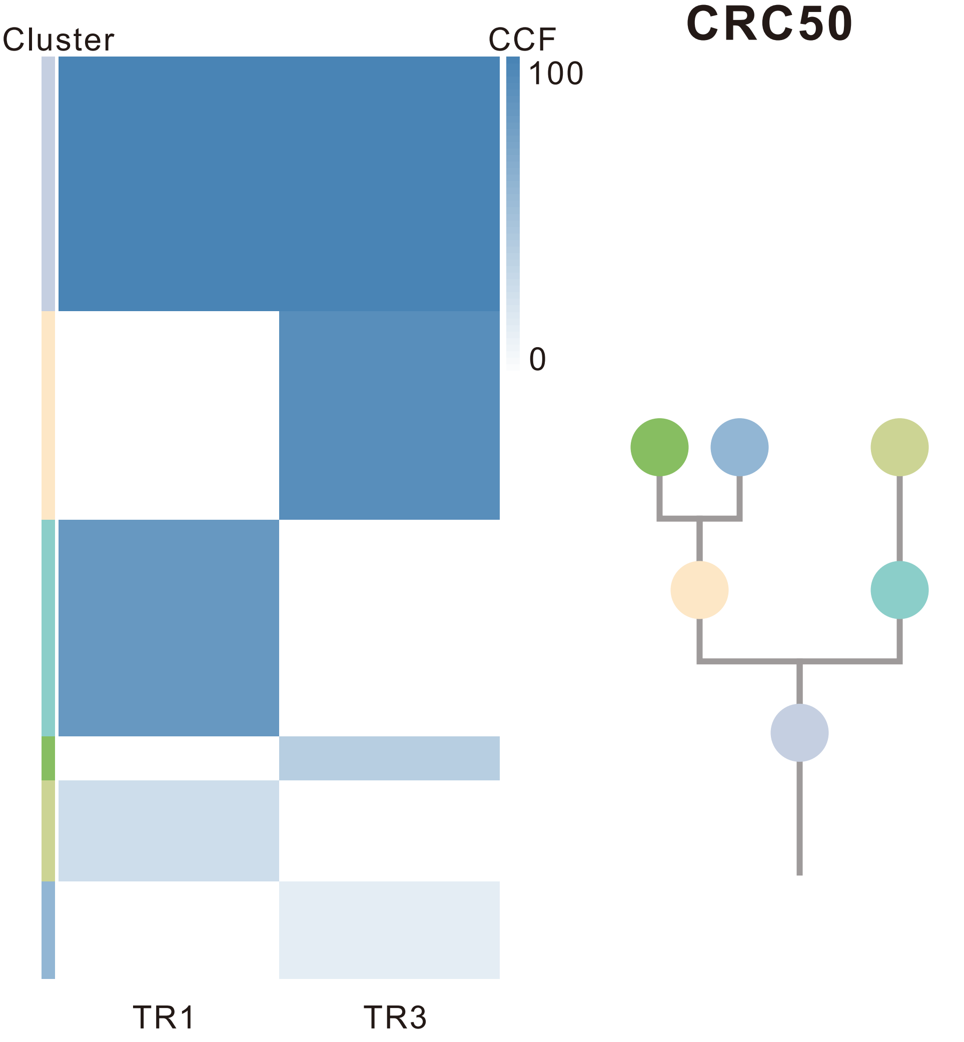

| Gene | Cytoband | Type | Cluster |
|---|---|---|---|
| *ARID1A* | 1p36.11 | SNV | 1 |
| *PIK3CA* | 3q26.32 | SNV | 1 |
| *KRAS* | 12p12.1 | SNV | 1 |
| *BAP1* | 3p21.1 | InDel | 1 |
| *APC* | 5q22.2 | InDel | 1 |
| *FBXW7* | 4q31.3 | SNV | 2 |
| *FAT4* | 4q28.1 | SNV | 3 |
| *DICER1* | 14q32.13 | InDel | 3 |
| *TSC1* | 9q34.13 | SNV | 5 |

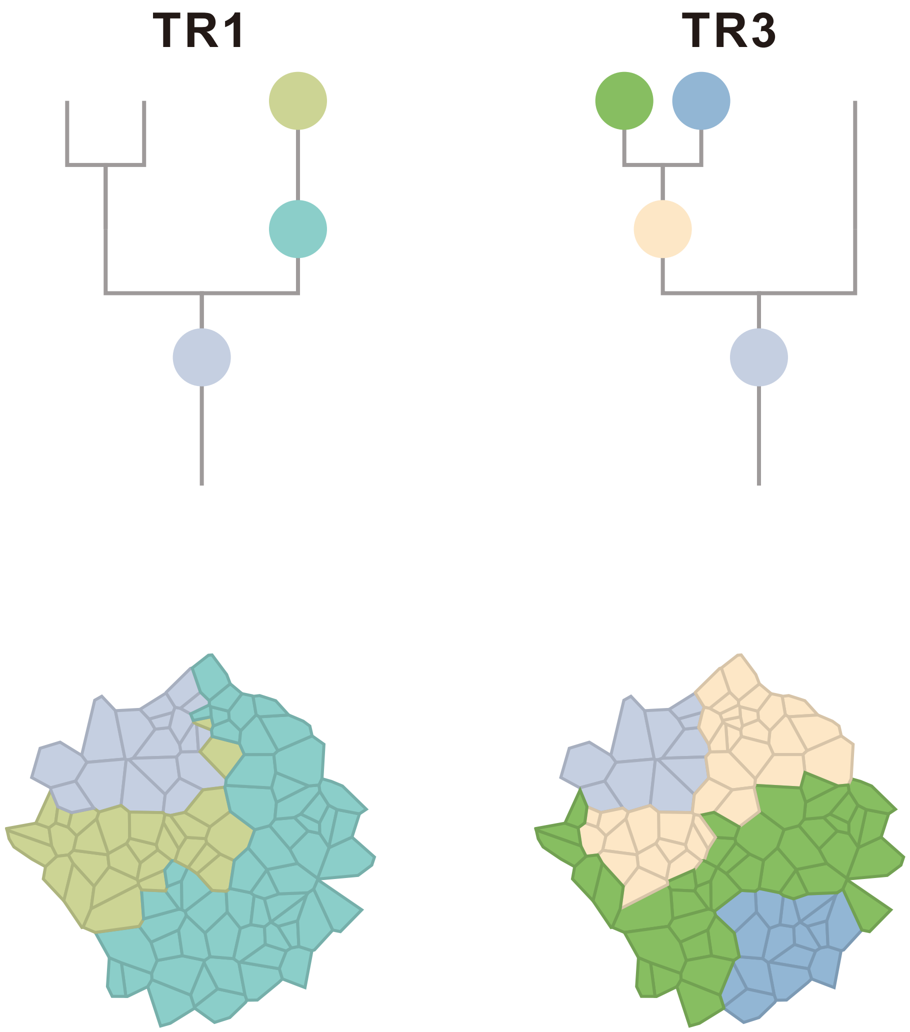

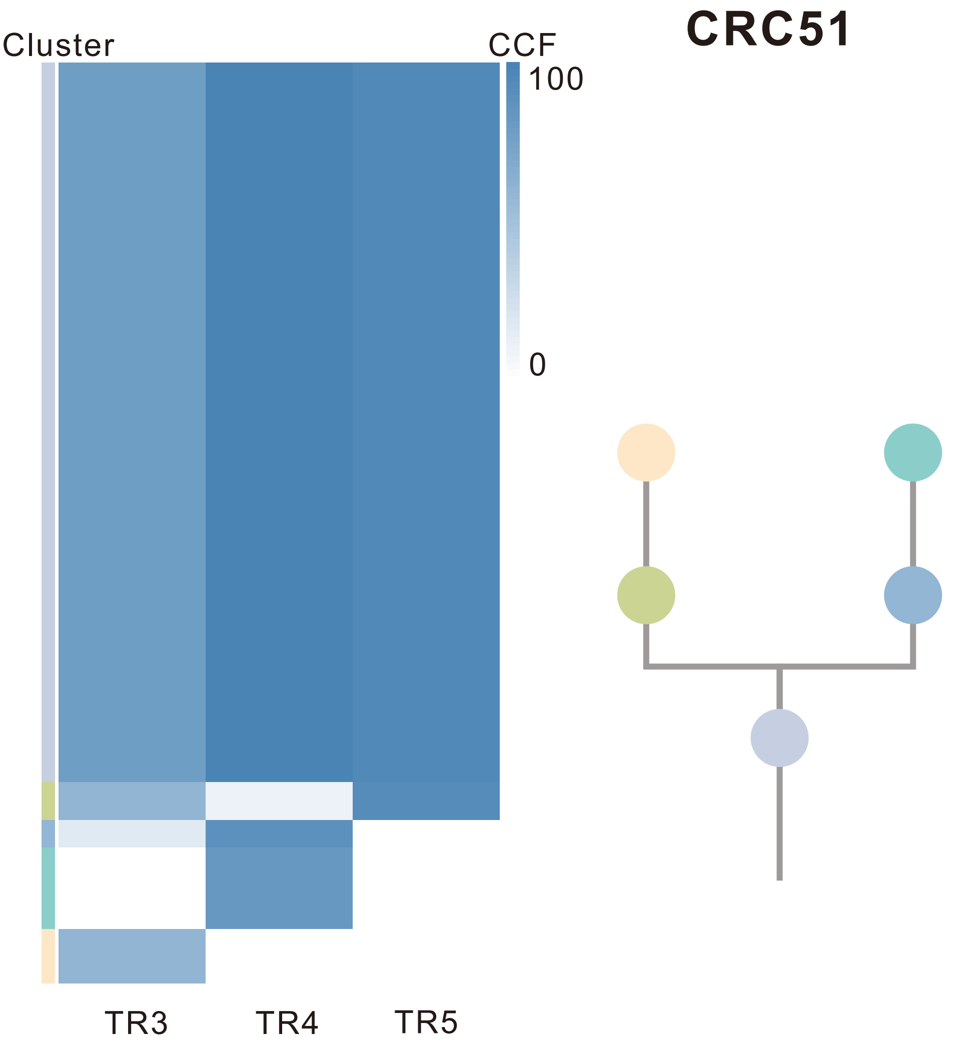

| Gene | Cytoband | Type | Cluster |
|---|---|---|---|
| *CAMTA1* | 1p36.23 | SNV | 1 |
| * *FBXW7* | 4q31.3 | SNV | 1 |
| * *APC* | 5q22.2 | SNV | 1 |
| *KRAS* | 12p12.1 | SNV | 1 |
| *TP53* | 17p13.1 | SNV | 1 |
| *SMAD4* | 18q21.2 | SNV | 1 |
| * *APC* | 5q22.2 | InDel | 1 |
| *AXIN2* | 17q24.1 | InDel | 1 |
| *LZTR1* | 22q11.21 | InDel | 1 |
| *MUC4* | 3q29 | Amp | 1 |
| *RUNX1T1* | 8q21.3 | Amp | 1 |
| *CDH17* | 8q22.1 | Amp | 1 |
| *PABPC1* | 8q22.3 | Amp | 1 |
| *UBR5* | 8q22.3 | Amp | 1 |
| *RAD21* | 8q24.11 | Amp | 1 |
| *MYC* | 8q24.21 | Amp | 1 |
| *RECQL4* | 8q24.3 | Amp | 1 |
| *ARNT* | 1q21.3 | Del | 3 |
| * *FBXW7* | 4q31.3 | InDel | 5 |
| *FOXP1* | 3p13 | Amp | NA |
| *MITF* | 3p13 | Amp | NA |
| *PLAG1* | 8q12.1 | Amp | NA |
| *PREX2* | 8q13.2 | Amp | NA |
| *NCOA2* | 8q13.3 | Amp | NA |
| *HEY1* | 8q21.13 | Amp | NA |

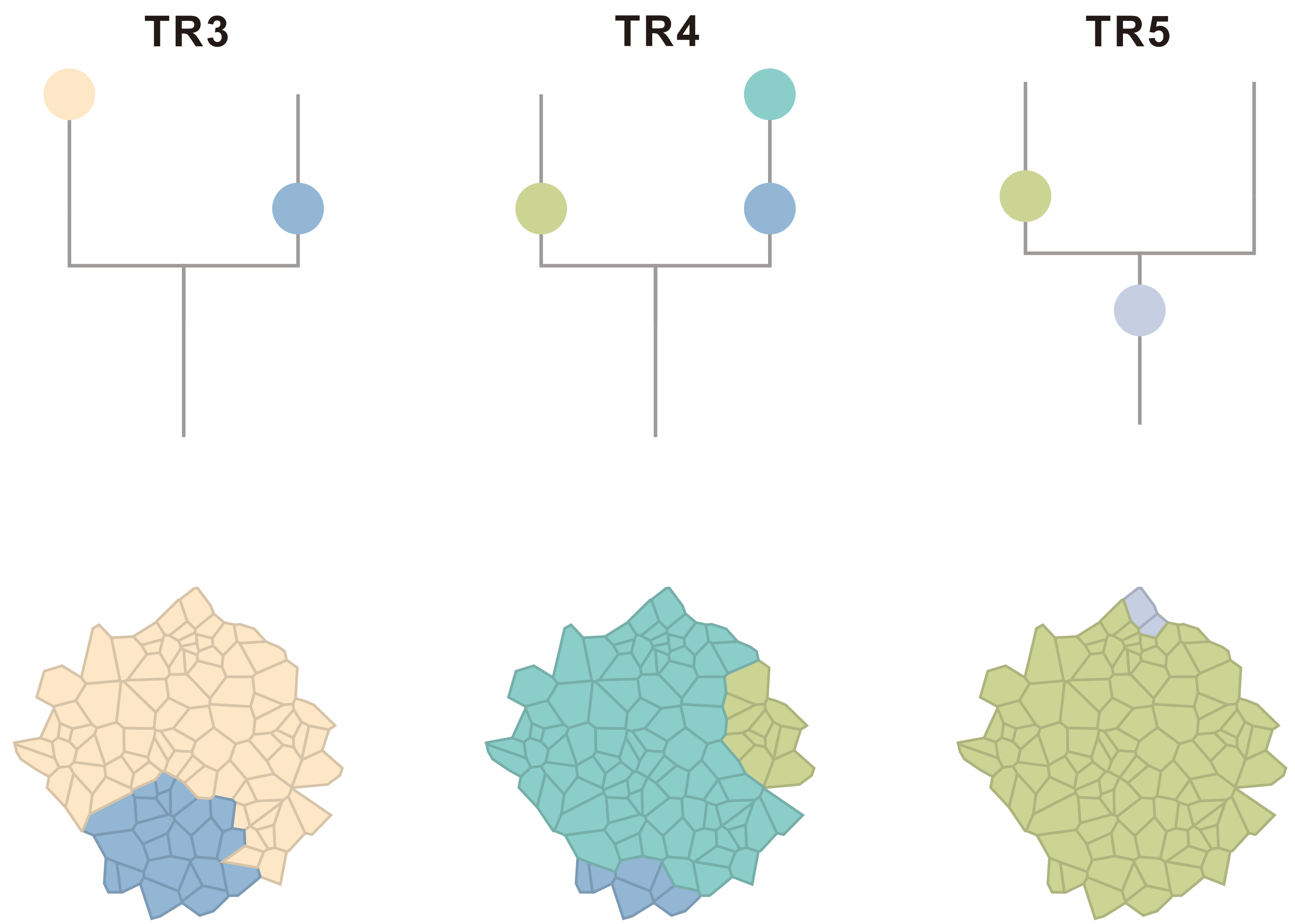

### CRC52

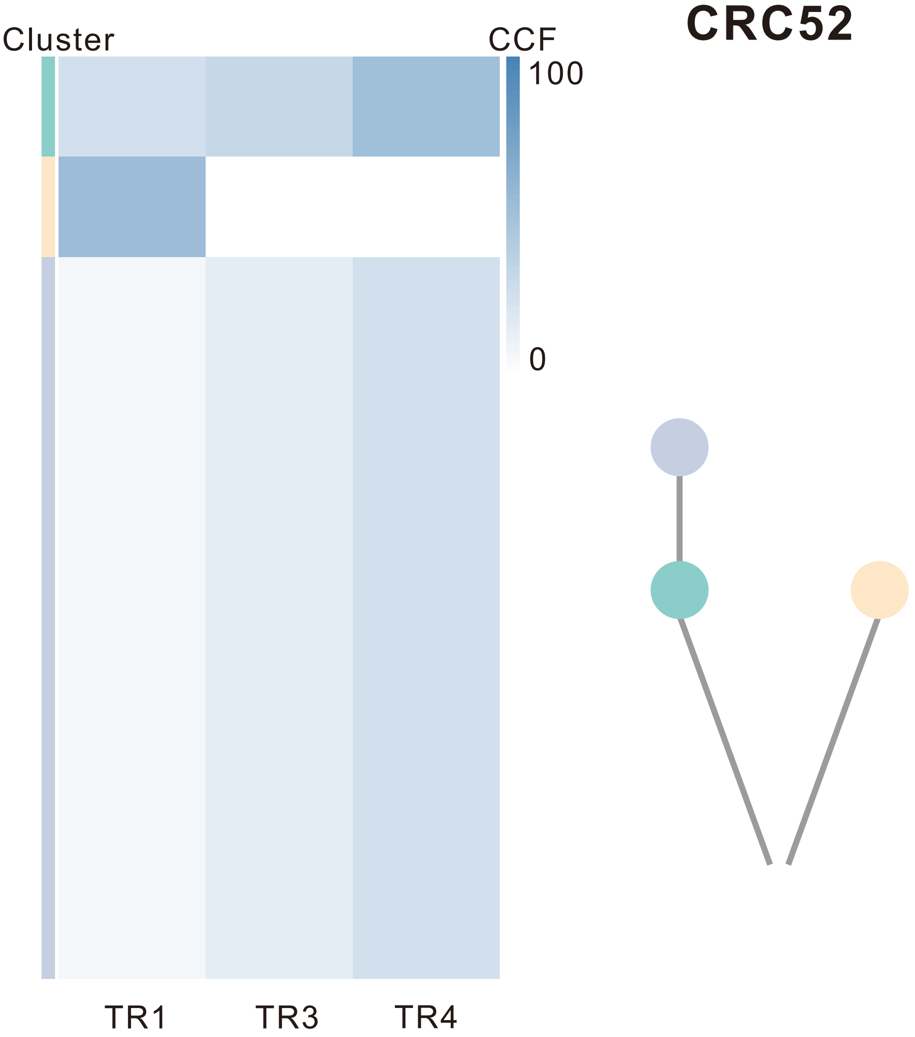

| Gene | Cytoband | Type | Cluster |
|---|---|---|---|
| *APC* | 5q22.2 | SNV | 1 |
| *TP53* | 17p13.1 | SNV | 1 |

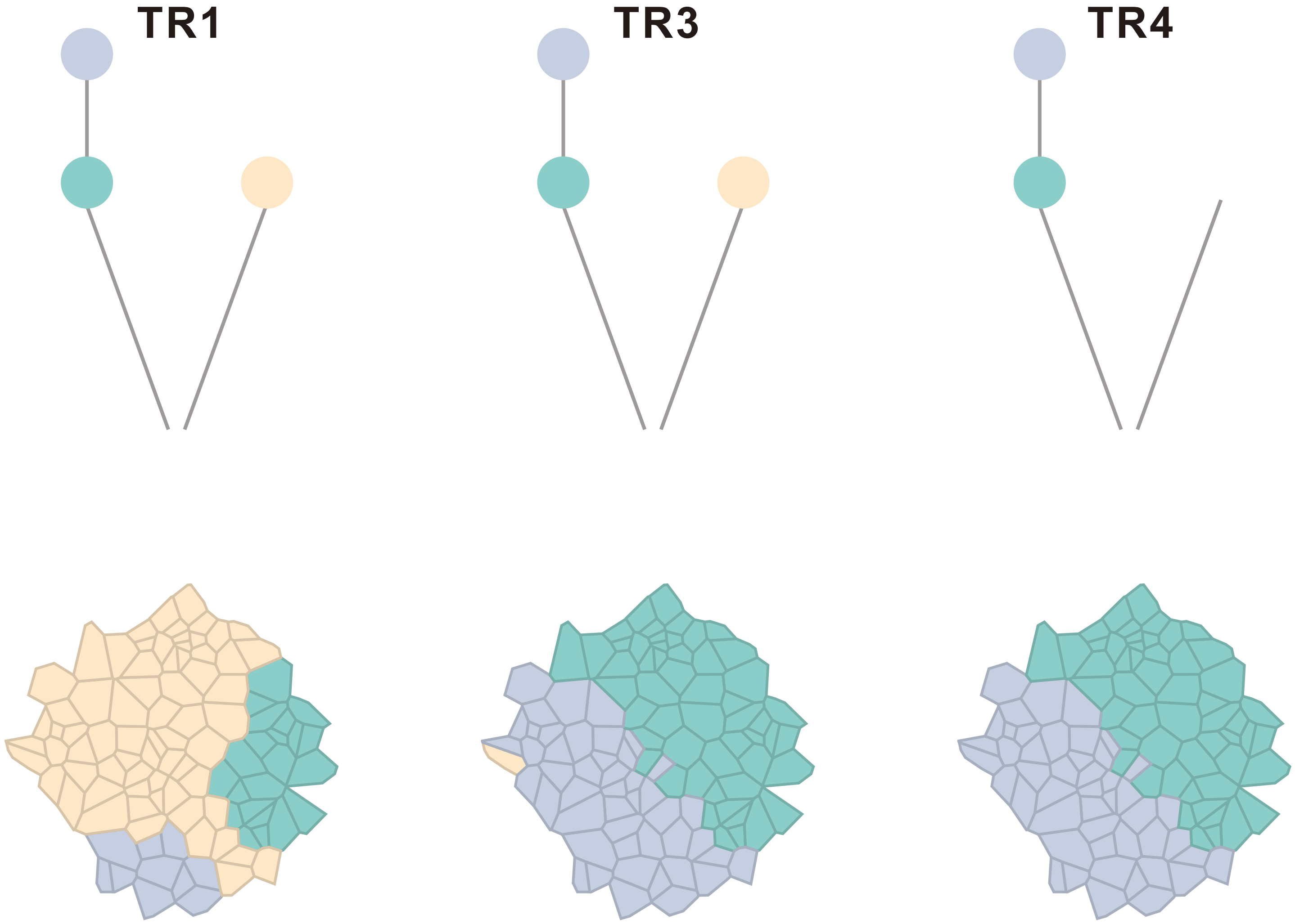

### CRC53

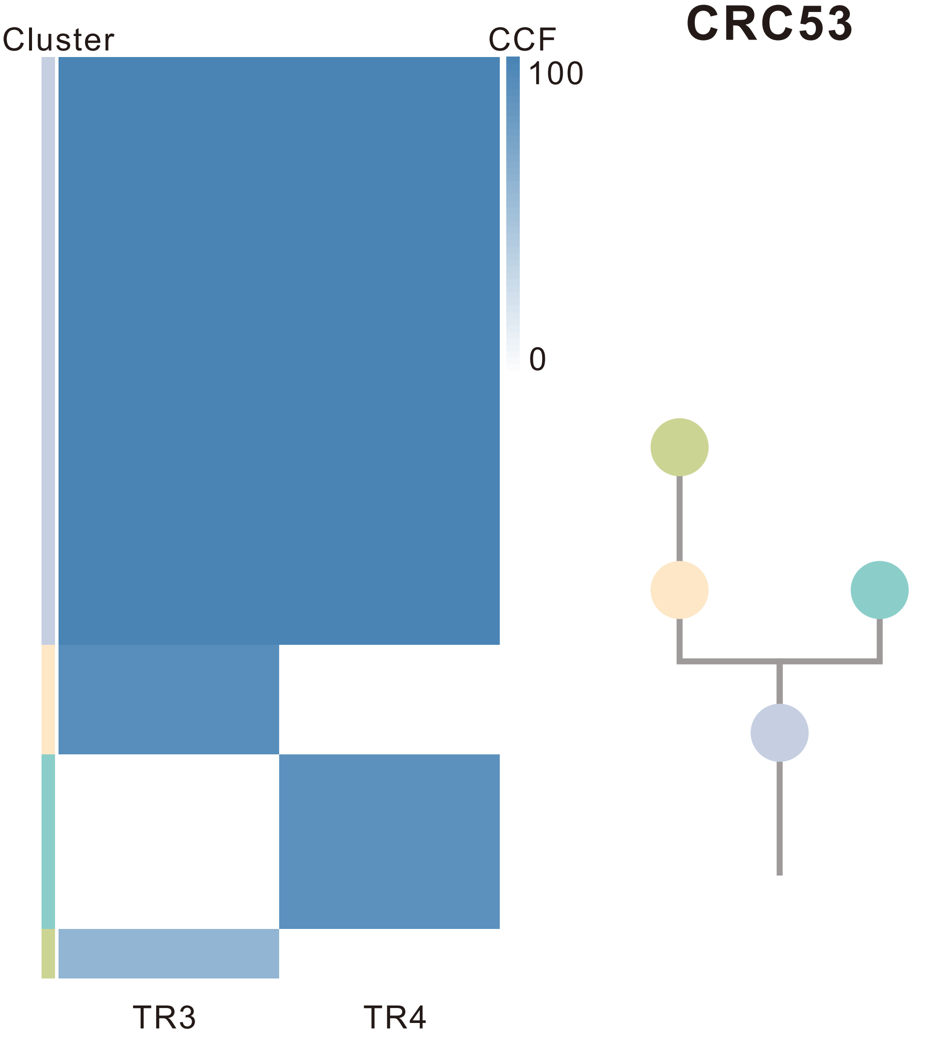

| Gene | Cytoband | Type | Cluster |
|---|---|---|---|
| *FBXW7* | 4q31.3 | SNV | 1 |
| *KRAS* | 12p12.1 | SNV | 1 |
| *TP53* | 17p13.1 | SNV | 1 |
| *SMARCA4* | 19p13.2 | SNV | 1 |
| *APC* | 5q22.2 | InDel | 1 |
| *LZTR1* | 22q11.21 | InDel | 1 |
| *KRAS* | 12p12.1 | Amp | 2 |
| *AMER1* | Xq11.2 | SNV | NA |

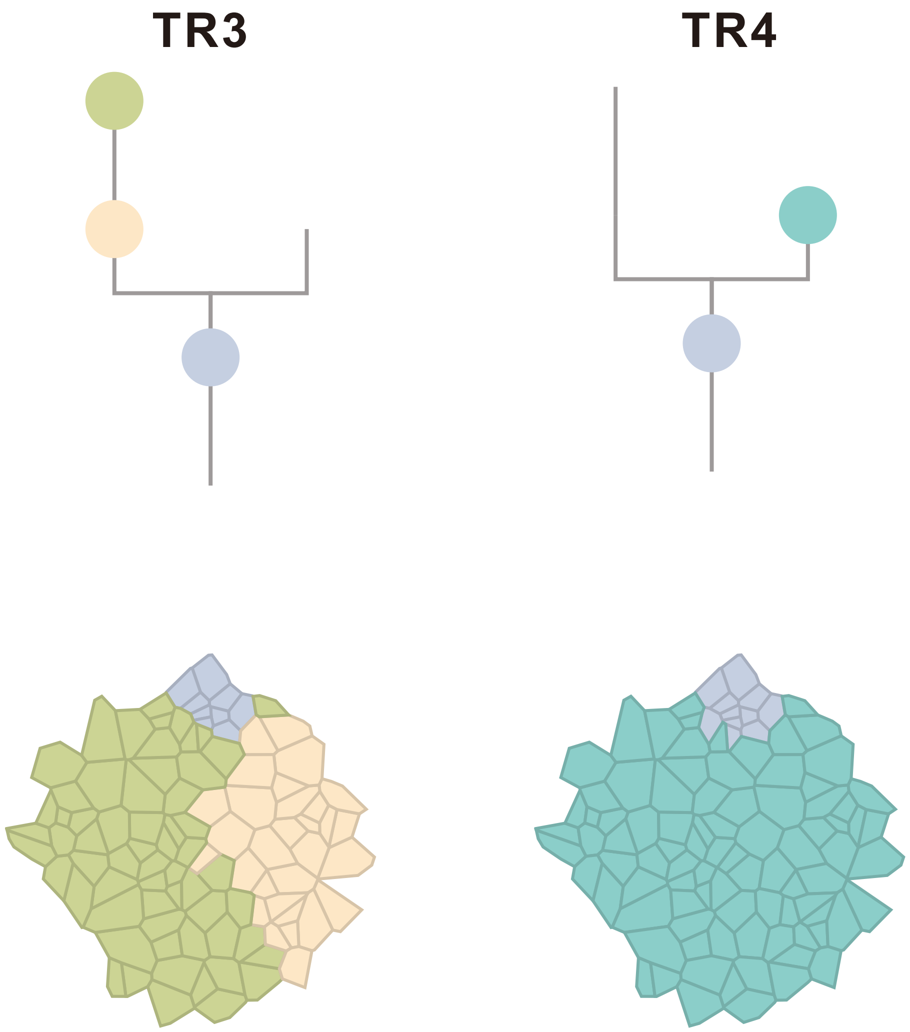

### CRC54

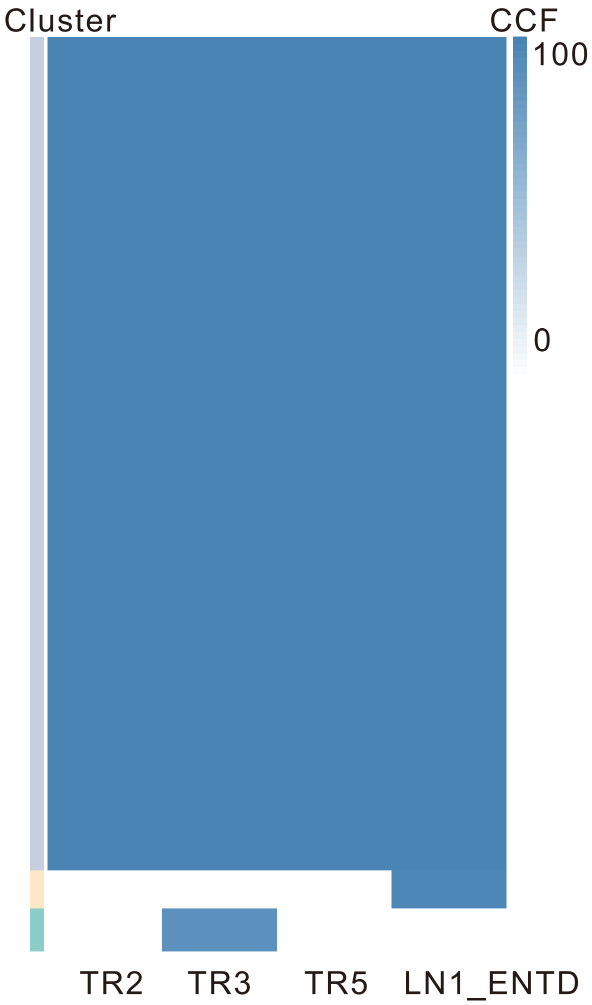

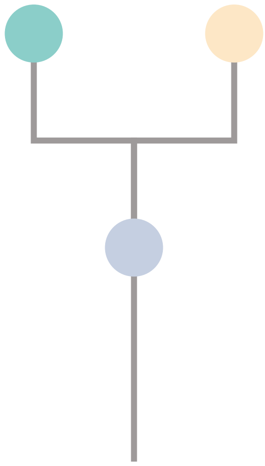

| Gene | Cytoband | Type | Cluster |
|---|---|---|---|
| * *APC* | 5q22.2 | SNV | 1 |
| *KRAS* | 12p12.1 | SNV | 1 |
| *MAX* | 14q23.3 | SNV | 1 |
| *TP53* | 17p13.1 | SNV | 1 |
| *SMAD4* | 18q21.2 | SNV | 1 |
| *LZTR1* | 22q11.21 | InDel | 1 |
| *SSX2* | Xp11.22 | Amp | 1 |
| *ARAF* | Xp11.23 | Amp | 1 |
| *GATA1* | Xp11.23 | Amp | 1 |
| *SSX1* | Xp11.23 | Amp | 1 |
| *SSX4* | Xp11.23 | Amp | 1 |
| *TFE3* | Xp11.23 | Amp | 1 |
| *WAS* | Xp11.23 | Amp | 1 |
| *KDM6A* | Xp11.3 | Amp | 1 |
| * *APC* | 5q22.2 | SNV | NA |
| *AMER1* | Xq11.2 | SNV | NA |
| *HRAS* | 11p15.5 | Amp | NA |

## TR2

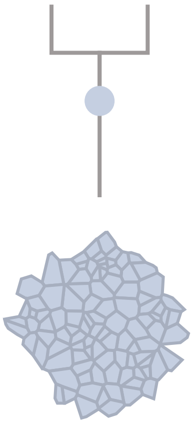

## TR3

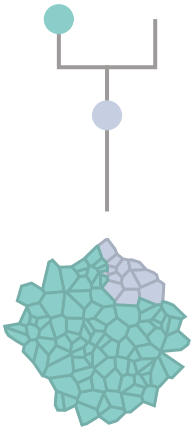

## TR5

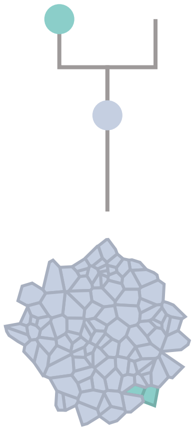

#### LN1_ENTD

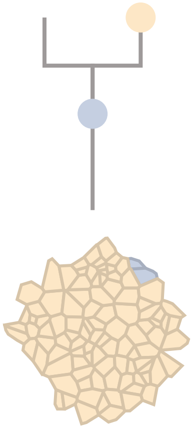

### CRC55

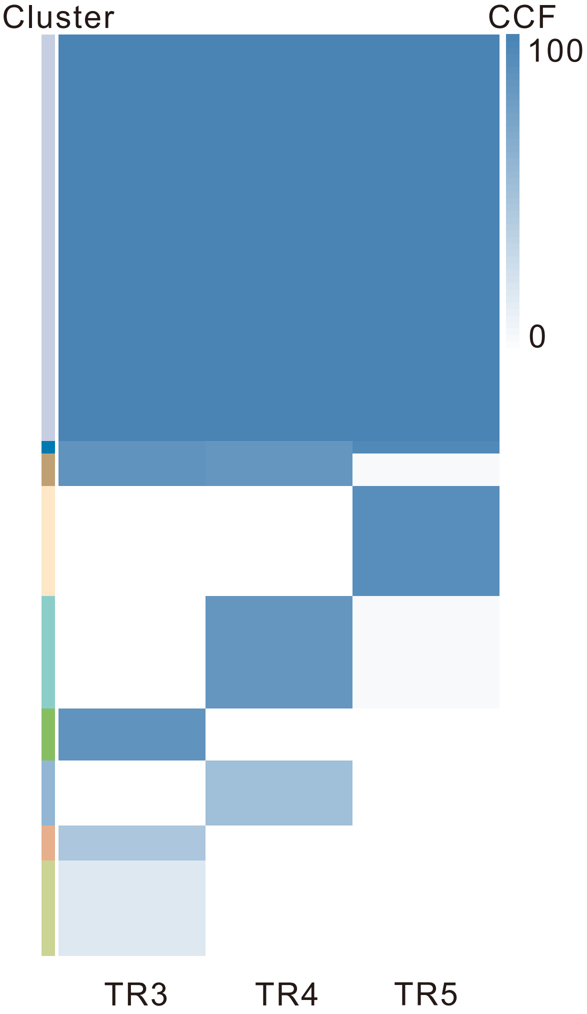

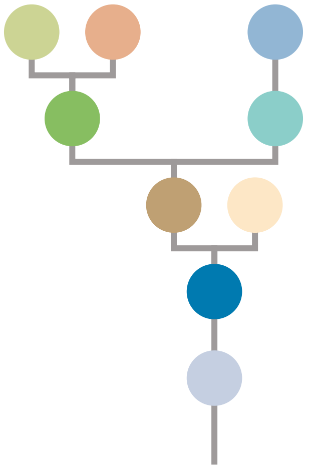

| Gene | Cytoband | Type | Cluster |
|---|---|---|---|
| *APC* | 5q22.2 | SNV | 1 |
| *CDH1* | 16q22.1 | SNV | 1 |
| *TP53* | 17p13.1 | SNV | 1 |
| *ARID1A* | 1p36.11 | InDel | 1 |
| *EGFR* | 7p11.2 | Amp | 1 |
| *HNRNPA2B1* | 7p15.2 | Amp | 1 |
| *HOXA11* | 7p15.2 | Amp | 1 |
| *HOXA13* | 7p15.2 | Amp | 1 |
| *HOXA9* | 7p15.2 | Amp | 1 |
| *MACC1* | 7p21.1 | Amp | 1 |
| *ETV1* | 7p21.2 | Amp | 1 |
| *RAC1* | 7p22.1 | Amp | 1 |
| *CARD11* | 7p22.2 | Amp | 1 |
| *FLT3* | 13q12.2 | Amp | 1 |
| *FOXO1* | 13q14.11 | Amp | 1 |
| *CYSLTR2* | 13q14.2 | Amp | 1 |
| *SRC* | 20q11.23 | Amp | 1 |
| *MUC4* | 3q29 | Amp | 2 |
| *NCOR1* | 17p12 | SNV | 3 |
| *MAFB* | 20q12 | Amp | 3 |
| *PLCG1* | 20q12 | Amp | 3 |
| *NFATC2* | 20q13.2 | Amp | 3 |
| *SALL4* | 20q13.2 | Amp | 3 |
| *GNAS* | 20q13.32 | Amp | 3 |
| *PTK6* | 20q13.33 | Amp | 3 |
| *AR* | Xq12 | SNV | NA |

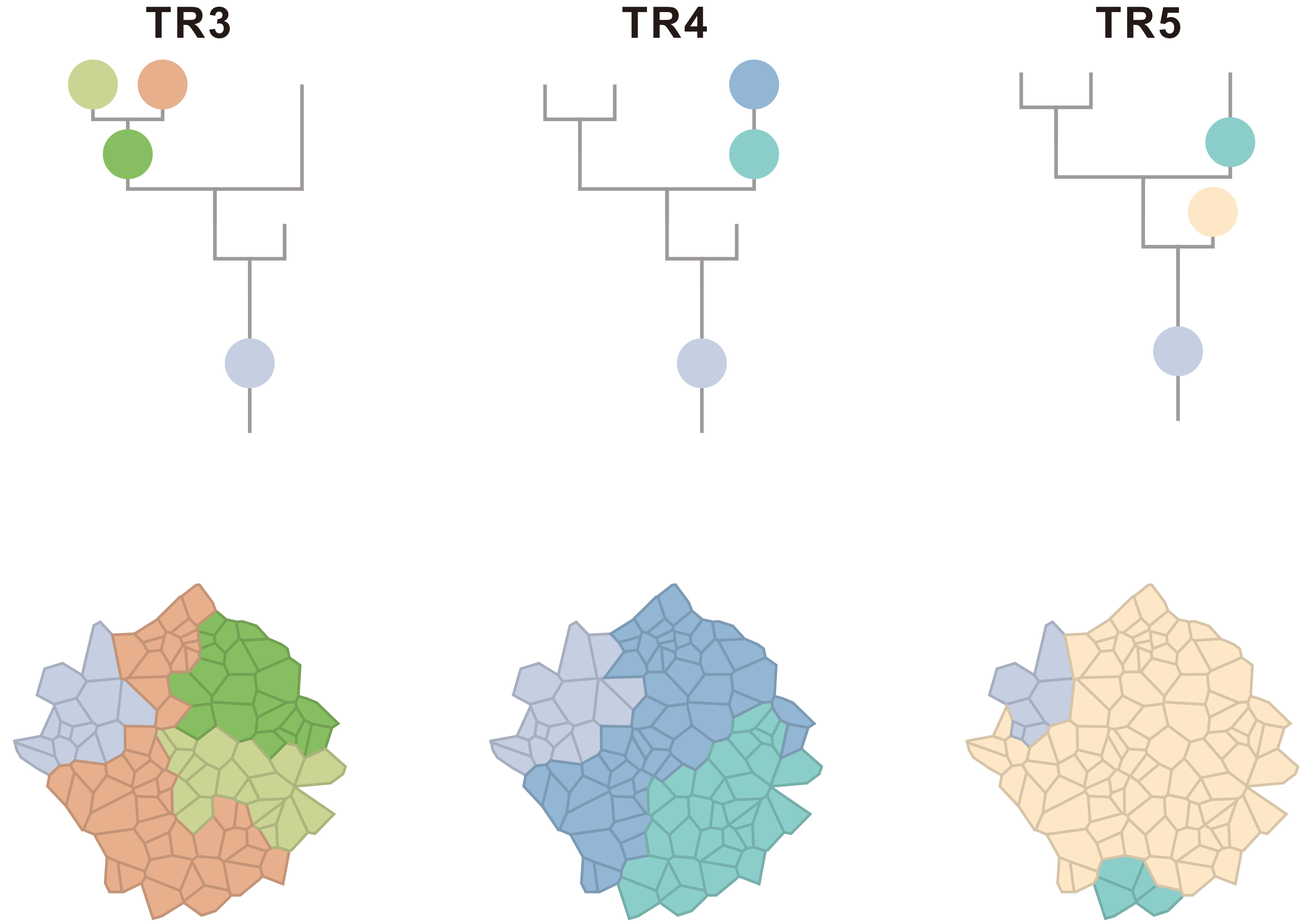

#### CRC56

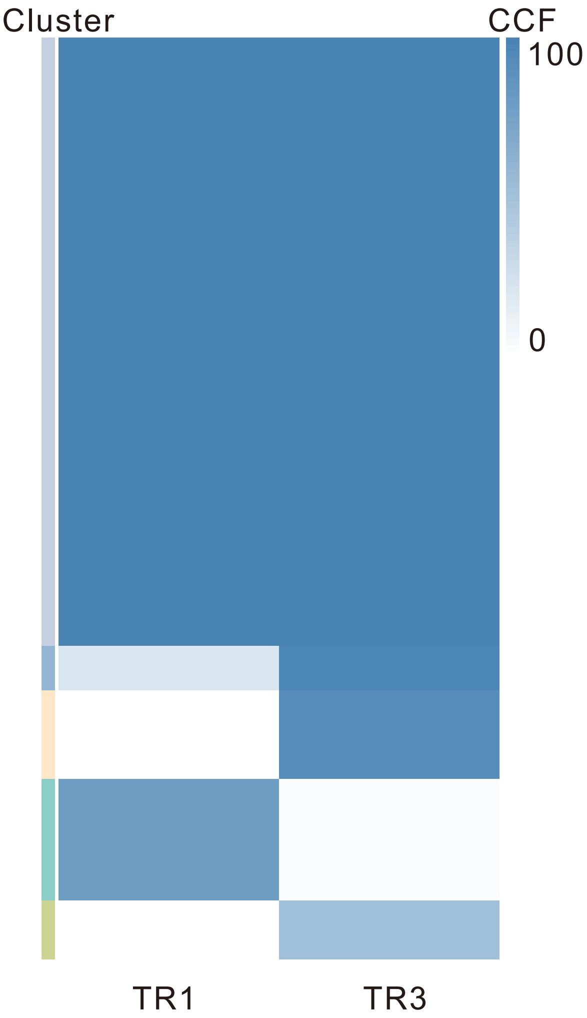

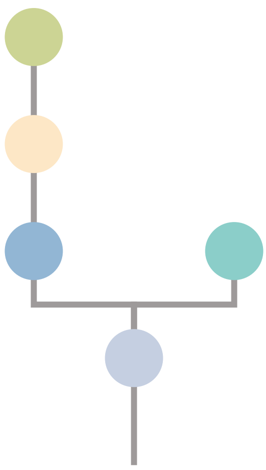

| Gene | Cytoband | Type | Cluster |
|---|---|---|---|
| *PIK3CA* | 3q26.32 | SNV | 1 |
| *KDR* | 4q12 | SNV | 1 |
| * *FBXW7* | 4q31.3 | SNV | 1 |
| * *APC* | 5q22.2 | SNV | 1 |
| *TP53* | 17p13.1 | SNV | 1 |
| *BCL9L* | 11q23.3 | InDel | 1 |
| *SRC* | 20q11.23 | Amp | 1 |
| *MAFB* | 20q12 | Amp | 1 |
| *PLCG1* | 20q12 | Amp | 1 |
| *NFATC2* | 20q13.2 | Amp | 1 |
| *SALL4* | 20q13.2 | Amp | 1 |
| *GNAS* | 20q13.32 | Amp | 1 |
| *PTK6* | 20q13.33 | Amp | 1 |
| *ARHGEF10* | 8p23.3 | Del | 1 |
| * *FBXW7* | 4q31.3 | SNV | 5 |
| * *APC* | 5q22.2 | SNV | 5 |
| *RBM10* | Xp11.23 | SNV | NA |

### TR1

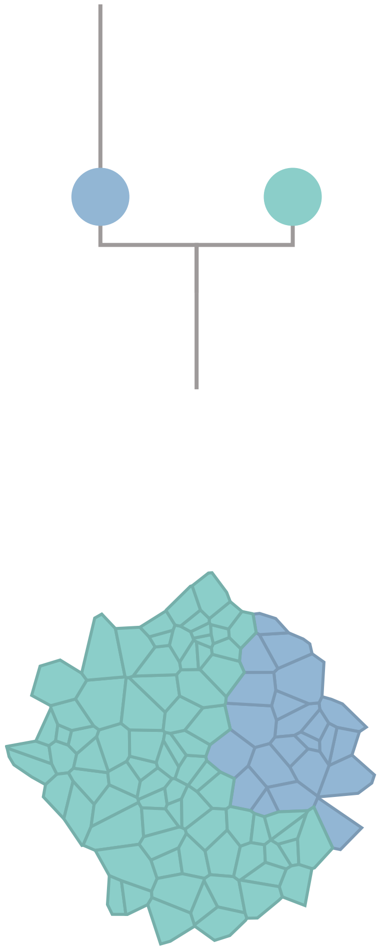

### TR3

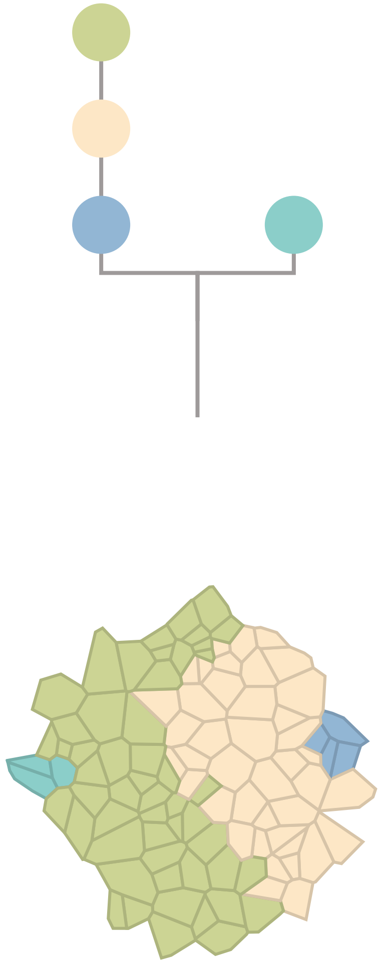

### CRC57

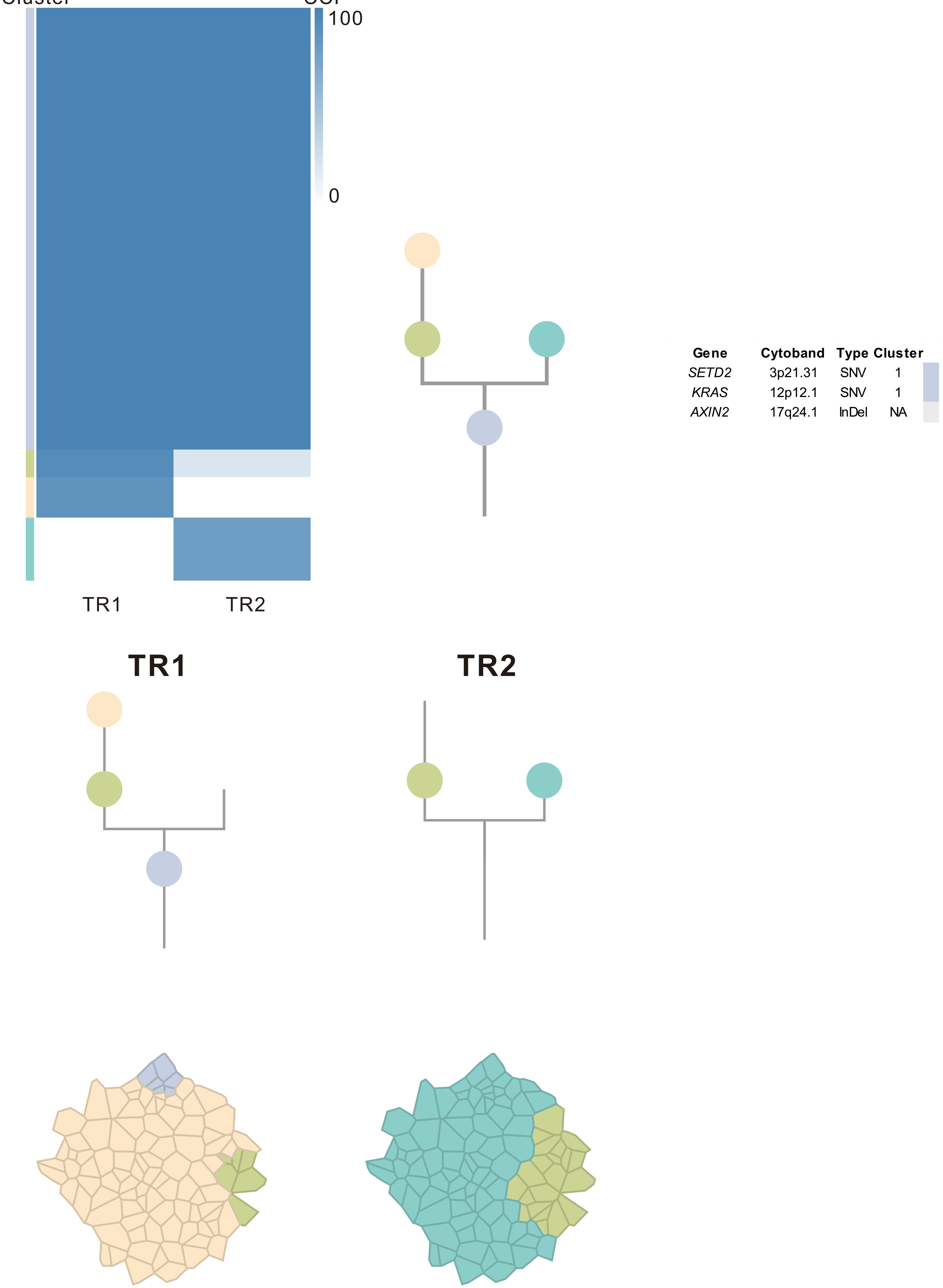

| Gene | Cytoband | Type | Cluster |
|---|---|---|---|
| *SETD2* | 3p21.31 | SNV | 1 |
| *KRAS* | 12p12.1 | SNV | 1 |
| *AXIN2* | 17q24.1 | InDel | NA |

### CRC58

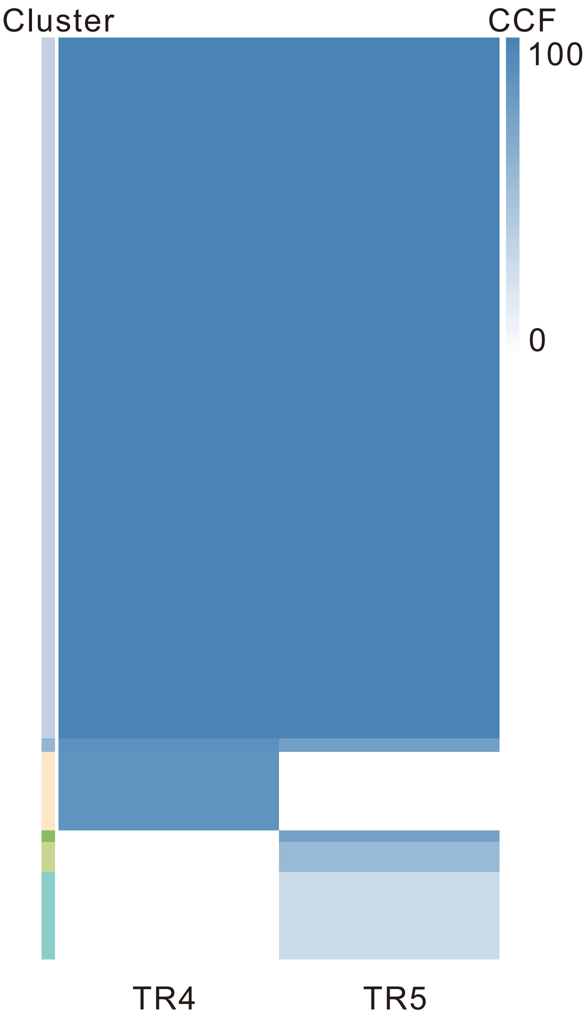

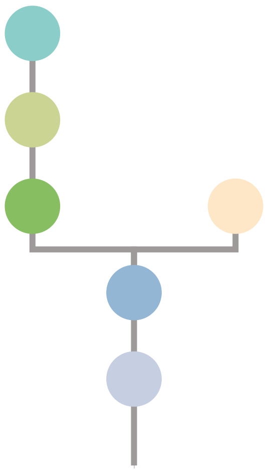

| Gene | Cytoband | Type | Cluster |
|---|---|---|---|
| *CTNNB1* | 3p22.1 | SNV | 1 |
| *ATR* | 3q23 | SNV | 1 |
| *TP53* | 17p13.1 | SNV | 1 |
| *EGFR* | 7p11.2 | Amp | 1 |
| *PLAG1* | 8q12.1 | Amp | 1 |
| *PREX2* | 8q13.2 | Amp | 1 |
| *NCOA2* | 8q13.3 | Amp | 1 |
| *RUNX1T1* | 8q21.3 | Amp | 1 |
| *HEY1* | 8q21.13 | Amp | 1 |
| *CDH17* | 8q22.1 | Amp | 1 |
| *PABPC1* | 8q22.3 | Amp | 1 |
| *UBR5* | 8q22.3 | Amp | 1 |
| *RECQL4* | 8q24.3 | Amp | 1 |
| *RAD21* | 8q24.11 | Amp | 1 |
| *MYC* | 8q24.21 | Amp | 1 |
| *FLT3* | 13q12.2 | Amp | 1 |
| *SRC* | 20q11.23 | Amp | 1 |
| *MAFB* | 20q12 | Amp | 1 |
| *PLCG1* | 20q12 | Amp | 1 |
| *NFATC2* | 20q13.2 | Amp | 1 |
| *SALL4* | 20q13.2 | Amp | 1 |
| *LZTR1* | 22q11.21 | InDel | 3 |
| *GNAS* | 20q13.32 | Amp | NA |

## TR4

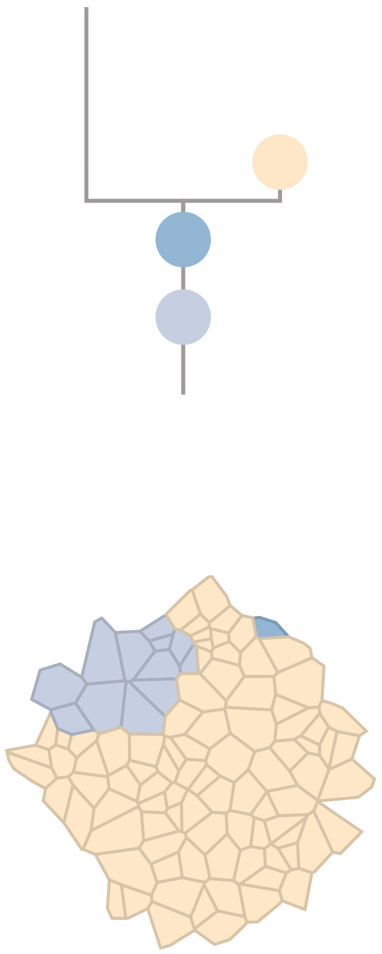

## TR5

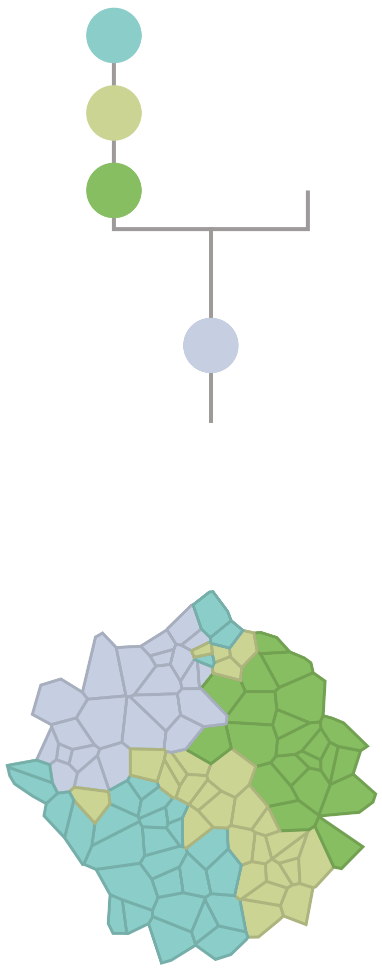

### CRC59

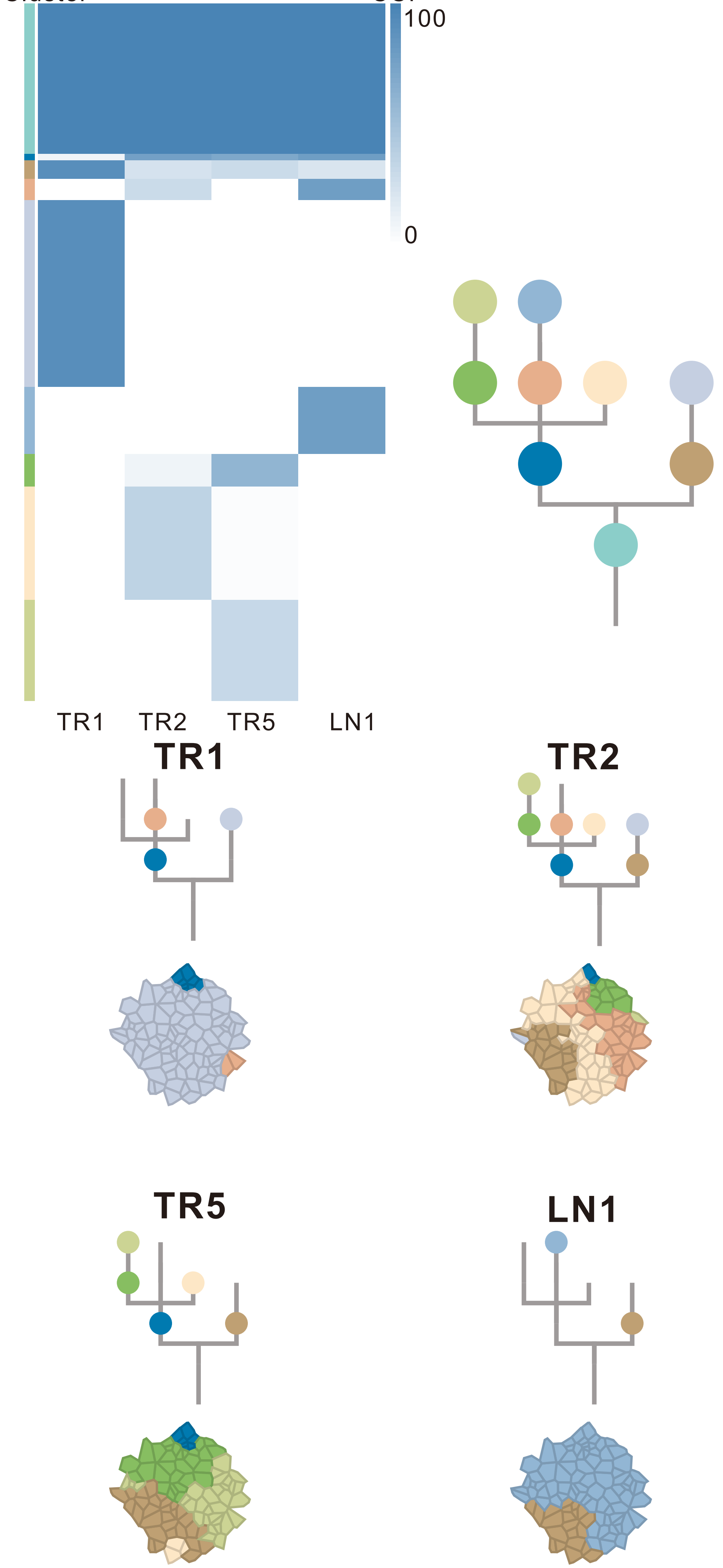

| Gene | Cytoband | Type | Cluster |
|---|---|---|---|
| *LRP1B* | 2q22.1 | SNV | 2 |
| * *APC* | 5q22.2 | SNV | 2 |
| *TCF7L2* | 10q25.3 | SNV | 2 |
| *TP53* | 17p13.1 | SNV | 2 |
| * *APC* | 5q22.2 | InDel | 2 |
| *FLT3* | 13q12.2 | Amp | 2 |
| *SRC* | 20q11.23 | Amp | 2 |
| *MAFB* | 20q12 | Amp | 2 |
| *PLCG1* | 20q12 | Amp | 2 |
| *NFATC2* | 20q13.2 | Amp | 2 |
| *SALL4* | 20q13.2 | Amp | 2 |
| *GNAS* | 20q13.32 | Amp | 2 |
| *PTK6* | 20q13.33 | Amp | 2 |
| *LEPROTL1* | 8p12 | Del | 2 |
| *WRN* | 8p12 | Del | 2 |
| *ARHGEF10* | 8p23.3 | Del | 2 |
| *SMAD2* | 18q21.1 | Del | 2 |
| *SMAD4* | 18q21.2 | Del | 2 |
| *RUNX1* | 21q22.12 | Del | 2 |
| *PIK3CA* | 3q26.32 | SNV | 3 |
| *CHD2* | 15q26.1 | SNV | 3 |
| *CREBBP* | 16p13.3 | SNV | 4 |
| *PBRM1* | 3p21.1 | InDel | 4 |
| *IKBKB* | 8p11.21 | Amp | 8 |
| *KAT6A* | 8p11.21 | Amp | 8 |
| *PLAG1* | 8q12.1 | Amp | 8 |
| *RUNX1T1* | 8q21.3 | Amp | 8 |
| *CDH17* | 8q22.1 | Amp | 8 |
| *PABPC1* | 8q22.3 | Amp | 8 |
| *UBR5* | 8q22.3 | Amp | 8 |
| *RAD21* | 8q24.11 | Amp | 8 |
| *MYC* | 8q24.21 | Amp | 8 |
| *NRG1* | 8p12 | Del | 8 |
| *HEY1* | 8q21.13 | Amp | NA |
| *OLIG2* | 21q22.11 | Amp | NA |

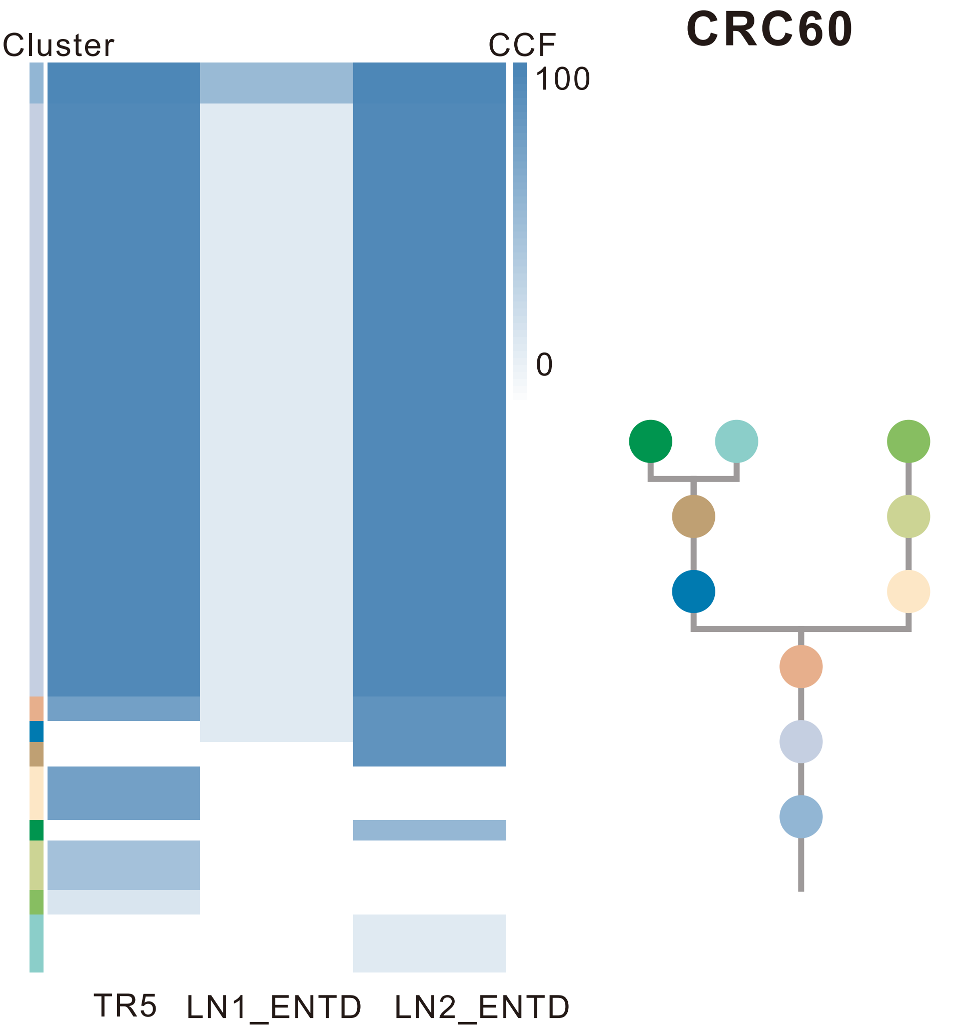

| Gene | Cytoband | Type | Cluster |
|---|---|---|---|
| *TGFBR2* | 3p24.1 | SNV | 1 |
| *KRAS* | 12p12.1 | SNV | 1 |
| *TP53* | 17p13.1 | SNV | 1 |
| * *APC* | 5q22.2 | InDel | 1 |
| * *APC* | 5q22.2 | InDel | 1 |
| *FLT3* | 13q12.2 | Amp | 5 |
| *CYSLTR2* | 13q14.2 | Amp | 5 |
| *FOXO1* | 13q14.11 | Amp | 5 |

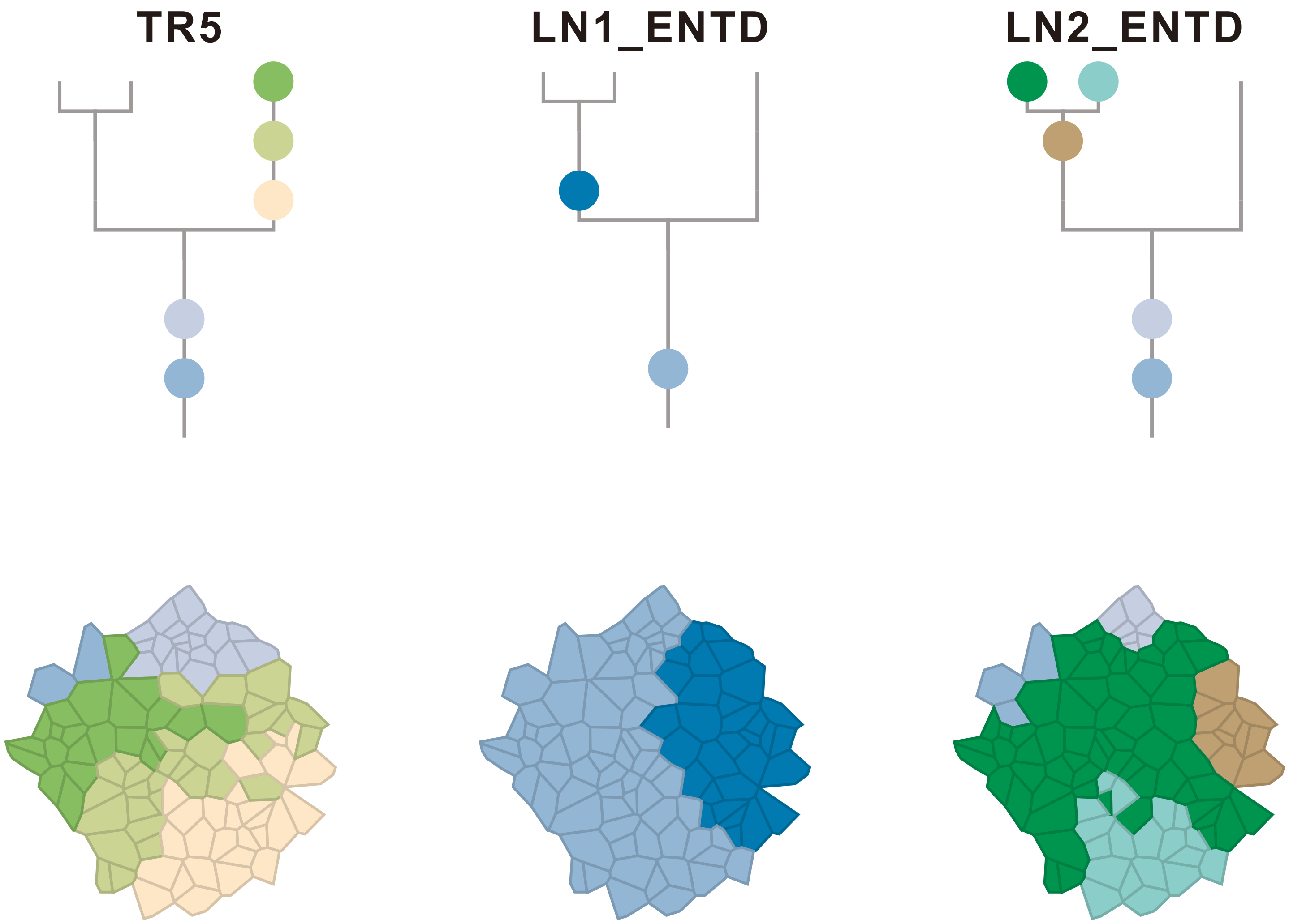

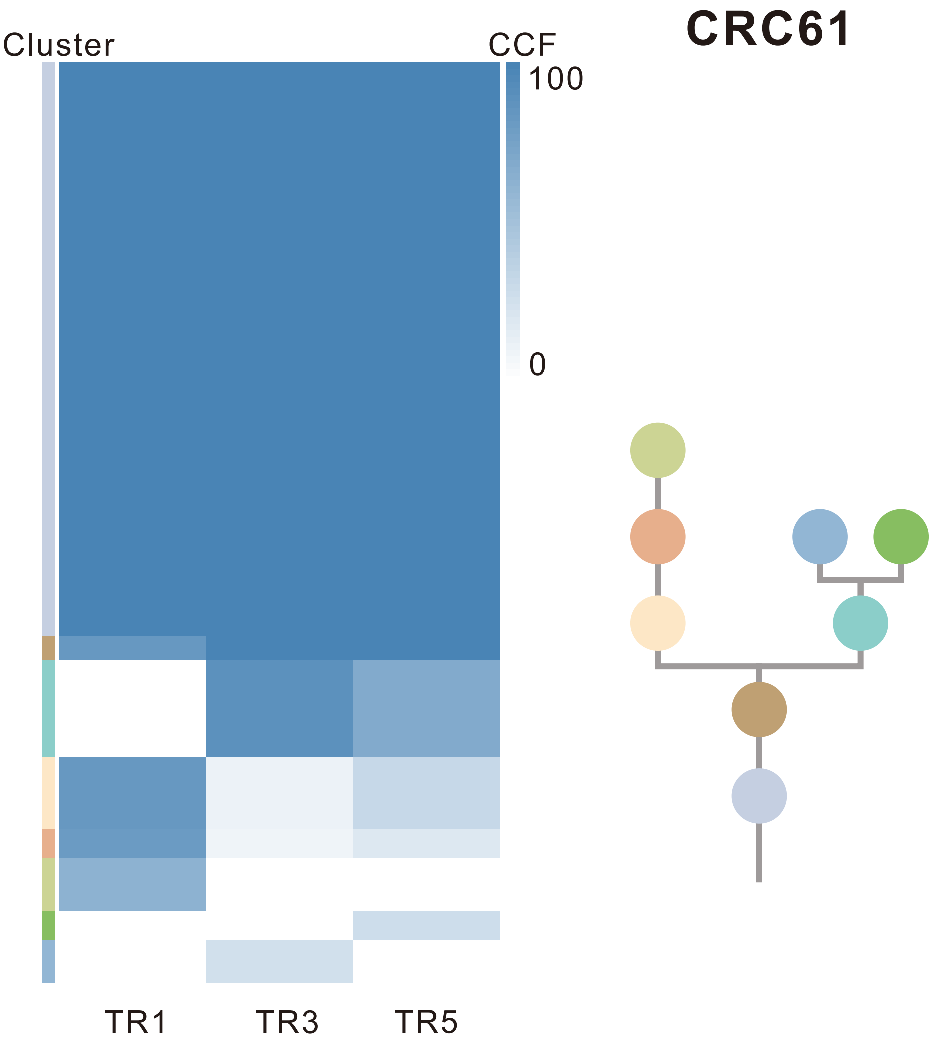

| Gene | Cytoband | Type | Cluster |
|---|---|---|---|
| *ARID1A* | 1p36.11 | SNV | 1 |
| *FBXW7* | 4q31.3 | SNV | 1 |
| *APC* | 5q22.2 | SNV | 1 |
| *TCF7L2* | 10q25.3 | SNV | 1 |
| *KRAS* | 12p12.1 | SNV | 1 |
| *TP53* | 17p13.1 | SNV | 1 |
| *AMER1* | Xq11.2 | SNV | NA |

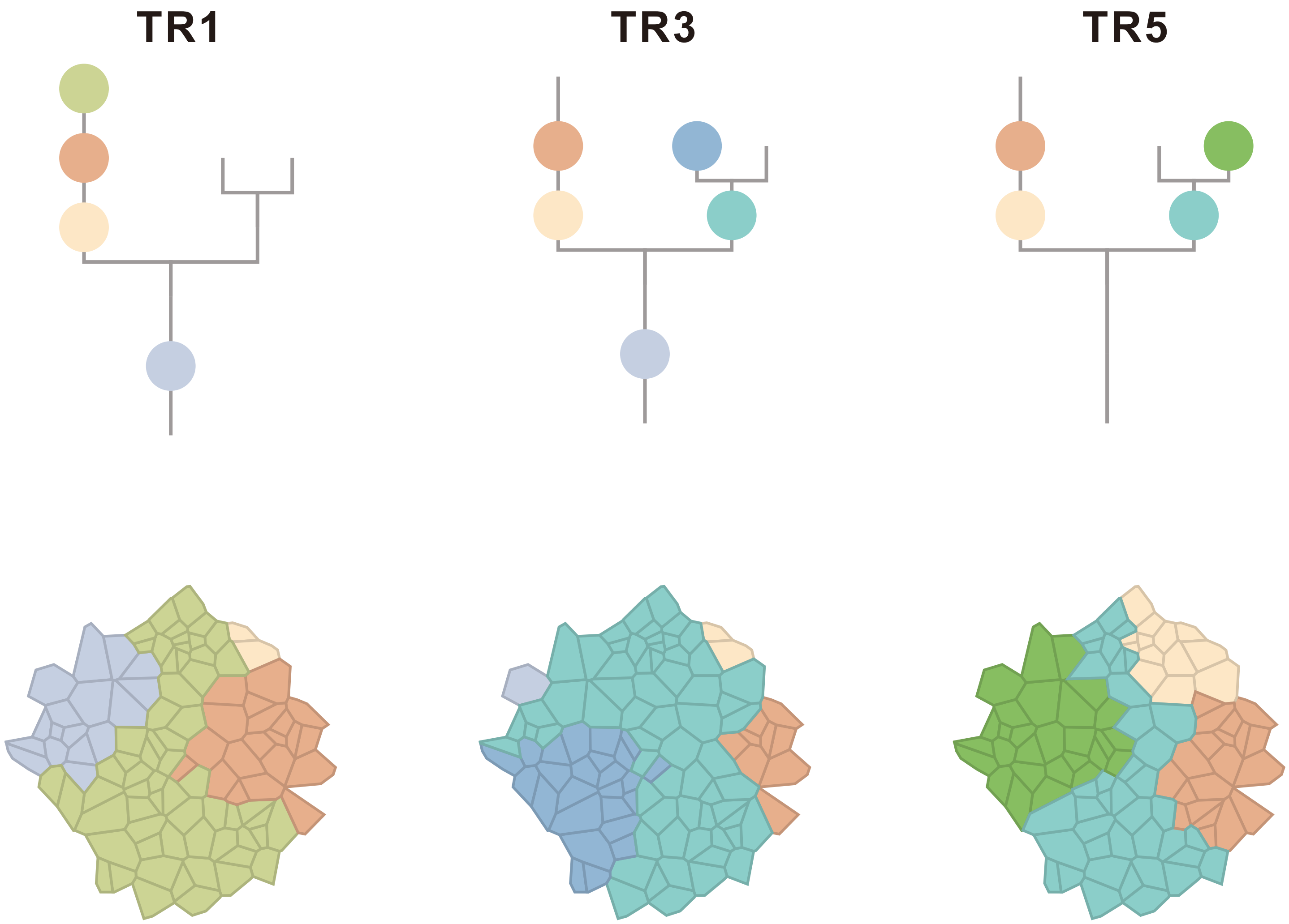

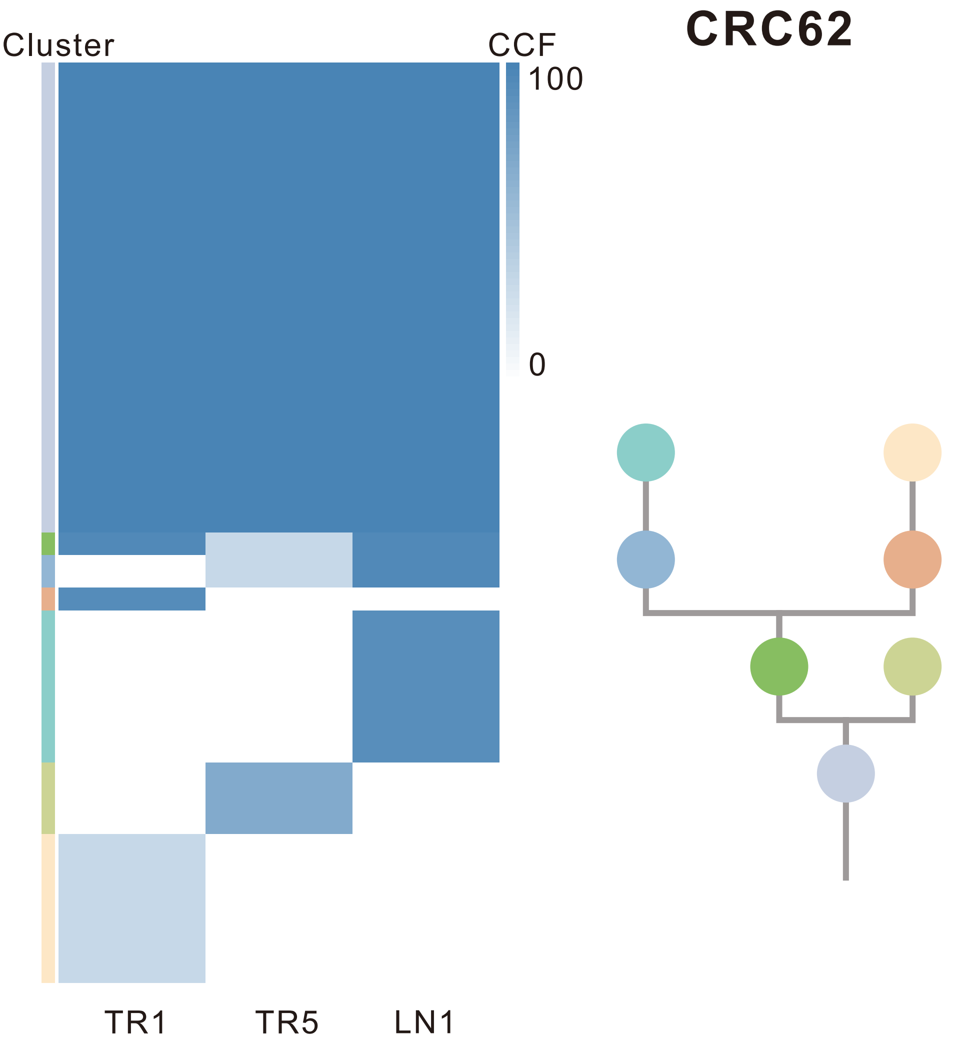

| Gene | Cytoband | Type | Cluster |
|---|---|---|---|
| *TP53* | 17p13.1 | SNV | 1 |
| *ARID1A* | 1p36.11 | InDel | 1 |
| *IL7R* | 5p13.2 | Amp | 1 |
| *CTNND2* | 5p15.2 | Amp | 1 |
| *TERT* | 5p15.33 | Amp | 1 |
| *SRC* | 20q11.23 | Amp | 1 |
| *MAFB* | 20q12 | Amp | 1 |
| *PLCG1* | 20q12 | Amp | 1 |
| *NFATC2* | 20q13.2 | Amp | 1 |
| *SALL4* | 20q13.2 | Amp | 1 |
| *GNAS* | 20q13.32 | Amp | 1 |
| *PTK6* | 20q13.33 | Amp | 1 |
| *GPC5* | 13q31.3 | SNV | NA |

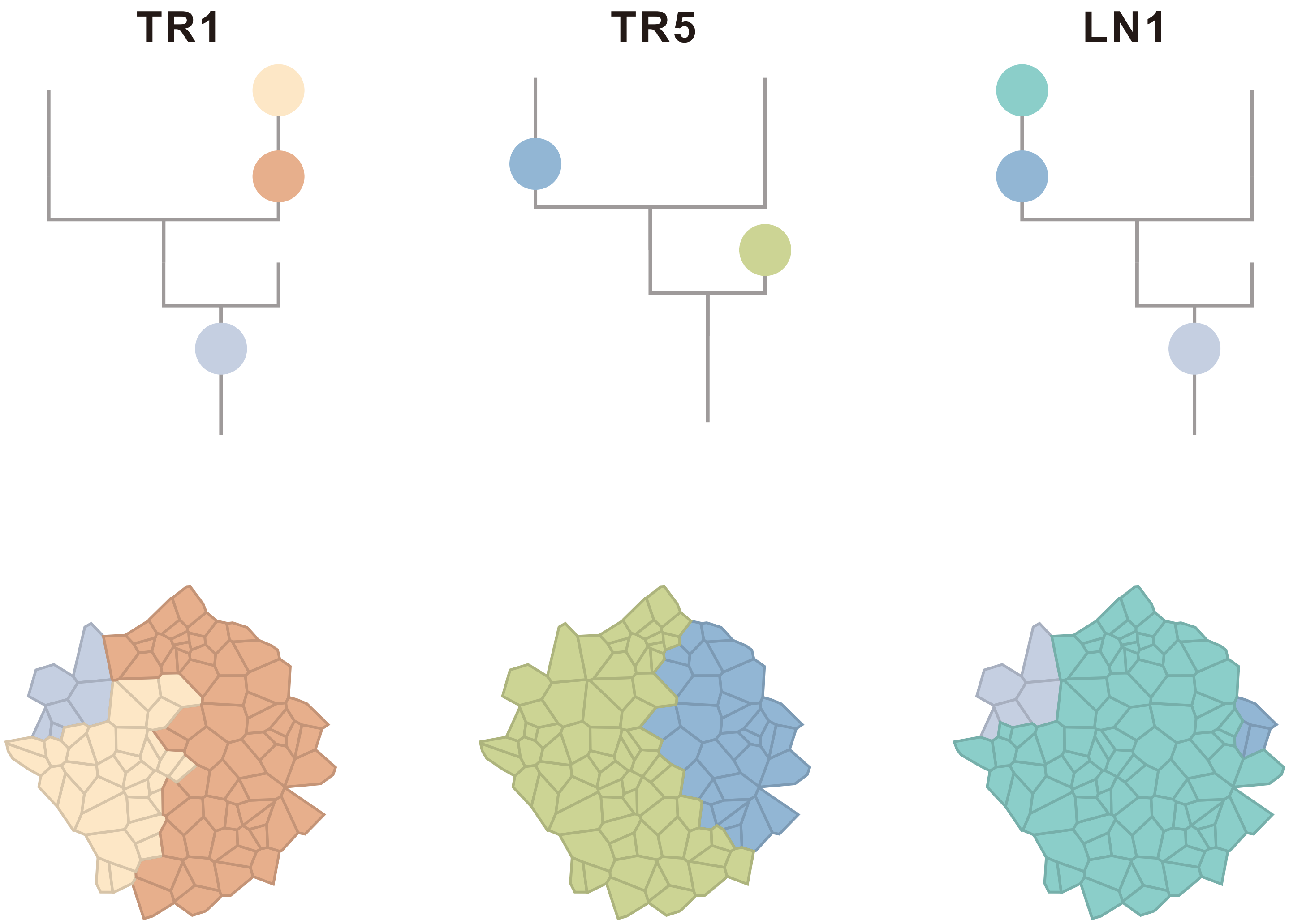

### CRC63

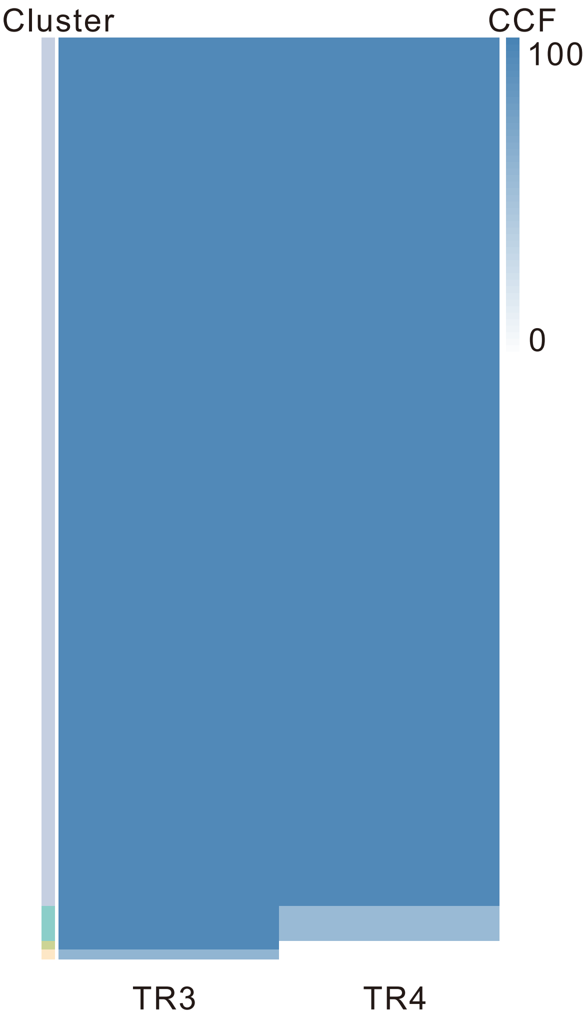

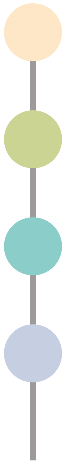

| Gene | Cytoband | Type | Cluster |
|---|---|---|---|
| *ARNT* | 1q21.3 | SNV | 1 |
| *ASXL2* | 2p23.3 | SNV | 1 |
| *FBXW7* | 4q31.3 | SNV | 1 |
| * *APC* | 5q22.2 | SNV | 1 |
| *TSC1* | 9q34.13 | SNV | 1 |
| * *TSC2* | 16p13.3 | SNV | 1 |
| * *TSC2* | 16p13.3 | SNV | 1 |
| *ERBB2* | 17q12 | SNV | 1 |
| *ARHGAP26* | 5q31.3 | SNV | 2 |
| * *APC* | 5q22.2 | InDel | 2 |
| *TP53* | 17p13.1 | SNV | NA |

## TR3

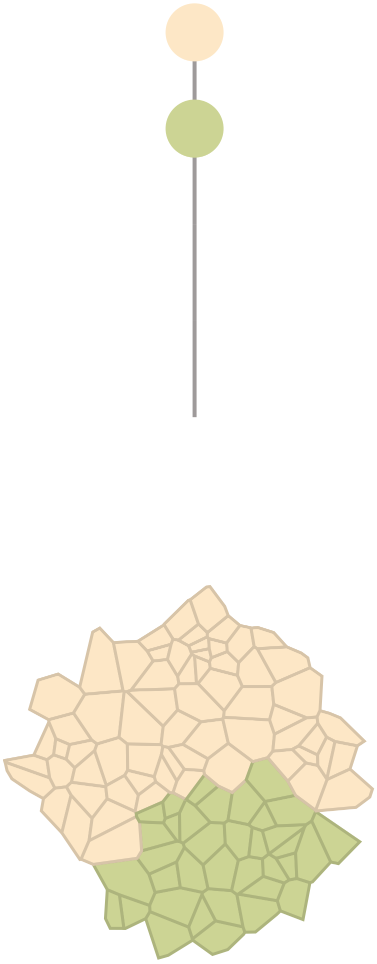

## TR4

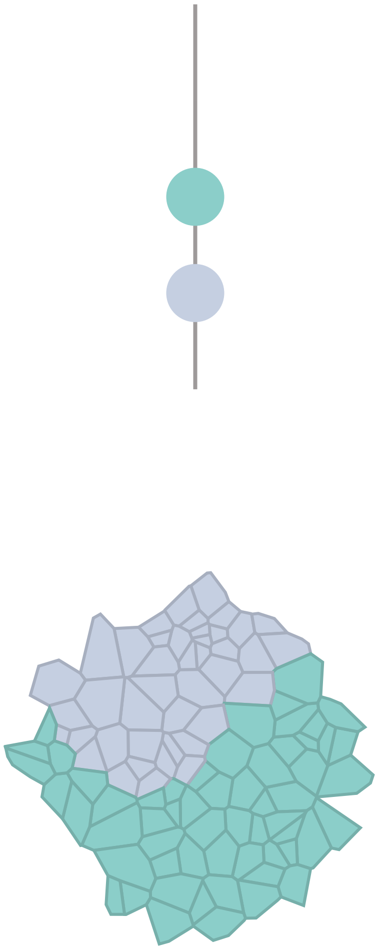

### CRC64

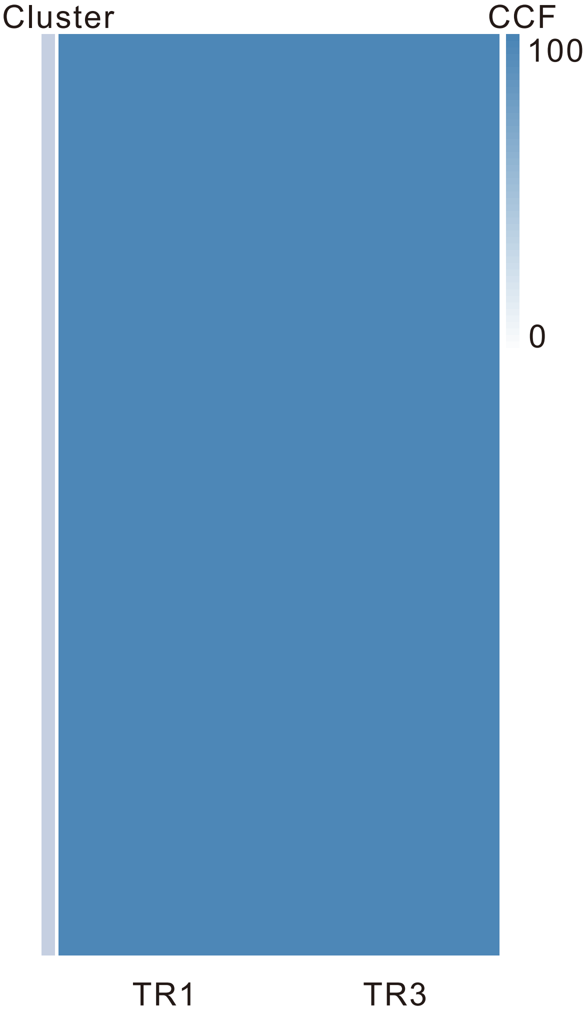

| Gene | Cytoband | Type | Cluster |
| --- | --- | --- | --- |
| *PIK3CA* | 3q26.32 | SNV | NA |
| *FBXW7* | 4q31.3 | SNV | NA |
| *KRAS* | 12p12.1 | SNV | NA |
| *SMAD2* | 18q21.1 | SNV | NA |
| *APC* | 5q22.2 | InDel | NA |
| *CBFA2T3* | 16q24.3 | Del | NA |

## TR1

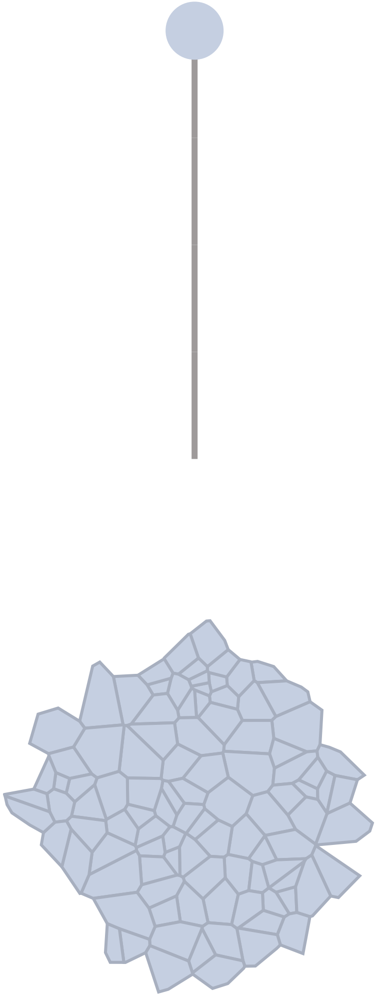

## TR3

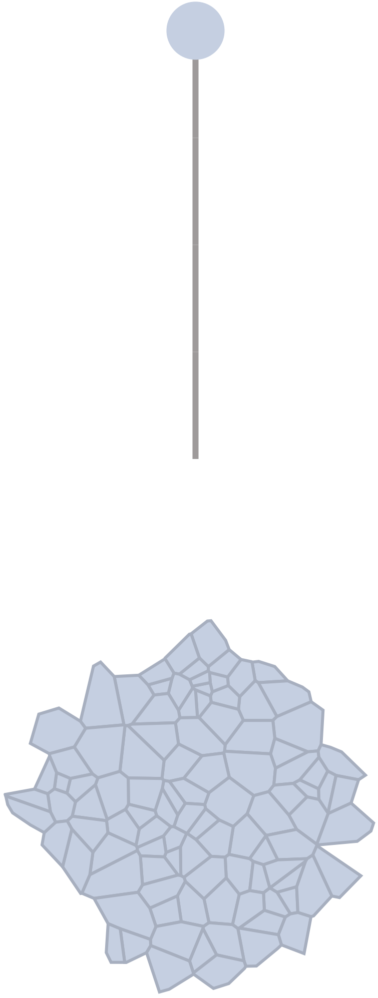

### CRC65

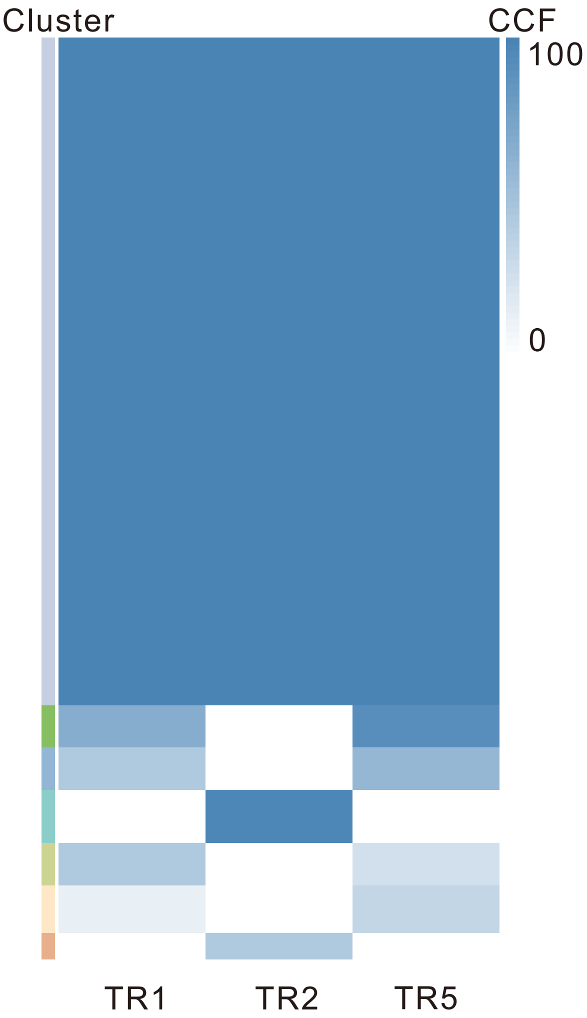

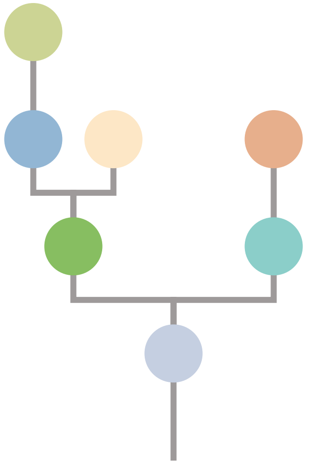

| Gene | Cytoband | Type | Cluster |
|---|---|---|---|
| *FAT4* | 4q28.1 | SNV | 1 |
| *CSMD3* | 8q23.3 | SNV | 1 |
| *KRAS* | 12p12.1 | SNV | 1 |
| *TP53* | 17p13.1 | SNV | 1 |
| *APC* | 5q22.2 | InDel | 1 |
| *SRC* | 20q11.23 | Amp | 1 |
| *MAFB* | 20q12 | Amp | 1 |
| *PLCG1* | 20q12 | Amp | 1 |
| *NFATC2* | 20q13.2 | Amp | 1 |
| *SALL4* | 20q13.2 | Amp | 1 |
| *PTK6* | 20q13.33 | Amp | 1 |
| *SIRPA* | 20p13 | Del | 1 |
| *CLTCL1* | 22q11.21 | Del | 1 |
| *LZTR1* | 22q11.21 | Del | 1 |
| *SMARCB1* | 22q11.23 | Del | 1 |
| *CHEK2* | 22q12.1 | Del | 1 |
| *ZNRF3* | 22q12.1 | Del | 1 |
| *NF2* | 22q12.2 | Del | 1 |
| *MYH9* | 22q12.3 | Del | 1 |
| *APOBEC3B* | 22q13.1 | Del | 1 |
| *EP300* | 22q13.2 | Del | 1 |
| *MKL1* | 22q13.2 | Del | 1 |
| *TBX3* | 12q24.21 | InDel | NA |
| *LZTR1* | 22q11.21 | InDel | NA |
| *GNAS* | 20q13.32 | Amp | NA |

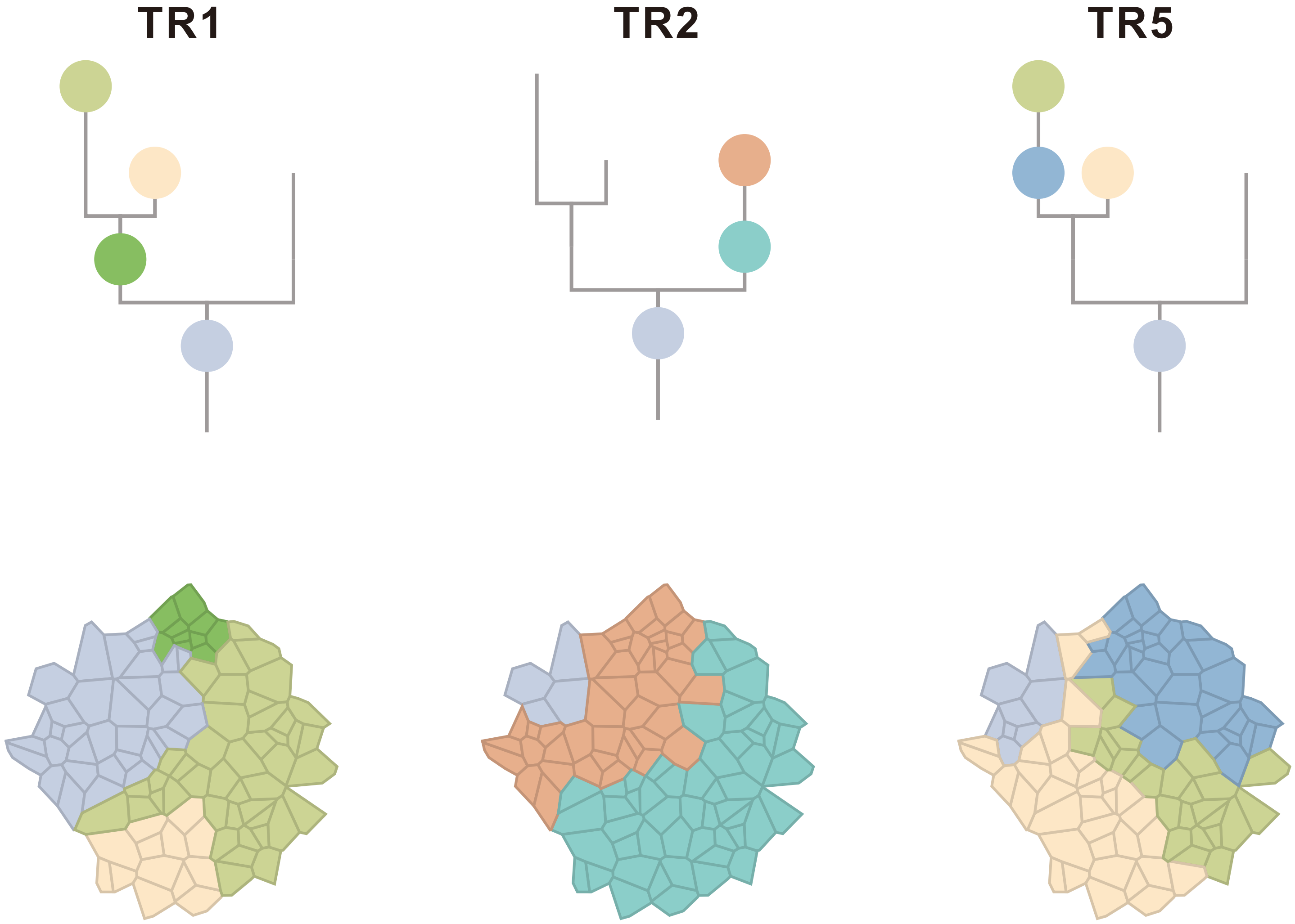

### CRC66

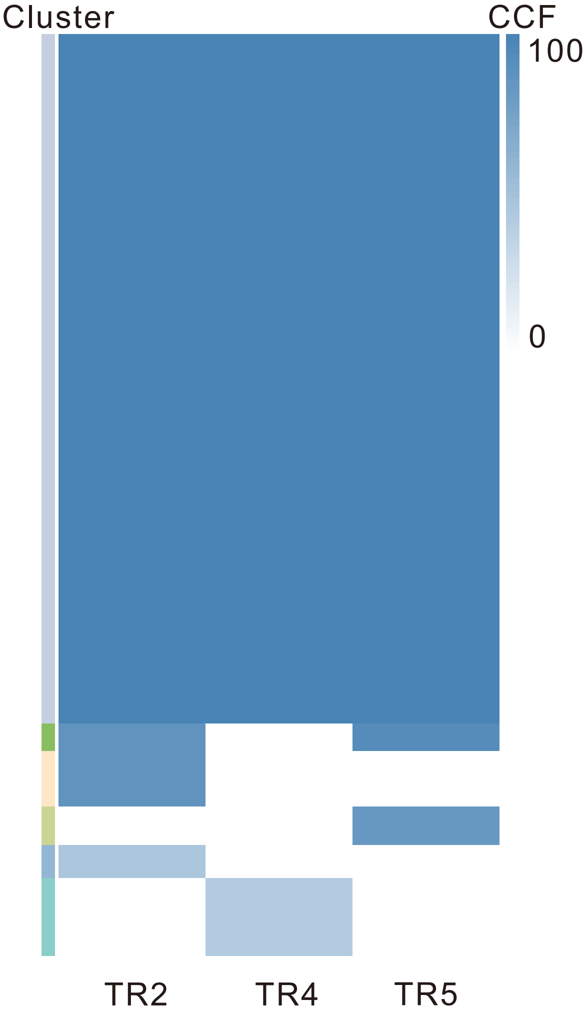

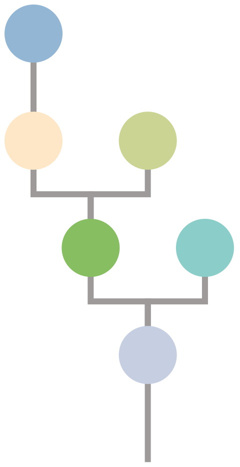

| Gene | Cytoband | Type | Cluster |
|---|---|---|---|
| * *FBXW7* | 4q31.3 | SNV | 1 |
| * *FBXW7* | 4q31.3 | SNV | 1 |
| * *FBXW7* | 4q31.3 | SNV | 1 |
| *APC* | 5q22.2 | SNV | 1 |
| *CDH11* | 16q21 | SNV | 1 |
| *TP53* | 17p13.1 | SNV | 1 |
| *SRC* | 20q11.23 | Amp | 1 |
| *MAFB* | 20q12 | Amp | 1 |
| *PLCG1* | 20q12 | Amp | 1 |
| *NFATC2* | 20q13.2 | Amp | 1 |
| *SALL4* | 20q13.2 | Amp | 1 |
| *GNAS* | 20q13.32 | Amp | 1 |
| *PTK6* | 20q13.33 | Amp | 1 |

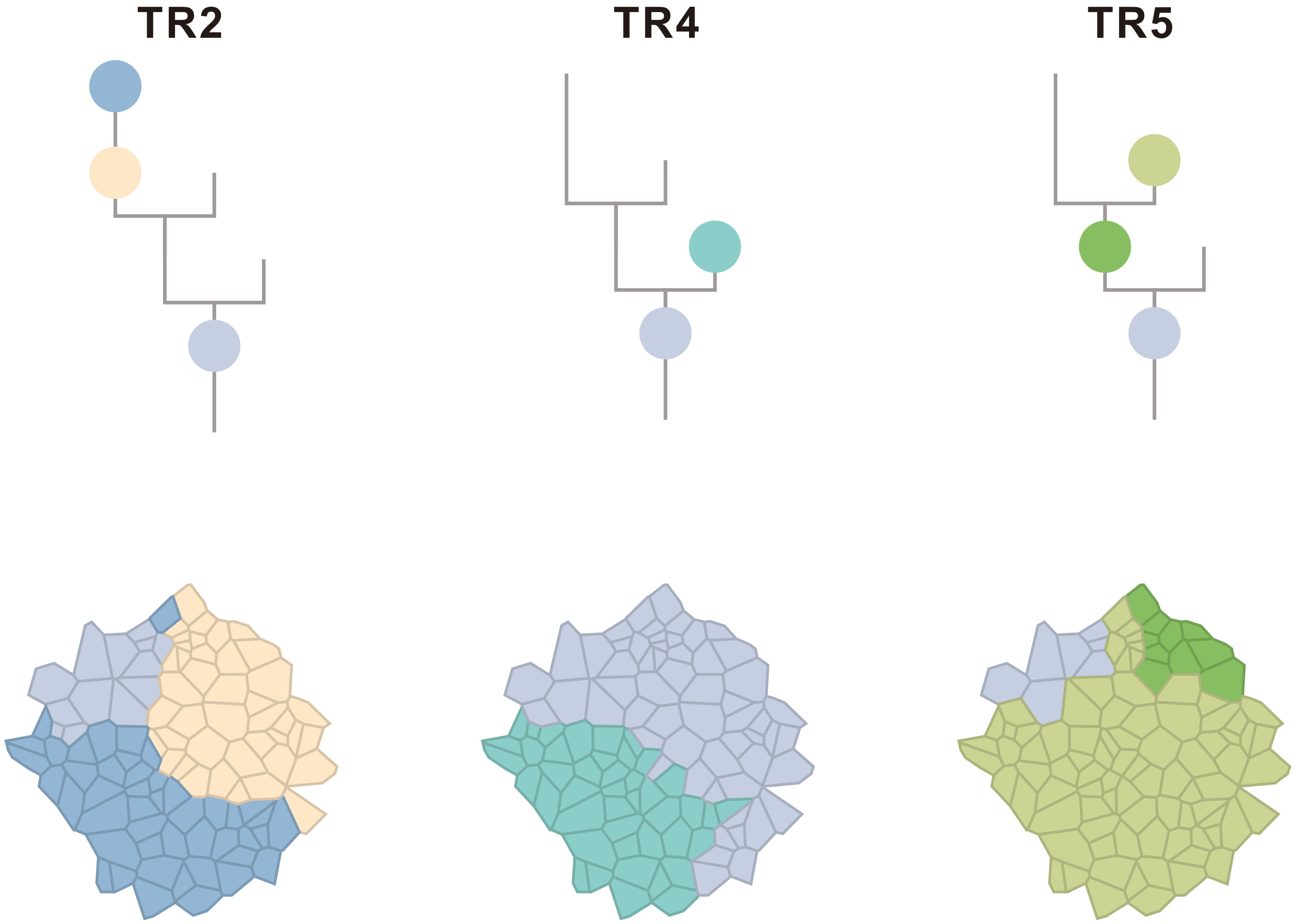

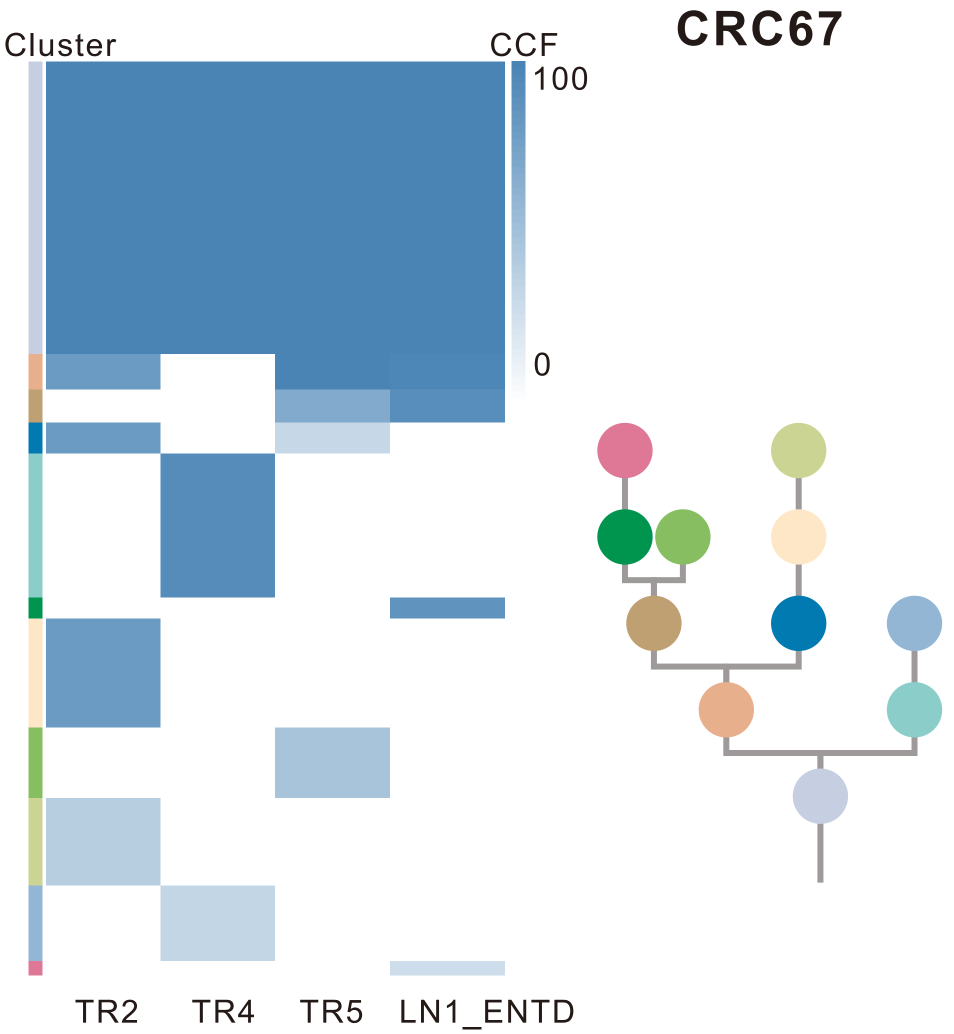

| Gene | Cytoband | Type | Cluster |
|---|---|---|---|
| * *LRP1B* | 2q22.1 | SNV | 1 |
| *PIK3CA* | 3q26.32 | SNV | 1 |
| *APC* | 5q22.2 | SNV | 1 |
| *EZH2* | 7q36.1 | SNV | 1 |
| *TP53* | 17p13.1 | SNV | 1 |
| *SMAD4* | 18q21.2 | SNV | 1 |
| *ARHGAP26* | 5q31.3 | SNV | 3 |
| * *LRP1B* | 2q22.1 | SNV | 6 |
| * *LRP1B* | 2q22.1 | SNV | 9 |

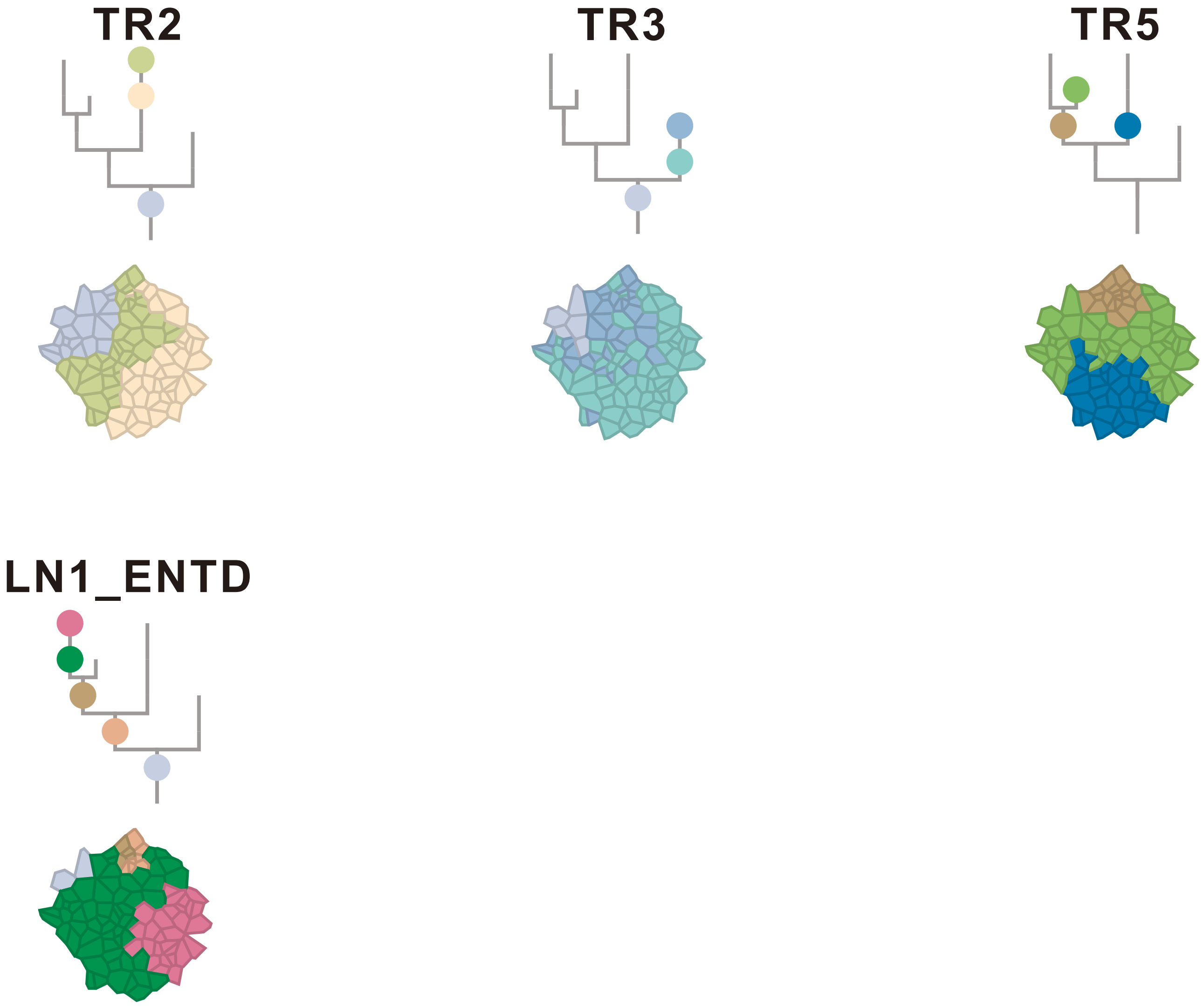

### CRC68

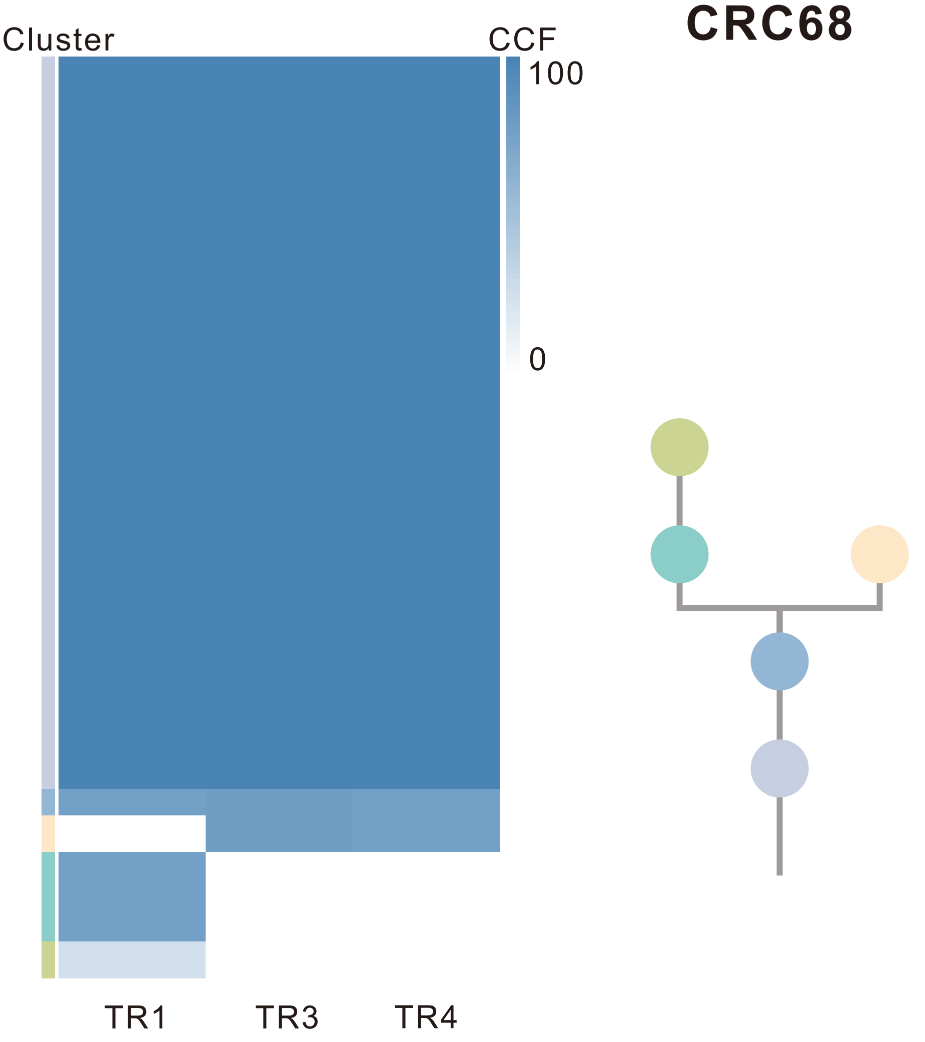

| Gene | Cytoband | Type | Cluster |
|---|---|---|---|
| *NRAS* | 1p13.2 | SNV | 1 |
| *PIK3CA* | 3q26.32 | SNV | 1 |
| * *APC* | 5q22.2 | SNV | 1 |
| *KRAS* | 12p12.1 | SNV | 1 |
| *CDH11* | 16q21 | SNV | 1 |
| *TP53* | 17p13.1 | SNV | 1 |
| * *APC* | 5q22.2 | InDel | 1 |
| *SRC* | 20q11.23 | Amp | 1 |
| *MAFB* | 20q12 | Amp | 1 |
| *PLCG1* | 20q12 | Amp | 1 |
| *NFATC2* | 20q13.2 | Amp | 1 |
| *SALL4* | 20q13.2 | Amp | 1 |
| *GNAS* | 20q13.32 | Amp | 1 |
| *PTK6* | 20q13.33 | Amp | 1 |
| *PRKAR1A* | 17q24.2 | SNV | NA |

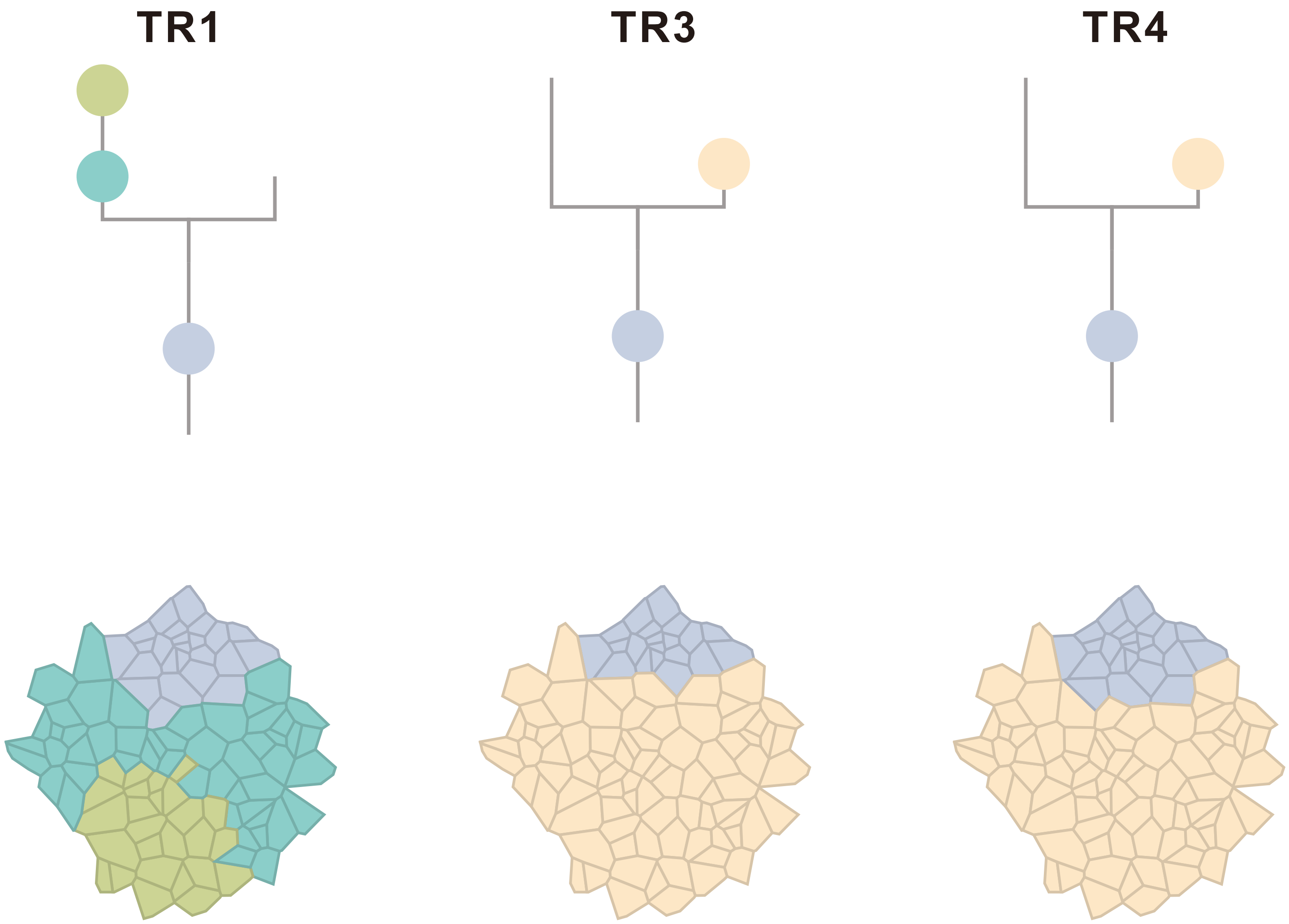

### Figure S7

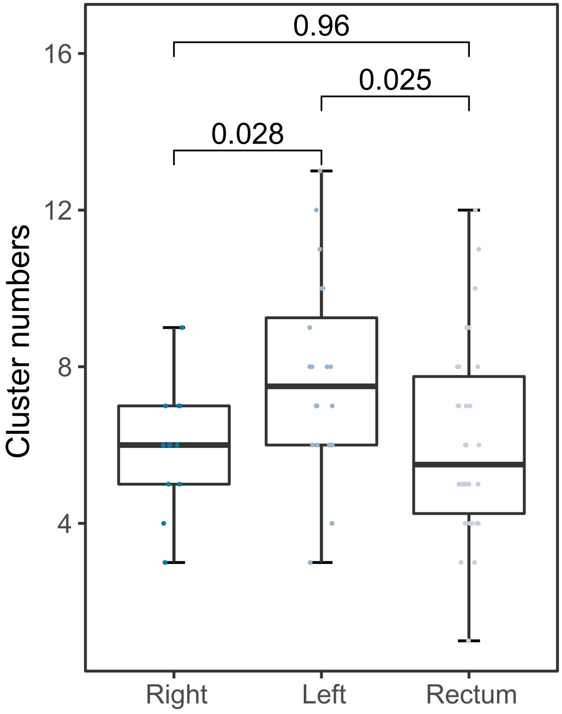

**Figure S7. Difference of cluster numbers in CRC tumors between right-sided colon, left-sided colon and rectal cancers.**
Box plots of cluster numbers in CRC tumors by position.

#### Figure S8

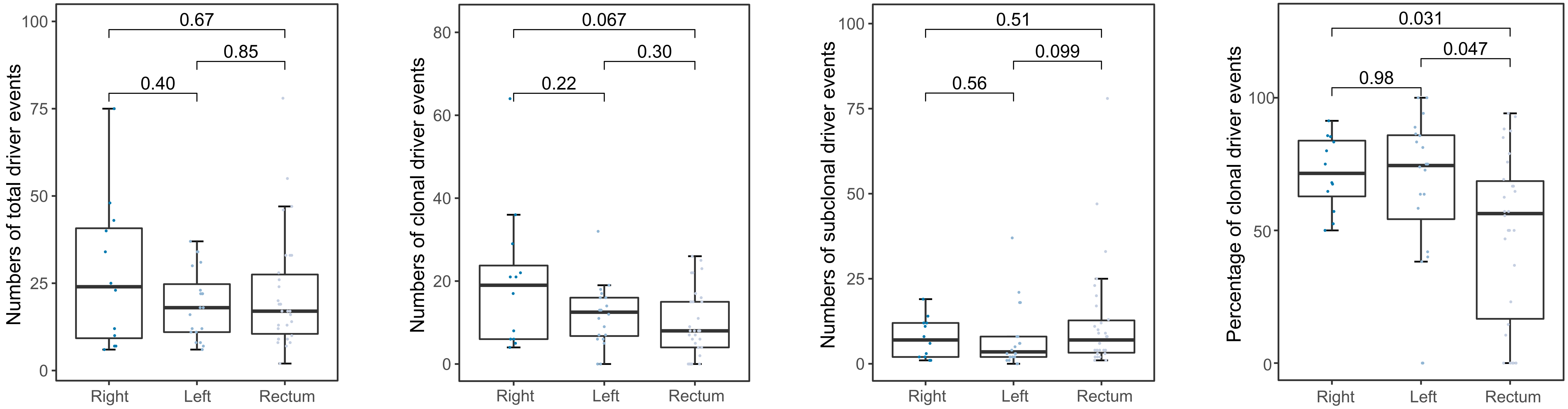

**Figure S8. Difference in numbers of driver events in CRC tumors among right-sided colon, left-sided colon and rectal cancers.** Box plots of total, clonal, subclonal and percentage of clonal numbers of driver events by tumor position.

### Figure S9

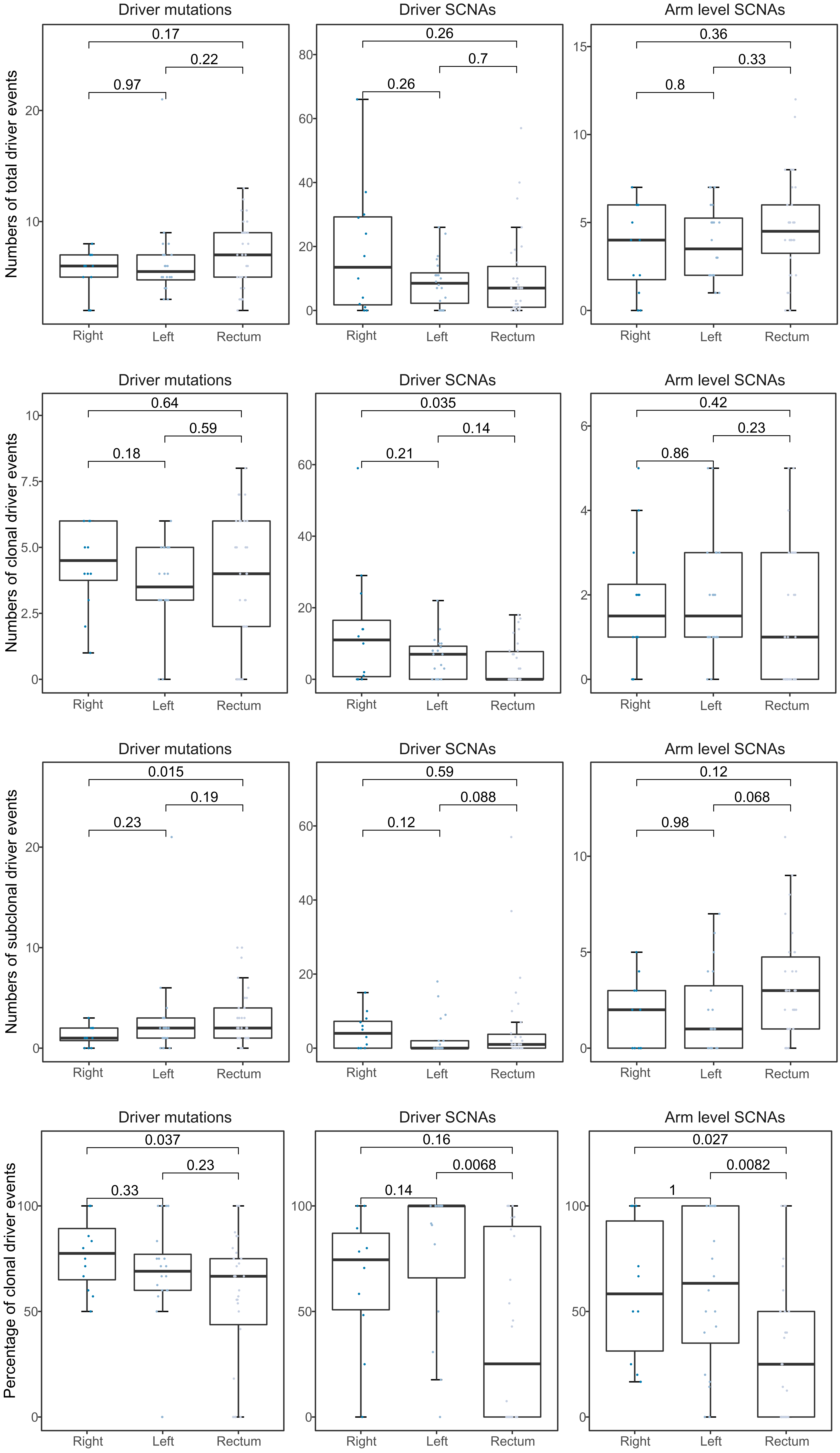

**Figure S9. Difference in types of driver events in CRC tumors among right-sided colon, left-sided colon and rectal cancers.**

Box plots of total, clonal, subclonal and percentage of clonal numbers of different driver events types by tumor position.

#### Figure S10

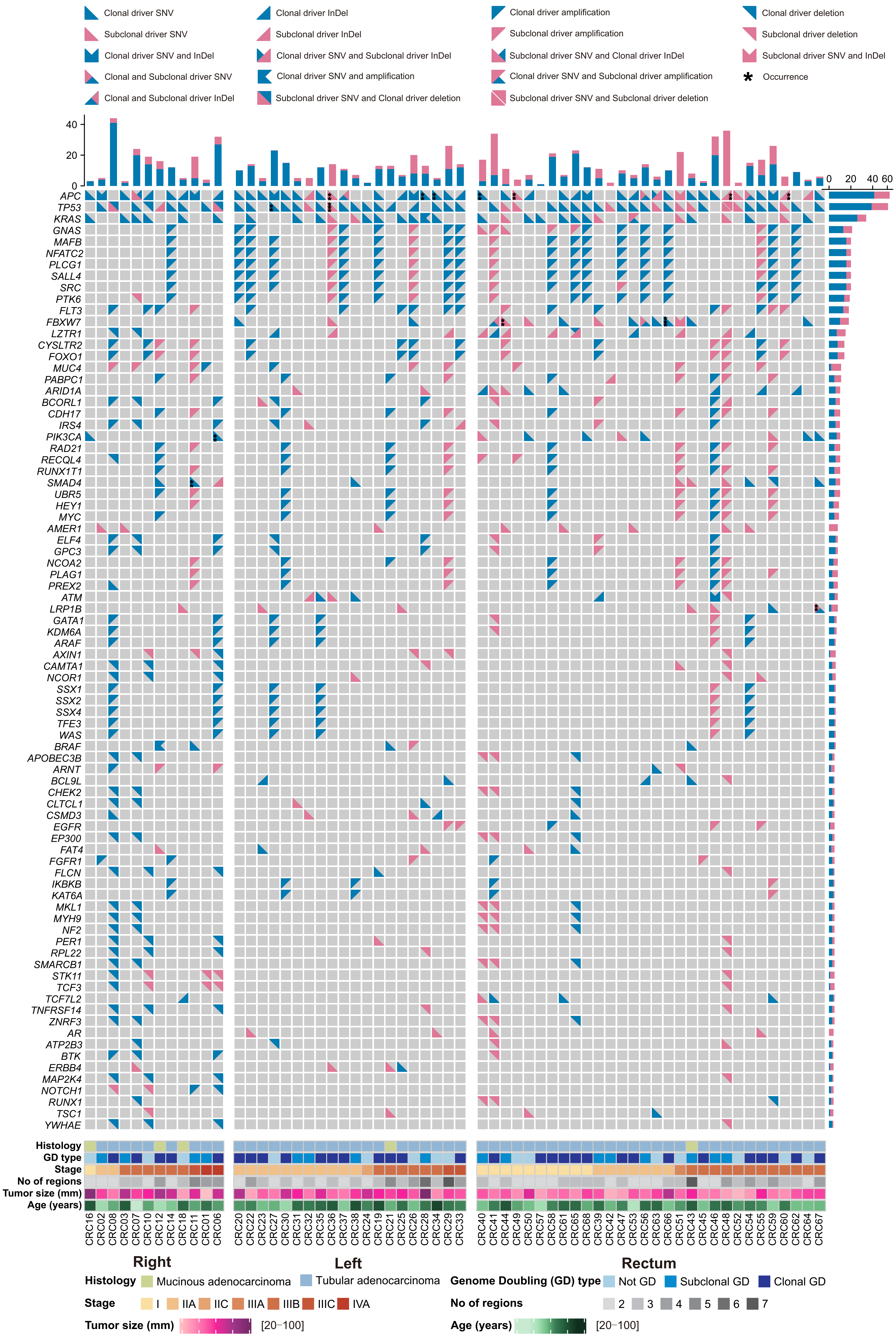

**Figure S10. Heterogeneity of driver alterations in CRC tumors.**
Driver alterations detected in each tumor. Only genes containing ≥ 4 driver alterations across the CRC patients are included. The shape and color represented the alteration type and clonal status as shown in the top panel. The second top panel displayed the number of driver alterations identified across individual CRC tumors and the bar plots to the right showed the number of the variants in right-sided colon, left-sided colon and rectal cancers for each gene. The lower panel showed the demographic and clinical characteristics of the 62 CRC patients in this study (divided by histology; whole genome doubling status; stage; number of regions; tumor size; age and tumor location).

### Figure S11

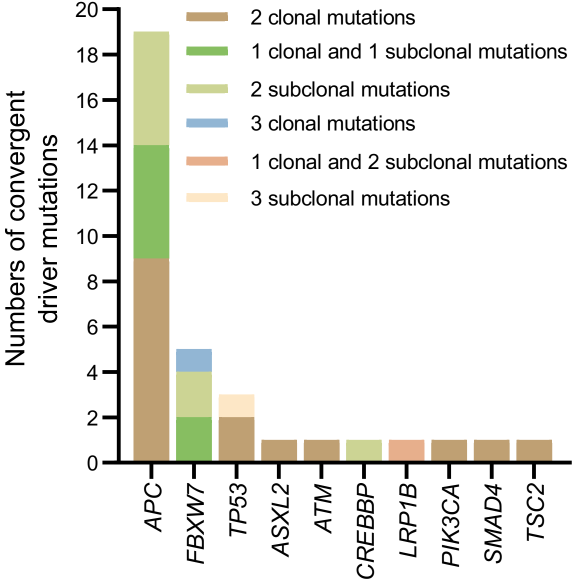

**Figure S11. Convergent driver mutations in CRC tumors.** All convergent driver mutation events were summarized in 62 CRC tumors.

#### Figure S12

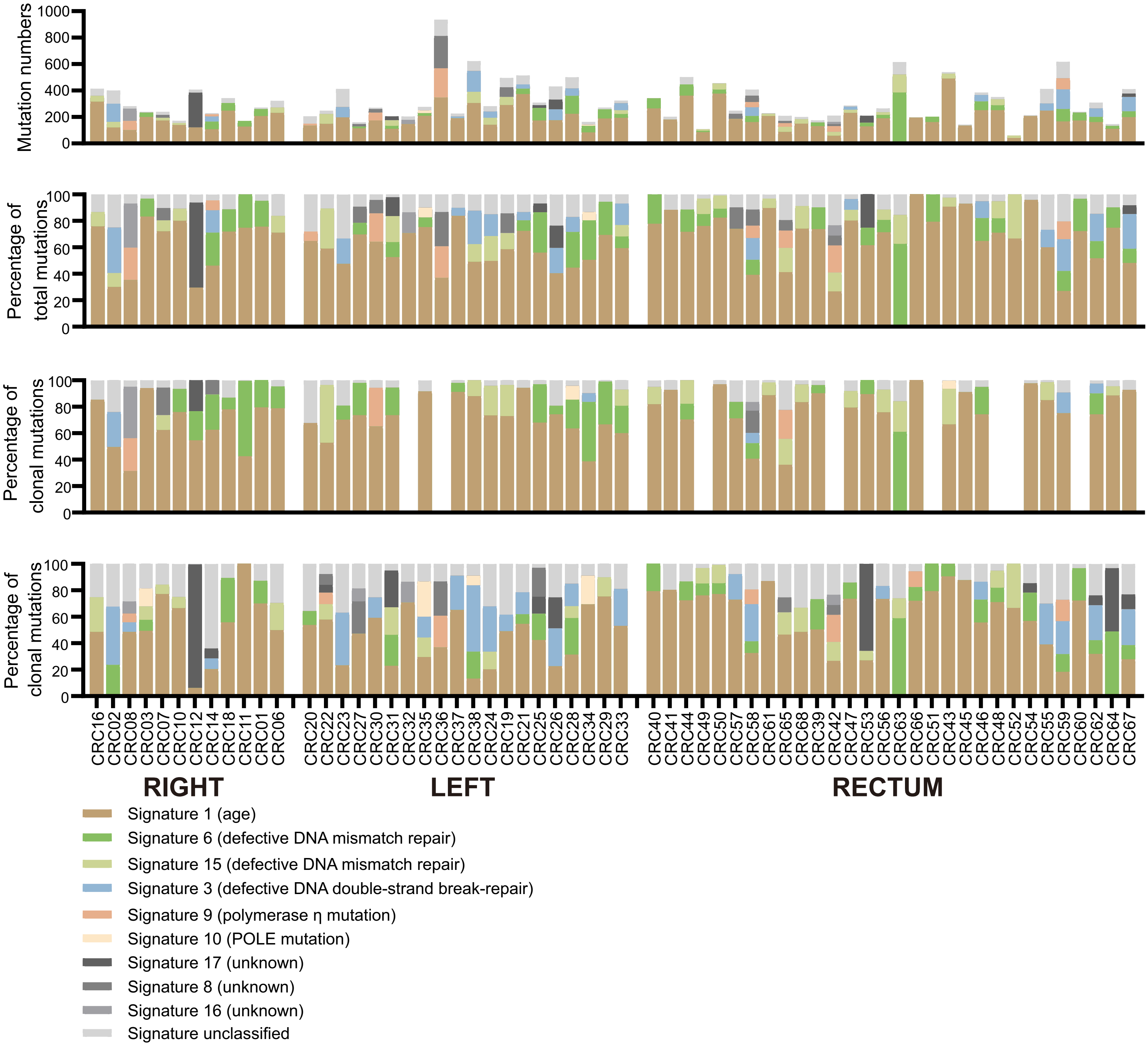

**Figure S12. Mutation signature in CRC tumors.**
Mutation signatures identified in CRC tumors, split according to tumor location and clonality of mutations.

### Figure S13

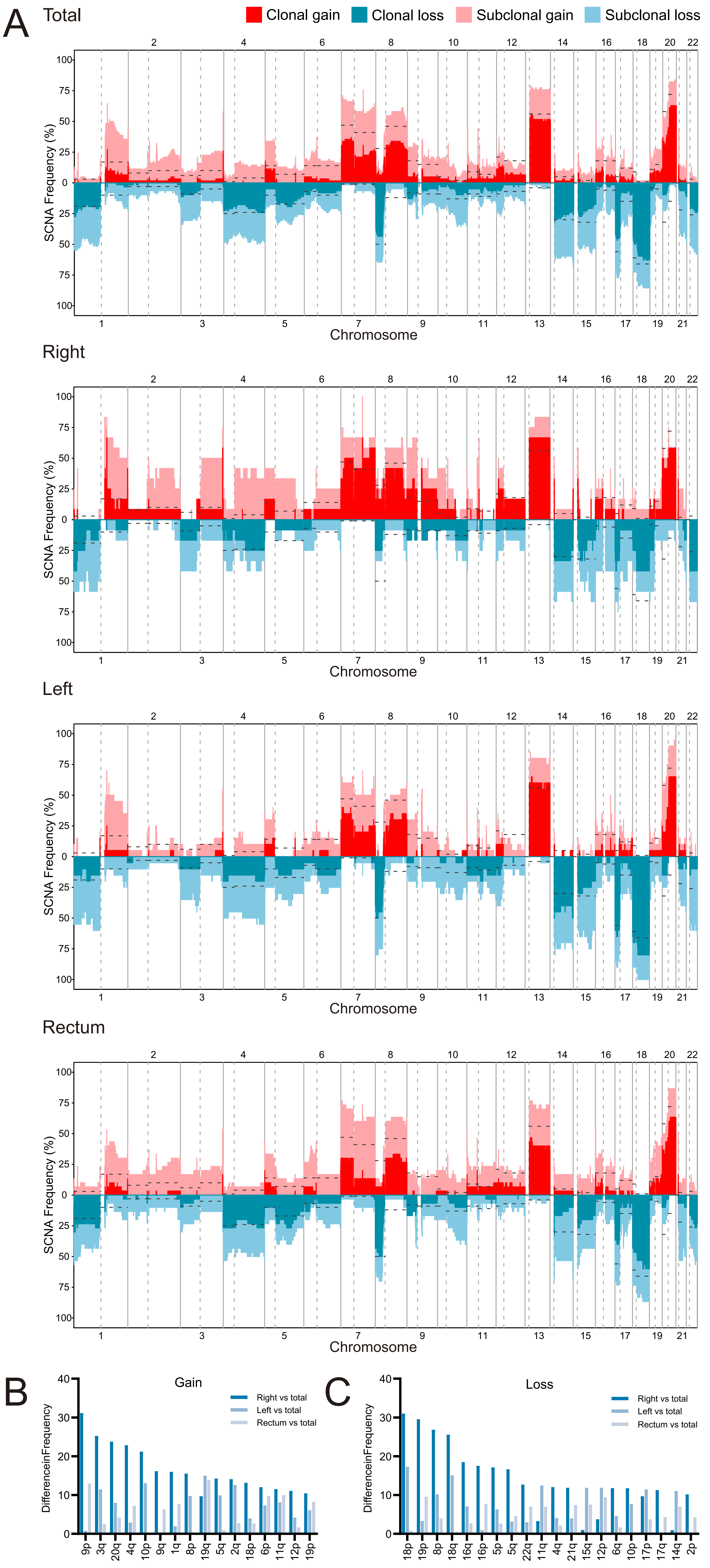

**Figure S13. Differences in SCNA frequencies among total, right-sided colon, left-sided colon and rectal cancers.**

(A) SCNA frequency of CRC tumors based on position. The dotted lines were frequency of SCNAs in TCGA CRC samples.

(B) and (C) The mean frequency of each chromosome arm by tumor position were calculated and only the chromosome arm with absolute difference of more than 10% were shown.

### Figure S14

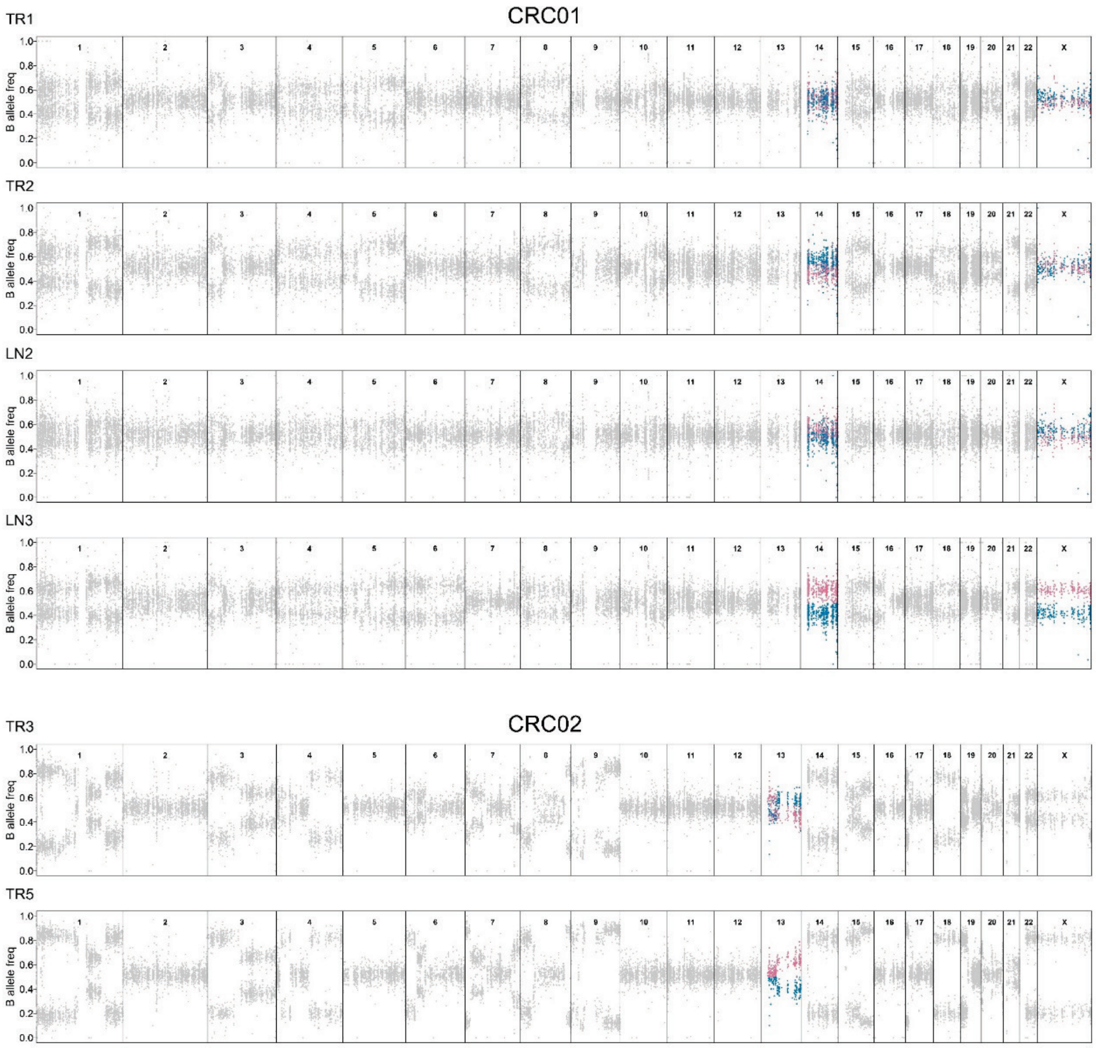

**Figure S14. Mirrored subclonal allelic imbalance (MSAI) events in CRC tumors.** B-allele frequency (BAF) profile of heterozygous SNPs across the genome (chromosomes 1-22, X) from all regions of different tumors. Sections of BAF in regions that have MSAI were highlighted in blue or red.

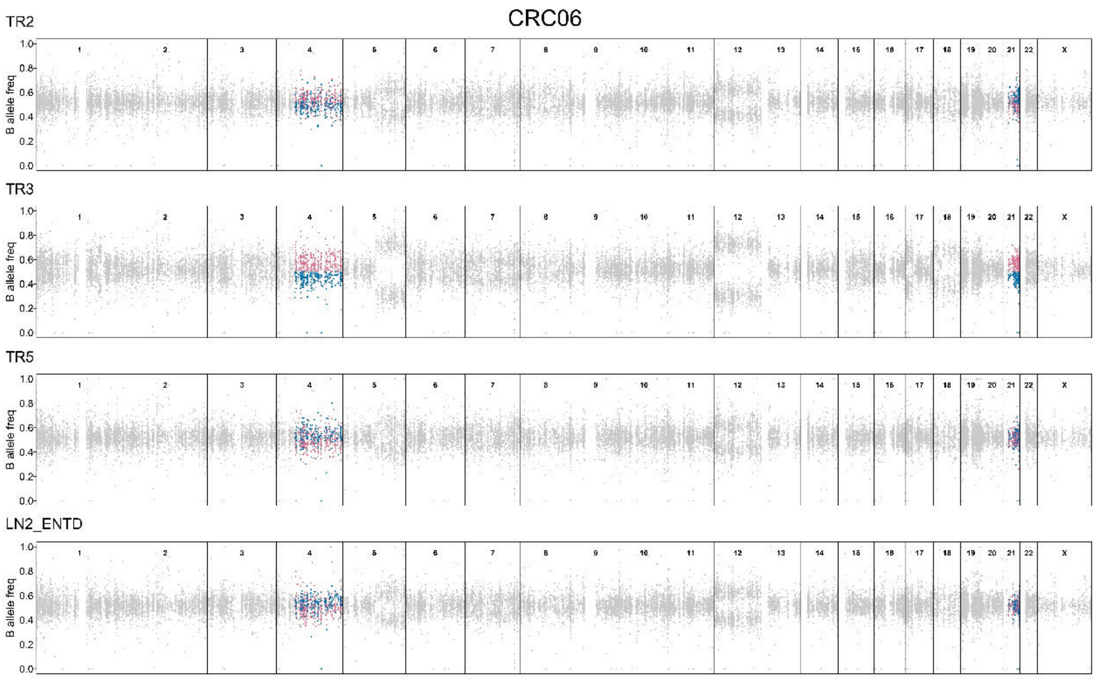
CRC06
TR2
TR3
TR5
LN2_ENTD
B allele freq
1 2 3 4 5 6 7 8 9 10 11 12 13 14 15 16 17 18 19 20 21 22 X

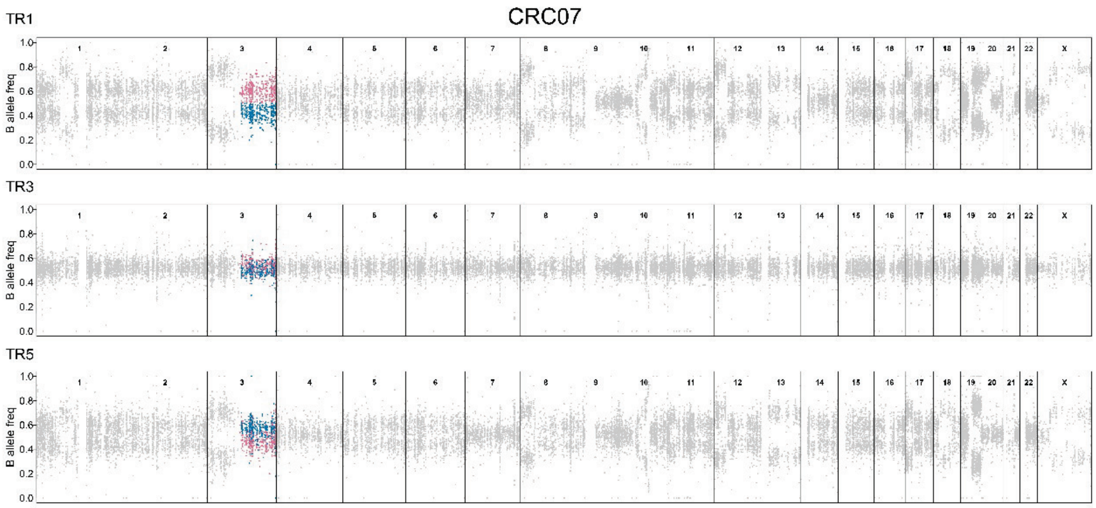
CRC07
TR1
TR3
TR5
B allele freq
1 2 3 4 5 6 7 8 9 10 11 12 13 14 15 16 17 18 19 20 21 22 X

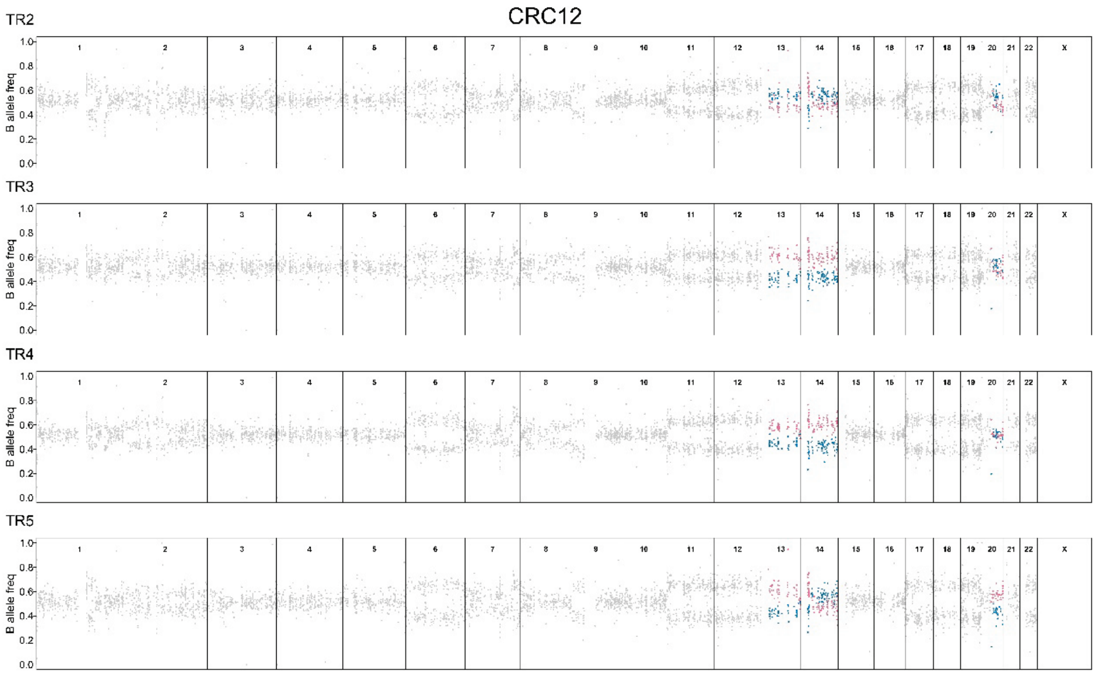
CRC12
TR2
TR3
TR4
TR5
B allele freq
1.0
0.8
0.6
0.4
0.2
0.0
1 2 3 4 5 6 7 8 9 10 11 12 13 14 15 16 17 18 19 20 21 22 X

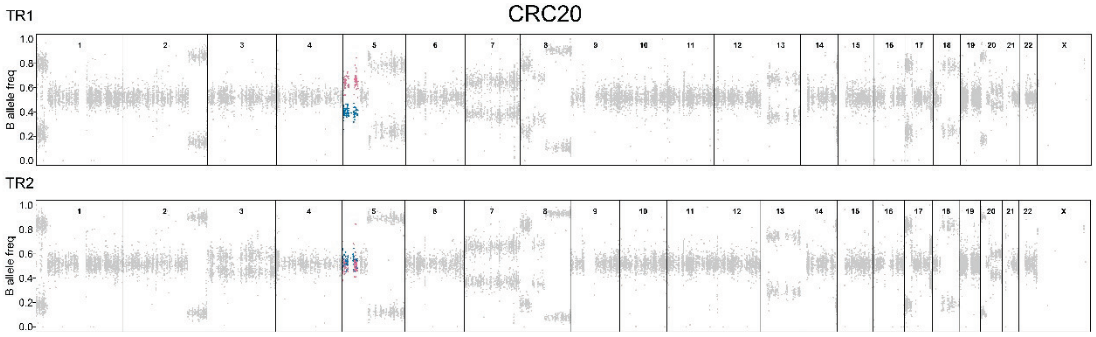
CRC20
TR1
TR2
B allele freq
1.0
0.8
0.6
0.4
0.2
0.0
1 2 3 4 5 6 7 8 9 10 11 12 13 14 15 16 17 18 19 20 21 22 X

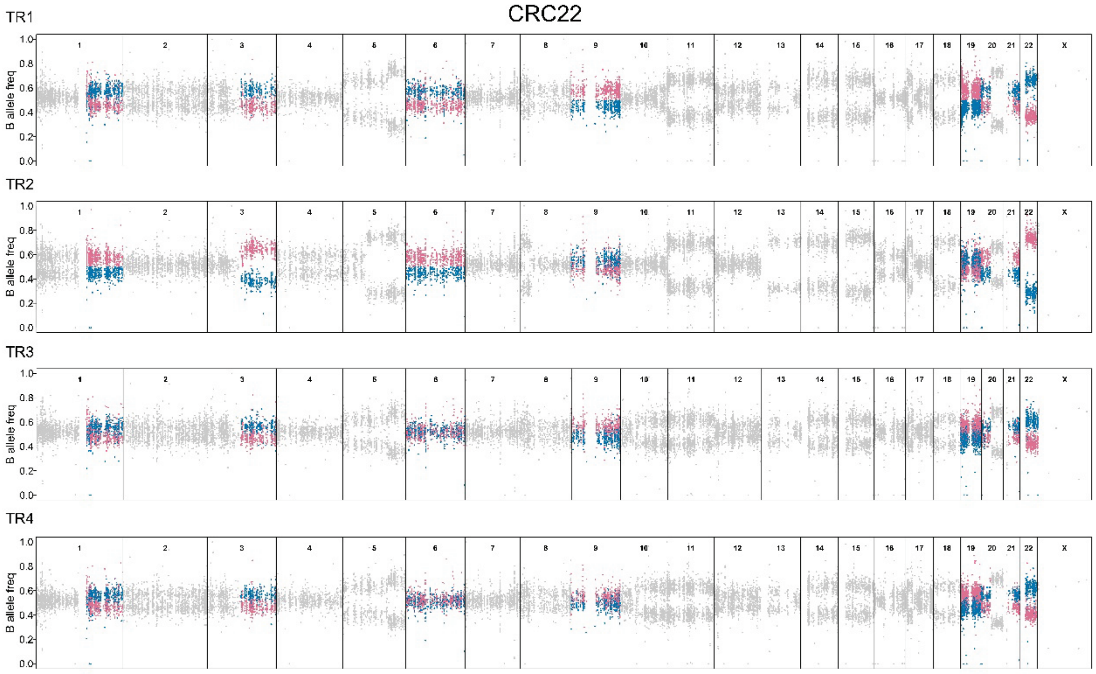
CRC22
TR1
TR2
TR3
TR4
B allele freq
1.0
0.8
0.6
0.4
0.2
0.0
1 2 3 4 5 6 7 8 9 10 11 12 13 14 15 16 17 18 19 20 21 22 X

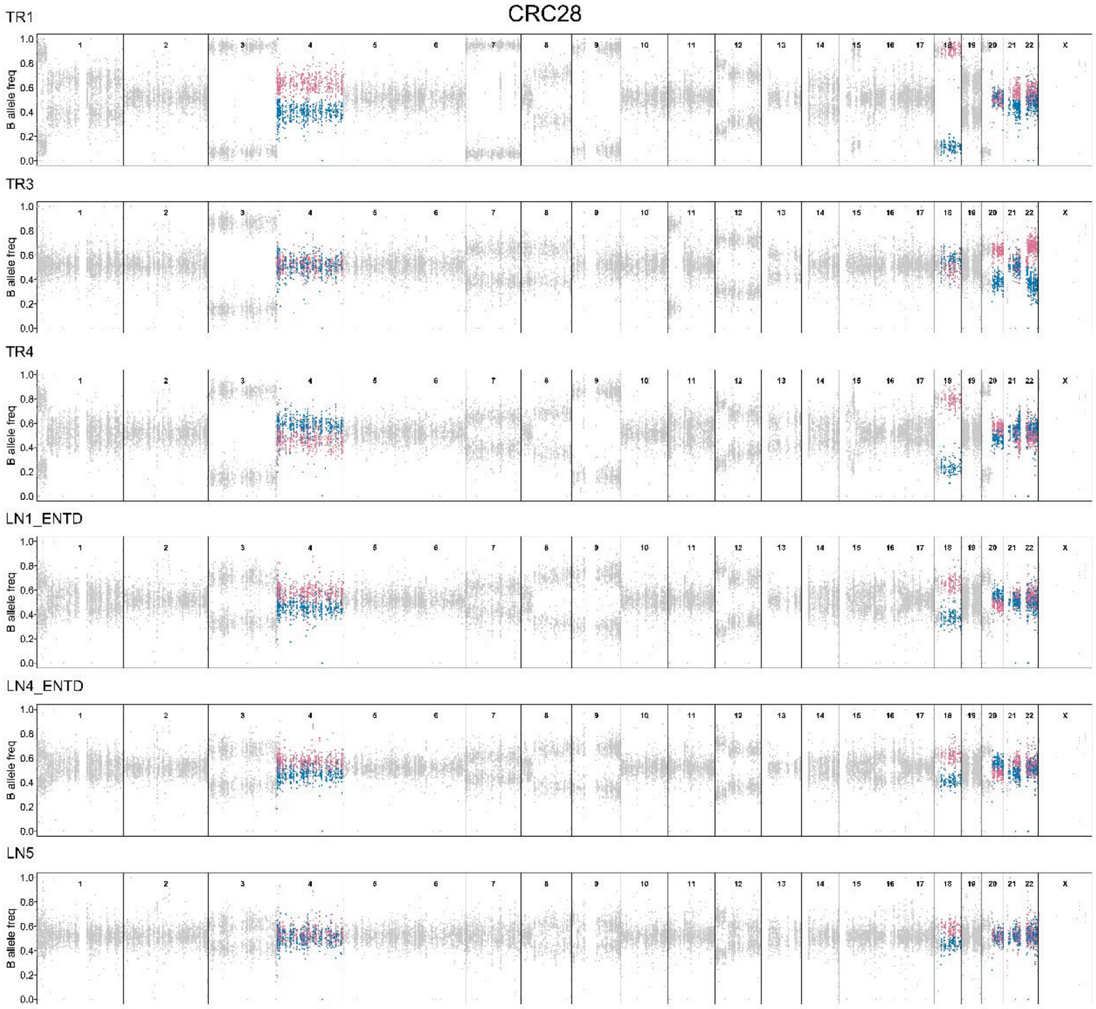
CRC28
TR1
TR3
TR4
LN1_ENTD
LN4_ENTD
LN5
B allele freq
1.0
0.8
0.6
0.4
0.2
0.0
1 2 3 4 5 6 7 8 9 10 11 12 13 14 15 16 17 18 19 20 21 22 X

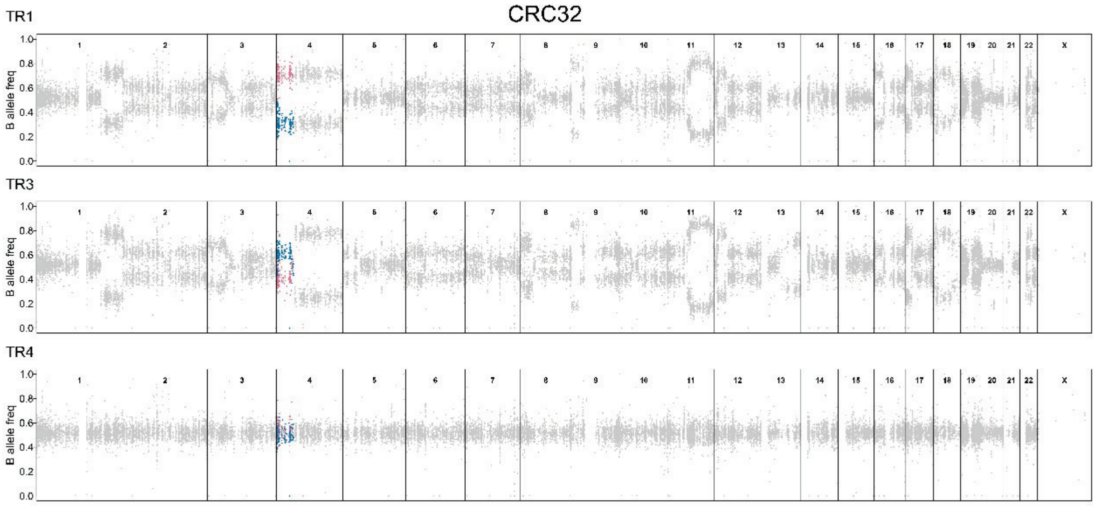
CRC32
TR1
TR3
TR4
B allele freq
1.0
0.8
0.6
0.4
0.2
0.0
1 2 3 4 5 6 7 8 9 10 11 12 13 14 15 16 17 18 19 20 21 22 X

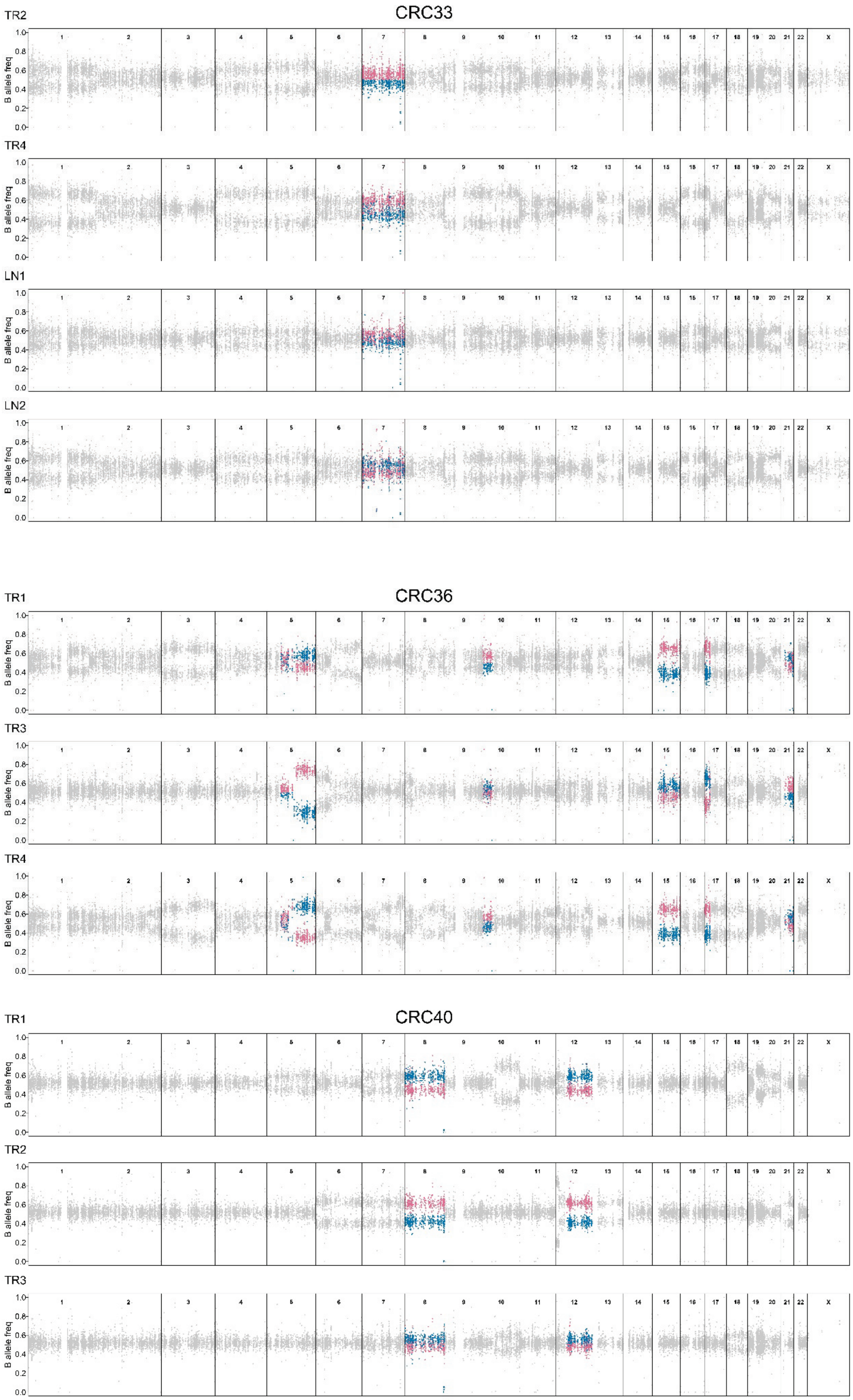
CRC33
TR2
TR4
LN1
LN2
CRC36
TR1
TR3
TR4
CRC40
TR1
TR2
TR3
B allele freq
1.0
0.8
0.6
0.4
0.2
0.0
1 2 3 4 5 6 7 8 9 10 11 12 13 14 15 16 17 18 19 20 21 22 X

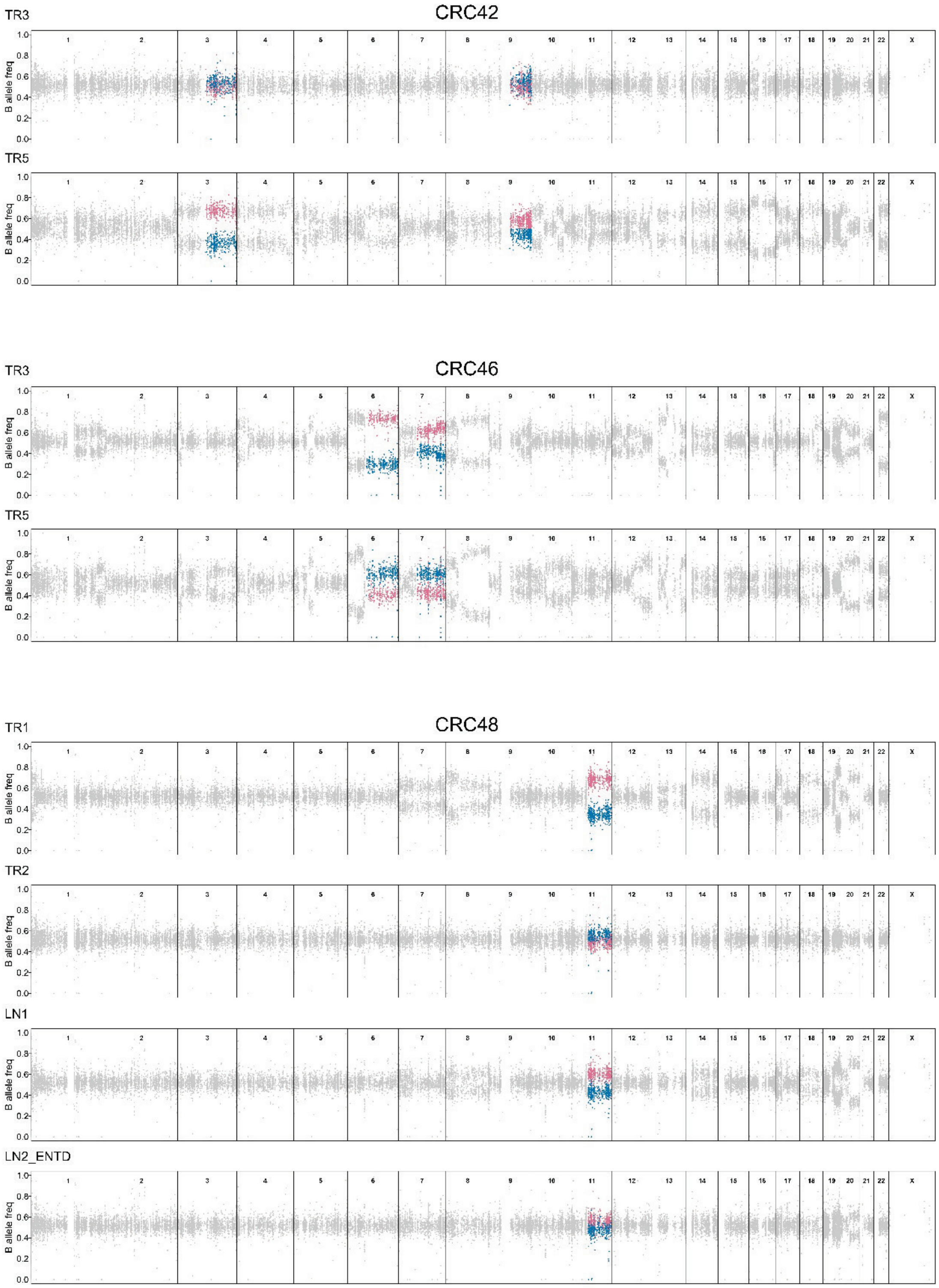
CRC42
TR3
TR5
CRC46
TR3
TR5
CRC48
TR1
TR2
LN1
LN2_ENTD
B allele freq
X
0.0
0.2
0.4
0.6
0.8
1.0

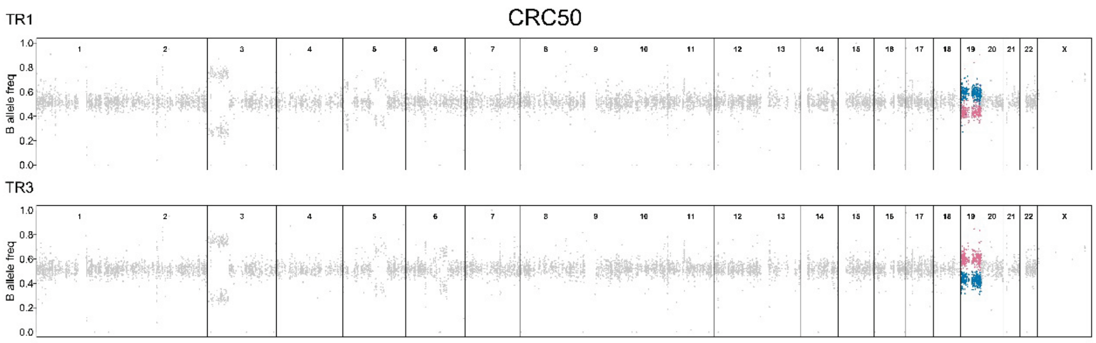
CRC50
TR1
TR3
B allele freq
1 2 3 4 5 6 7 8 9 10 11 12 13 14 15 16 17 18 19 20 21 22 X

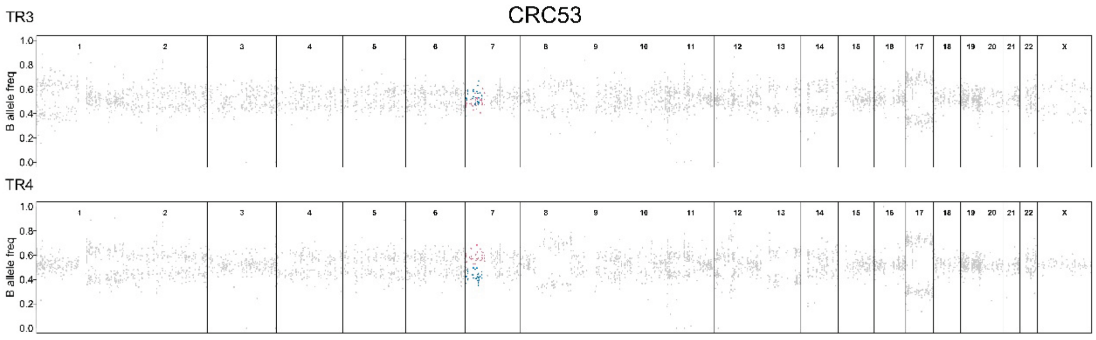
CRC53
TR3
TR4
B allele freq
1 2 3 4 5 6 7 8 9 10 11 12 13 14 15 16 17 18 19 20 21 22 X

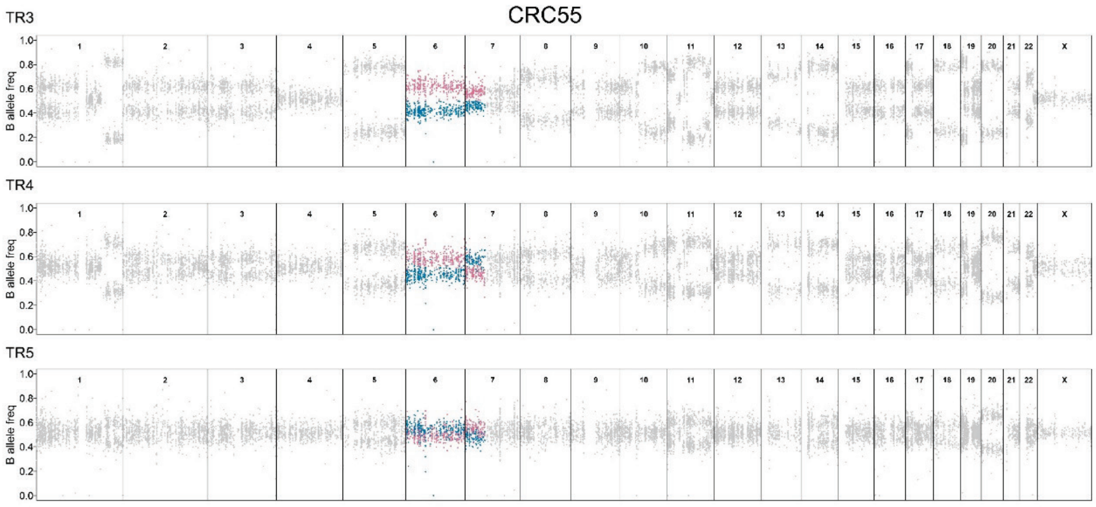
CRC55
TR3
TR4
TR5
B allele freq
1 2 3 4 5 6 7 8 9 10 11 12 13 14 15 16 17 18 19 20 21 22 X

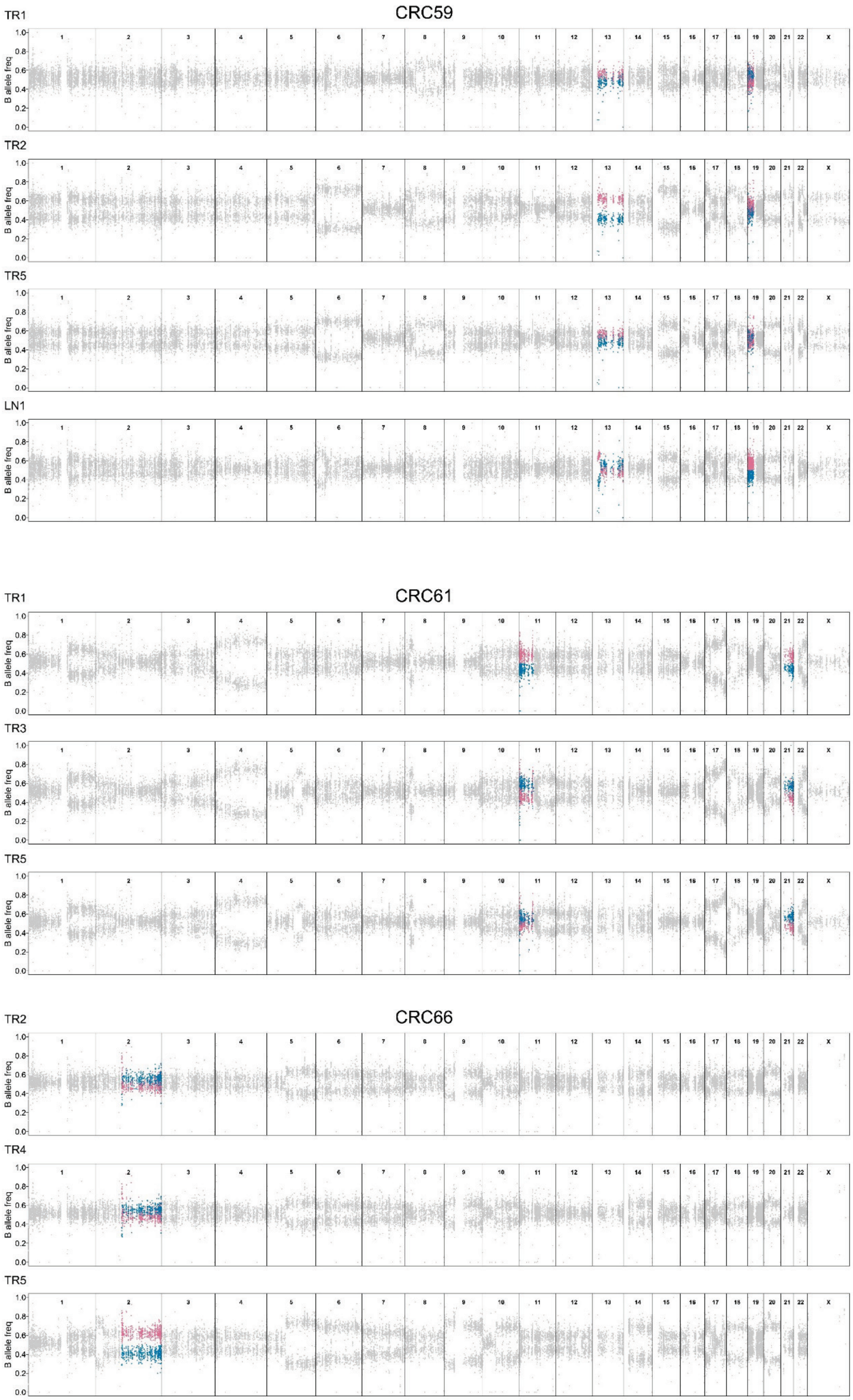
CRC59
TR1
TR2
TR5
LN1
CRC61
TR1
TR3
TR5
CRC66
TR2
TR4
TR5
B allele freq
0.0
0.2
0.4
0.6
0.8
1.0
1 2 3 4 5 6 7 8 9 10 11 12 13 14 15 16 17 18 19 20 21 22 X

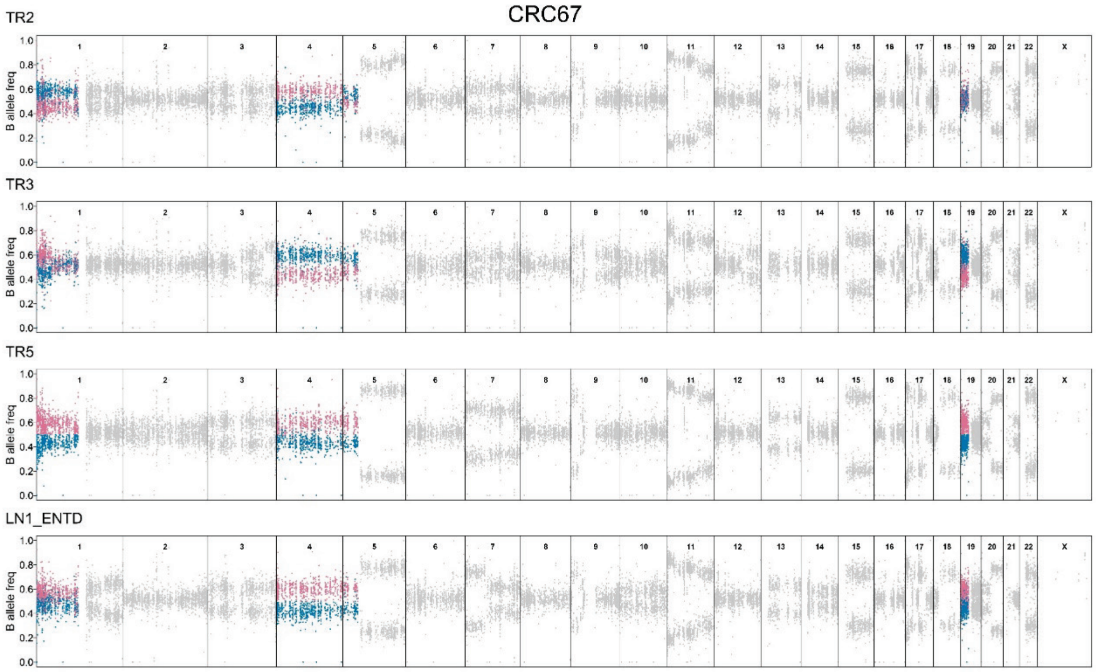
CRC67
TR2
TR3
TR5
LN1_ENTD
B allele freq
1 2 3 4 5 6 7 8 9 10 11 12 13 14 15 16 17 18 19 20 21 22 X

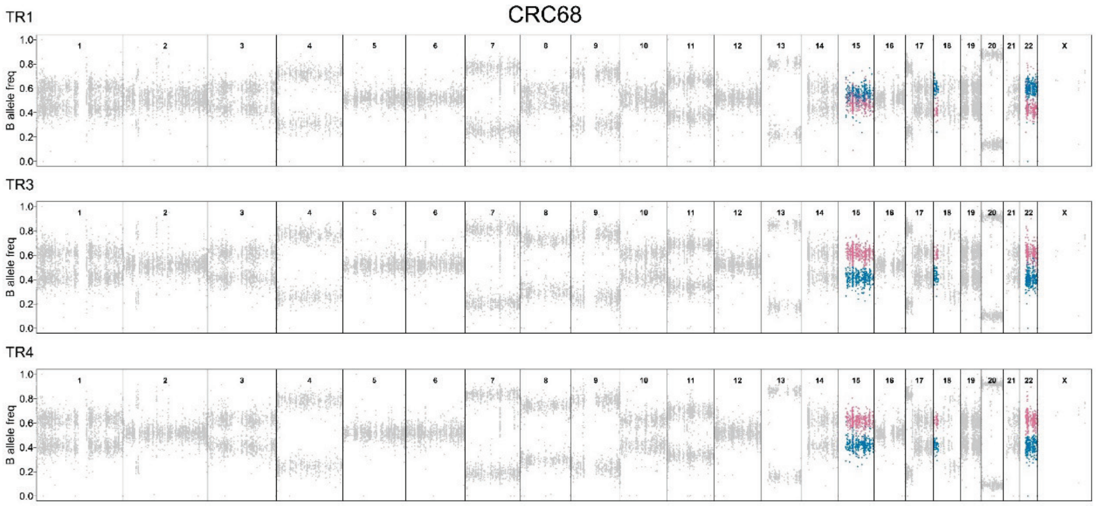
CRC68
TR1
TR3
TR4
B allele freq
1 2 3 4 5 6 7 8 9 10 11 12 13 14 15 16 17 18 19 20 21 22 X

### Figure S15

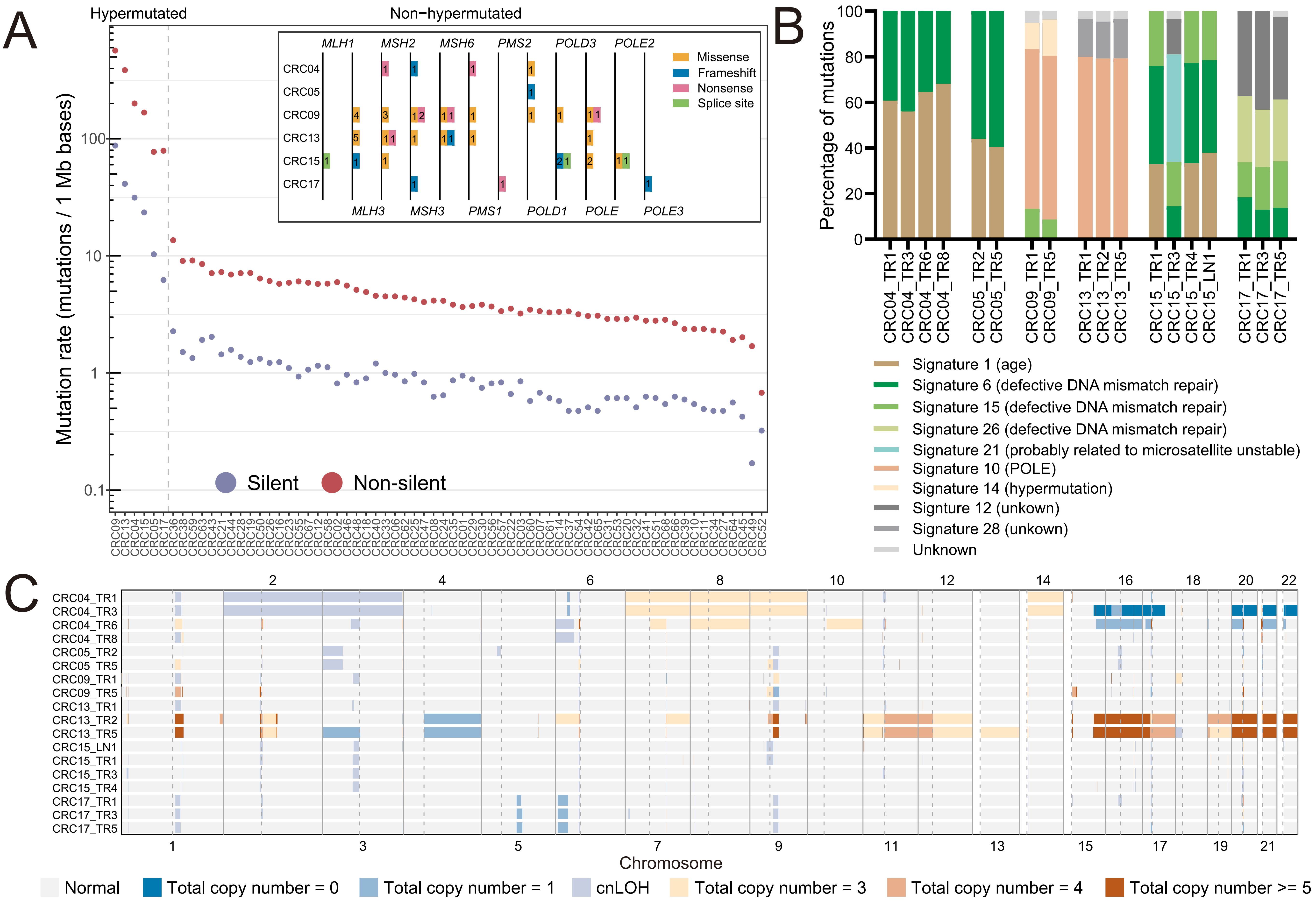

**Figure S15. Intratumor heterogeneity landscape of hypermutated CRC tumors.**

(A) Mutation rate of 68 CRC tumors. Inset, mutations in mismatch-repair genes, *POLE* and *POLD* gene family among the hypermutated tumors. The definition of hypermutated tumors was that all the sample in that tumor had more than 10 mutations/1 Mb bases.

(B) Mutation signature of all the samples from 6 hypermutated CRC tumors.

(C) Heatmap of genome-wide view SCNAs in 6 hypermutated CRC tumors.

### Figure S16

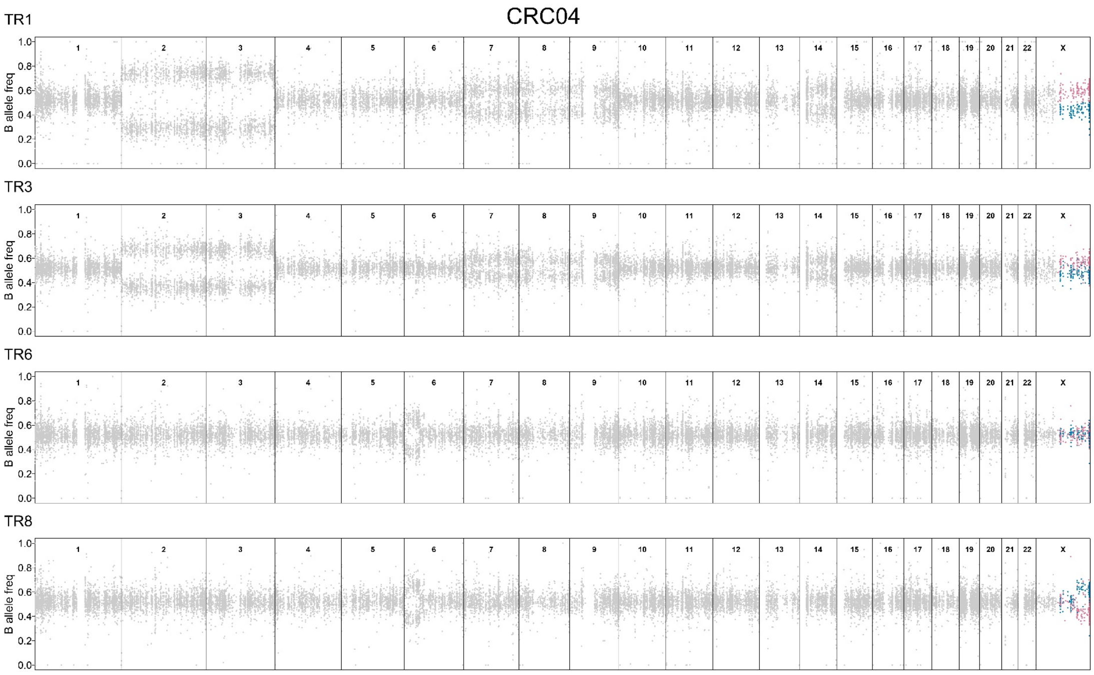

**Figure S16. MSAI events in hypermutated CRC tumor.**

B-allele frequency (BAF) profile of heterozygous SNPs across the genome (chromosomes 1-22, X) from all the regions of CRC04. Sections of BAF in regions that had MSAI were highlighted in blue or red.
